# Cell Fate Simulation Reveals Cancer Cell Features in the Tumor Microenvironment

**DOI:** 10.1101/508705

**Authors:** Sachiko Sato, Ann Rancourt, Masahiko S. Satoh

## Abstract

To elucidate the dynamic evolution of cancer cell characteristics within the tumor microenvironment (TME), we developed an integrative method combining single-cell tracking, cell fate simulation, and three-dimensional (3D) TME modeling. We began our investigation by analyzing the spatiotemporal behavior of individual cancer cells in cultured pancreatic and cervical cancer cell lines, with a focus on the α2-6 sialic acid (2-6Sia) modification on glycans, which is associated with cell stemness. Our findings revealed that pancreatic cancer cells exhibited significantly higher levels of 2-6Sia modification, correlating with enhanced reproductive capabilities, whereas cervical cancer cells showed less prevalence of this modification. To accommodate the *in vivo* variability of 2-6Sia levels, we employed a cell fate simulation algorithm that digitally generates cell populations based on our observed data, simulating cell growth patterns. Subsequently, we constructed a 3D TME model incorporating these deduced cell populations along with specific immune cell landscapes derived from 193 cervical and 172 pancreatic cancer cases. Our analysis suggests that pancreatic cancer cells are less influenced by the immune cell landscape within the TME compared to cervical cancer cells, highlighting that the fate of cancer cells is shaped by both the surrounding immune landscape and the intrinsic characteristics of the cancer cells.

## Introduction

Cancer cell populations exhibit a range of phenotypical characteristics stemming from genetic sequence alterations and chromosomal variations (*1–8*). Notably, missense mutations in p53 are prevalent in over 50% of human cancer cases (*9–15*). Aneuploidy, characterized by an abnormal number of chromosomes, arises through mechanisms such as DNA damage, impaired mitotic phase checkpoints (*16, 17*), and multipolar cell division, which is frequently observed in cancer tissues (*18–20*). Moreover, cancer cells often retain stemness-like properties to varying degrees within a cancer cell population (*21–24*), contributing to the phenotypical diversity that accumulates during cancer development.

These heterogeneous cancer cell populations coexist within the tumor microenvironment (TME), comprising various cell types such as infiltrated immune cells, tumor-associated fibroblasts, endothelial cells, and the cancer cells themselves (*25–28*). This ecosystem is characterized by conditions like hypoxia and abnormal levels of cytokines, growth factors, and metabolites (*25–28*). Insights into the cellular composition of the TME in primary tumors have been provided by The Cancer Genome Atlas Program through genomic, epigenetic, and protein-level molecular profiling (*29–33*), depicting a snapshot of the TME’s landscape at a specific stage in cancer development. However, both the landscape and the attributes of cancer cells within the TME are subject to evolution, influencing the fate of cancer.

To gain insight into the spatiotemporal dynamics of cancer cell evolution and its interplay with immune cells in the TME, we developed a three-dimensional (3D) TME simulation. This simulation positions cancer and immune cells to investigate the fate of cancer cells within the TME, considering the spatiotemporal aspects of their interactions. Our approach began with acquiring spatiotemporal data on individual cell behavior through empirical single-cell tracking of established cultured cancer cell lines (*20, 34*). We developed a computerized single-cell tracking system to generate data suitable for bioinformatics analysis, involving long-term live cell imaging, video generation, individual cell tracking, and cell lineage creation. Our focus was particularly on aspects related to stemness, notably the α2-6 sialic acid structure (2-6Sia) (*35–40*), and multipolar cell division leading to altered chromosome numbers (*18–20*). Our findings revealed that cells derived from pancreatic cancer exhibit higher 2-6Sia expression, correlating with enhanced reproductive capacity. In contrast, a smaller subset of cervical cancer-derived cells shows elevated 2-6Sia expression, highlighting distinct characteristics from pancreatic cancer cells. Subsequently, using single-cell data from cancer cell lines and considering the *in vivo* variation of 2-6Sia expression levels (*41–44*), we generated deduced cell populations digitally using a cell fate simulation algorithm (*20*). These deduced populations, embodying the attributes of cancer cells in cell lines, reflect specific 2-6Sia expression levels observed *in vivo*. These populations were then integrated into the 3D TME simulation alongside immune cells, enabling us to simulate the fate of each cancer cell within the TME. We conducted a 3D TME simulation using immune cell landscapes characteristic of 193 cervical and 172 pancreatic cancers. Our 3D TME simulation, utilizing immune cell landscape data from 193 cervical and 172 pancreatic cancer cases, indicates that the fate of pancreatic cancer cells is less influenced by the immune cell landscape within the TME, compared to cervical cancer cells. These findings suggest that the fate of cancer cells within the TME is affected not only by the immune cell landscape but also by the intrinsic characteristics of the cancer cells themselves, underscoring the importance of understanding the spatiotemporal context of interactions between immune cells and cancer cells with diverse characteristics within the TME.

## Results

### Overview of single-cell tracking and 3D TME simulation

The cancer cell population comprises cells with diverse phenotypic characteristics that evolve over time (Fig. 1A). The fate of these cells is influenced by cells residing in the TME, including immune cells. These immune cells can either suppress the growth of cancer cells or create a permissive microenvironment for the proliferation of cancer cells (*25–28*). To simulate the fate of cancer cells within the TME, we required detailed information about the spatiotemporal behavior of individual cancer cells.

**Fig. 1.**
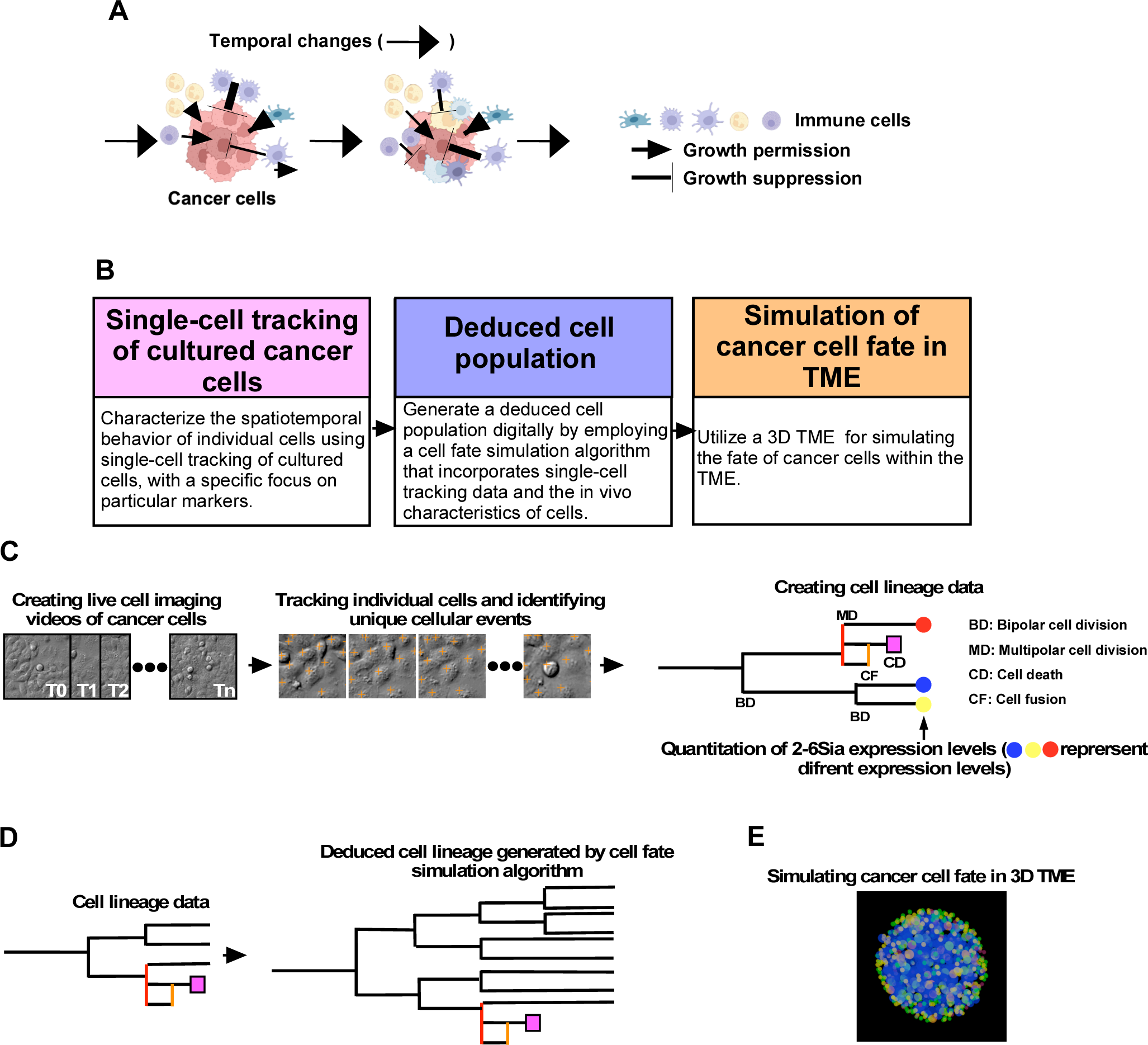
Conceptual Workflow for Performing a 3D TME Simulation. **A.** The cancer cell population within the TME exhibits inherent phenotypic diversity and dynamically evolves over time. **B.** To uncover the temporal dynamics of cancer cells within the TME, a conceptual workflow for performing a 3D TME simulation was developed. It starts with the acquisition of spatiotemporal information about individual cancer cells using a computerized single-cell tracking system. Subsequently, deduced cell populations that signify particular attributes of cells *in vivo* are generated using a cell fate simulation algorithm, followed by the execution of a 3D TME simulation. **C.** Single-cell tracking is performed using established cancer cell lines. This involves creating live cell imaging videos of cultured cancer cells and tracking individual cells. The resulting data resources include cell lineage information. Quantitative analysis of 2-6Sia expression levels is conducted at the end of imaging and integrated into the cell lineage data. **D.** Utilizing the cell lineage data, deduced cell populations are created by focusing on specific characteristics of cancer cells using a cell fate simulation algorithm. **E.** The deduced cancer cell population, along with immune cells, is introduced into a 3D TME generated by artificial intelligence for the simulation. In this simulation, deduced cancer cells (depicted in blue) coexist with immune cells (depicted in yellow, green, and red) within a sphere.

To accomplish this, we conducted single-cell tracking analysis (*20, 34*) on established cancer cell lines (Fig. 1B). In order to generate data suitable for bioinformatic analysis, we devised a computerized single-cell tracking system (Supplementary Fig. 1). This procedure involved live cell imaging using near-infrared-illuminated Differential Interference Contrast (NER-DIC) microscopy (*20, 34*), which allowed us to create grayscale live cell imaging videos without introducing any phototoxicity. Then, we developed an algorithm to segment the grayscale DIC images with a wide range of cell densities, from sparse to densely populated populations (Supplementary Fig. 2A and B). This approach allowed us to track individual cell behaviors within these videos, enhancing our understanding of cellular dynamics. Subsequently, automated cell tracking was executed using an algorithm that analyzes the surrounding areas of the tracking-target region. It identifies the tracking-target area in the subsequent image frame by considering information from the surrounding areas (Supplementary Fig. 3). Through this tracking process, various cellular events, including multipolar cell division, can be detected. This results in the establishment of accurate cell lineage data, including the spatiotemporal behaviors of a cell that commence with the tracking process and encompass all of its progeny (Fig. 1C). After live cell imaging, cells can be labeled with specific antibodies or lectins targeting biomolecules or post-translational modifications, such as oligosaccharides bound to membrane proteins, for fluorescent imaging. The quantified fluorescent values associated with each cell were then integrated into the cell lineage data (Fig. 1C).

While cancer cell lines are derived from a single cell within the original cancer cell population, they may only partially represent the characteristics of the original population. To address this, we employed a cell fate simulation algorithm (*20*) (Fig. 1B and D). This algorithm calculates the probabilistic likelihood of a specific cellular event occurring subsequent to a preceding event, considering the duration between these events (*20*). For instance, it can assess the likelihood of multipolar cell division occurring after bipolar cell division, along with the associated duration. Consequently, the algorithm generates cell lineage data based on these probabilistic values. The growth dynamics of each cell population can be defined by combining these probabilistic values. Therefore, the cell fate simulation algorithm has the capability to simulate the growth of various cell populations when the requisite information to define these probabilistic values is available. For example, if we possess knowledge that the expression of 2-6Sia is notably high in highly reproductive cells, the algorithm can generate a deduced cell population in which reproductive cells exhibit a high level of 2-6Sia expression. Similarly, if we know the levels of variation of 2-6Sia expression *in vivo*, a digitally generated cell population (deduced cell population) that can reflect the attributes of growth patterns of cancer cells *in vivo* can be created. Subsequently, this deduced cell population was introduced into a 3D TME (Fig. 1B and E) alongside immune cells, allowing us to simulate the fate of these cells within the TME. This integration into a 3D TME model facilitates a comprehensive simulation of cancer cell dynamics, offering predictions on cell fate in response to the complex interplay within the TME.

### Distinct Characteristics of Cell Lines Derived from Cervical and Pancreatic Cancer

We conducted single-cell tracking using HeLa (cervical cancer) and MiaPaCa2 (pancreatic cancer) cell lines. The cell population size at each time point was determined through single-cell tracking data (Fig. 2A). Additionally, Supplementary Data 1, Supplementary Data 2, Supplementary Video 1, and Supplementary Video 2 provide cell lineage maps and videos illustrating the tracking processes. Our results reveal an increase in the population size of HeLa and MiaPaCa2 cells, with average cell doubling times of 26.6 and 22.7 hours, respectively.

**Fig. 2.**
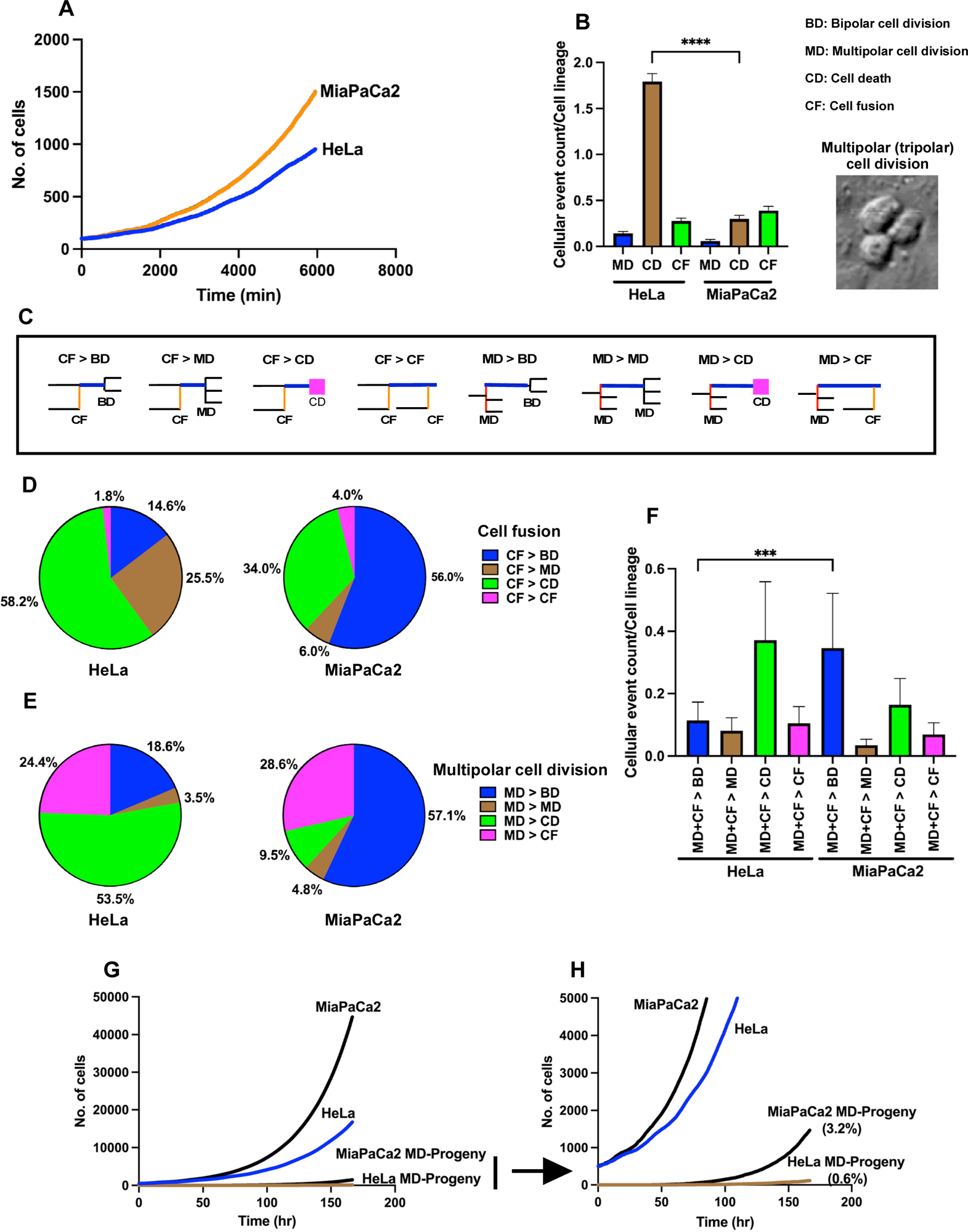
Characteristics of cervical cancer cell line (HeLa) and pancreatic cancer cell line (MiaPaCa2). Single-cell tracking was conducted with HeLa and MiaPaCa2 cells, plotting the number of cells at each time point to determine cell population expansion curves. B. The counts of multipolar cell divisions (MD), cell deaths (CD), and cell fusions (CF) occurring within cell lineages were recorded. Statistical analysis was performed using ordinary one-way ANOVA (Tukey’s), with ****P < 0.0001 indicating statistical significance. An example of a cell undergoing tripolar cell division is illustrated in a DIC image. C. The fate of cells resulting from cell fusion and the progeny produced by multipolar cell division were explored, with possible patterns illustrated (e.g., ‘CF > BD’ indicates bipolar cell division (BD) occurs following CF). D. The fate of cells resulting from cell fusion is detailed. E. The fate of progeny produced through multipolar cell division is presented. F. The total numbers of BDs, MDs, CDs, and CFs that occurred following CF or MD were determined, with statistical analysis conducted using ordinary one-way ANOVA (Kruskal-Wallis test) and ****P < 0.0001 indicating statistical significance. G. Cell fate simulation, based on single-cell tracking data of HeLa and MiaPaCa2 cells, determined the number of reproductive progeny derived from cells produced through multipolar cell division. H. A magnified view of the reproductive progeny results is shown, with ‘MD-Progeny’ referring to reproductive progeny produced through multipolar cell division. The percentage in parentheses indicates the proportion of reproductive progeny within the total HeLa or MiaPaCa2 cell population.

Next, we assessed the occurrence of multipolar cell divisions, cell death, and cell fusion in HeLa and MiaPaCa2 cells (Fig. 2B). Notably, MiaPaCa2 cells exhibited a lower frequency of multipolar cell divisions, cell death, and cell fusion compared to HeLa cells. Subsequently, we delved into the cellular events following multipolar cell divisions and cell fusion. In Fig. 2C, we have listed the patterns we analyzed. For instance, ‘CF (cell fusion) > BD (bipolar cell division)’ illustrates that bipolar cell division occurs following cell fusion. In the case of HeLa cells, over 50% of the progeny arising from multipolar cell division (MD) and cells resulting from cell fusion ultimately underwent cell death (CD) (Fig. 2D and E, HeLa). In contrast, among MiaPaCa2 cells generated through multipolar cell division (MD) or cell fusion (CF), more than 50% retained the capability to undergo bipolar cell division (BD) (Fig. 2D and E, MiaPaCa2). This difference between HeLa and MiaPaCa2 cells suggests that while MiaPaCa2 cells undergo multipolar cell division and cell fusion less frequently, the progeny resulting from these events have a higher probability of survival compared to HeLa cells (Fig. 2F, MD+CF > BD, HeLa vs. MiaPaCa2). Consequently, MiaPaCa2 cells have a greater chance of accumulating cells with altered chromosome numbers due to multipolar cell division and cell fusion, potentially making up 3.2% of the total cell population of MiaPaCa2 cells after 170 hours of growth (Fig. 2G and H).

### Expression of 2-6Sia in Cervical and Pancreatic Cancer Cell Lines

HeLa and MiaPaCa2 cell lines were subjected to immunostaining for the cancer marker TAG-72 (*45–47*), the stem cell marker CD133 (*48–50*), and the stem cell glycan marker fucose α1-2galactose α1-3-structure (*40*), recognized by the rBC2LCN bacterial lectin (Fig. 3A and B, respectively). The majority of HeLa cells exhibited broad expression of these markers, with most MiaPaCa2 cells also displaying these markers, except for the rBC2LCN lectin staining.

**Fig. 3.**
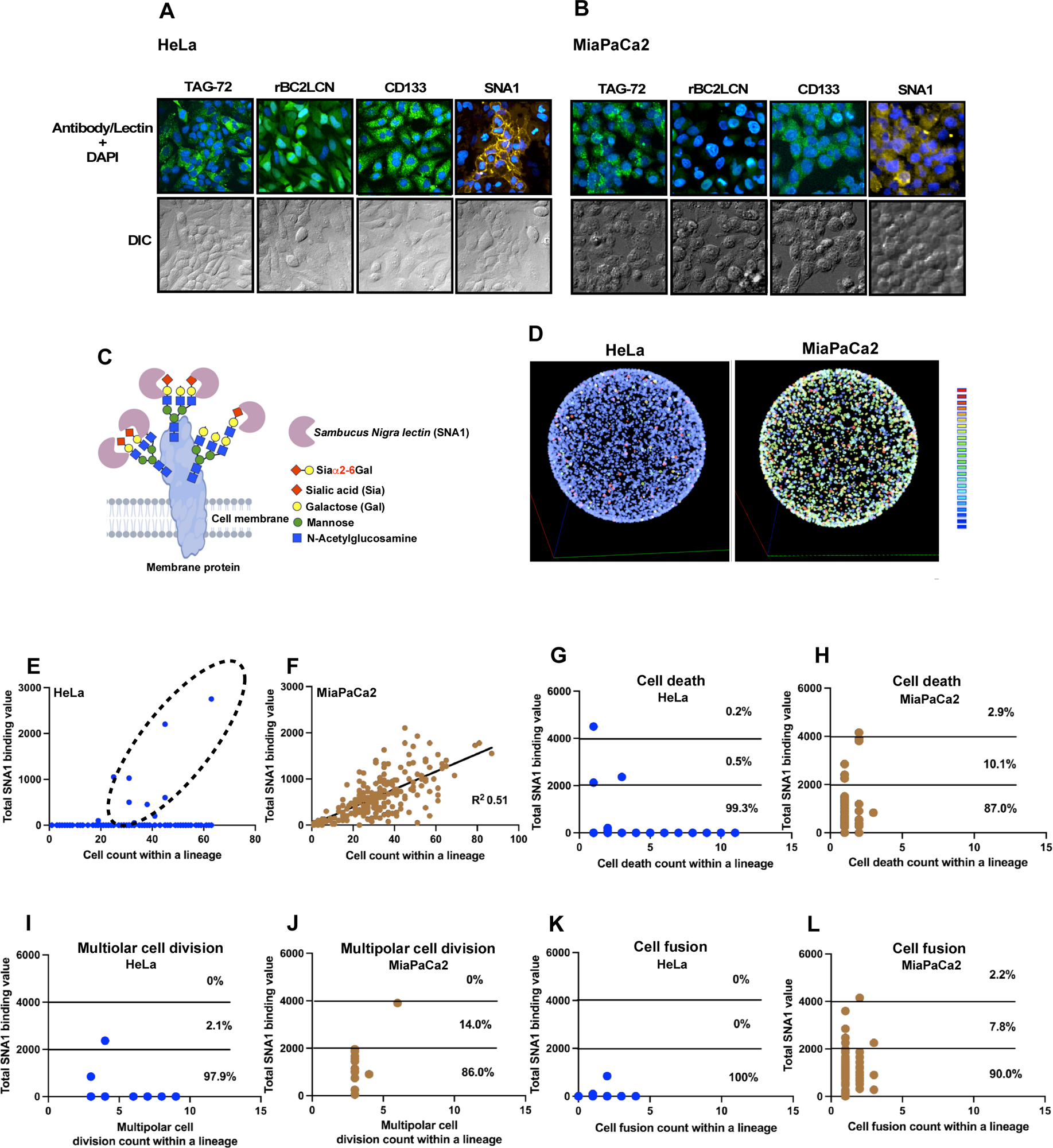
Analysis of 2-6Sia expression using SNA1. Fluorescence-tagged rBC2LCN, SNA1, anti-CD133, and anti-TAG-72 staining was performed on HeLa (**A**) and MiaPaCa2 (**B**) cells, followed by Alexa 488-labeled secondary antibody visualization and fluorescence imaging. Nuclei were counterstained with DAPI (blue), and DIC images were included. **C**. Illustration of membrane proteins with sialic acid α2-6 galactose (2-6Sia) termini oligosaccharides, recognized by *Sambucus Nigra* lectin (SNA1). **D**. Visualization of 2-6Sia expression in individual cells arranged on a spherical surface, color-coded by a heatmap scale. **E-F**. Cell counts within HeLa (**E**) and MiaPaCa2 (**F**) cell lineages, respectively, with total SNA1 binding values highlighted by a dotted circle for lineages with higher values. Statistical analysis used simple linear regression (R^2^ = 0.51). **G-H**. Cell death counts within HeLa (**G**) and MiaPaCa2 (**H**) lineages plotted against total SNA1 binding, categorized into groups by SNA1 values (<2000, 2000-3999, >4000) with percentages. **I-J**. Multipolar cell division counts within HeLa (**I**) and MiaPaCa2 (**J**) lineages against SNA1 binding, categorized similarly with percentages. **K-L**. Cell fusion counts within HeLa (**K**) and MiaPaCa2 (**L**) lineages against SNA1 binding, are also categorized with percentages shown.

The oligosaccharide structure attached to membrane proteins and terminated by α2-6 sialic acid structure, recognized by *Sambucus Nigra* lectin (Fig. 3C; SNA1), is suggested to be associated with the maintenance of stemness features and malignancy (*35–40, 43*). Interestingly, only a subset of HeLa cells expressed 2-6Sia (Fig. 3A, SNA1 staining), indicating that 2-6Sia-related stemness differs from other markers, such as CD133. Conversely, the majority of MiaPaCa2 cells displayed positive 2-6Sia expression (Fig. 3B, SNA1 staining). We employed single-cell tracking data linked to the SNA1 binding levels of individual cells to create a visual heatmap representation of these levels. In this visualization (Fig. 3D), individual cells were positioned on the surface of a sphere, illustrating that MiaPac2 cells exhibit elevated levels of 2-6Sia-related stemness compared to HeLa cells.

In Fig. 3E (HeLa) and F (MiaPaCa2), we plotted the total number of cells in a cell lineage on the horizontal axis, where a higher value represents a cell’s ability to produce a larger number of progeny (indicative of reproductive ability), and the total SNA1 binding levels of progeny within the cell lineage on the vertical axis. Consequently, if a cell lineage comprises a large number of cells, all expressing high levels of 2-6Sia, it will be positioned towards the upper-right quadrant of the graph. Notably, only a subset of the HeLa cell population expressed 2-6Sia, and these cells exhibited a higher reproductive ability, while 2-6Sia-negative cells showed varied reproductive ability (Fig. 3E). In contrast, the majority of MiaPaCa2 cells expressed 2-6Sia, and the levels of expression displayed a linear relationship with their reproductive ability (Fig. 3F). We also conducted similar analyses by plotting the cell death count (Fig. 3G for HeLa and Fig. 3H for MiaPaCa2), multipolar cell division count (Fig. 3I for HeLa and Fig. 3J for MiaPaCa2), and cell fusion count (Fig. 3K for HeLa and Fig. 3L for MiaPaCa2). In all instances, it was observed that cell death, multipolar cell division, and cell fusion predominantly occurred in cells exhibiting lower 2-6Sia expression levels (below 2000 of total SNA1 binding). These findings not only underscore the connection between 2-6Sia expression and the maintenance of cellular reproductive ability but also highlight its role in reducing the likelihood of cell death, multipolar cell division, and cell fusion. Taken together, these results suggest that only a subset of HeLa cells expressing 2-6Sia retain their stemness, contributing to a higher reproductive ability to maintain the cell population (Fig. 3E), consistent with our previous observation (*34*). The presence of 2-6Sia-negative cells increases the likelihood of cell death, multipolar cell division, and cell fusion (Fig. 3G, I, and K). In contrast, the majority of MiaPaCa2 cells maintain 2-6Sia-related stemness (Fig. 3F), resulting in fewer occurrences of cell death, multipolar cell division, and cell fusion (Fig. 3H, J, and L). This implies that 2-6Sia-related stemness plays a role in preserving the integrity of the cell population.

### Generation of Deduced Cell Populations and Designing the 3D TME Simulation

Our findings thus far indicate distinct patterns of 2-6Sia expression in HeLa and MiaPaCa2 cells, which could be related to the reproductive capability of these cells and their role in maintaining the integrity of the cell population. Specifically, our results suggest that 2-6Sia expression reduces the occurrence of cell fusion and multipolar cell division, events that can lead to the production of cells with altered chromosome numbers. In cancer tissues, the expression levels of 2-6Sia among cancer cells vary (*41–44*), implying that each cancer cell within a tissue may possess a unique reproductive ability and a chance to produce cells with altered chromosome numbers. This variation falls within the range of 10 to 200% of the average expression levels (*41–44*). To explore the effects of varying levels of 2-6Sia expression on cancer cell fate, we created deduced cell populations with 1.5, 1.0, 0.5, and 0.25-fold 2-6Sia expression relative to the levels found in HeLa and MiaPaCa2 cells, using the cell fate simulation algorithm (*20*) with some modifications. An overview of the deduced cell population generation is provided in Supplementary Fig. 4 and the Supplementary Text for Supplementary Fig. 4. The created cell populations are designated as follows: Cervical 2-6Sia 1.5, 1.0, 0.5, and 0.25, and Pancreatic 2-6Sia 1.5, 1.0, 0.5, and 0.25. In essence, the cell fate simulation algorithm constructs a digitally generated cell population (deduced cell population), where events that occur within each cell composing the population and its progeny, such as cell division, cell death, and cell fusion, are determined. This allows for the generation of cell lineage maps that visually represent the fate and genealogical relationships of each cell within the population. Cell lineage maps of Cervical 2-6Sia 1.5 and Pancreatic 2-6Sia 1.5 are provided in Supplementary Data 3 and Supplementary Data 4, respectively. Fig. 4A (HeLa) and Fig. 4B (MiaPaCa2) depict the cell population expansion curves, which include empirical tracking results (light blue line) and deduced cell populations (blue lines). The deduced cell populations generated using single-cell tracking data of HeLa cells show a ∼5% slower rate of population expansion compared to actual HeLa cells, as the cell fate simulation algorithm reflects the averaged proliferation pattern of the given cell population. In Fig. 4C, the levels of 2-6Sia in the deduced cell populations are depicted. We used the deduced cell populations for 3D TME simulation, carried out by positioning these cells and immune cells within a 3D space (sphere) (see details in Supplementary Materials and Methods).

**Fig. 4.**
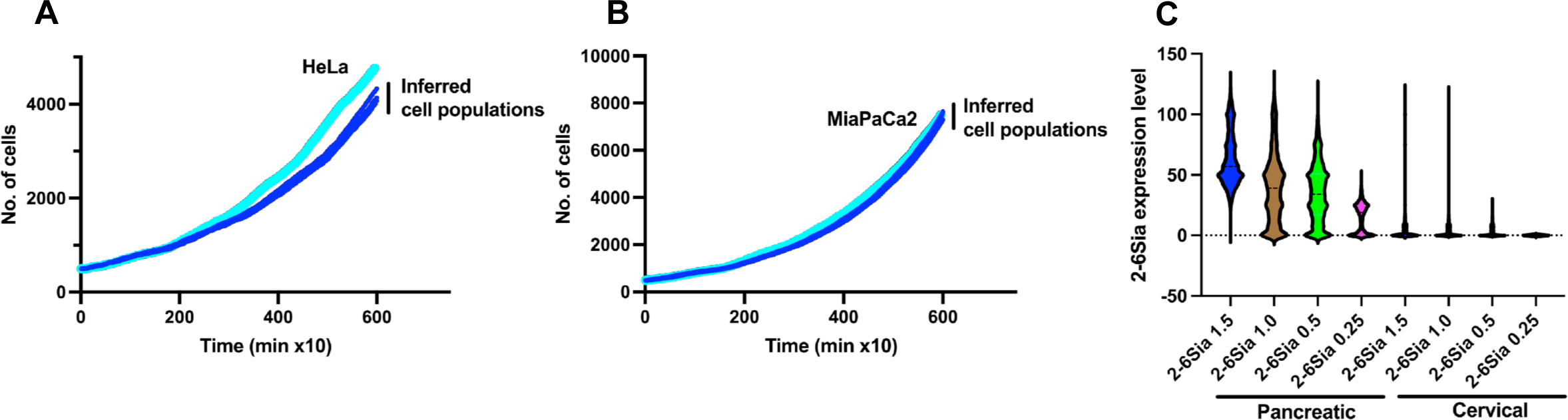
Generation of deduced cell populations. **A** (HeLa; light blue line and Cervical 2-6Sia; blue lines) and **B** (MiaPaCa2; light blue line and Pancreatic 2-6Sia; blue lines). Deduced cell populations were generated using a cell fate simulation algorithm by varying 2-6Sia expression levels. The cell population expansion curves for these cells are shown. **C.** The expression levels of 2-6Sia in the deduced cell population are presented. As the cell fate simulation algorithm incorporated the recorded 2-6Sia expression patterns from cell lineage data, the deduced cell population’s 2-6Sia expression levels reflect those of HeLa and MiaPaCa2 cells. These cell populations were designated as Cervical 2-6Sia 1.5, 1.0, 0.5, and 0.25, and Pancreatic 2-6Sia 1.5, 1.0, 0.5, and 0.25.

### Categorization of TME-Resident Cells for 3D TME Simulation

To streamline the development of the 3D TME simulation algorithm, we categorized TME-resident cells into three groups: Suppressive, Permissive, and Lethal (Fig. 5A). Suppressive cells, which include CD4^+^ T cells, CD8^+^ T cells, and B cells, contribute to an environment that inhibits cancer cell growth. Permissive cells include macrophage M2, regulatory T cells, and endothelial cells, all of which create a microenvironment favorable to cancer cell proliferation. Lethal cells consist of cytotoxic T cells and natural killer cells. Some cell types, such as M1 macrophages and neutrophils, can exhibit dual roles, either suppressing or promoting cancer cell growth depending on the context (*25–28*). Due to this complexity, we chose not to include these cells in the 3D TME simulation.

**Fig. 5.**
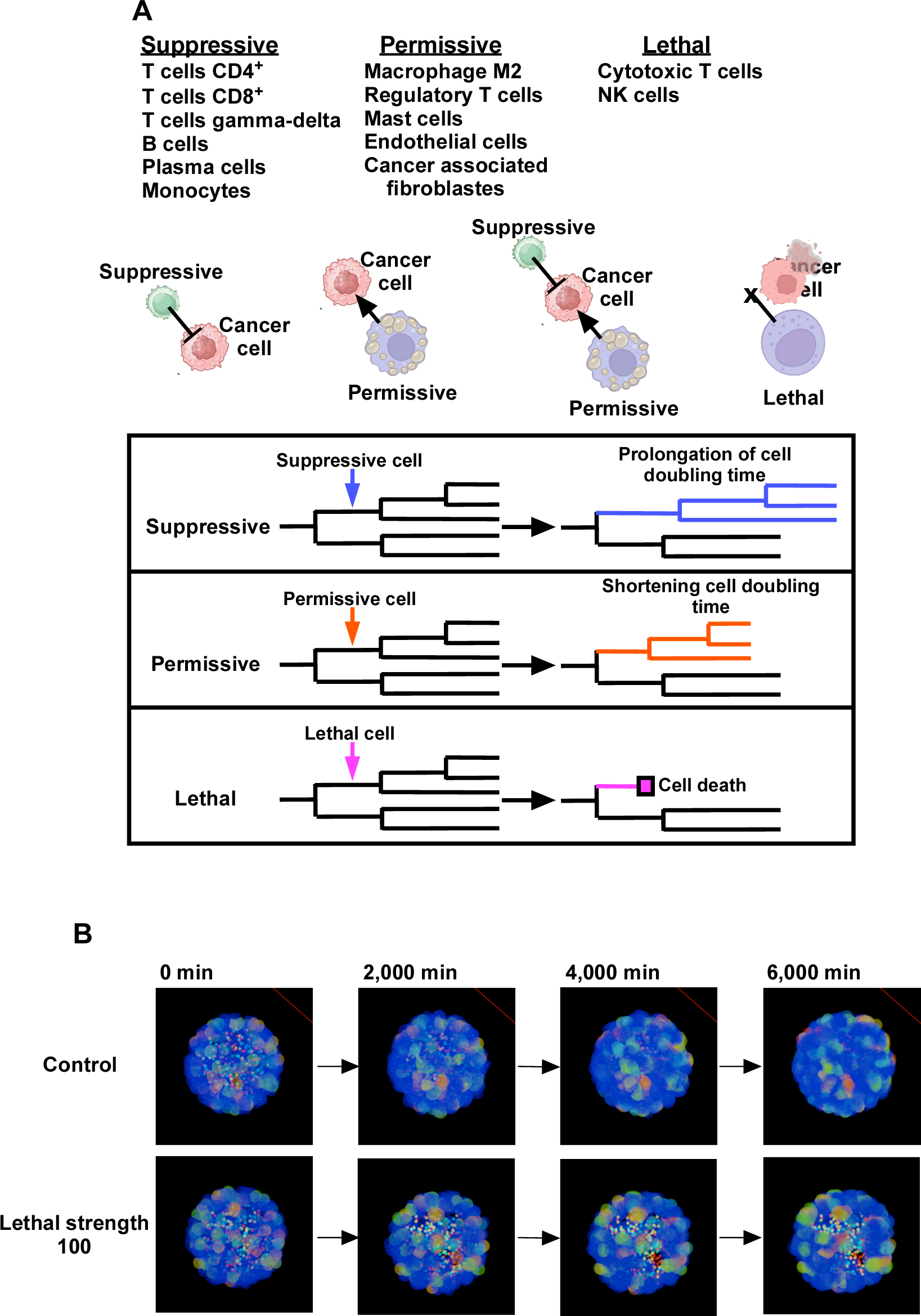
Categorization of immune cells. **A.** Immune cells within the TME were classified into Suppressive, Permissive, and Lethal cells. When a cancer cell is located within a defined spatial distance of a Suppressive cell, it experiences an extended doubling time. The doubling time of its progeny is also extended. When a cancer cell is within a specified spatial distance of a Permissive cell, it experiences a shortened doubling time. The doubling time of its progeny is also shortened. When both Suppressive and Permissive cells are within a specified distance, their impact on cancer cells depends on their respective distances from the cancer cell. When a cancer cell is within a specified spatial distance of a Lethal cell, cell death is marked to the cell, and its progeny are removed from the cell lineage database. **B.** A 3D TME simulation was conducted using Cervical 2-6Sia 1.5 cells in the presence of Suppressive, Permissive, and Lethal cells, represented by red, light blue, and yellow spheres, respectively (denoted as smaller spheres). The expression levels of 2-6Sia were visualized using a heatmap scale, ranging from blue to red (denoted as a larger size of the spheres).

Within the 3D TME simulation, Suppressive, Permissive, and Lethal cells were placed alongside deduced cancer cells that were generated digitally. These cancer cells undergo cell division, cell death, or cell fusion at specific moments, as represented by cell lineage maps (Supplementary Data 3 and Supplementary Data 4). In scenarios where immune cells do not influence cancer cells, the latter proliferate in 3D space according to their cell lineage map data (Fig. 5A). The effects of encountering Suppressive, Permissive, or Lethal cells are recorded by modifying the data. When a deduced cancer cell encounters a Suppressive cell, the doubling times for the cell and its progenies are lengthened, slowing their proliferation. Conversely, when a deduced cell encounters a Permissive cell, the doubling times of the deduced cells and their progenies are reduced, enhancing proliferation rates. Interactions with a Lethal cell result in the removal of the deduced cell and its progenies from the simulation (see details in Supplementary Materials and Methods).

To perform the 3D TME simulation, several parameters are considered, including: [1] the resistance levels of deduced 2-6Sia-expressing cells against Suppressive and Lethal cells, [2] cell ratios: establishing the relative numbers of Suppressive, Permissive, and Lethal cells in relation to deduced cancer cells, and [3] impact strength: the strength of the influence exerted by Suppressive, Permissive, and Lethal cells on deduced cancer cells. Details for setting these parameters, along with control data, are provided in Supplementary Pseudo Code 1, Supplementary Text for Pseudo Code 1, and Supplementary Fig. 5. Graphical representations of the 3D TME, depicting views at 0, 2000, 4000, and 6000 minutes, are shown in Fig. 5B. The simulation was conducted with Cervical 2-6Sia 1.5 cells under Control conditions or the influence of Lethal cells.

### Simulations with Pancreatic and Cervical Cancer TME

We conducted 3D TME simulations using TME cell landscape data from 193 cervical and 172 pancreatic cancer patients sourced from the TIMEDB database (*51*). Leveraging the ConsensusTME dataset, we classified immune cells, cancer-associated fibroblasts, and endothelial cells into three categories: Suppressive, Permissive, and Lethal cells. We calculated the percentage of each cell category and introduced the corresponding number of cells into the 3D TME, based on percentages unique to each cancer case (refer to Supplementary Data 5 for the calculated percentages). For example, if the Suppressive cell percentage was determined to be 45%, we placed 225 Suppressive cells alongside 500 deduced cancer cells in the simulation. We selected specific setting values for Suppressive, Permissive, and Lethal strength, confirmed to impact the population expansion of deduced cancer cells, as detailed in Supplementary Fig. 5. Four deduced cell populations were employed for both cervical (2-6Sia 1.5, 1.0, 0.5, and 0.25) and pancreatic (2-6Sia 1.5, 1.0, 0.5, and 0.25) cancers. Accordingly, simulations were conducted for each of the 193 cervical and 172 pancreatic cancer cases, using their respective cell populations. In Figs. 6-8, each dot represents the outcome of simulations using the TME cell landscape data for each cancer case. Specifically, in Figs. 7-8, the blue, brown, light blue, and green dots indicate results obtained with 2-6Sia 1.5, 1.0, 0.5, and 0.25, respectively, illustrating the impact of varying 2-6Sia expression levels on simulation outcomes.

**Fig. 6.**
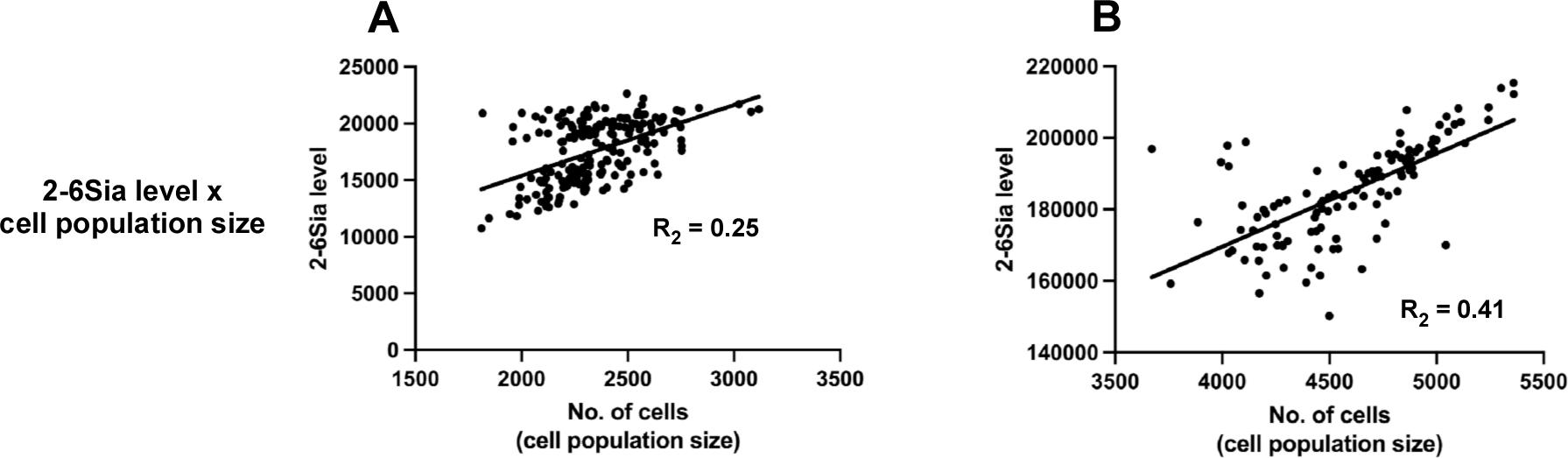
Relationships of simulation results with 2-6Sia expression. Simulation results are presented for Cervical 2-6Sia 1.0 (**A**) and Pancreatic 2-6Sia 1.0 (**B**). The simulated cell population size is plotted against 2-6Sia expression levels. The resulting R^2^ values were shown. Each dot signifies the outcome of simulations using the TME cell landscape data of each cancer case.

**Fig. 7.**
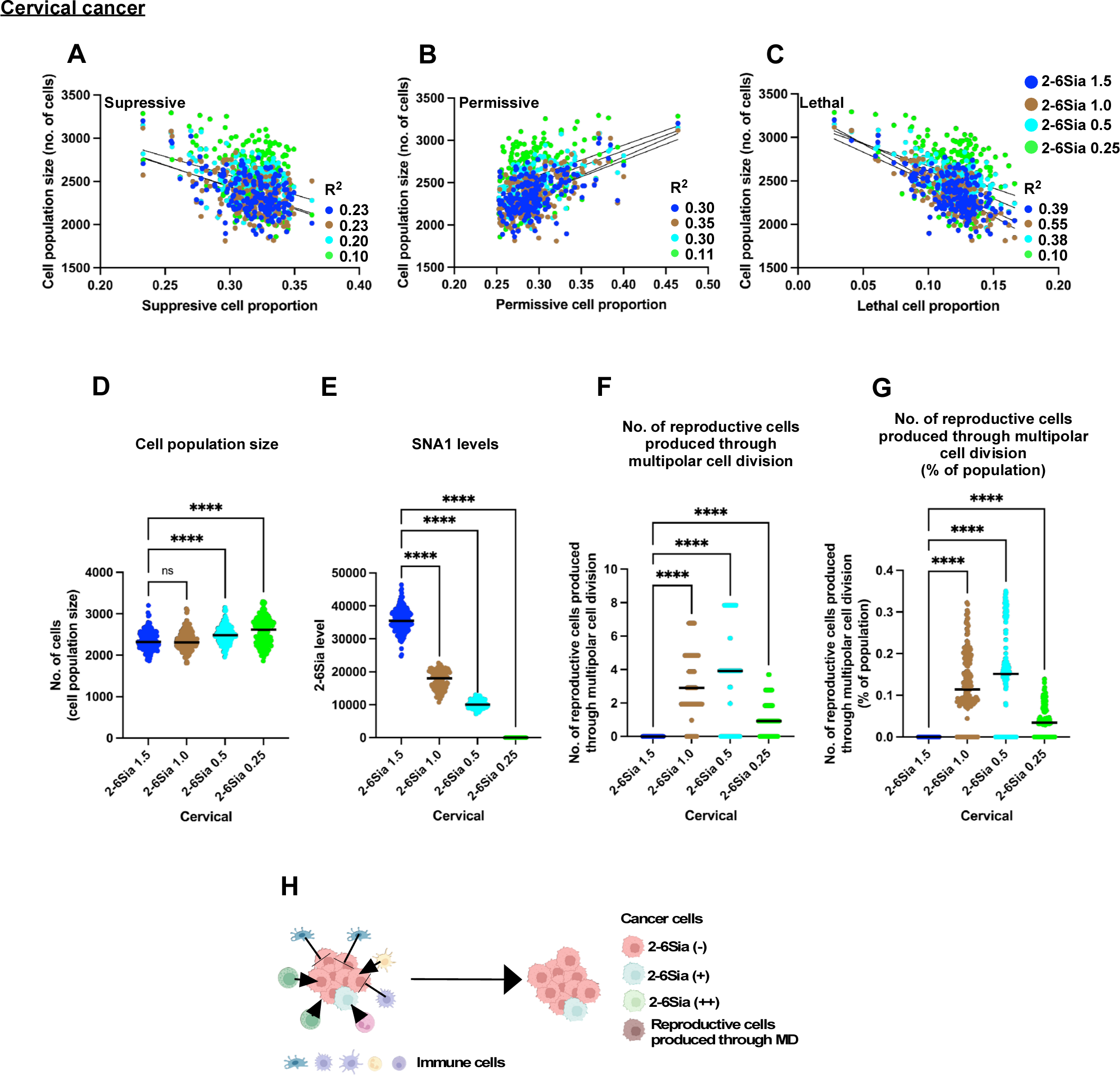
Characteristics of cervical cancer cells within the TME. Four different populations of Cervical 2-6Sia cells: 2-6Sia 1.5, 2-6Sia 1.0, 2-6Sia 0.5, and 2-6Sia 0.25, were employed in 3D TME simulations using TME cell landscape data from 193 cervical cancer cases. **A-C**. The simulated population size of Cervical 2-6Sia cell populations is plotted against the initial proportion of Suppressive cells (**A**), Permissive cells (**B**), and Lethal cells (**C**). Variation of each data relative to the linear regression line was calculated. The resulting R2 values were shown. **D-G**. Cell population sizes (**D**), 2-6Sia expression levels (**E**), and the number of reproductive cells produced through multipolar cell division (**F**) of Cervical 2-6Sia (1.5, 1.0, 0.5, and 0.25) were analyzed. The percentage of the number of reproductive cells produced through multipolar cell division relative to the total cell population is also shown (**G**). Statistical analysis was conducted using ordinary one-way ANOVA (Tukey’s), and significance is denoted as ****P < 0.0001, while “ns” indicates no significance. The results of the analysis are shown between Cervical 2-6Sia 1.5 and the other populations. **H**. The summarized characteristics of cervical cancer cells within the TME context are presented. Each dot signifies the outcome of simulations using the TME cell landscape data of each cancer case.

**Fig. 8.**
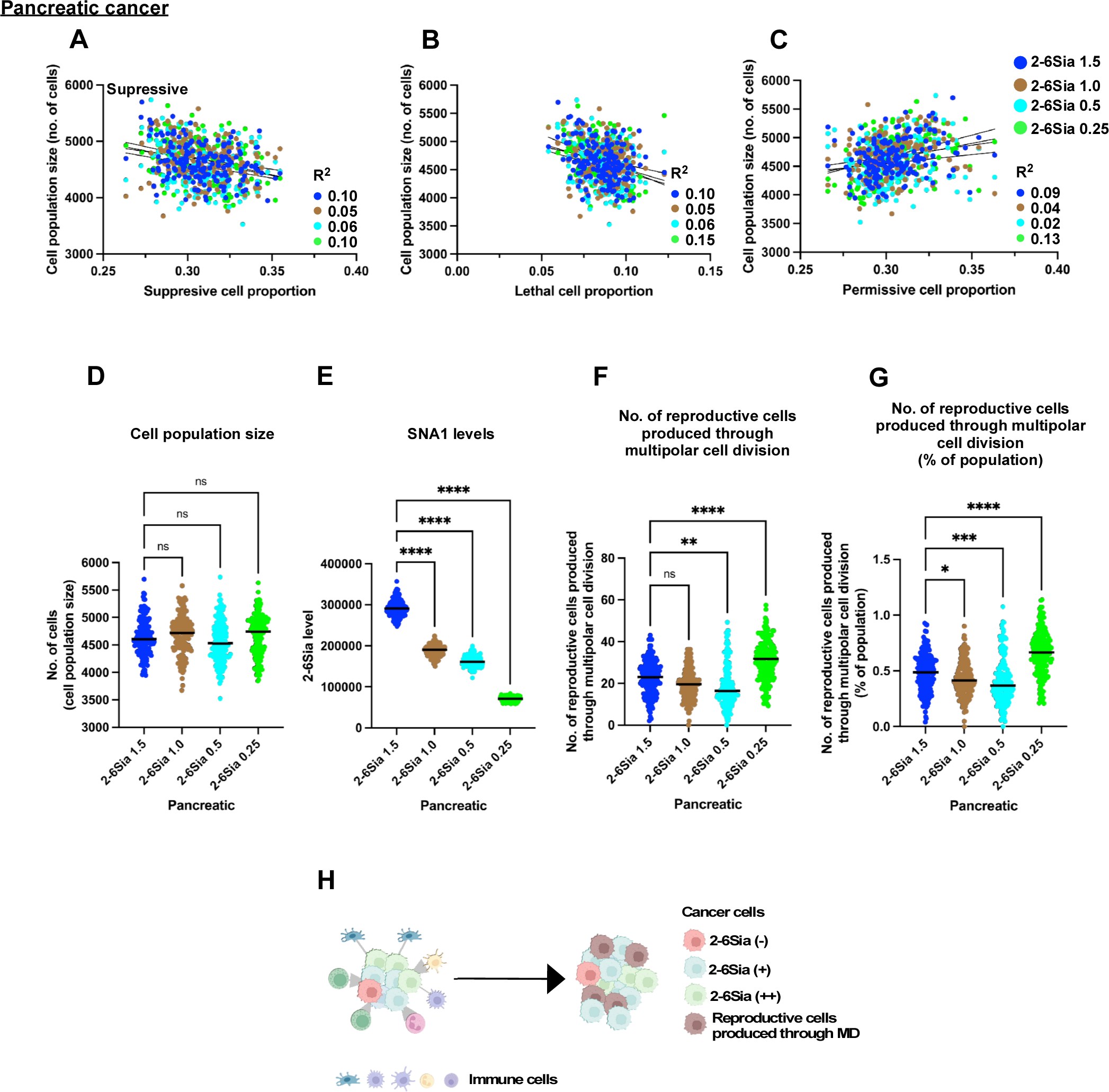
Characteristics of pancreatic cancer cells within the TME. Four different populations of Pancreatic 2-6Sia cells: 2-6Sia 1.5, 2-6Sia 1.0, 2-6Sia 0.5, and 2-6Sia 0.25, were employed in 3D TME simulations using TME cell landscape data from 172 pancreatic cancer cases. **A-C.** The simulated population size of Pancreatic 2-6Sia cell populations is plotted against the initial proportion of Suppressive cells (**A**), Permissive cells (**B**), and Lethal cells (**C**). Variation of each data relative to the linear regression line was calculated. The resulting R^2^ values were shown. **D-G.** Cell population sizes (**D**), 2-6Sia expression levels (**E**), and the number of reproductive cells produced through multipolar cell division (**F**) of Pancreatic 2-6Sia (1.5, 1.0, 0.5, and 0.25) were analyzed. The percentage of the number of reproductive cells produced through multipolar cell division relative to the total cell population is also shown (**G**). Statistical analysis was conducted using ordinary one-way ANOVA (Tukey’s), and significance is denoted as **P < 0.01 and ****P < 0.0001, while “ns” indicates no significance. The results of the analysis are shown between Pancreatic 2-6Sia 1.5 and the other populations. **H**. The summarized characteristics of pancreatic cancer cells within the TME context are presented. Each dot signifies the outcome of simulations using the TME cell landscape data of each cancer case.

### Relationship between simulated cell population size and 2-6Sia expression levels

To examine the relationship between simulated cell population size and 2-6Sia expression levels, we performed a comparative analysis using Cervical 2-6Sia 1.0 cells and Pancreatic 2-6Sia 1.0 cells within TME cell landscapes. We observed a positive correlation between the simulated cell population size and 2-6Sia expression levels, as shown in Fig. 6A (R^2^ = 0.25 for Cervical) and Fig. 6B (R^2^ = 0.41 for Pancreatic). This finding suggests the significant role of 2-6Sia expression in maintaining the cell population.

### Characteristics of cervical cancer cells within the TME

We investigated the relationship between the simulated population size of Cervical 2-6Sia cells and the proportions of Suppressive (Fig. 7A), Permissive (Fig. 7B), or Lethal (Fig. 7C) cells, based on the cell landscape data of cervical cancer TME from the TIMEDB database (*51*). The results revealed correlations between the simulated cell population size of Cervical 2-6Sia cells and the proportions of Suppressive, Permissive, and Lethal cells within the TME of cervical cancer (Fig. 7A-C; R^2^ > 0.1-0.55, where each dot represents the outcome of a simulation for each cancer case, see Supplementary Data 5 for the proportions). Notably, negative linear correlations were observed with the proportions of Suppressive (Fig. 7A) and Lethal (Fig. 7C) cells, while a positive correlation was found with the proportion of Permissive cells (Fig. 7B). These findings suggest that the proliferation of Cervical 2-6Sia cells is significantly influenced by the composition of immune cells within the TME for each cancer case. Among the Cervical 2-6Sia cell populations, the average cell population size of Cervical 2-6Sia 0.25 was observed to be 10% larger than that of Cervical 2-6Sia 1.5 (Fig. 7D). As expected, Cervical 2-6Sia 1.5 cells exhibited the highest levels of 2-6Sia expression, which gradually decreased in the Cervical 2-6Sia 1.0, 0.5, and 0.25 populations (Fig. 7E). The contribution of reproductive progeny produced through multipolar cell division was marginal (Fig. 7F), with such progeny accounting for less than 0.4% of the Cervical 2-6Sia cell population (Fig. 7G).

These findings underscore the distinctive characteristics of cervical cancer cells within the TME, highlighting the significant influence of the immune cell landscape on the proliferation of cervical cancer cells (Fig. 7H).

### Characteristics of pancreatic cancer cells within the TME

We then explored the relationship between the simulated population size of Pancreatic 2-6Sia cells and the proportions of Suppressive (Fig. 8A), Permissive (Fig. 8B), or Lethal (Fig. 8C) cells. Unlike with Cervical 2-6Sia cells, we found almost no significant correlations (R^2^ < 0.15, where each dot represents the outcome of a simulation for each cancer case, see Supplementary Data 5 for the proportions) for Pancreatic 2-6Sia cells. This suggests that the proliferation of Pancreatic 2-6Sia cells was relatively unaffected by the immune cell composition within the TME. Among different Pancreatic 2-6Sia cell populations, we observed no significant differences in population size (Fig. 8D). Pancreatic 2-6Sia 1.5 cells exhibited the highest levels of 2-6Sia expression, which gradually decreased in the Pancreatic 2-6Sia 1.0, 0.5, and 0.25 populations, as expected (Fig. 8E). Concerning the number of reproductive cells produced through multipolar cell division, there was a reduction in the number for Pancreatic 2-6Sia 0.5 cells but an increase for the Pancreatic 2-6Sia 0.25 cell population compared to Pancreatic 2-6Sia 1.5 cells (Fig. 8F and G, averaging from 0.48 in Pancreatic 2-6Sia 1.5 cells to 0.66 in Pancreatic 2-6Sia 0.25 cells). This suggests that although varying levels of 2-6Sia did not substantially impact the simulated cell population size, cells with altered chromosome numbers expanded within the Pancreatic 2-6Sia cell population, particularly in those with 0.25-fold levels of 2-6Sia expression.

These results underscore the unique characteristics of pancreatic cancer cells (Fig. 8H). The majority of pancreatic cells expressing 2-6Sia seemed to be less influenced by the TME cell landscape. In addition, cells with altered chromosome compositions accumulated in populations expressing lower levels of 2-6Sia, introducing genetic diversity within the pancreatic cell population.

## Discussion

In this study, we empirically obtained data related to the spatiotemporal behavior of individual cells using a computerized single-cell tracking system with two established cancer cell lines. We then deduced cell populations, signifying particular characteristics of cancer cells *in vivo*, using the cell fate simulation algorithm we previously developed (*20*). Additionally, we simulated the impact of the immune cell landscape in the TME on these deduced cells using a 3D TME simulation, focusing on the role of 2-6Sia-related stemness and the reproductive cells produced through multipolar cell division. To date, extensive data have been accumulated at the gene, protein, cell, and tissue levels regarding cancer cells and the TME. However, these data often represent the status of cancer tissue or cells at a specific moment within the long process of cancer development. Thus, we believe that incorporating a temporal context based on empirical data, coupled with simulations that account for the heterogeneous nature of the cancer cell population, provides a deeper understanding of the dynamics of cancer development.

In our study on cervical cancer cells, we have demonstrated a significant influence of the immune cell landscape within the TME on the fate of cervical cancer cells, mainly due to the lack of the expression of 2-6Sia, which is known to be associated with stemness (*35–40*). These results allow us to predict that the majority of cervical cancer cell populations lacking 2-6Sia-related stemness may exhibit sensitivity to cytotoxic treatments, reflecting the general response of cervical cancer to treatment. Unlike cervical cancer cells, our study reveals that the fate of pancreatic cancer cells is relatively less influenced by the immune cell landscape within the TME. This difference can be attributed to the fact that the majority of MiaPaCa2 cells express 2-6Sia, which is known to be related to resistance to cell killing (*52–54*). Consequently, cells with high 2-6Sia expression levels show reduced sensitivity to immune cells in the TME. With regard to 2-6Sia expression, it has been reported that the glycan modification status with 2-6Sia is subject to dynamic regulation. For example, circulating α2-6Sialyltransferase 1 can modify cell surface glycoproteins with 2-6Sia when CMP-sialic acid is released from inflammation-activated platelets (*55*). This implies that cancer cells that do not express 2-6Sia can be converted into 2-6Sia-expressing cells by this platelet-related mechanism. The levels of 2-6Sia expression on pancreatic cancer cells thus may be subject to various regulatory mechanisms.

With regard to 2-6Sia expression, we found that a high level of expression contributes to maintaining the integrity of cells by preventing them from undergoing cell death, cell fusion, and multipolar cell division. In addition, cells expressing 2-6Sia exhibit stable growth, suggesting that once such a population is established, it can proliferate stably. This stable proliferation may, in turn, be recognized clinically as malignant cancer. However, while the accumulation of abnormalities through aberrant events is characteristic of cancer, high levels of stemness may counteract these events. Thus, the coexistence of high stemness and malignant characteristics presents a paradox. Perhaps cells with initially high stemness undergo a phase of reduced stemness, allowing for the acquisition of abnormalities, or cells with moderate stemness—sufficient to permit aberrant events but not prevent them—may be pivotal in cancer progression. Indeed, it has been demonstrated that invasive cancer cells can arise from populations with reduced 2-6Sia expression levels (*56*). Therefore, identifying cancer cells that balance the propensity for change with maintaining integrity may be crucial for understanding cancer progression.

Finally, we have showcased the framework of the 3D TME simulation here. By extending this framework to consider various other cancer-associated characteristics, we believe this framework will be able to shed light on aspects that have been overlooked.

## Materials and Methods

See Supplementary Materials and Methods

## Supporting information

Supplementary Text for Pseudocode 1

Supplementary Text for Supplementary Fig 4

Supplementary Pseudocode 1

Legends for Supplementary Videos-Figs

## ACKNOWLEDGMENTS

We give our special thanks to Philip C. Hanawalt (Stanford University) for encouraging us to continue developing single-cell tracking technology. We thank the Bioimaging Platform of the Research Centre for Infectious Diseases, Axe of Infectious and Inflammatory Diseases at the CHU de Quebec Research Centre for their invaluable technical support with the microscopes. We also acknowledge the Developmental Studies Hybridoma Bank for providing the monoclonal antibody against CD133, which is created by the U.S. National Institutes of Health, Institute of Child Health and Human Development, and maintained at the Department of Biology, University of Iowa (Iowa City, IA, USA). This research received financial support from the Canadian Foundation for Innovation and the Canadian Institutes for Health Research.

## AUTHOR CONTRIBUTIONS

M.S.S. and S.S. designed the study, conducted experiments, and contributed to the manuscript’s writing. M.S.S. developed the software for the computerized single-cell tracking system, cell fate simulation, and 3D TME simulation. A.R. conducted experiments, analyzed single-cell tracking data, and contributed to the manuscript’s editing.

## DATA AND CODE AVAILABILITY STATEMENT

All data generated or analyzed during this study are included in the paper and supporting files.

