## Supplementary Materials and Methods for "Cell Fate Simulation Reveals Cancer Cell Features in the Tumor Microenvironment"

### Cell culture and cell plating.

HeLa cells were cultured in Dulbecco's modified Eagle's medium (DMEM) supplemented with 10% fetal bovine serum (FBS) in a humidified atmosphere with 5% $CO^2$. MiaPaCa2 cells were maintained in DMEM supplemented with 10% FBS and 2% horse serum under a 5% $CO_2$-humidified atmosphere. For cell plating, approximately 3500 cells in 50 µl of cell suspension were added to the center of each well of a Lab-Tek™ II 8 Chamber Slide. Subsequently, 0.75 ml of culture medium was gently added to each well. The Lab-Tek™ II 8 Chamber Slide was placed on a microscope stage 24 hours after plating.

### Culture conditions on the microscope stage.

Cells were maintained in an environmental chamber (Live Cell Instruments, Korea) set at 37°C with 80% relative humidity. To prevent an increase in medium pH within the Lab-TekII 8-well chamber, the $CO_2$ concentration was controlled at 7.5%. To minimize medium evaporation, a NIR-DIC optimal glass lid (Live Cell Instruments) was placed on the Lab-Tek™ II 8 Chamber Slide. The typical medium evaporation rate was 10 µl/24 hours, and as a result, 700–800 µl of medium was added per well, ensuring cells could be cultured for at least one week without the need for medium changes.

For monitoring pH during long-term live cell imaging, phenol red was included in the medium. Importantly, this inclusion did not interfere with NIR-DIC imaging.

### Live cell imaging microscopy.

A custom microscope was constructed using an Olympus IX81 microscope frame (Quorum Technologies, Ontario, Canada). NIR-DIC imaging was chosen to minimize phototoxicity and reduce sensitivity to light distortion caused by plastic partitions and surface irregularities in the culture medium. The microscope featured dual imaging capabilities, combining NIR-DIC and fluorescent imaging for both live and fixed samples. Two distinct light paths were established within the microscope. For NIR-DIC imaging, NIR light was generated using light-emitting diodes, and passed through a polarizer, Nomarski prism, and condenser before illuminating the cells. The returning light passed through an objective lens and Nomarski prism, ultimately reaching the first charge-coupled device (CCD) camera (Camera 1, Hamamatsu Photonics, Image EM, 512×512 pixels). In the case of fluorescent imaging, laser light was directed onto cells stained with fluorescence-conjugated antibodies or proteins. This was accomplished by passing the laser light through a Nipkow disk (Yokokawa-Quorum Technologies) and an objective lens to excite fluorophores. The emitted light was collected by the Nipkow disk and captured using the second CCD camera (Camera 2, Hamamatsu, Image EM, 512×512 pixels). The system allowed the use of ×10–×40 objectives. In this study, an Olympus ×10 dry objective (UPlanSApo, 10×/0.40 NA, α/0.17/FN26.5) or ×20 dry objective (UPlanSApo, 20×/0.75 NA, α/0.17/FN2G.5) was employed. To achieve equivalent magnifications of ×15 and ×30, respectively, a ×1.5 coupler (Quorum Technologies) was inserted into the light path leading to the CCD cameras. A precision piezo XY stage with absolute measurement capabilities was utilized. An environmental chamber (Live Cell Instrument) was mounted on the piezo stage. The entire microscope system, along with the piezo XY stage, was controlled using Metamorph software (Quorum Technologies) on a Windows computer. Images were acquired using Metamorph's multi-dimensional acquisition mode (MDA). For NIR-DIC imaging, a 34 ms exposure time was used. The exposure time for fluorescence imaging was adjusted based on the fluorescence intensity. Typically, 20–80 z-planes at 1 μm intervals were acquired for NIR-DIC imaging. The resulting z-plane NIR-DIC images, captured by Camera 1, were saved as 512×512 pixel multi-layer TIFF files. For fluorescence imaging, z-plane NIR-DIC and fluorescence images generated by CCD cameras 1 and 2, respectively, were merged using a macro program in Metamorph. This produced 512×1024 pixel multi-layer TIFF files. To manage data, Computer 1 (Windows computer) was connected to Computer 2 (Macintosh) via an Ethernet cable. Computer 2 was responsible for creating live cell videos. Image files generated by Computer 1 were promptly transferred to Computer 2 using an in-house *File Transfer* software program, with subsequent deletion of the image files from Computer 1.

**Developed software for a single-cell tracking system.**

Image files produced by Metamorph underwent processing through a series of in-house software tools (Supplementary Fig. 1). The computerized single-cell lineage tracking analysis system consisted of two computers: Computer 1 (Windows) controlled the microscope, while Computer 2 (Macintosh) handled image processing, single-cell tracking, and data analysis tasks. Computer 1 was equipped with commercially available image acquisition software (Metamorph) to oversee microscope control and image file creation. Computer 2 utilized several in-house software programs (italic names indicate the in-house software). *Image Processing Controller* managed other software programs. *File Transfer* and *Map* facilitated communication between Computers 1 and 2. *File Converter* imported images generated by other microscopes. *Name Assignment*, *Focal Image Selection*, and *Contrast Set* were responsible for creating live cell videos. *Data Backup* controlled file archiving and backup procedures. *Outline Drawing* performed image segmentation and established the object segmentation library. *Movie Viewer* played movies and fine-tuned image quality. *Object Tracking Controller*, in conjunction with *Automatic Object Tracking*, created the cell lineage database and facilitated data verification. *Data Analysis* offered various options for data analysis. The data generated through this process was subsequently used for cell fate simulation and 3D TME simulation.

**Setting FOVs and adjustment of focus.**

Cells cultured in each well of the Lab-Tek™ II 8 Chamber Slide were simultaneously monitored. To cover the area of interest with multiple fields of view (FOVs), 2D image acquisition arrays were established in each well (*1*). The configuration of FOVs for these arrays and the necessary focus adjustments were executed using *Map* software in conjunction with Metamorph. *Map* software was specifically designed to coordinate and share information on the xyz positions between Metamorph on Computer 1 and software on Computer 2. To initiate this coordination, the objective lens was initially positioned at the left corner of Well 2 of the Lab-Tek™ II 8 Chamber Slide, and this position was registered in *Map* software. Subsequently, the outline of a Lab-Tek™ II 8 Chamber Slide was traced on the screen. The objective lens was then directed to a FOV of interest within a well using the outline as a guide. To view the cells in the surrounding area, 5x5 FOV dimensions were captured using Metamorph, which generated xyz position data for each FOV. A suitable area for live cell imaging was searched by repeating this process. When the suitable area was found, xyz position data saved on Computer 1 was transferred to Computer 2 using *Map* software. Then, the selection of the area to be imaged was finalized by setting the 2D image acquisition arrays that define each position of FOV using *Map* software. Next, the optimal objective z position for each FOV was determined, and the xyz position information created by Map software was compiled into a multi-dimensional acquisition (MDA) file format. This MDA file was then used by Metamorph to acquire an image of each FOV. The complied MDA file, created by *Map* software, was transmitted to Computer 1 and uploaded to Metamorph. Subsequently, Metamorph performed MDA image acquisition to generate images for the corresponding FOVs.

**Starting image acquisition.**

The first round of image acquisition was initiated by Metamorph through MDA image acquisition (Supplementary Fig. 1). Subsequently, *File Transfer* transmitted the multi-layer TIFF files (512×512 pixels) generated by Metamorph (Computer 1) to Computer 2. To facilitate file archiving, each multi-layer TIFF file was assigned a specific file name using *Name Assignment*. The image files underwent processing using *Focus Image Selection*, which involved selecting a focal image from among the images within a multi-layer TIFF file or generating an all-in-focus image. The all-in-focus image was constructed by selecting the optimal grayscale value from the z-axis data for each pixel. Following this, the contrast of the resulting images was adjusted, and image backgrounds were corrected using *Contrast Set*. Since the background patterns varied for each FOV, custom background patterns were generated using data from single-layer TIFF files. Each of these background patterns was then applied to the corresponding FOVs. Ultimately, stitched images were created using *Contrast Set* after contrast adjustment, background correction, and fine-tuning of the position for each FOV.

**Automated long-term live cell imaging video creation.**

Following the initial setup, automated image acquisition commenced. Metamorph captured images of each FOV at 10-minute intervals. Subsequently, after each round of image acquisition, the process involving *File Transfer*, *Name Assignment*, *Focus Image Selection*, and *Contrast Set* was coordinated by *Image Processing Controller*. The status of the images could be monitored using *Image Viewer*, and *Data Backup* generated backup files as needed. This process continued until the completion of live cell imaging.

**Staining of cells with SNAI-TRITC.**

Fluorescent imaging was conducted at the end of the live cell imaging process. Before removing the Lab-Tek™ II 8 Chamber Slide from the microscope stage, a snapshot was captured using Map software to document the position of a unique object, such as a cell with a distinct shape. Subsequently, the chamber slide was removed from the stage, and the cells underwent three washes with phosphate-buffered saline (PBS). Following this, the cells were fixed with 3.7% paraformaldehyde for 15 minutes at room temperature and then washed three additional times with PBS. Next, the cells were treated with Carbo-Free Blocking Solution (×1) (Vector) for 1 hour at room temperature and subsequently incubated with TRITC-labelled SNAI (E-Y Laboratories) diluted with Carbo-Free Blocking Solution (×1) to a concentration of 50 μg/ml for 1 hour at 4°C. Following this incubation, the cells were washed with PBS three times and exposed to DAPI. This was performed by diluting two drops of NucBlue Fixed Cell ReadyProbes Reagent with 1 ml Carbo-Free Blocking Solution (×1) for 15 minutes at room temperature, followed by three additional washes with PBS. The Lab-Tek™ II 8 Chamber Slide was then returned to the microscope stage, and its position was adjusted using the snapshot, which was taken before removing the chamber slide from the microscope stage. As returning it to the identical position as during live cell imaging proved to be challenging, the shift in the x and y position from the live cell imaging setup was calculated, and a new MDA file, accounting for this shift in position, was generated. Following the upload of this new MDA file, fluorescent image acquisition was conducted by Metamorph. Fluorescent imaging was conducted with a 250 ms exposure using lasers with wavelengths of 403 nm and 491 nm for DAPI and TRITC, respectively.

**Segmentation of NIR-DIC images.**

Grayscale live cell imaging videos were then subjected to image segmentation. To this end, we developed a method referred to as Stepwise Area Expansion, which was performed by *Outline Drawing* software (Supplementary Fig. 2). Four images were created from the original grayscale image by applying four different thresholds (TH) values (TH1–4, Supplementary Fig. 2A). First, pixels above the threshold value (e.g. 200, 170, 140, and 110 of 0–255 grayscale for TH1, 2, 3, and 4, respectively) were extracted from the grayscale image. Segmentation was then performed step by step, starting from the TH1 image (Supplementary Fig. 2B). Connectivity analysis was performed on the TH1 image to identify groups of pixels that were attached and an edge circle that surrounded the group of pixels was then determined (Supplementary Fig. 2B, Panel a, pink circles). The TH1 edge circles were overlaid on the corresponding grayscale image (Supplementary Fig. 2B, Panel b). The pixels composing the circles were referred to as the original pixels. Using the overlaid grayscale image as a reference, the software examined the pixel values for each original pixel. The values of pixels located in a circle from 12 o'clock from the original pixels were examined (Supplementary Fig. 2B, Panel b and Line extension (magnified)). The edges of the pixels were extended until the values became either 100 (representing the grayscale background) or 50 above or below the original pixel, as long as those extensions were towards the outside of the edge circles. The final pixel positions were determined by this technique. Similar examinations were performed for the other three directions (Supplementary Fig. 2B, Panel b: 3, 6, and 9 o'clock directions; yellow, light blue, and blue lines, respectively) and the final pixel positions were then linked to create four directional lines corresponding to the 12, 3, 6, and 9 o'clock directions (Supplementary Fig. 2B, Panel c and Edge linking (magnified), L 12', L3', L6', and L9'). Finally, the four lines were linked to make an edge circle (Supplementary Fig. 2B, Panel d, green circles), which completed the initial segmentation for the first TH1 image. The TH1 edge circles were then overlaid on the TH2 image and another connectivity analysis was performed, except for the TH1 edge circle areas (Supplementary Fig. 2B, Panel e, green circles). Two types of edge circles (Supplementary Fig. 2B, Panel e, green and pink circles) were thus created at this stage. The pink circles that emerged in the TH2 image were overlaid on the corresponding grayscale image, the pixel value of each original pixel in the pink circles was examined, the final pixel positions were determined (Supplementary Fig. 2B, Panel f), and the positions were linked to create four lines (Supplementary Fig. 2B, Panel g) and a new edge circle was created (Supplementary Fig. 2B, Panel h), as described for TH1. The software applied a different expansion approach for the green edge circles that had been determined during TH1 processing (Supplementary Fig. 2B, Panel e, green circles): the location of each pixel on the edge of the circles (Supplementary Fig. 2B, Panel k) was moved by 1–4 pixels outside the original (Supplementary Fig. 2B, Panel l, Edge expansion (magnified)) to create the red circles (Supplementary Fig. 2B, Panel l). Both the green TH2 (Supplementary Fig. 2B, Panel h) and red TH2 circles (Supplementary Fig. 2B, Panel

l) were then overlaid onto the TH2 image (Supplementary Fig. 2B, Panel i), and the circle overlaps were removed to create new edge circles (Supplementary Fig. 2B, Panel j). Those new edge circles were then applied to TH3, and the above-mentioned processes were repeated for TH3 and TH4 images. Areas surrounded by edge circles were numbered.

This approach proves valuable in scenarios where cell segmentation encounters challenges. For instance, when a flat cell coexists with larger, bright cells that are easily distinguishable in the NIR-DIC image, the flat cell may become obscured by the brightness of the larger cells, making segmentation difficult. In the Stepwise Area Expansion method, the initial step involves defining the boundaries of the brighter objects within the TH1 image. Since the TH1 image typically does not encompass the darker flat cell, there is a possibility that the flat cell lies outside this boundary. In subsequent steps utilizing images created with lower threshold levels, the darker flat cell can be included in the image. By excluding the area within the previously determined boundary during segmentation, the region corresponding to the flat cell can be segmented. This approach facilitates accurate segmentation of grayscale images that contain cells with varying brightness levels and shapes. Supplementary Fig. 2A (depicted by blue lines) provides examples of segmentation outcomes using both high and low-density HeLa cell cultures.

**Assignment of cell lineage number and cell number.**

To assign cell lineage numbers and cell identifiers to the cells within the Time 1 image, a verification and correction process was carried out for the segmentation results using the *Object Tracking Controller* software. In cases where multiple segments were incorrectly associated with a single cell, one segment was chosen to represent the cell, and segments were merged as needed. Conversely, if multiple cells were grouped within a single segment, the segment was divided to accurately represent each individual cell. The cells identified within the Time 1 image were classified as progenitor cells and a unique cell lineage number was assigned to each progenitor cell. Progenitor cell numbers were designated as 0. Data was then recorded in the Cell Lineage Database.

**Automated object tracking (single-cell tracking).**

After assigning cell lineage numbers to segmented areas, the automatic cell tracking process was initiated using the *Automatic Cell Tracking* software. Supplementary Fig. 3 provides an overview of the automatic tracking process for the segmented area, outlined by green lines. In Supplementary Fig. 3A (Time A), orange characters represent segmented areas identified as cell representations. The blue area with a white asterisk denotes the cell being tracked, while the yellow and magenta areas indicate segmented areas representing neighboring cells. In the subsequent time point (Supplementary Fig. 3B, Time A+1), the positions of cells and segmentation patterns changed. To perform single-cell tracking, it was essential to determine the segmented area corresponding to the cell being tracked. To achieve this, the blue area from Time A was overlaid onto the Time A+1 image (Supplementary Fig. 3C). However, the blue segmented area from Time A did not exist in Time A+1; instead, it overlapped with the light blue and white segmented areas. The *Automatic Cell Tracking* software couldn't ascertain which area corresponded to the tracked cell. To resolve this, the yellow and magenta areas were overlaid on the Time A+1 image (Supplementary Fig. 3D). This process revealed that the light blue area was related to the yellow area. Furthermore, the magenta area overlapped with the white area, but the larger portion of the white area overlapped with the blue area. Consequently, the *Automatic Cell*

*Tracking* software determined that the white area represented the cell being tracked (Supplementary Fig. 3E). This process was systematically repeated for all cells recorded in the database.

**Tracking data verification.**

Since achieving 100% accuracy in automatic single-cell tracking is not always feasible in practice, the tracking data underwent verification using the interactive features of the *Object Tracking Controller* software. If errors were identified, the single-cell tracking data were corrected, ultimately enabling the generation of nearly 100% accurate single-cell tracking data.

**Data analysis software.**

Bioinformatics analysis was conducted using the *Data Analysis* software, which encompasses a range of functions such as generating cell lineage maps, calculating the individual cell doubling time, and determining cell population expansion curves. Notably, cell fate simulation is one of the features offered by this software.

**Staining of cells with antibodies against a stem cell marker.**

HeLa and MiaPaCa2 cells were fixed with 3.7% paraformaldehyde for 15 minutes at room temperature, followed by three additional washes with PBS. Subsequently, the cells underwent treatment with Carbo-Free Blocking Solution (×1) (Vector) for 1 hour at room temperature. After this blocking step, the cells were incubated with the following primary antibodies or lectin: Anti-CD133 antibody (Developmental Studies Hybridoma Bank) at a dilution of 1:50, which corresponds to 5 μg/ml, in Carbo-Free Blocking Solution (×1) for 30 minutes at room temperature, Anti-TAG-72 antibody (B72.3) at a 1:50 dilution in Carbo-Free Blocking Solution (×1) for 1 hour at room temperature, or FITC-labelled rBC2LCN (Wako) at a 1:100 dilution in Carbo-Free Blocking Solution (×1) for 30 minutes at room temperature. Following the primary antibody or FITC-labelled rBC2LCN incubation, the cells were washed three times with PBS. For cells incubated with primary antibodies (anti-CD133 and anti-TAG-72), a secondary antibody (Invitrogen, goat anti-mouse IgG Alexa Fluor 488) diluted 1:1000 was applied for 1 hour at room temperature. After the secondary antibody incubation, the cells were subjected to three additional washes with PBS. Subsequently, the cells were stained with DAPI and underwent three final washes with PBS. Fluorescent imaging was conducted with a 250 ms exposure using lasers with wavelengths of 403 nm for DAPI and 491 nm for Alexa Fluor 488 and FITC.

**Generation of deduced cell populations with varied 2-6Sia expression levels.**

The initial step in generating the deduced cell population involved creating a 2-6Sia expression list (Supplementary Fig. 4A). In this list, we considered examples from Lineage 1 and Lineage 2. Indirect immunofluorescence using TRITC-tagged SNA1 was performed at the end of live cell imaging, allowing us to determine the SNA1 binding levels of cells present at the end of the imaging period. Lineage 1 exhibited no expression of 2-6Sia in any of the cells present at the end of live cell imaging, while Lineage 2 showed various levels of 2-6Sia expression in its cells. Based on the 2-6Sia expression levels of cells at the end of live cell imaging, we traced back the 2-6Sia expression levels along the cell lineage map to determine the expression levels of their parent cells. This was done by calculating the average expression of daughter cells, thus estimating the evolution of 2-6Sia expression levels within a cell lineage. Subsequently, we compiled a 2-6Sia expression list that included the 2-6Sia expression levels of all tracked HeLa or MiaPaCa2 cells. For the Lineages 1 and 2 examples, the list contained 17 data items with a value of 0 and 17 data items with varying levels of expression values.

Next, we generated the deduced cell population using the cell fate simulation algorithm (*1*), as illustrated in Supplementary Fig. 4B. This algorithm analyzed cell lineage data and calculated the probability of certain types of cell events occurring following a given event, along with the time intervals between events. The combinations of events that the algorithm analyzed were previously described (*1*). The cell fate simulation algorithm employed these probabilistic values to generate a deduced cell population. Initially, the algorithm assigned the length of time to the First event (the event that occurred in the progenitor cell) to the progenitors. In the actual process, 500 progenitors were generated, each at a different stage of the cell cycle, resulting in varying time intervals until the First event.

Subsequently, a cellular event was assigned to a progenitor cell based on the probabilistic values. If a bipolar cell division was assigned, two daughter cells were created. In Example 1, a longer time interval until the next event was assigned compared to Example 2. After the assignment, the algorithm referred to the cell doubling time (the time between bipolar cell divisions) of 2-6Sia-expressing cells. As 80% of the cell doubling times for 2-6Sia-expressing cells fell within ±10% of the average cell doubling time for these cells, the algorithm checked whether any of the 2-6Sia-expressing cells had cell doubling times within ±10% of the time assigned to daughter cells. If no such cells were found, 2-6Sia expression levels were not assigned to the daughter cells (Example 1). However, if some cells met this criterion, 2-6Sia expression levels were assigned to the daughter cells (Example 2). This approach allowed us to generate deduced cells with reproductive abilities similar to those of 2-6Sia-expressing cells.

Subsequently, the algorithm assigned 2-6Sia expression levels to the daughter cells by referring to the 2-6Sia expression level list (Supplementary Fig. 4C). A random value was generated to select one value from the list, which was then assigned to one of the daughter cells. In the case of the Lineage 1 and 2 examples (Supplementary Fig. 4A), there was a 50% chance of selecting value 0. As the variation in 2-6Sia expression levels typically fell within ±15%, the algorithm determined the 2-6Sia expression level of the second daughter cell based on the first daughter cell's level, accounting for the ±15% variation. Finally, 2-6Sia expression levels were modified to create deduced cell populations with various levels of 2-6Sia expression. This was achieved by multiplying the value of the 2-6Sia expression by factors such as 1.5, 0.5, and 0.25. In cases where the calculated value exceeded the highest 2-6Sia expression value observed in the HeLa or MiaPaCa2 cell population, the value was capped at the highest observed level.

**The 3D TME simulation methodology.**

The outline of the 3D TME simulation was presented in Supplementary Persuade Code 1. The codes for the 3D TME were developed using C, C++, and Objective-C with the utilization of SceneKit (macOS 12). To execute the 3D TME simulation, various parameters pertaining to graphic display and simulation execution were configured. Graphic display parameters for both cancer cells and immune cells included options such as cell shape (sphere, box, or capsule), displayed cell size, color, alpha value, motility of cells, nucleus size, and nucleus color. These parameters could be individually adjusted for cancer cells, Suppressive cells, Permissive cells, and Lethal cells. Additionally, for fluorescence display reflecting the quantitated fluorescent value of each cancer cell, the level of expression could be visualized using either a heatmap scale (ranging from blue to red, indicating low to high expression) or manually set colors.

Concerning Suppressive, Permissive, and Lethal cells, the ratio of each cell type relative to cancer cells can be specified. In a typical simulation, 500 progenitors of the deduced cell population were employed, and the numbers of Suppressive, Permissive, and Lethal cells were determined based on the specified ratio relative to the total of 500 cells. For instance, if the ratio was set to 1:0.5 (cancer cells to either Suppressive, Permissive, or Lethal cells), then 250 of the chosen cell types were placed accordingly. These ratios could be customized independently for each category of cells, and the range of the ratio as well as the increment could be set. For instance, the 3D TME simulation could be automatically conducted by accounting for the defined range and increment. As an example, if the range was set from 1:0 to 1:1 for each category with an increment of 0.1, this would result in 1331 simulations being carried out automatically.

The strength of the impact could also be configured. In the case of Suppressive and Permissive cells, the setting values, such as 10, represented the percentage for prolonging or shortening the cell doubling time. For example, if a deduced cell had a doubling time of 25 hours and encountered a Suppressive cell, the doubling time would be extended by 10%, making it 27.5 hours. In the case of encountering Permissive cells, the cell doubling time for deduced cells and their progeny would be reduced by 10%. However, this reduced cell doubling time could not fall below the shortest cell doubling time observed within the deduced cell population, and in such cases, the shortest time was assigned.

As for Lethal cells, the setting value represented the probability of inducing cell death. For instance, a setting of 100 indicated a 100% chance of inducing cell death when cancer cells encountered a Lethal cell. When cell death was triggered for a cancer cell, all its progeny were removed from the cell lineage database. The range of these setting values, along with the increment, could be customized, and the 3D TME simulation could be automatically executed, considering the specified increment value. The resistance of cancer cells to Suppressive and Lethal effects could also be set. For example, if the maximum resistance level was defined as 5-fold, then cancer cells subjected to Suppressive or Lethal effects would have a 20% chance (1 in 5 chance) of experiencing these effects.

These cells were positioned within spheres, each with a radius of 50 pixels. Cells were randomly distributed within the space, typically with a radius of 6 pixels from the cancer cell to search for Suppressive, Permissive, and Lethal cells. Under this setting, neighboring cancer cells were found within an 80% chance within the 6-pixel radius. Multiple cell lineage data of the deduced cell population can be loaded and the primary simulation process commenced with cancer cells identified at Time 1 and involved searching within a specified radius. If Suppressive, Permissive, and/or Lethal cells were discovered, the two nearest cells were selected. If the nearest cell was a Lethal cell, the resistance level of a cancer cell was initially considered, followed by the strength factor. If a cancer cell was determined to be killed, the cell death process was executed. If the nearest cell was a Suppressive cell, and the second nearest cell was not a Permissive cell, the resistance level of a cancer cell was initially taken into account, followed by the strength factor. If the cancer cell doubling time was prolonged, the process for prolongation was executed. If the nearest cell was a Permissive cell, and the second nearest cell was not a Suppressive cell, the strength factor was taken into account. If the cancer cell doubling time was determined to be reduced, the process for reduction was executed. If the nearest cell was a Suppressive cell, and the second nearest cell was a Permissive cell, the distances of both cells from a cancer cell were calculated, and the Suppressive strength was adjusted, considering the presence of Permissive cells. For instance, if the initial strength of both Suppressive and

Permissive cells was set to 50 and the distance of Suppressive and Permissive cells from a cancer cell was 3 and 6 pixels, respectively, the effect of Permissive strength was considered as half of the Suppressive strength, resulting in an adjusted Suppressive strength of 25 (50 (Suppressive) - 25 (Permissive)). If the nearest cell was a Permissive cell, and the second nearest cell was a Suppressive cell, a similar adjustment process was applied. These processes were iteratively carried out for each cancer cell identified at each time point and continued until reaching the final time point.

**Parameters for the Methodology of 3D TME Simulation.**

In the 3D TME simulation, the motility of cancer cells, Suppressive cells, Permissive cells, and Lethal cells was restricted to a maximum displacement of 2 pixels from their initial positions within the 3D TME sphere. This constraint was crucial to confine cancer cells and immune cells within the simulation sphere. However, this constraint also fixed the relative positions of cancer cells concerning Suppressive, Permissive, and Lethal cells. For example, once a cancer cell was positioned within the 3D TME, the spatial relationship between that cancer cell and nearby Suppressive, Permissive, or Lethal cells remained constant throughout the cell's simulation. As a result, if Suppressive, Permissive, and/or Lethal cells located within 6 pixels of a cancer cell were monitored at each time point, the same combination of Suppressive, Permissive, and/or Lethal cells would persist. This implied that if a Suppressive cell was initially the closest to a cancer cell, it would always remain the closest cell throughout the cancer cell's lifetime.

During the simulation, when a cancer cell experienced a Suppressive or Permissive effect, it did not receive additional Suppressive or Permissive effects. In other words, if a cell encountered a Suppressive cell and had the effect induced, it would not undergo another Suppressive effect. This rule applied uniformly to all progeny cells. However, these progeny cells could subsequently encounter additional Suppressive, Permissive, or Lethal effects.

When cancer cells produced daughter cells, the locations of these daughter cells within the 3D space were randomly determined within 6 pixels from the parent cancer cell. This randomization altered the relative positions of Suppressive, Permissive, and Lethal cells concerning the daughter cells compared to their parent cell. Consequently, if a Suppressive effect was applied to a daughter cell due to its parent cell's previous interaction with a Suppressive cell, and the daughter cell encountered a Permissive cell, the daughter cell would be subject to the Permissive effect. Similarly, daughter cells could receive a Suppressive or Lethal effect if either of these cells became the closest to a daughter cell.

**Other Parameters for 3D TME Simulation Methodology.**

Upon placing Suppressive, Permissive, and Lethal cells in the 3D TME, individual cell lineages were established, each characterized by its specific attribute (e.g., suppressing, permissive, or lethal). The 3D TME simulation algorithm executed simulation processes by referencing these attributes. This flexibility allowed the creation of comprehensive simulation scenarios by modifying attributes, including alterations at specific stages of the simulation to reflect changes in cell behavior. \

Each cancer cell was associated with attributes, such as 2-6 Sia expression levels, and by incorporating multiple attributes, the simulation could encompass various characteristics of cancer cells. Thus, the 3D TME simulation, based on lineage data and associated attributes for each cell, facilitated the execution of complex simulations, provided that the biological implications of each cell's attributes were defined.

**Statistical analysis.**

Statistical analyses were conducted using Prism 10 software.
