## Supplementary Text for Pseudocode 1 for "Cell Fate Simulation Reveals Cancer Cell Features in the Tumor Microenvironment"

### Supplementary Text for Pseudo Code 1

To perform the 3D TME simulation, several parameters needed to be defined. These parameters included: [1] Resistance levels of 2-6Sia-expressing Cells: Determining the resistance levels of cells expressing 2-6Sia to suppressive and lethal effects, [2] Cell ratios: Establishing the relative numbers of Suppressive, Permissive, and Lethal cells in relation to cancer cells, and [3] Impact strength: The strength of the influence exerted by Suppressive, Permissive, and Lethal cells on cancer cells.

As for [1], cells expressing 2-6Sia demonstrate approximately 2 to 4.5-fold resistance to TNF, cisplatin, and gemcitabine compared to non-expressing cells (*1-3*). To take into account this, we assigned a 5-fold resistance factor to cancer cells expressing the highest level of 2-6Sia when subjected to Suppressive or Lethal effects, relative to non-expressing cells. In this scenario, when a cancer cell encounters, for instance, a Lethal cell, there is a 20% chance of the cancer cell being eliminated (a 1 in 5 chance). Conversely, if a cancer cell lacks 2-6Sia expression and encounters a Lethal cell, it is eliminated with a 100% probability. The degree of resistance for other cells was calculated based on their 2-6Sia expression levels.

Regarding parameter [2], we conducted a 3D TME simulation involving cancer cells and either Suppressive, Permissive, or Lethal cells to determine the ratio that could effectively observe the impact of these cells on cancer cells (Supplementary Fig. 5A-H, using Cervical 2-6Sia 1.0 and Pancreatic 2-6Sia 1.0 cells). Within the ratios of deduced cancer cells to Suppressive cells, ranging from 1:0.1 to 1:0.9 (Supplementary Fig. 5A and E) and the ratios of cancer cells to Lethal cells, ranging from 1:0.1 to 1:0.46 (Supplementary Fig. 5C and G), we observed a decrease in the expansion of the cancer cell population, as expected. This reduction was primarily due to the prolongation of cell doubling time or cell elimination, with Lethal cells exerting a stronger effect compared to Suppressive cells. However, when we performed the simulation by varying the ratio of Permissive cells, it did not have a significant impact on the expansion of the cell population. This observation can be attributed to the fact that the minimal cell doubling time was restricted from falling below the shortest doubling time observed within the deduced cell population (Supplementary Fig. 5B and F). Deduced cancer cells, which had a doubling time close to the shortest observed, may have experienced a comparatively lower influence from the presence of Permissive cells. Consequently, we conducted the 3D TME simulation with a fixed ratio of 1:0.2 for deduced cancer cells to Suppressive cells and a varying ratio of Permissive cells (cancer cells: Suppressive cells: Permissive cells; 1:0.2:0.2 to 1:0.2:0.5, Supplementary Fig. 5D and H). This approach allowed the effects of Permissive cells to counteract the prolongation of cell doubling time caused by Suppressive cells, resulting in the reduction of cell doubling time. As anticipated, we observed that Permissive cells effectively mitigated the reduction in population expansion induced by Suppressive cells. These results indicate that the effects of Suppressive, Permissive, and Lethal cells can be observed within the ratio range of 1:0.1 to 1:0.9 for Suppressive and Permissive cells and of 1:0.1 to 1:0.46 for Lethal cells.

Regarding parameter [3], we performed a 3D TME simulation to determine the range of strength for Suppressive, Permissive, and Lethal cells' effects on cancer cells (Supplementary Fig. 5I and P). For Suppressive cells, when the strength was set to 10, the cell doubling time of the cells encountering the Suppressive cell and its progeny was extended by 10% (for instance, if the doubling time was 25 hours, it was extended to 27.5 hours). Similarly, with a strength setting of 10 for Permissive cells, the cell doubling time of the encountering cell and its progeny was reduced by 10%. If the reduced doubling time fell below the shortest doubling time within the deduced cell population, it was capped at the shortest time. For Lethal cells, the chance of inducing cell death was set. For example, setting values of 20 implied that cell death occurred with a 20% chance when cancer cells encountered a Lethal cell. We conducted 3D TME simulations by varying these setting values and found a similar trend for Suppressive, Permissive, Lethal, and the combination of Suppressive and Permissive cells, as analyzed by cancer cell to immune cell ratio (Supplementary Fig. 5I-P). These results suggest that the effects of Suppressive, Permissive, and Lethal cells can be analyzed within the range of 10% to 100% for Suppressive and Permissive cells and a 10% to 50% chance for Lethal cells.

The 3D TME simulation process involved the identification of Suppressive, Permissive, and Lethal cells within a specified spatial range around a cancer cell. At each time point, Suppressive, Permissive, and/or Lethal cells within this spatial range were located, and the nearest cell to the cancer cell was selected. If the selected cell was a Lethal cell, first, the resistance levels related to the 2-6Sia expression on cancer cells were considered, followed by the Lethal strength. If cell death was determined to be induced, the process of eliminating the cancer cell and its progeny was executed. If the nearest cell was a Suppressive cell, the cell doubling time was extended, after considering 2-6Sia expression levels followed by the Suppressive strength. When the nearest cell was a Permissive cell, the cell doubling time was reduced, accounting for the Permissive strength. In cases where the nearest and second nearest cells were a combination of Suppressive and Permissive cells, or vice versa, the relative distance between both cells was factored into the calculation of strength. This process was applied to each deduced cancer cell, and the iterative simulation continued until reaching the final time point. In Fig. 5B, a graphical representation of the 3D TME is presented, depicting views at 0, 2000, 4000, and 6000 minutes. The simulation was performed with Cervical 2-6Sia 1.5 cells in the presence of Suppressive, Permissive, and Lethal cells, with the influence strength of each category set to 0 (Control) or Lethal strength 100.
