## Supplementary Text for Supplementary Fig 4 for "Cell Fate Simulation Reveals Cancer Cell Features in the Tumor Microenvironment"

### (Deduced cell population generation)

The initial step involves creating a 2-6Sia expression list (**A** and **B**), which encompasses the expression levels of all analyzed HeLa or MiaPaCa2 cells obtained through single-cell tracking. This list serves as a reference when assigning 2-6Sia expression levels to individual cells. Subsequently, the cell fate simulation algorithm was used to generate deduced cell populations. The first step within the generation is to determine the length of time progenitor cells spend before experiencing their first event (**A** and **C**). Since each cell is in a different stage of the cell cycle, this length varies. The algorithm then assigns a cellular event to the progenitor cell. In the case of bipolar cell division, the algorithm generates two daughter cells. Next, 2-6Sia expression levels are assigned to each daughter cell using the 2-6Sia expression list (**A** and **D**). This is achieved by generating a random value to select one of the expression values stored in the list. For HeLa cells, where only a subset expresses 2-6Sia, the majority of values in the list are expected to be zero, resulting in a deduced cell population that mirrors HeLa cells' 2-6Sia expression pattern. Similarly, a deduced cell population resembling the 2-6Sia expression pattern of MiaPaCa2 cells can be generated. By assigning 2-6Sia expression to deduced cells with similar cell doubling time as HeLa or MiaPaCa2 cells, the 2-6Sia expression can be assigned to cells in a manner that reflects their reproductive capacity. Finally, 2-6Sia expression levels were modified by factors of 1.5, 1.0, 0.5, and 0.25. Although the expression levels may exceed the highest 2-6Sia expression levels of HeLa or MiaPaCa2 cells, they were capped at the highest observed value. We started with 500 progenitor cells resulting in the creation of 500 cell lineages per deduced cell populations.
