## Supplementary figures and images for "Cell Fate Simulation Reveals Cancer Cell Features in the Tumor Microenvironment"

### Supplementary Fig. 1

##### Sup Fig. 1

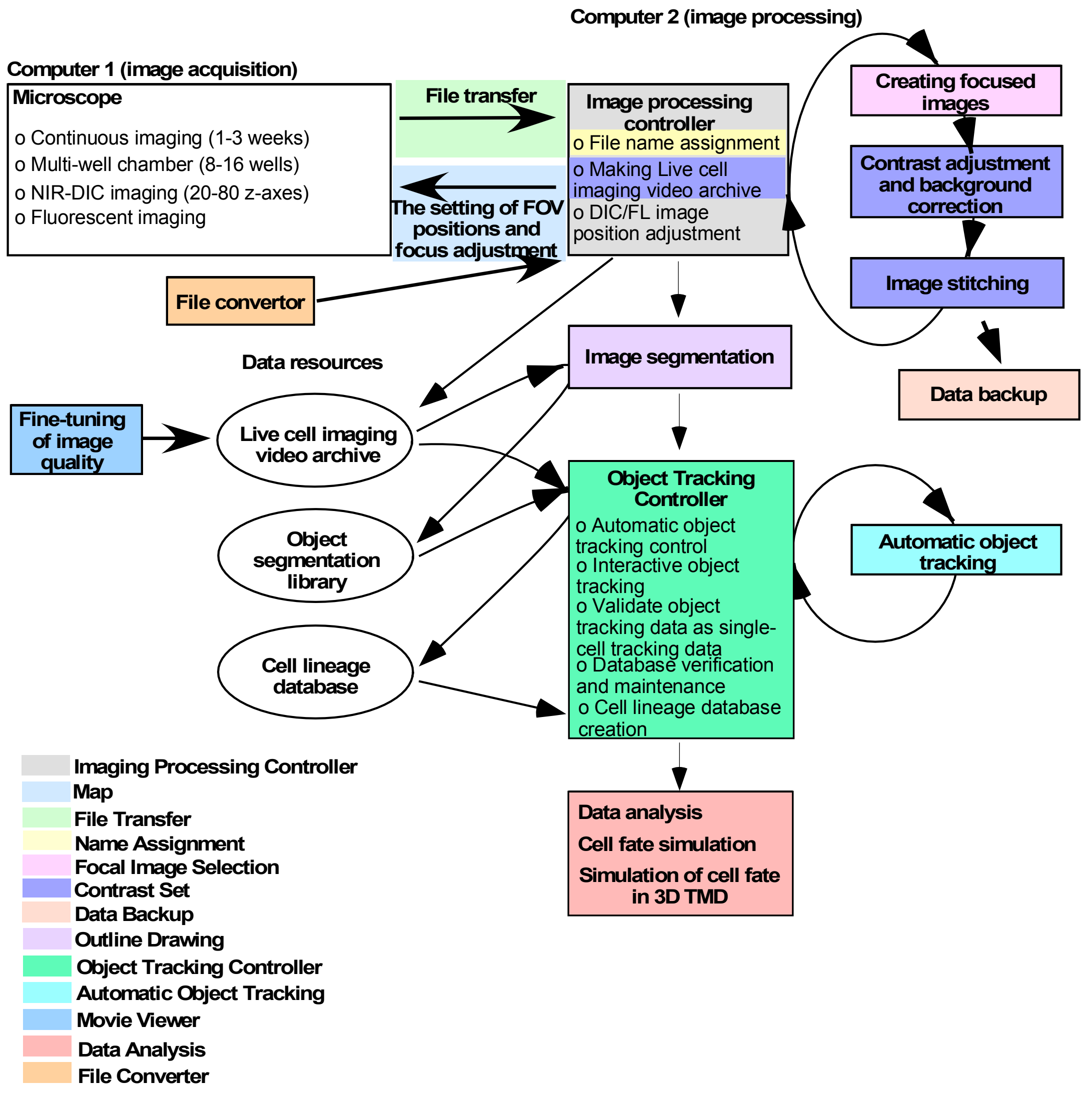


Sato et al.

### Supplementary Fig. 2

##### Sup Fig. 2

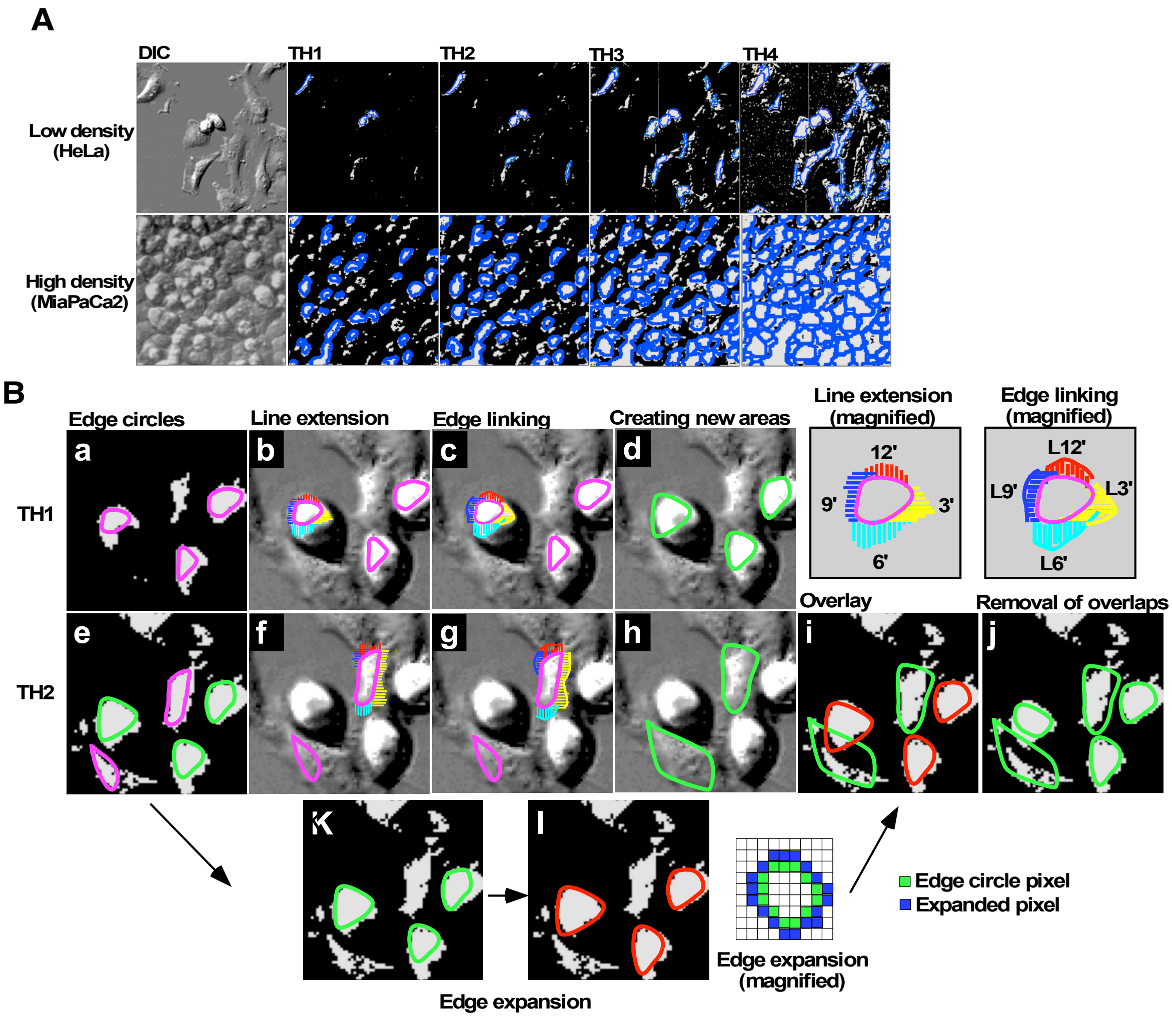


Sato et al.

### Supplementary Fig. 4

#### Sup Fig. 4

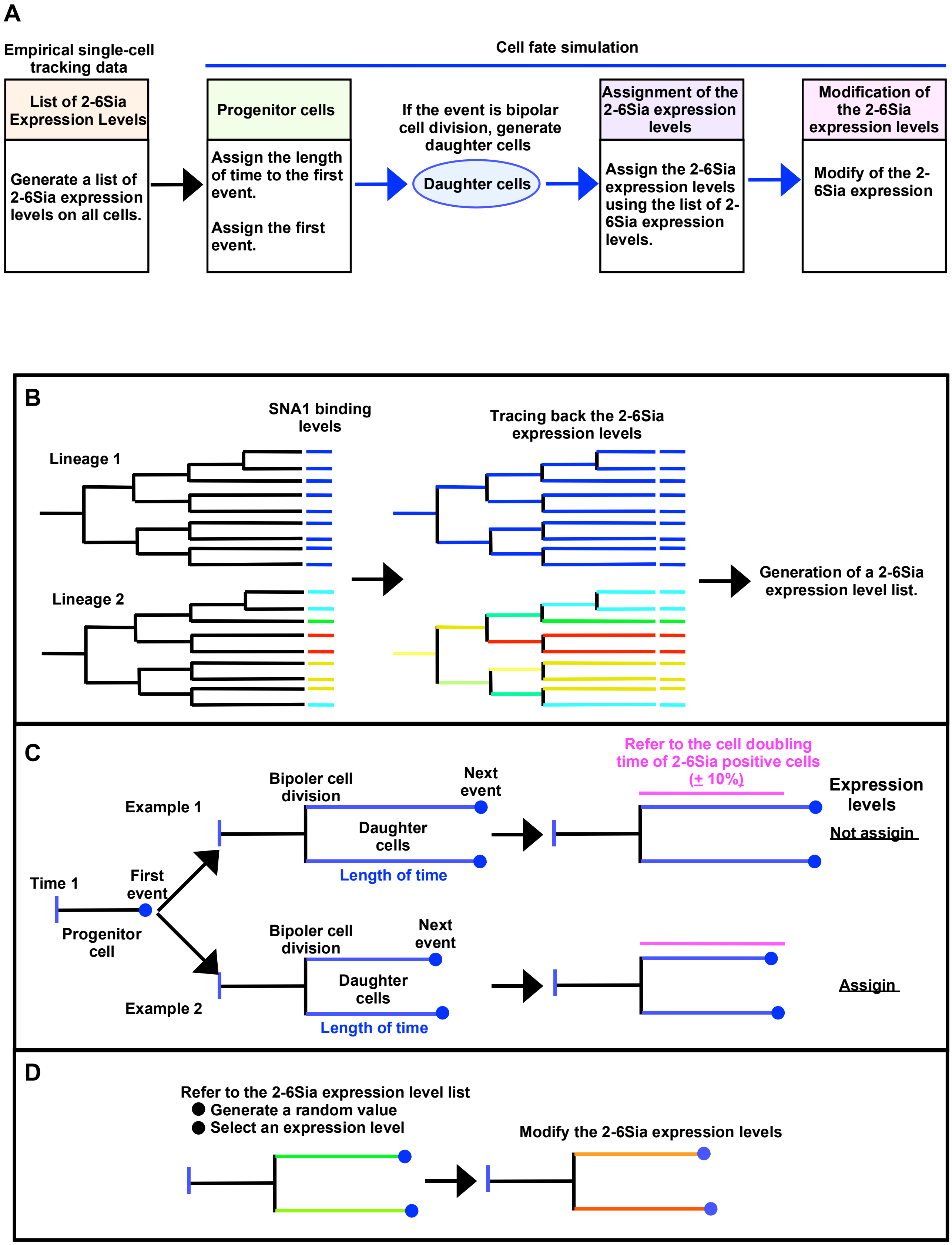


Sato et al.

### Supplementary Fig. 5

#### Sup Fig. 5

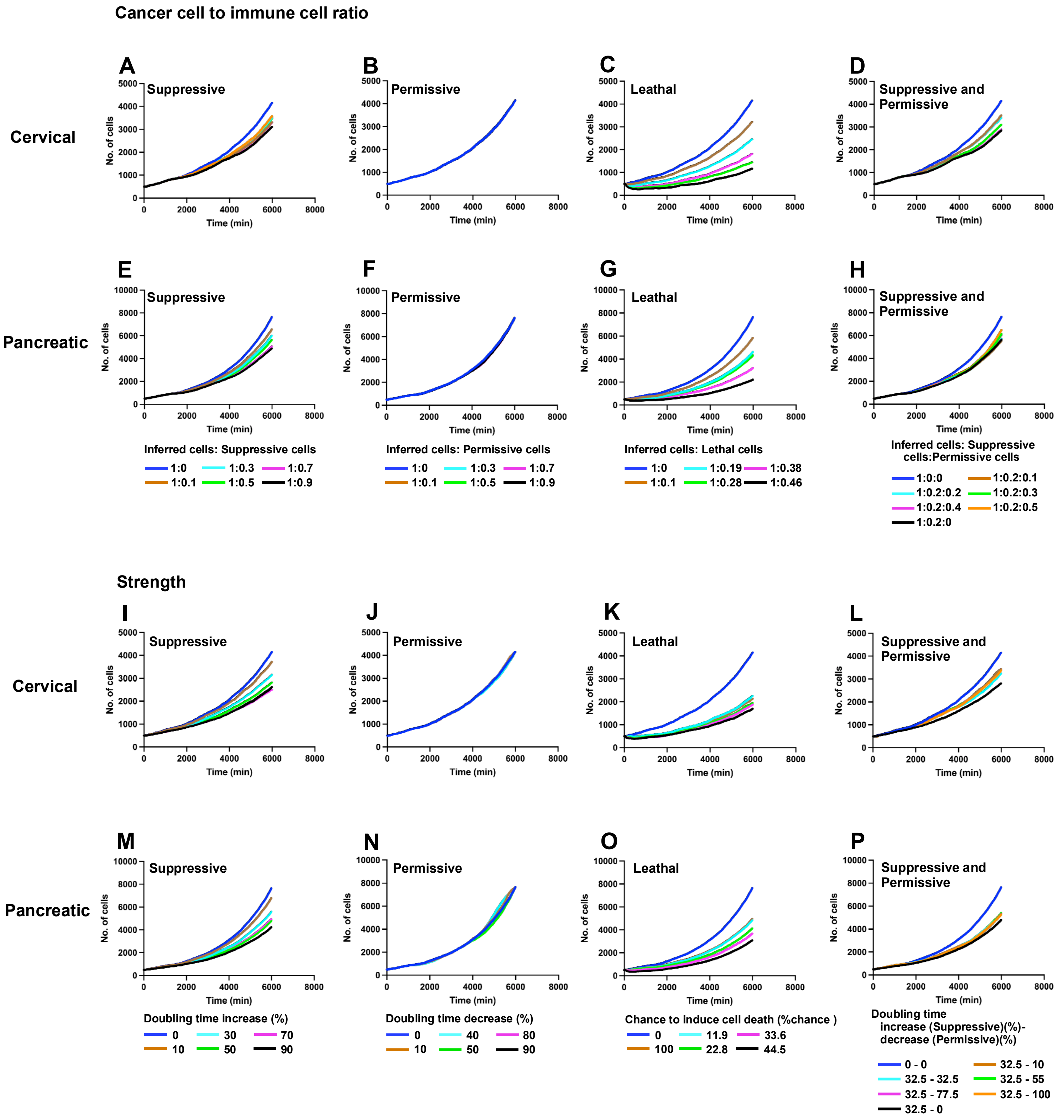


Sato et al.

### Supplementary Fig. 6

#### Sup Fig. 6

Cervical cancer

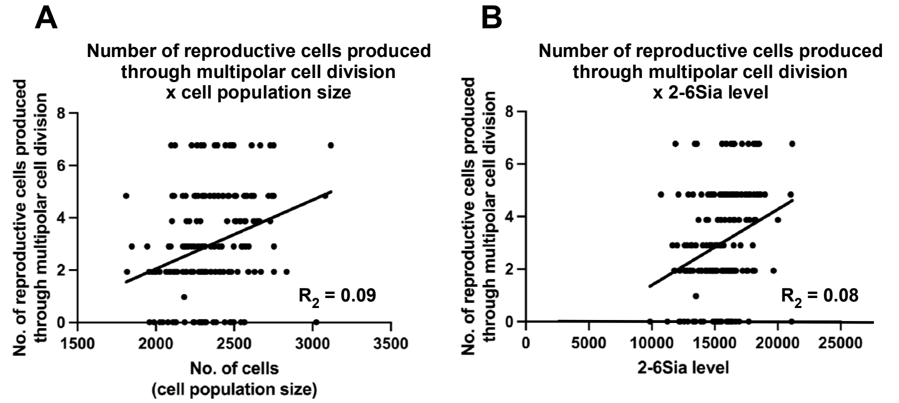


Pancreatic cancer

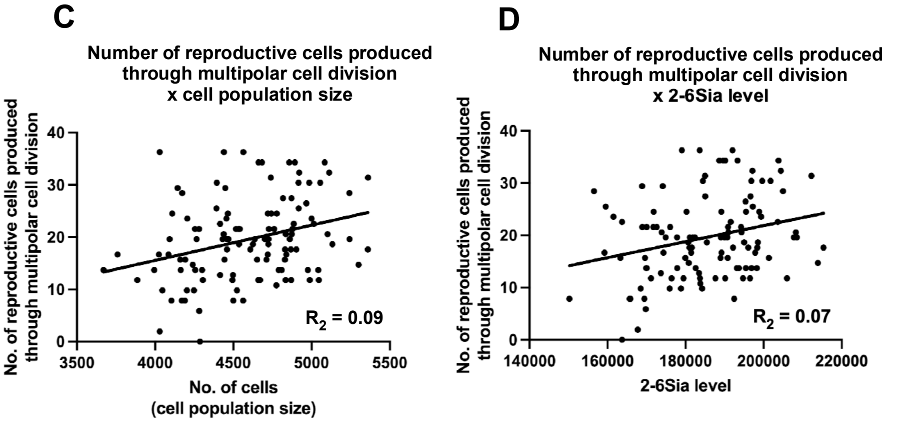


Sato et al.
