## Supplementary Fig. 3 for "Cell Fate Simulation Reveals Cancer Cell Features in the Tumor Microenvironment"

### Sup Fig. 3

Time A

Time A+1

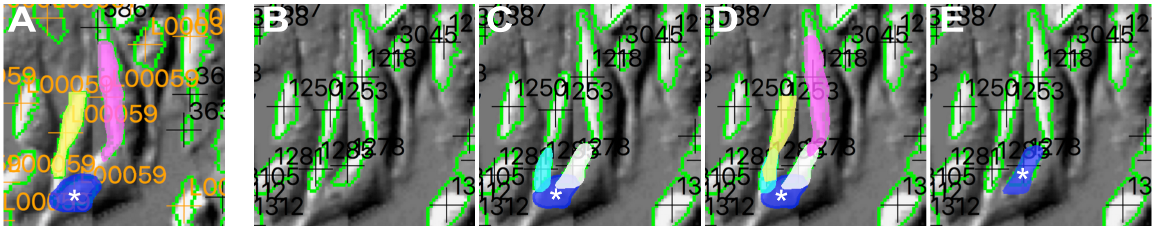


A: The cell under tracking ( blue with a white asterisk) and neighboring cells (yellow and magenta)

B: Time A+1 with segmented areas

C: Nearby segmented areas (light blue and white)

D: Adjacent segmented areas (light blue and white) and neighboring cells (yellow and magenta)

E: The segmented area that represents the cell being tracked.
