## Supplementary Pseudocode 1 for "Cell Fate Simulation Reveals Cancer Cell Features in the Tumor Microenvironment"

Setting simulation parameters:
  Display and setting parameters (cancer cells):
    * Cell Shape
    * Size
    * Motility
    * Alpha value
    * Fluorescent display mode (heat map or manual color selection):
    * Nucleus size
    * Nucleus color
  Display and setting parameters (TME cells):
    * Cell Shape
    * Size
    * Motility
    * Alpha value
    * Nucleus size
    * Nucleus color
    *Color for Suppressive, Permissive, and Lethal cells.
    * Ratio of cells to place for Suppressive, Permissive, and Lethal cells relative to cancer Cells; Each ratio can vary.
    * Impact strength of Suppressive, Permissive, and Lethal cells on cancer cells; Each value can vary.
    * Cancer cell resistance to suppressive and lethal cells based on stemness.
General parameters:
  * Diameter of spheres
  * Search distance

Place TME cells in the sphere.
Place cancer cells in the sphere.

For x = simulation time:
  * Calculate the distance of a cancer cell to Suppressive, Permissive, and Lethal cells in 3D TME.
  * Select the two nearest TME cells within the search distance.
  * Identify the type of cells: Suppressive, Permissive, and Lethal cells.

  * If a Lethal cell is the nearest cell:
    * Calculate the resistance of cancer cells to the Lethal effect (based on the stemness level)
    * Determine whether the cell death procedure is executed
    * If Yes, obtain the Lethal strength
    * Determine whether the cell death procedure is executed
    * If Yes, execute the cell death procedure

  * If a Suppressive cell is the nearest and the second nearest cell is not a Permissive cell:
    * Calculate the resistance of cancer cells to the Suppressive effect (based on the stemness

level)
* Determine whether the cell doubling time prolongation procedure is executed
* If Yes, obtain the Suppressive strength
* Determine whether the cell doubling time prolongation procedure is executed
* If Yes, execute the prolongation procedure

* If a Permissive cell is the nearest and the second nearest cell is not a Suppressive cell:
  * Determine whether the cell doubling time shortening procedure is executed
  * If Yes, execute the shortening

* If a Suppressive cell is the nearest and the second nearest cell is a Permissive cell or vice versa:
  * Obtain the Suppressive strength and Permissive Strength
  * Calculate the distance from a cancer cell to Suppressive and Permissive cells
  * Adjust Suppressive or Permissive strength
  * Execute the prolongation or the shortening procedure

Repeat

Export results
Display results in 3D

End
