## Supplementary Data 1 for "Cell Fate Simulation Reveals Cancer Cell Features in the Tumor Microenvironment"

Analysis: HeLa, Treat.: HeLa, Cell: HeLa

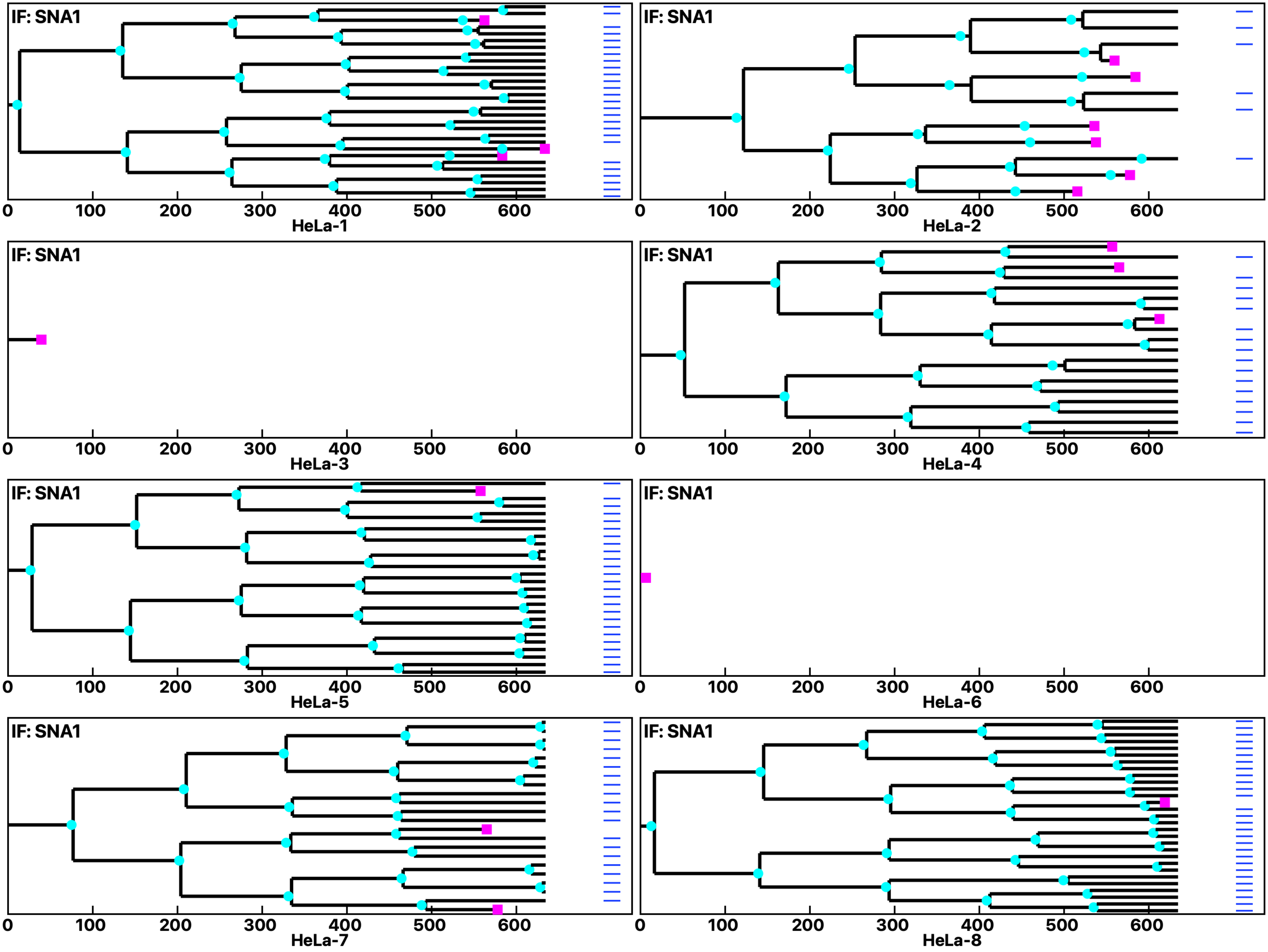

Analysis: HeLa, Treat.: HeLa, Cell: HeLa

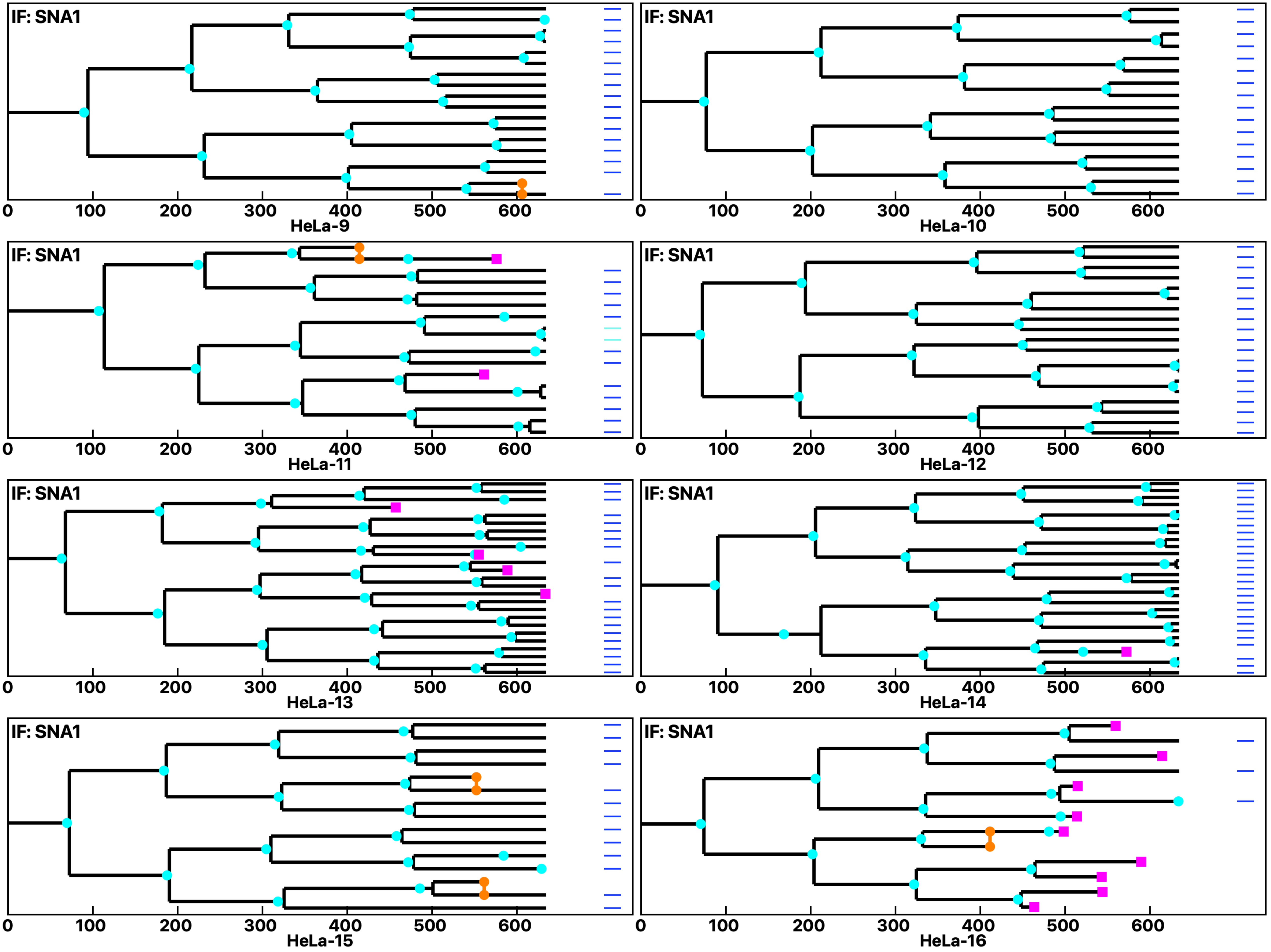

Analysis: HeLa, Treat.: HeLa, Cell: HeLa

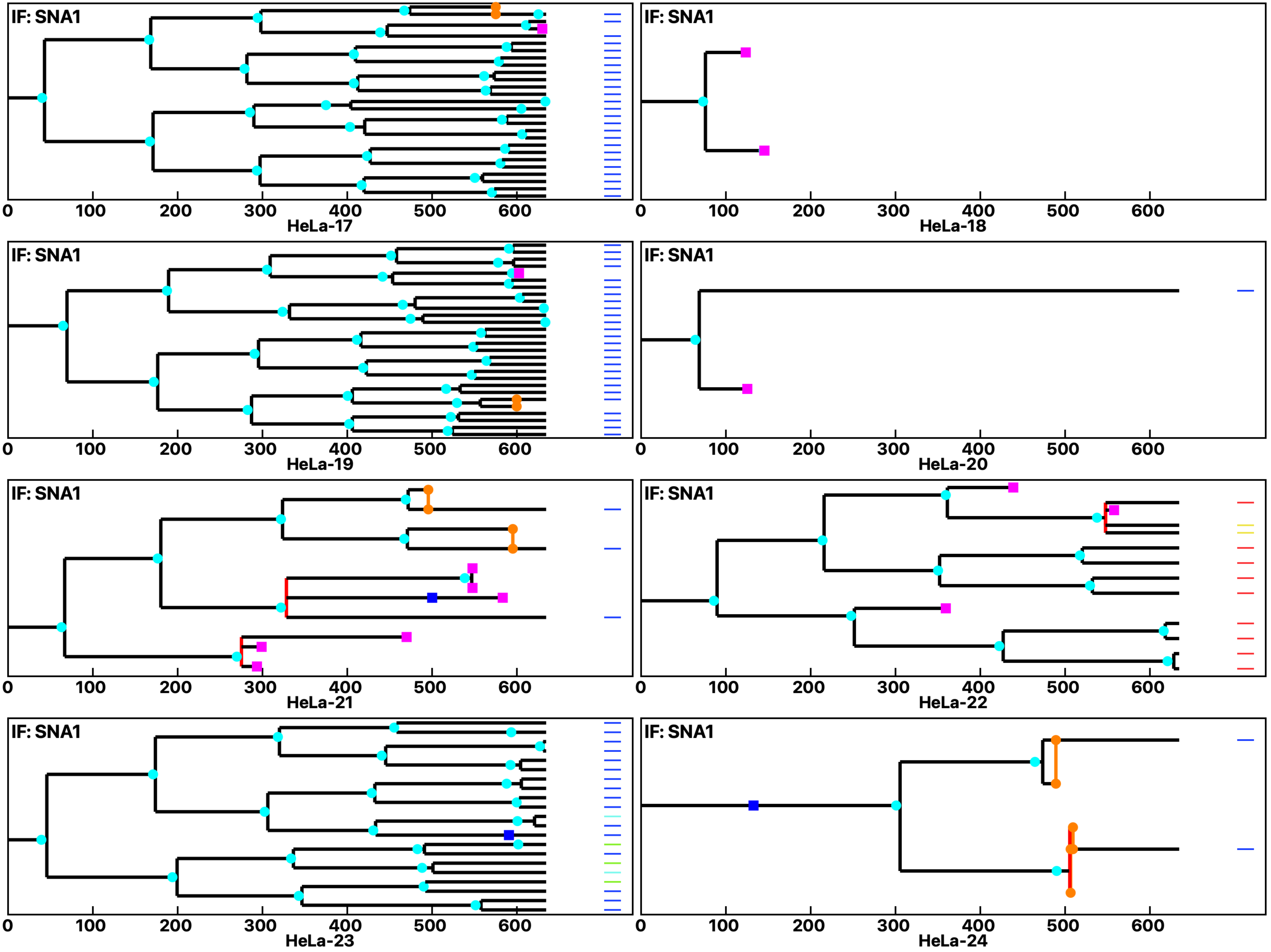

Analysis: HeLa, Treat.: HeLa, Cell: HeLa

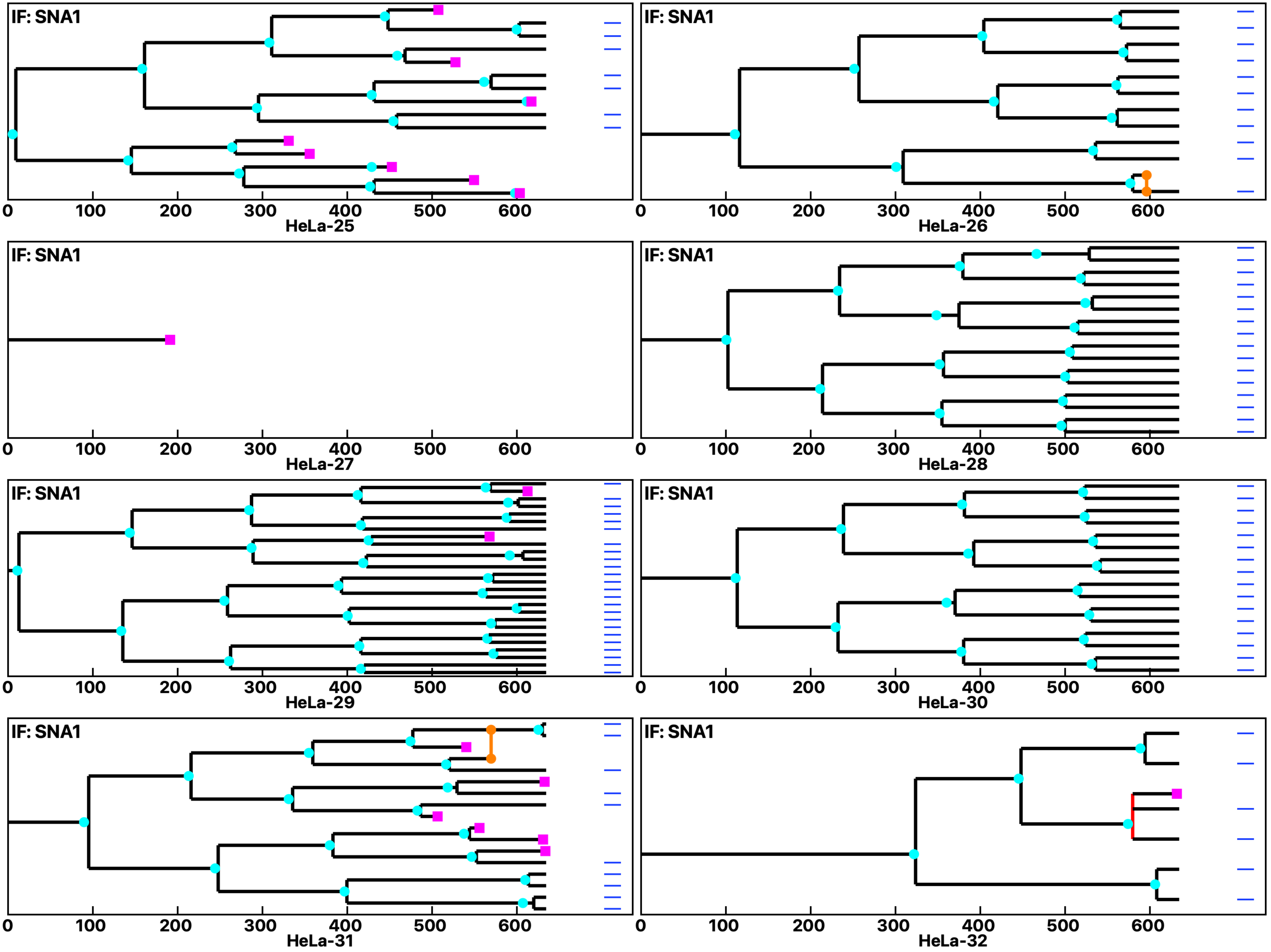

**Analysis: HeLa, Treat.: HeLa, Cell: HeLa**

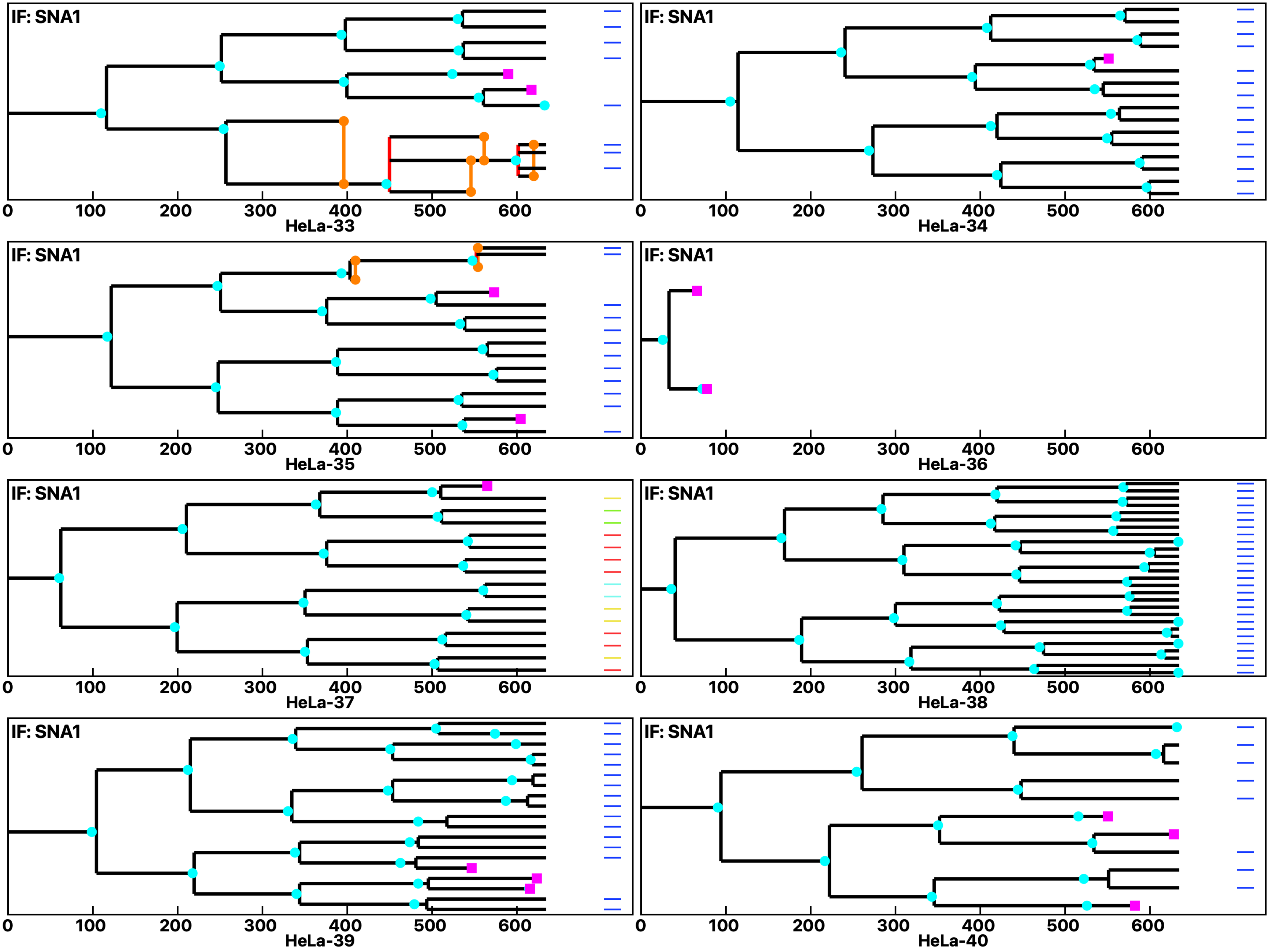

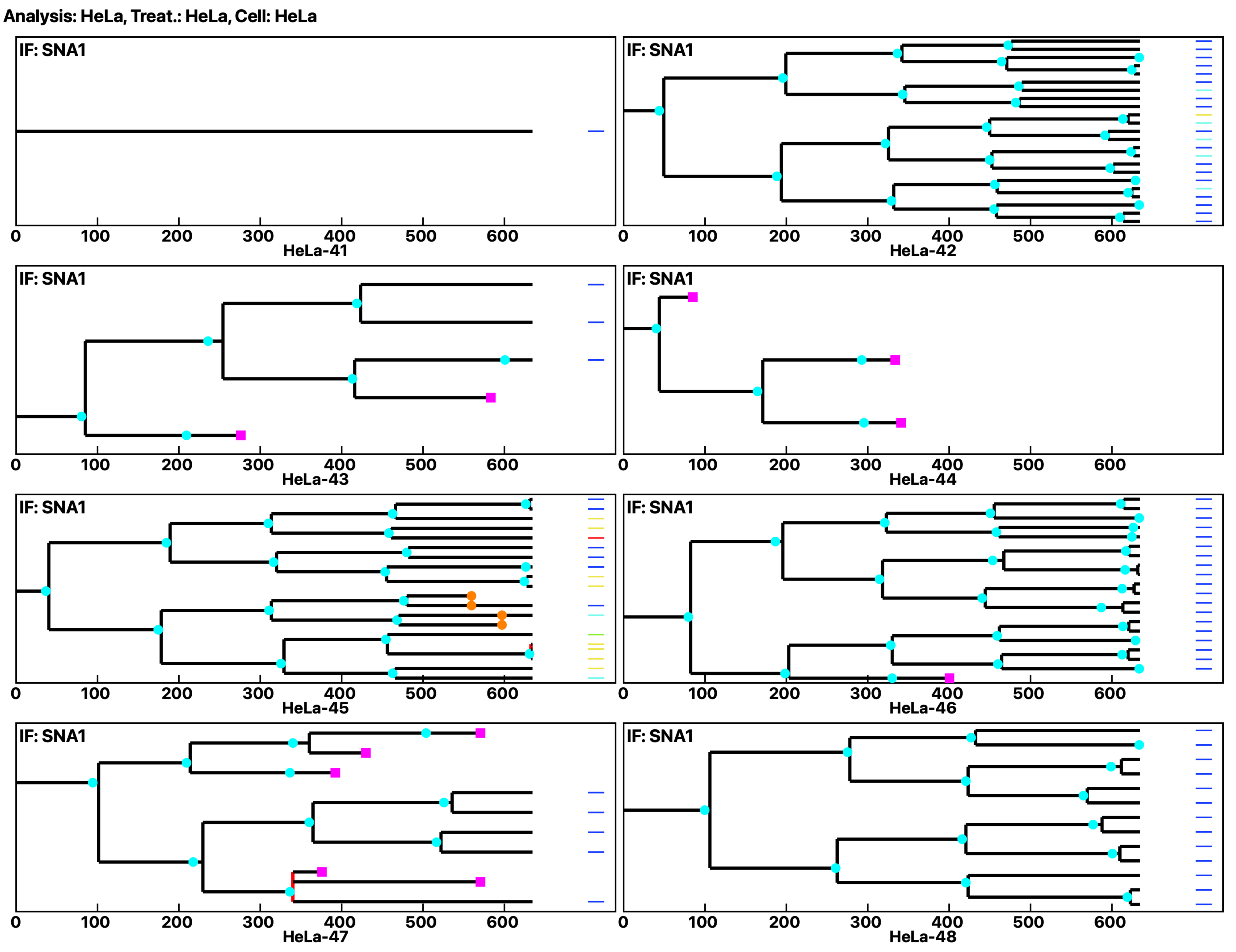
Analysis: HeLa, Treat.: HeLa, Cell: HeLa
IF: SNA1
0 100 200 300 400 500 600
HeLa-41
IF: SNA1
0 100 200 300 400 500 600
HeLa-42
IF: SNA1
0 100 200 300 400 500 600
HeLa-43
IF: SNA1
0 100 200 300 400 500 600
HeLa-44
IF: SNA1
0 100 200 300 400 500 600
HeLa-45
IF: SNA1
0 100 200 300 400 500 600
HeLa-46
IF: SNA1
0 100 200 300 400 500 600
HeLa-47
IF: SNA1
0 100 200 300 400 500 600
HeLa-48

Analysis: HeLa, Treat.: HeLa, Cell: HeLa

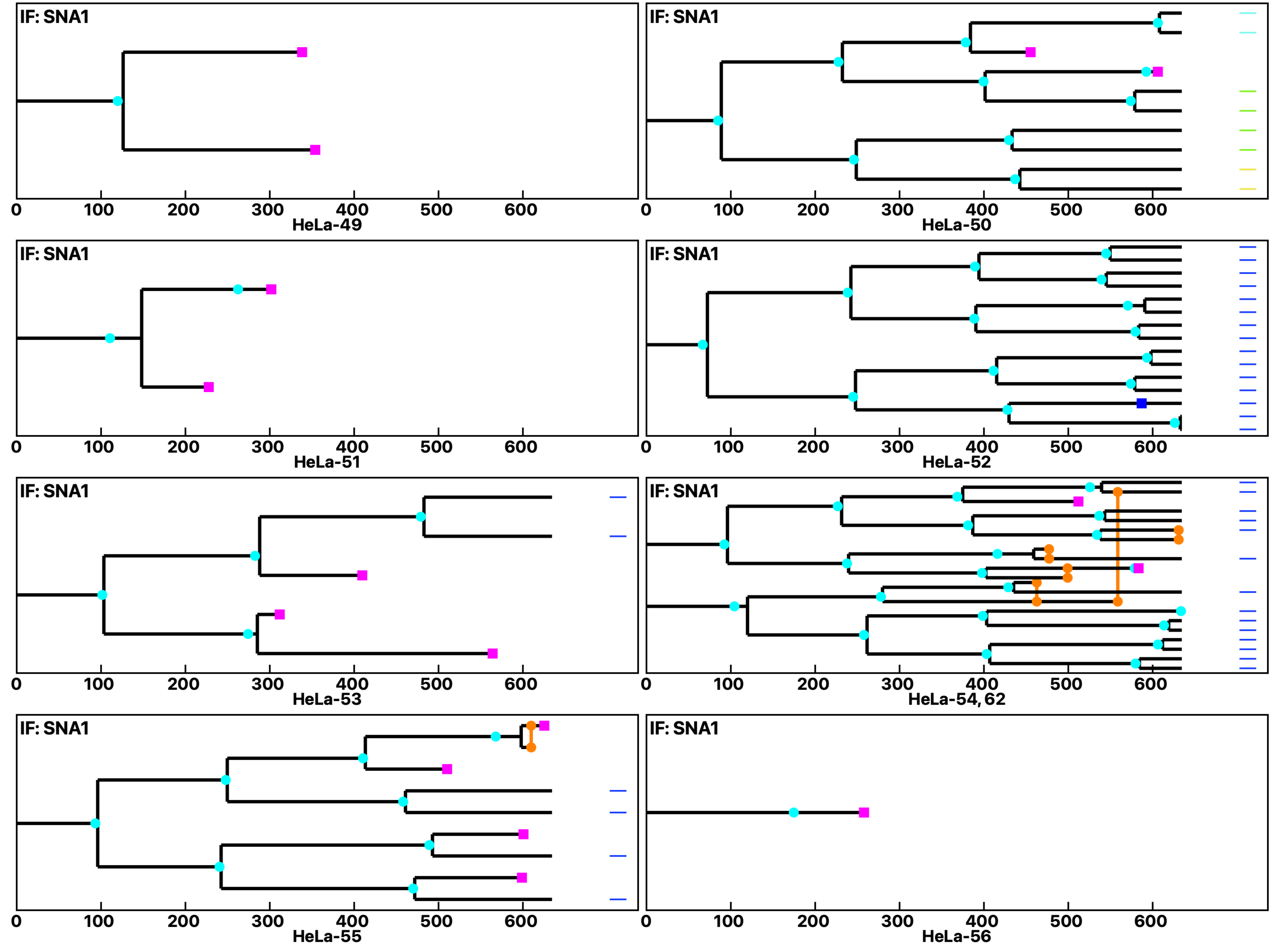

Analysis: HeLa, Treat.: HeLa, Cell: HeLa

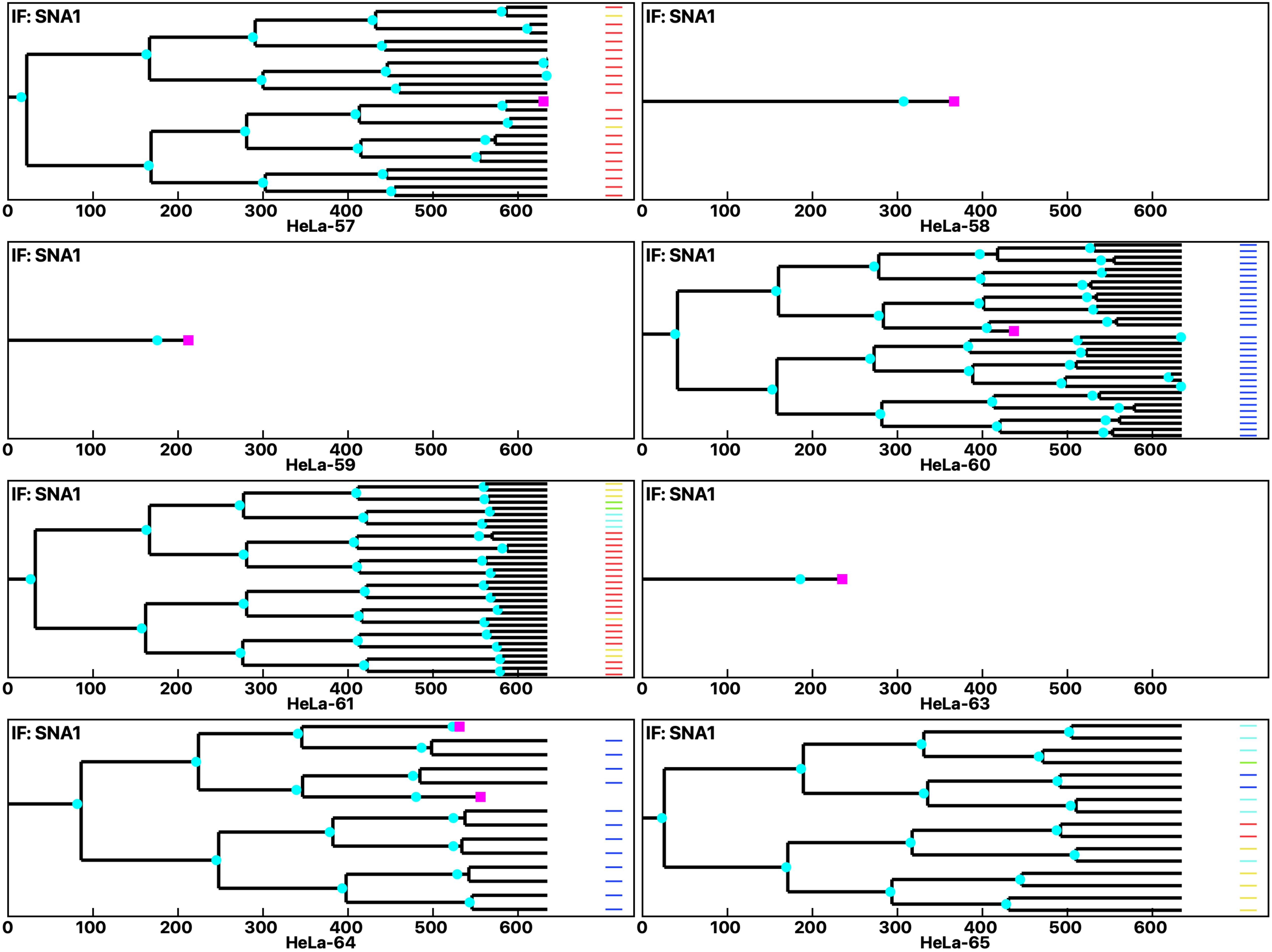

Analysis: HeLa, Treat.: HeLa, Cell: HeLa

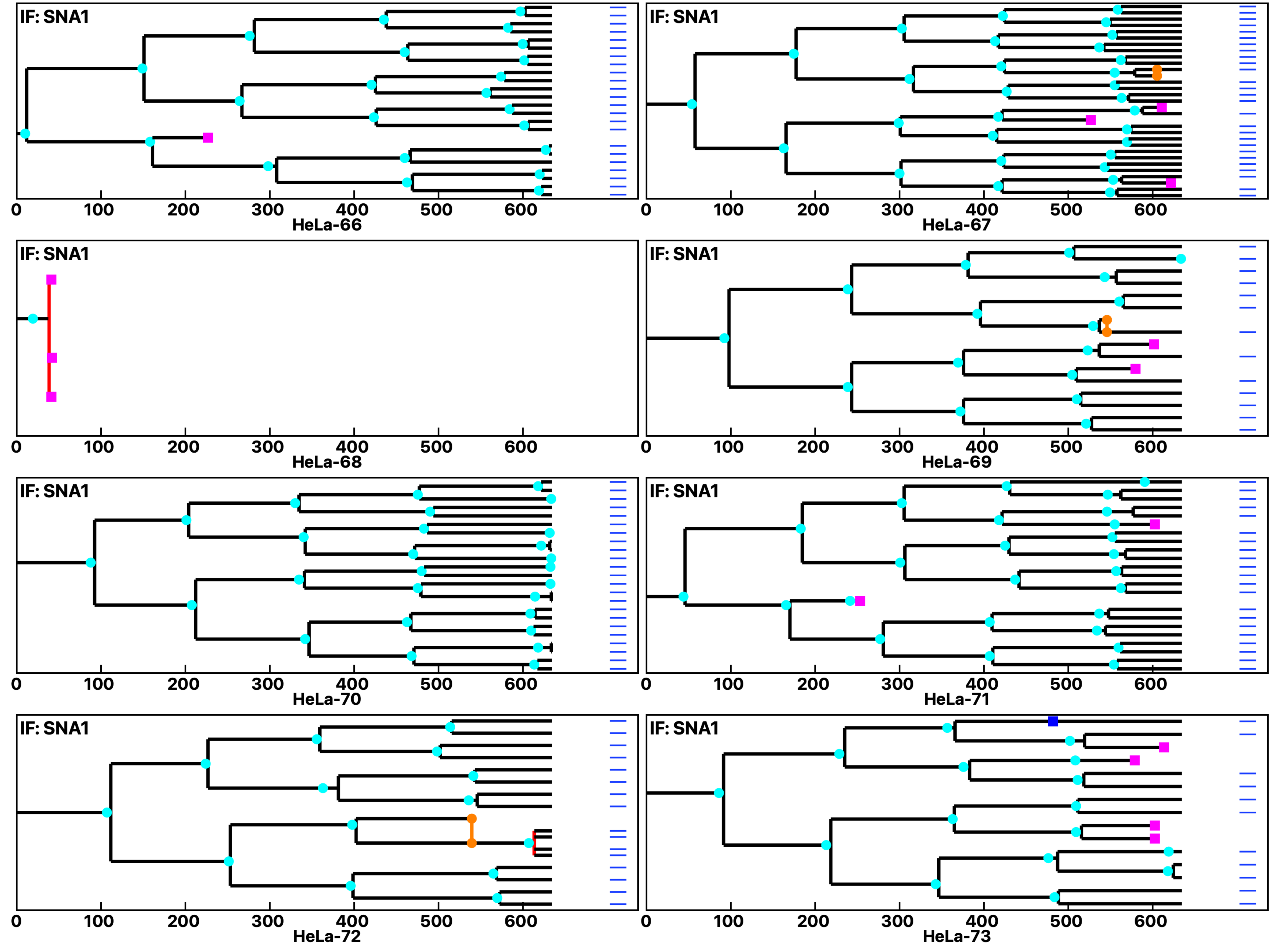

Analysis: HeLa, Treat.: HeLa, Cell: HeLa

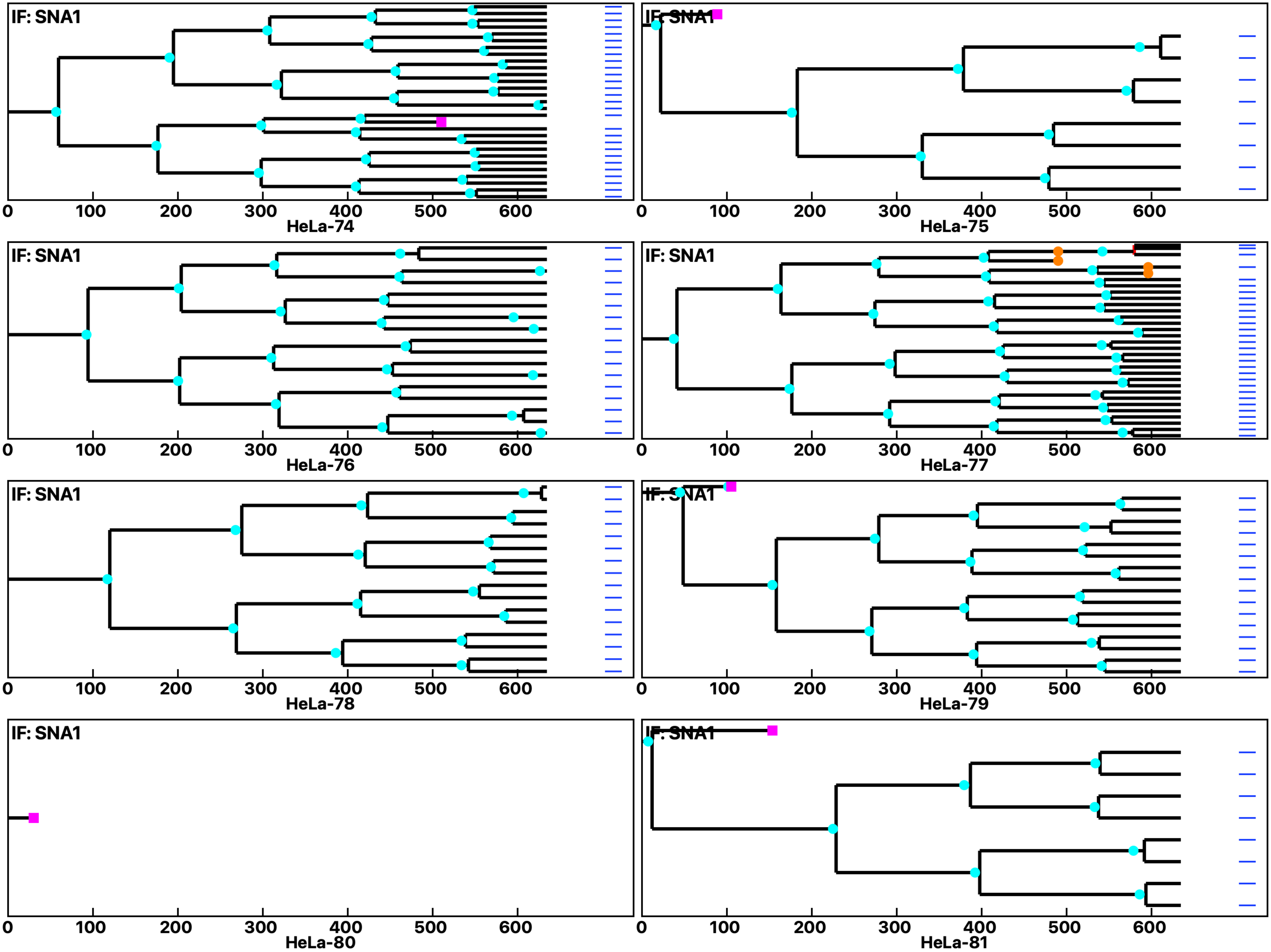

Analysis: HeLa, Treat.: HeLa, Cell: HeLa

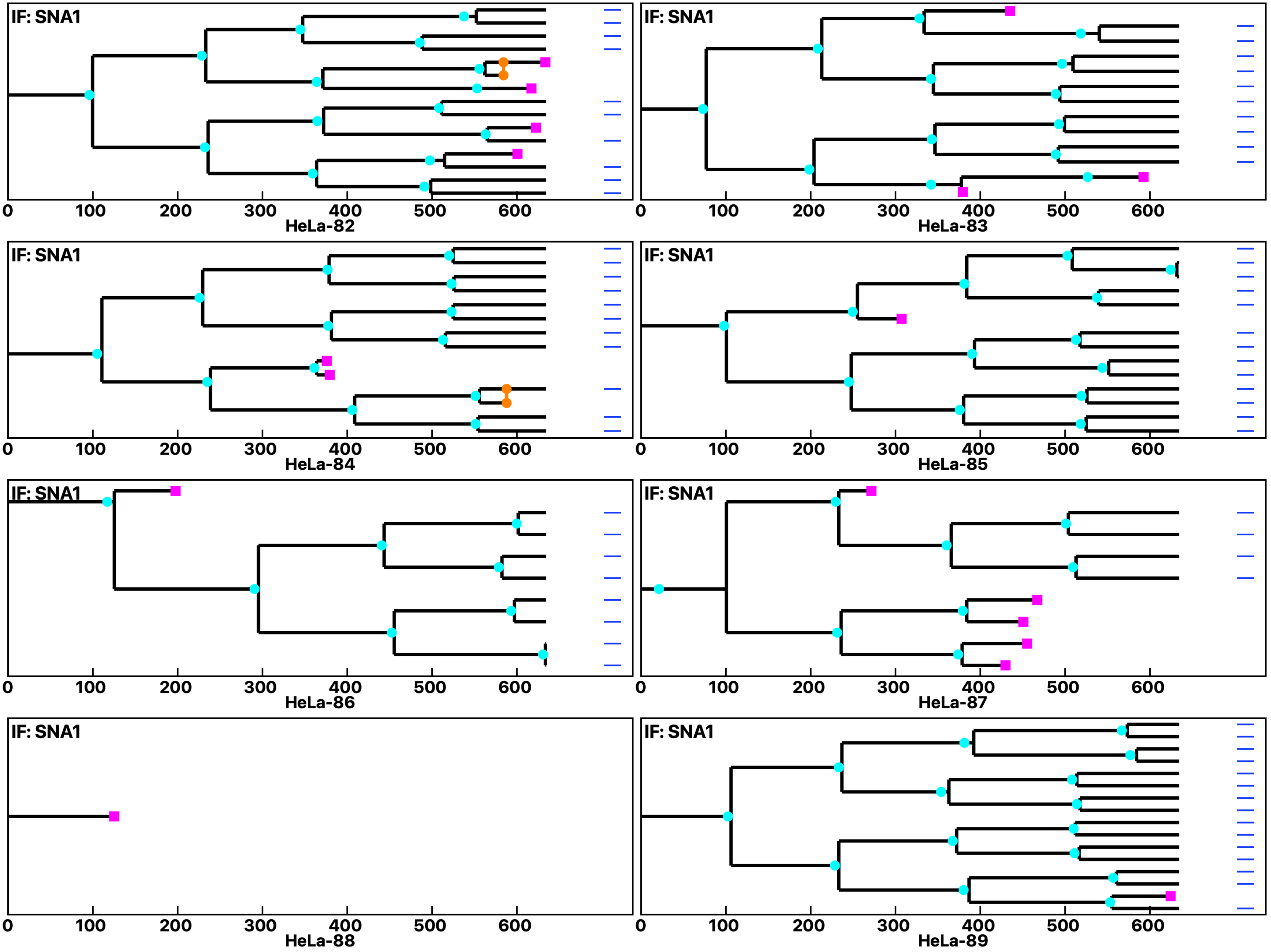

**Analysis: HeLa, Treat.: HeLa, Cell: HeLa**

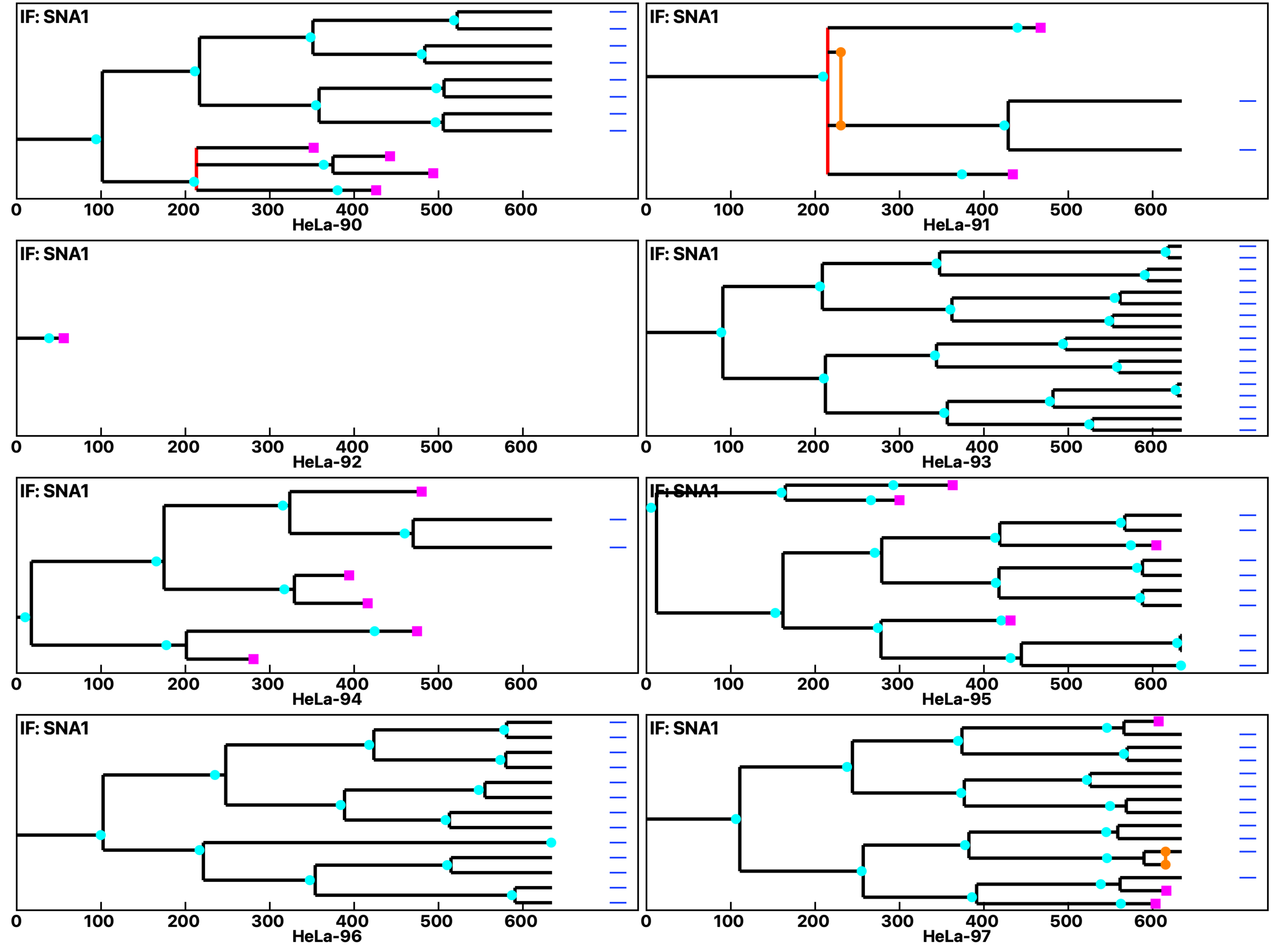

**Analysis: HeLa, Treat.: HeLa, Cell: HeLa**

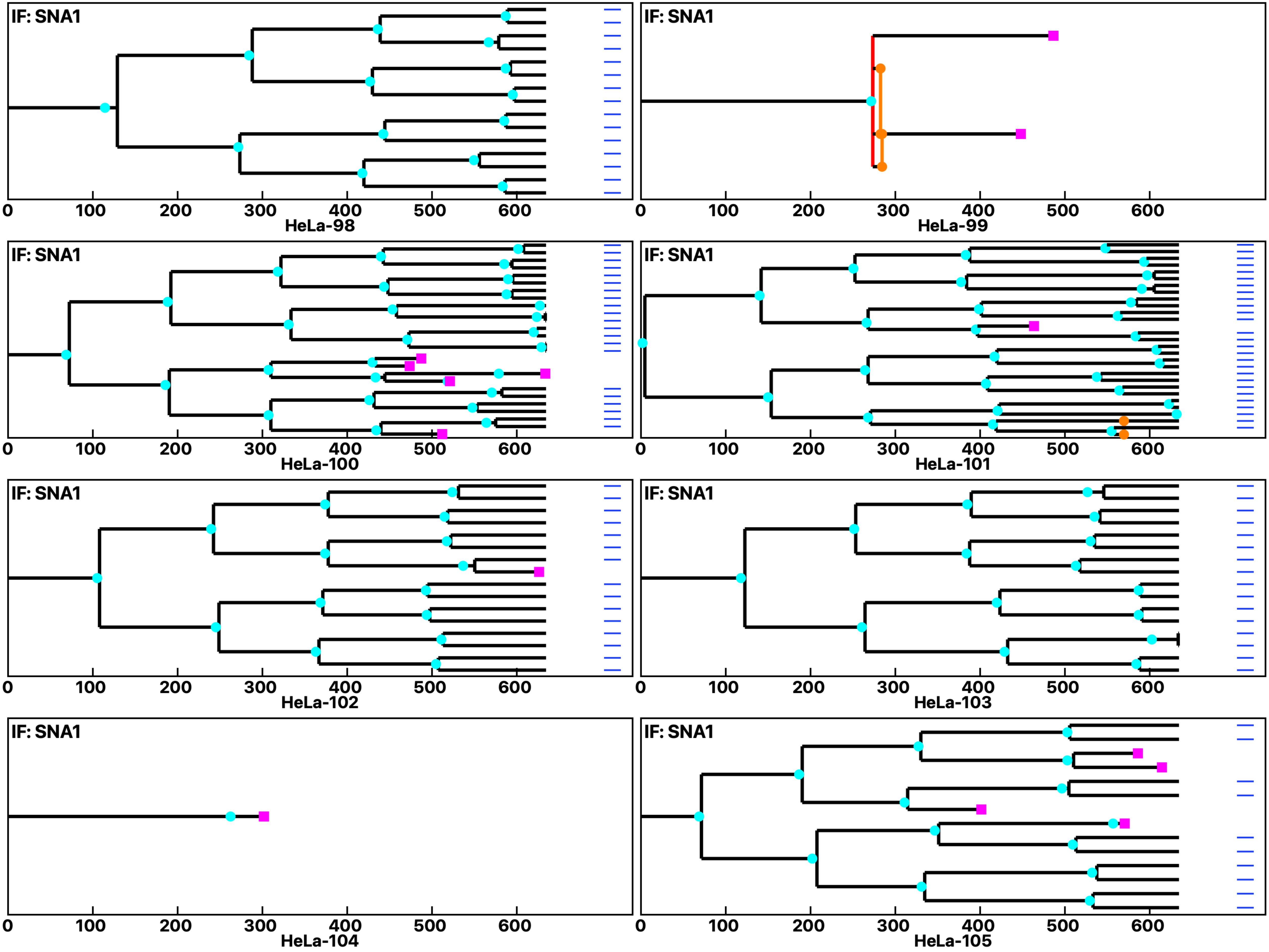

**Analysis: HeLa, Treat.: HeLa, Cell: HeLa**

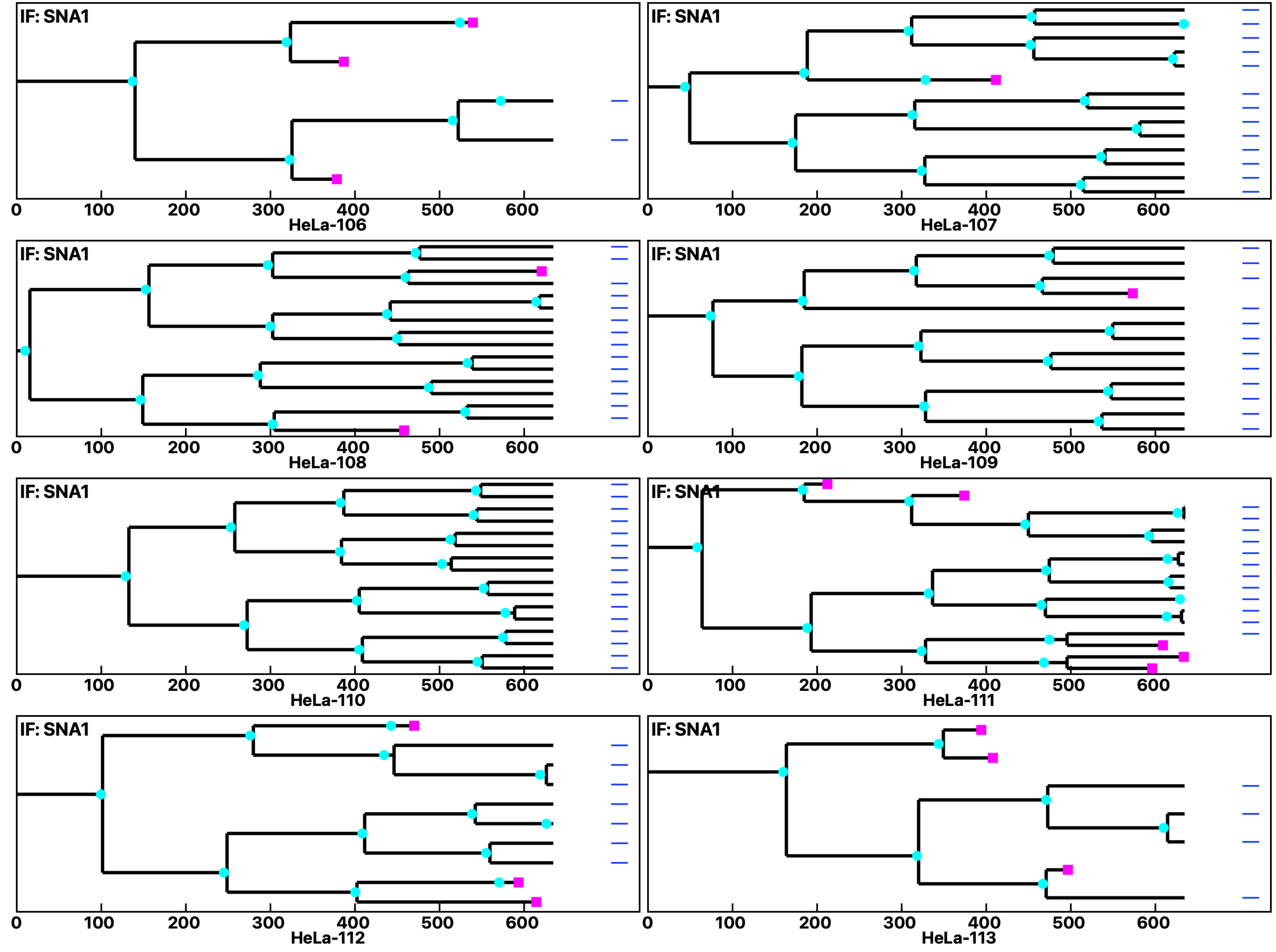

**Analysis: HeLa, Treat.: HeLa, Cell: HeLa**

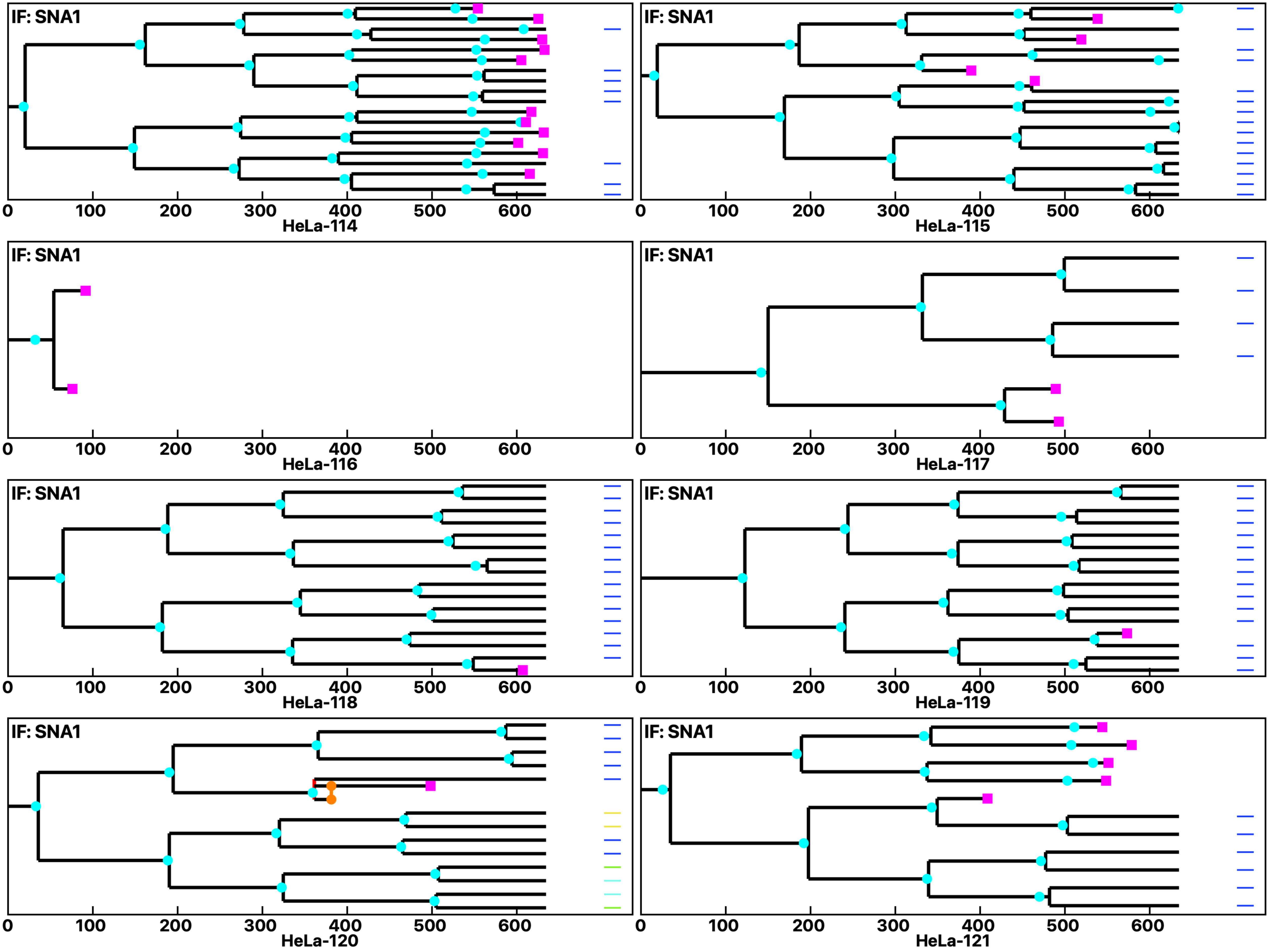

Analysis: HeLa, Treat.: HeLa, Cell: HeLa

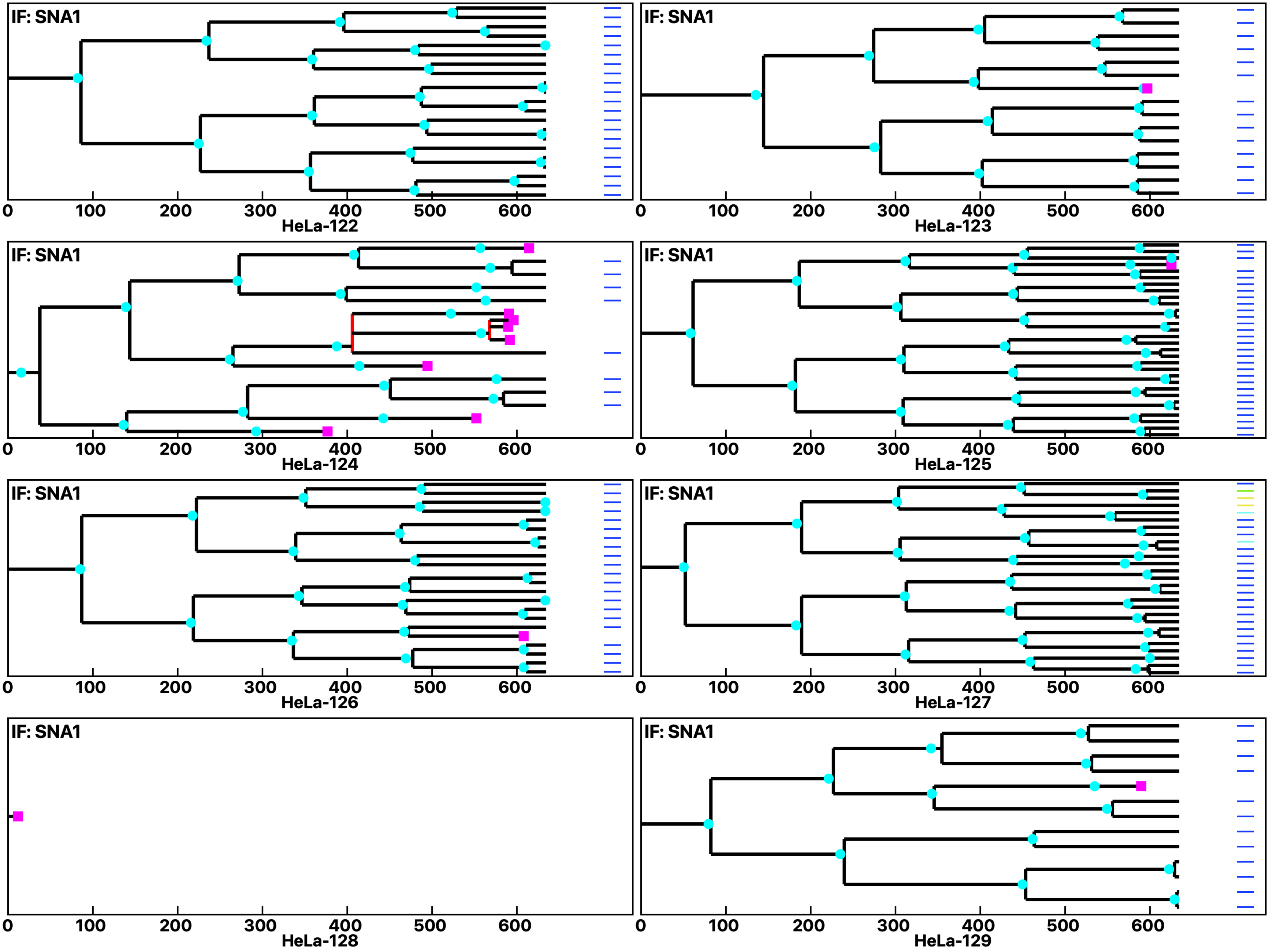

Analysis: HeLa, Treat.: HeLa, Cell: HeLa

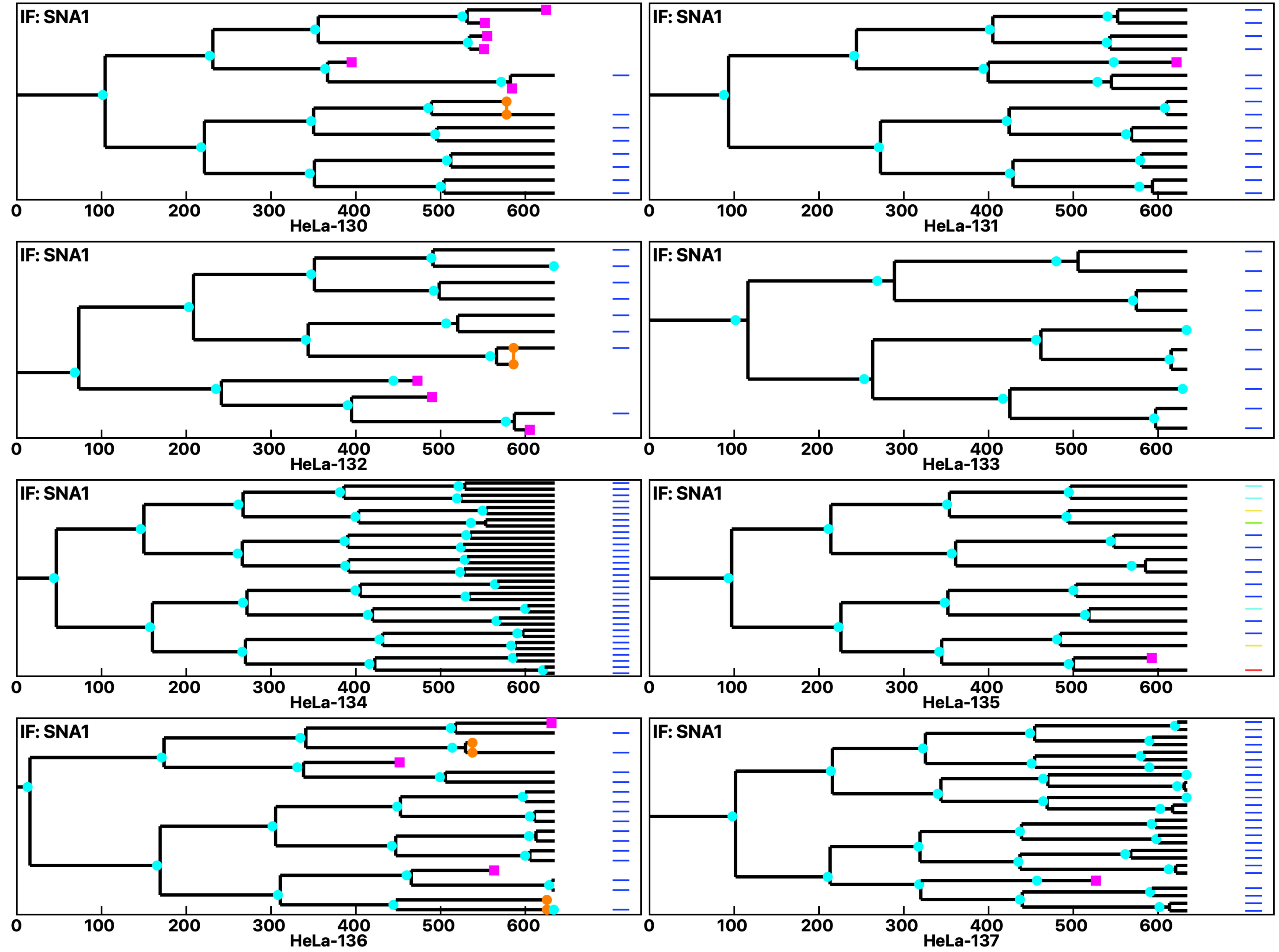

Analysis: HeLa, Treat.: HeLa, Cell: HeLa

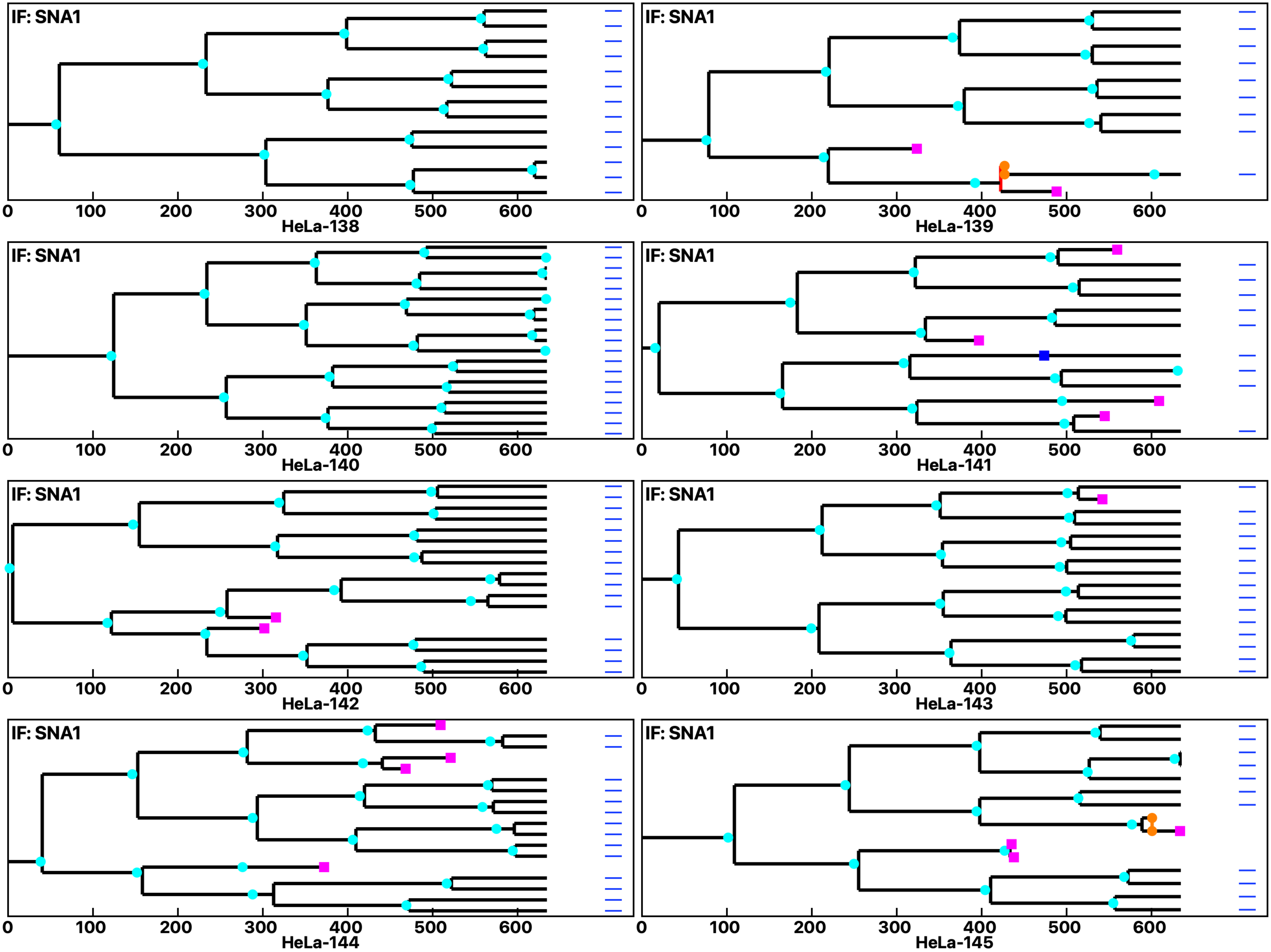

Analysis: HeLa, Treat.: HeLa, Cell: HeLa

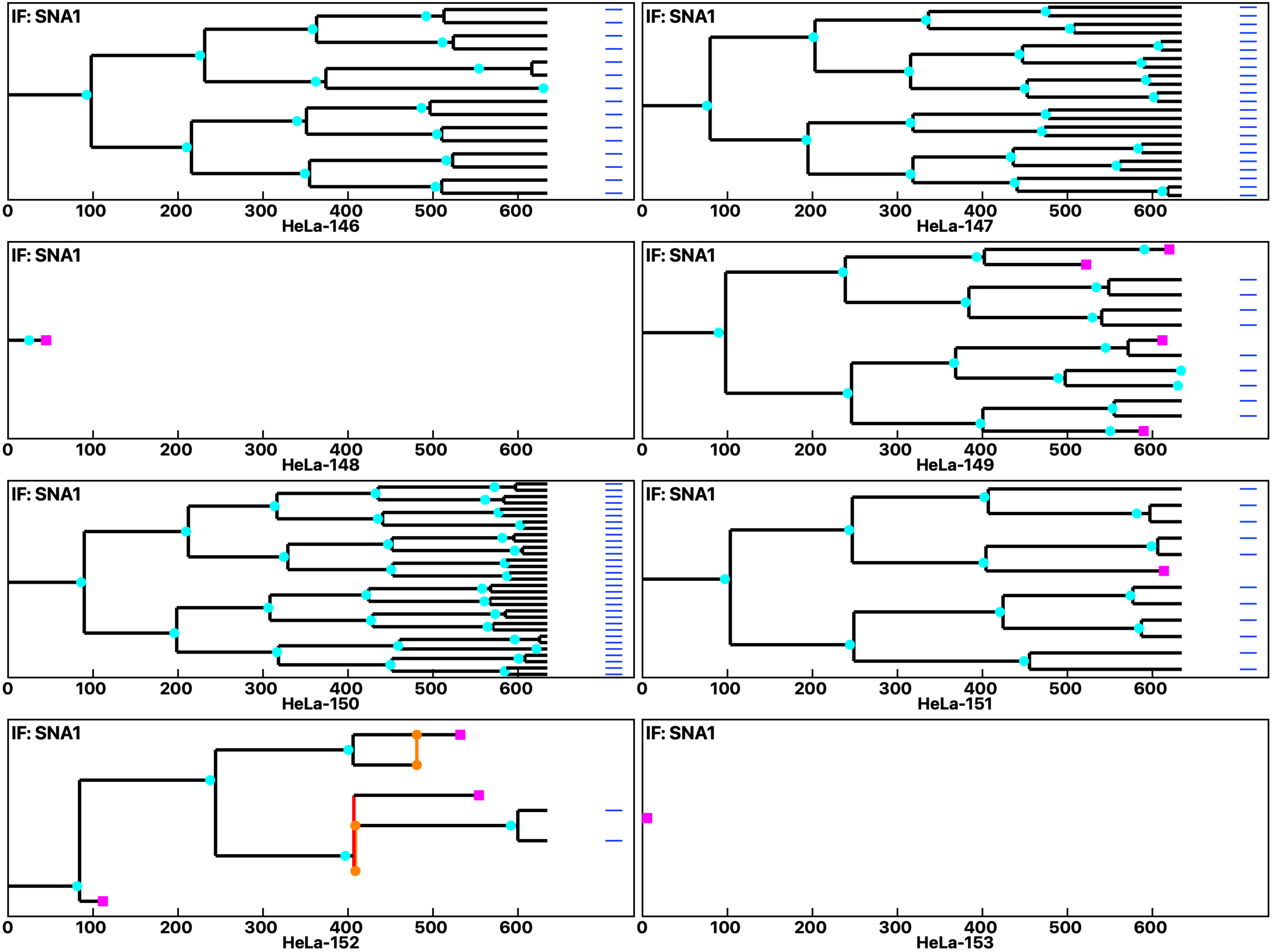

Analysis: HeLa, Treat.: HeLa, Cell: HeLa

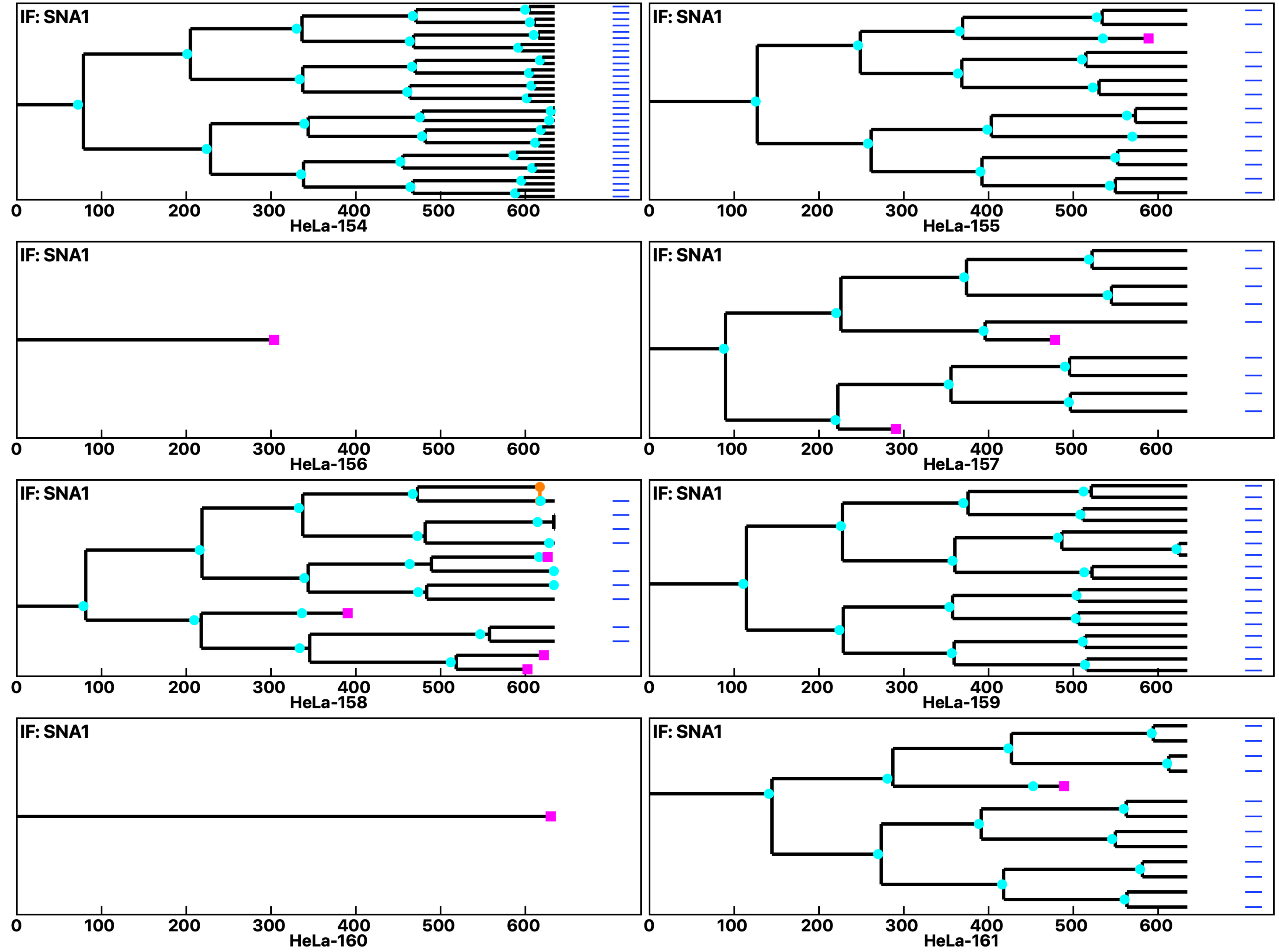

Analysis: HeLa, Treat.: HeLa, Cell: HeLa

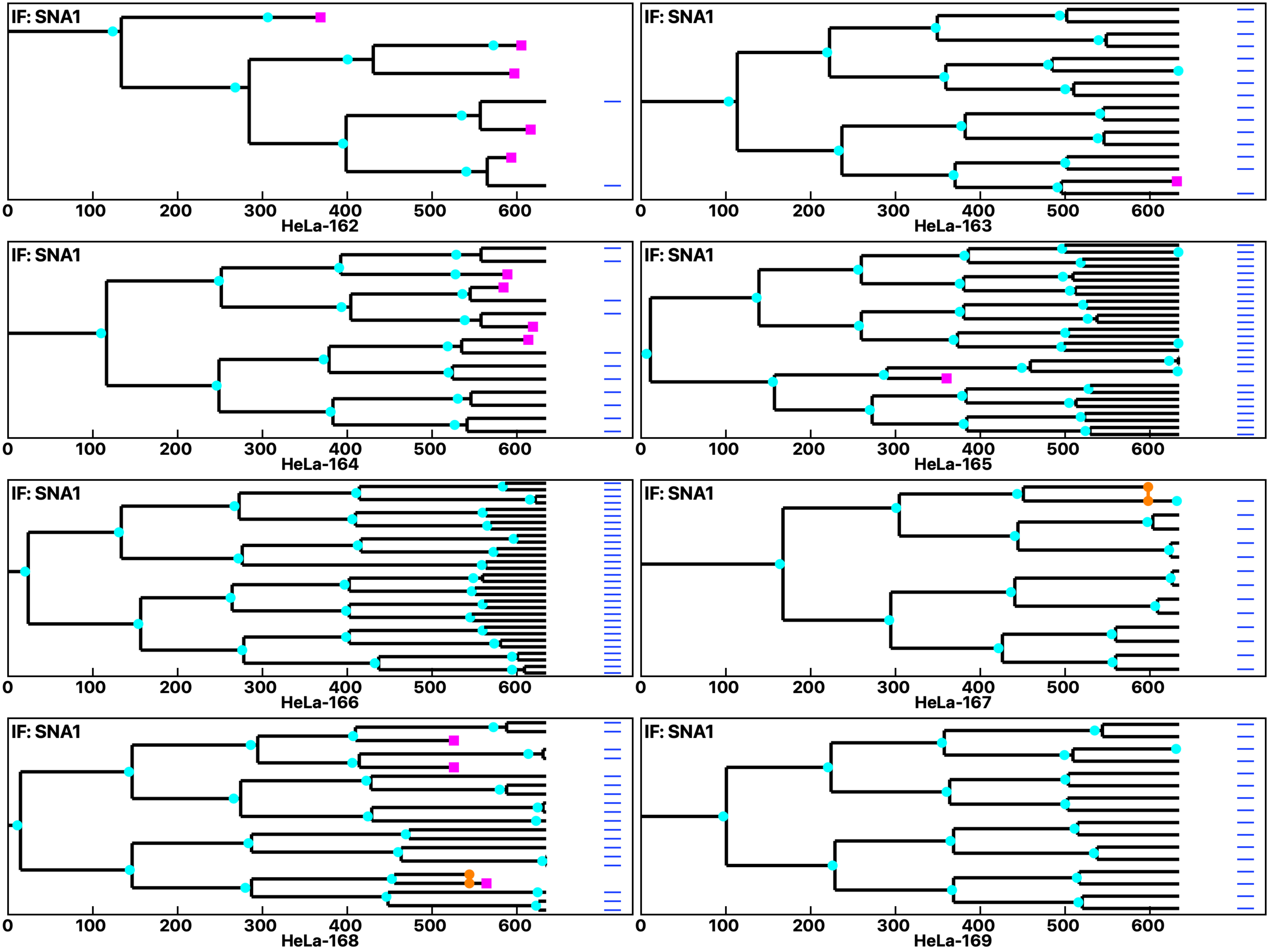

Analysis: HeLa, Treat.: HeLa, Cell: HeLa

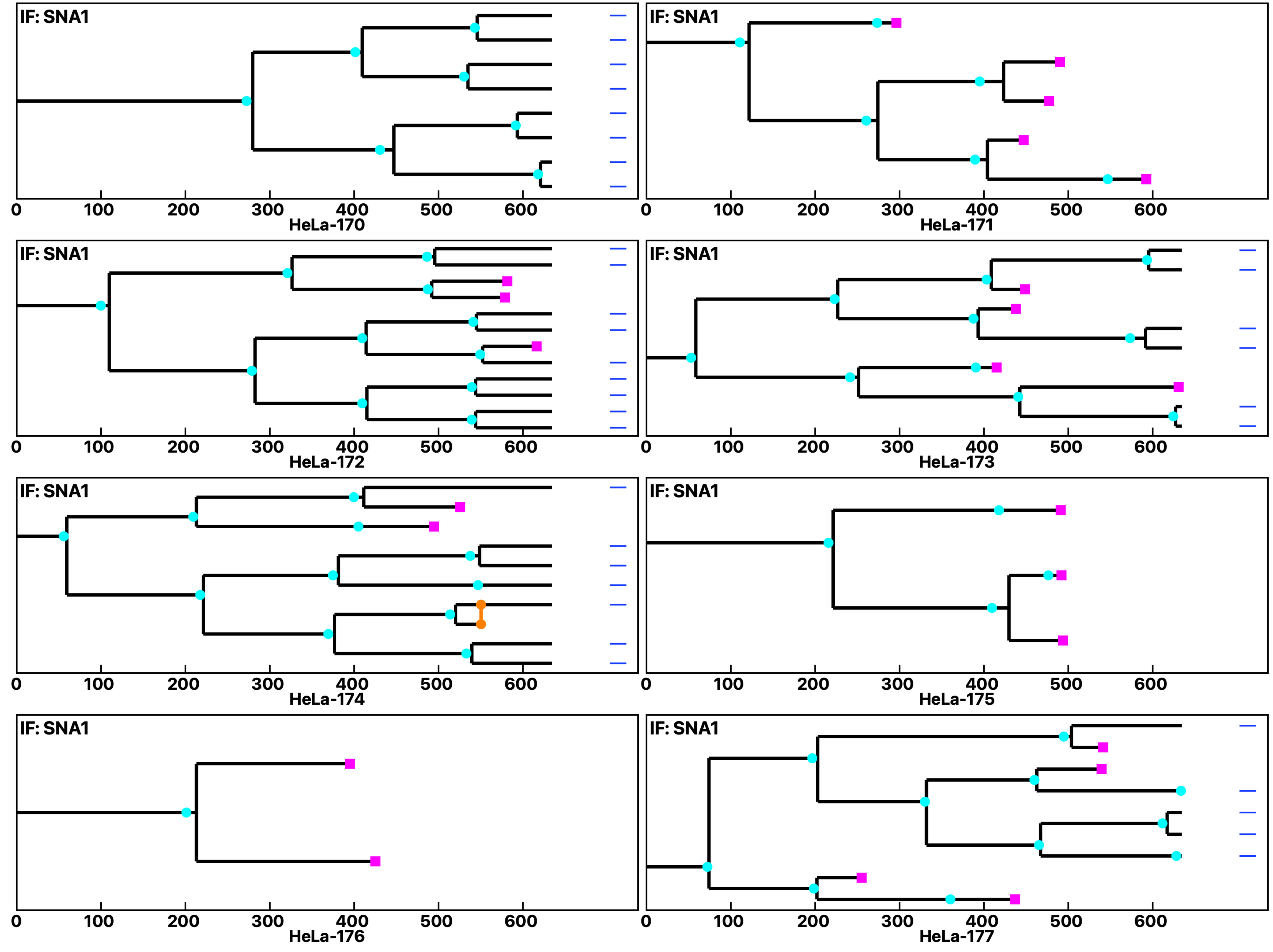

Analysis: HeLa, Treat.: HeLa, Cell: HeLa

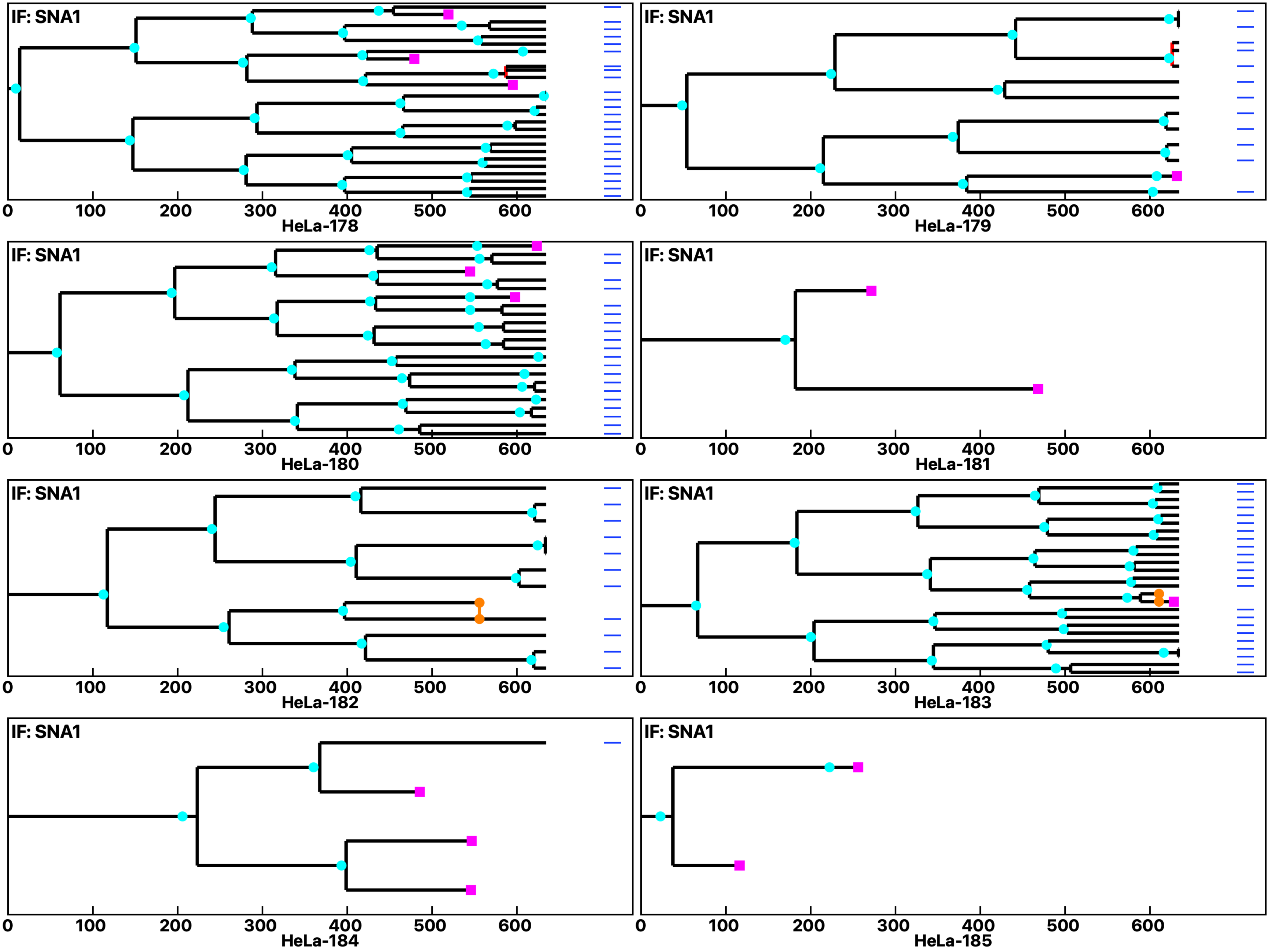

Analysis: HeLa, Treat.: HeLa, Cell: HeLa

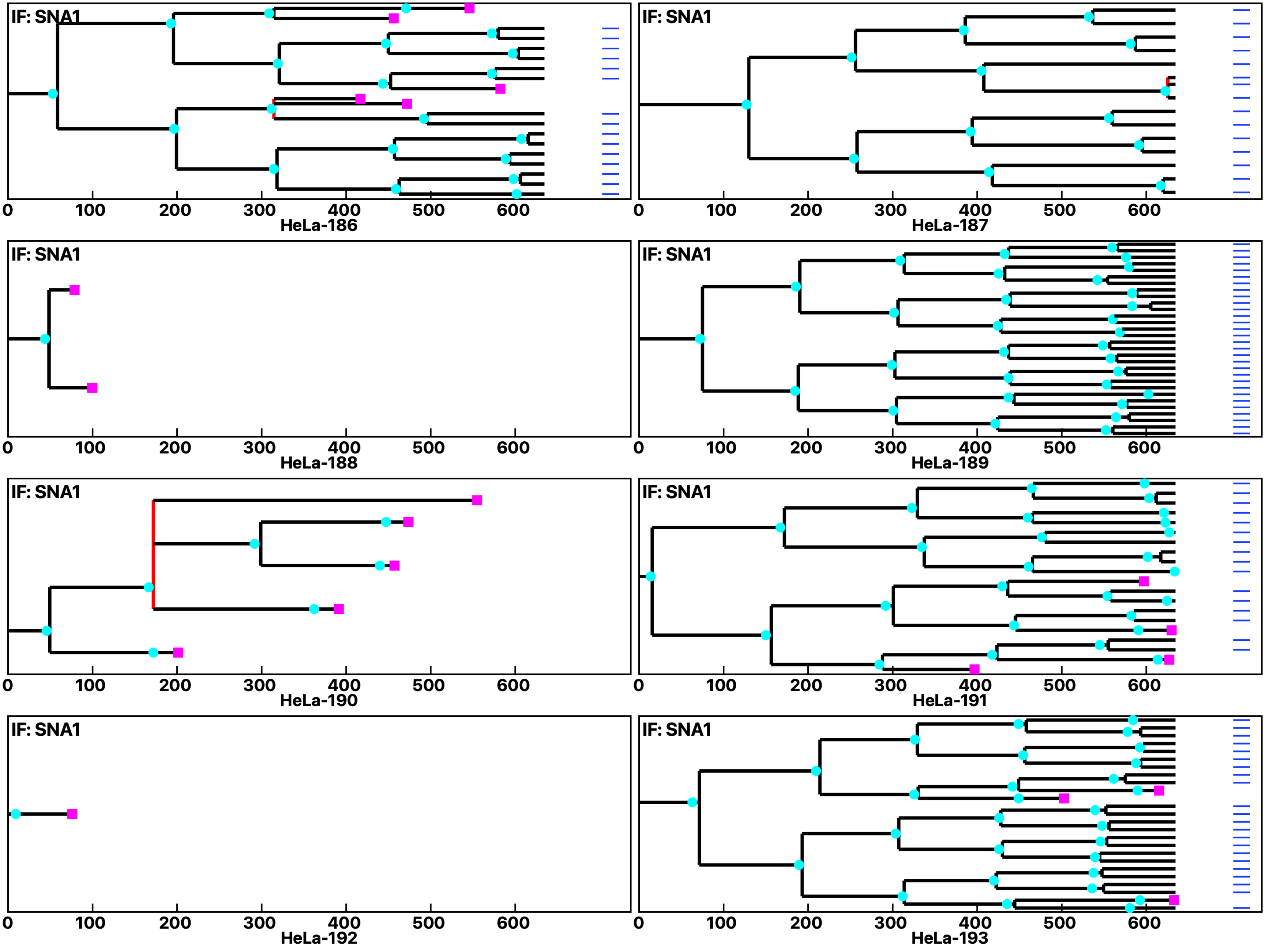

**Analysis: HeLa, Treat.: HeLa, Cell: HeLa**

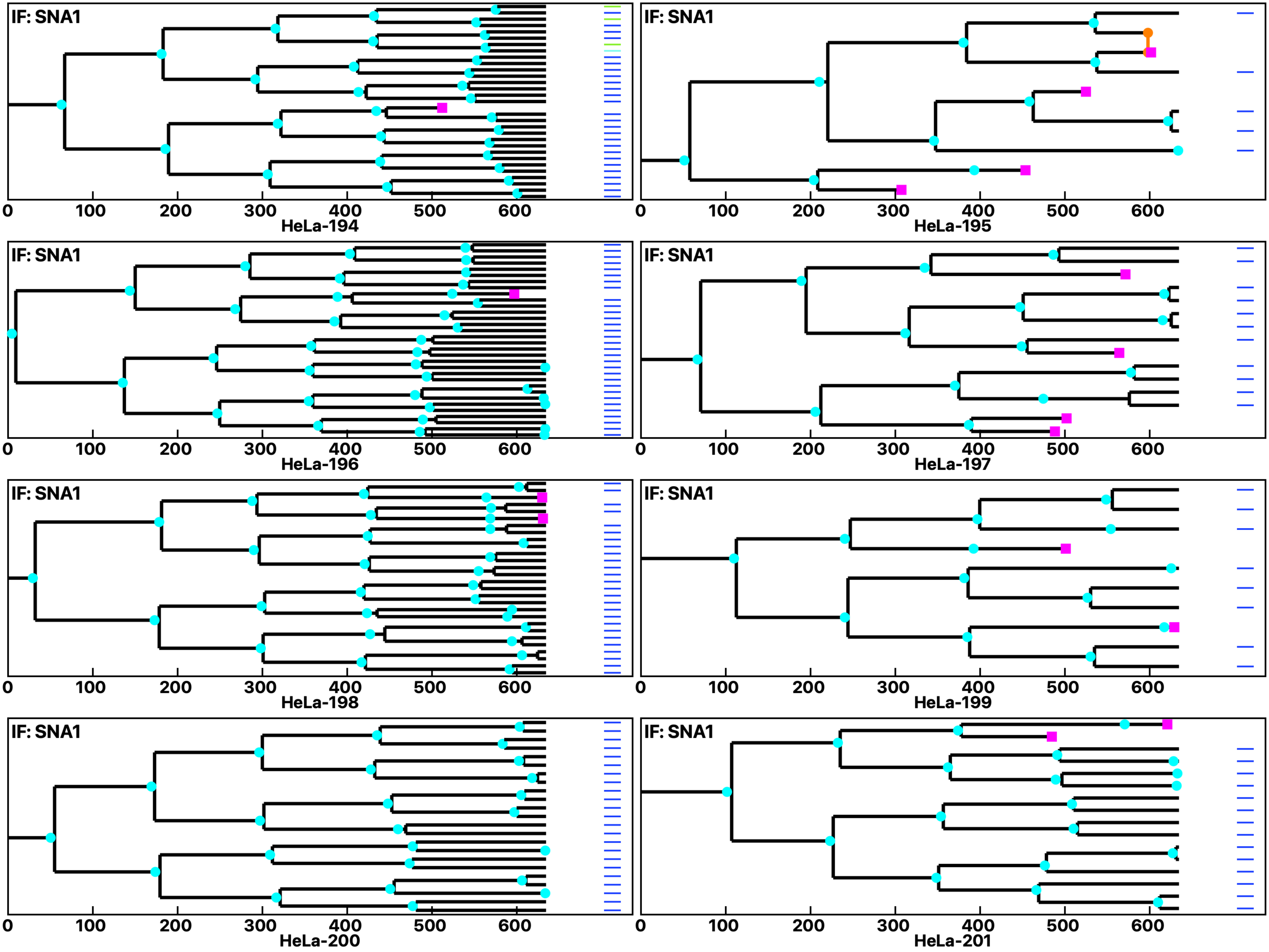

Analysis: HeLa, Treat.: HeLa, Cell: HeLa

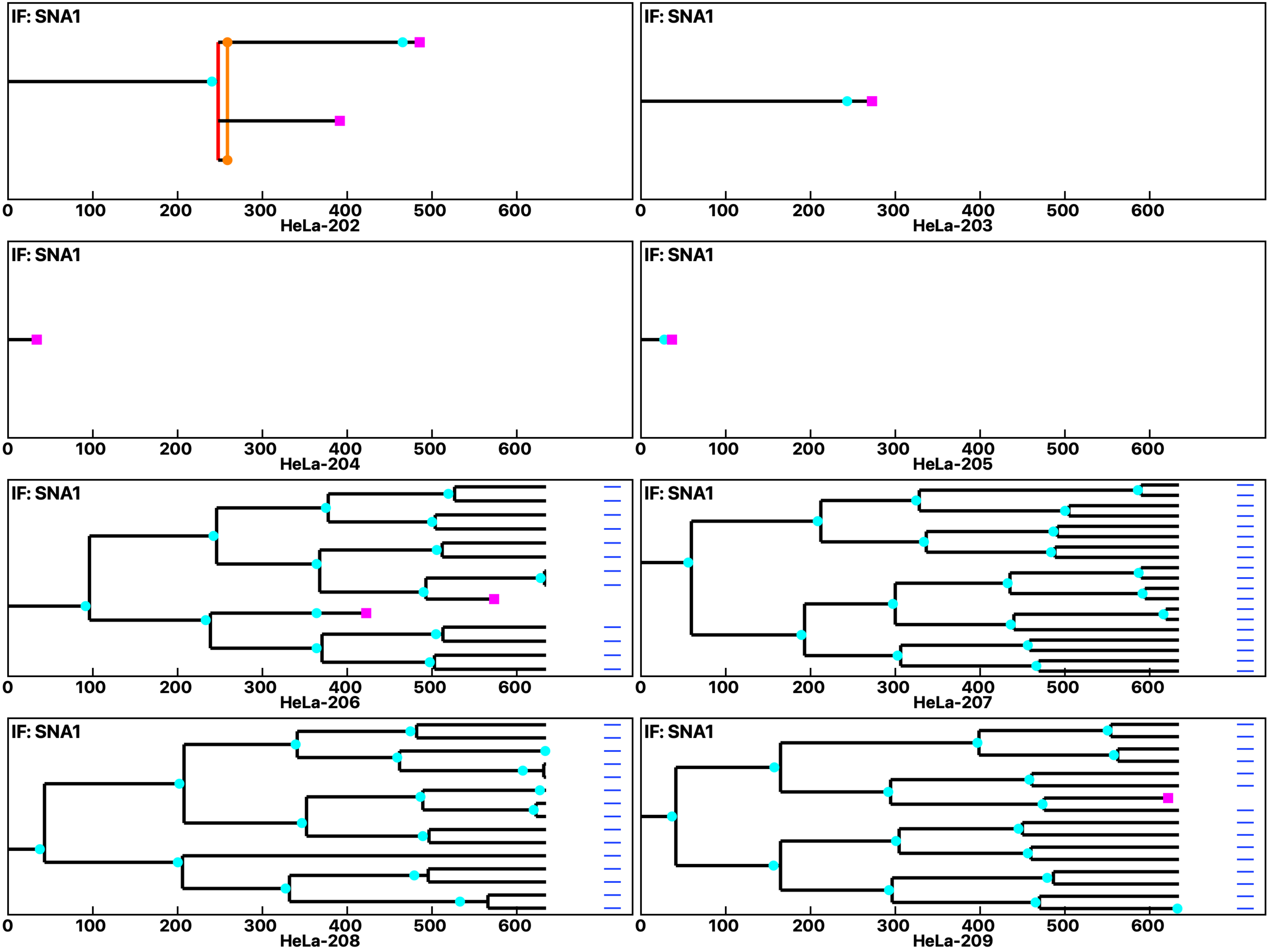

Analysis: HeLa, Treat.: HeLa, Cell: HeLa

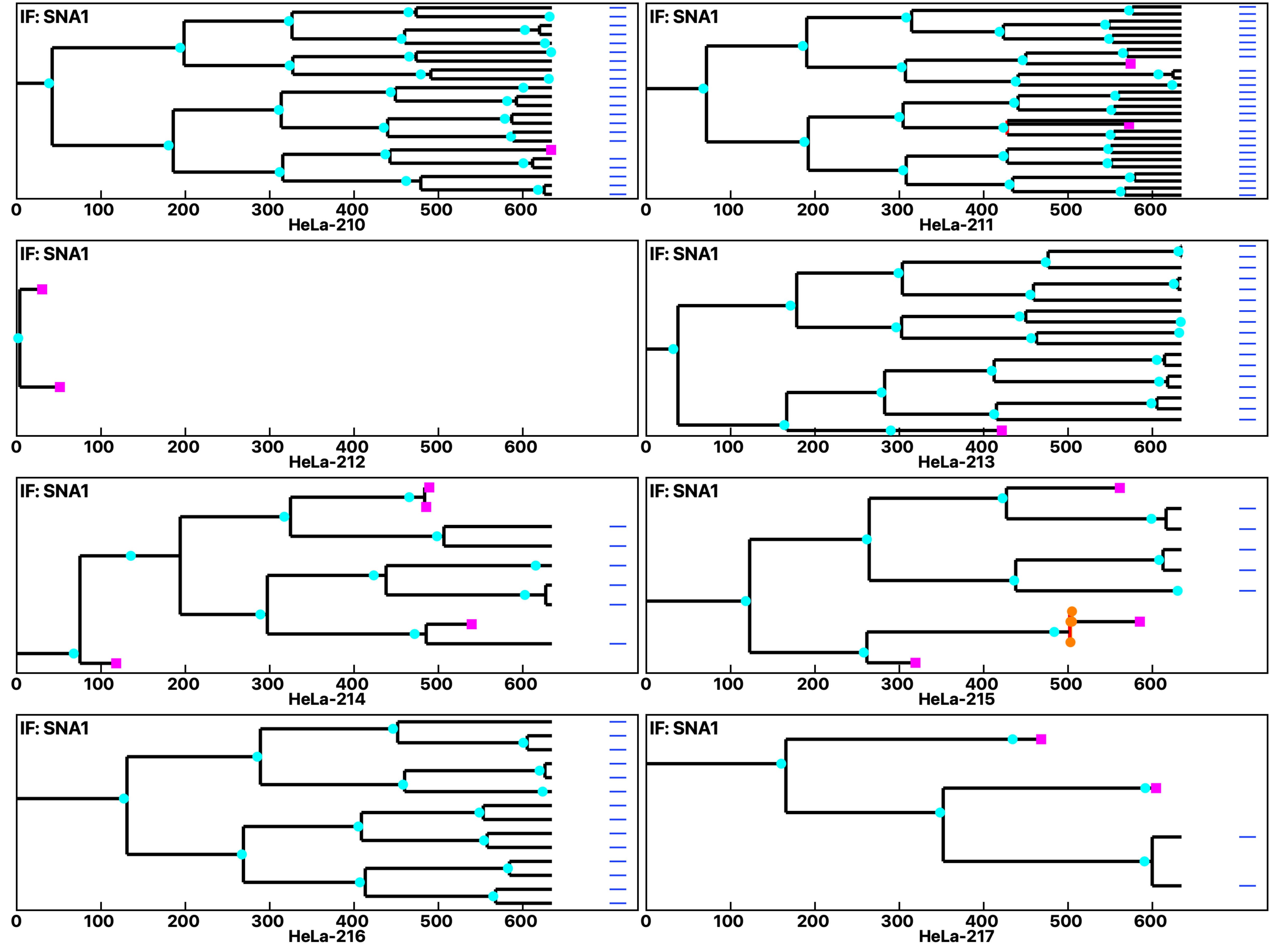

Analysis: HeLa, Treat.: HeLa, Cell: HeLa

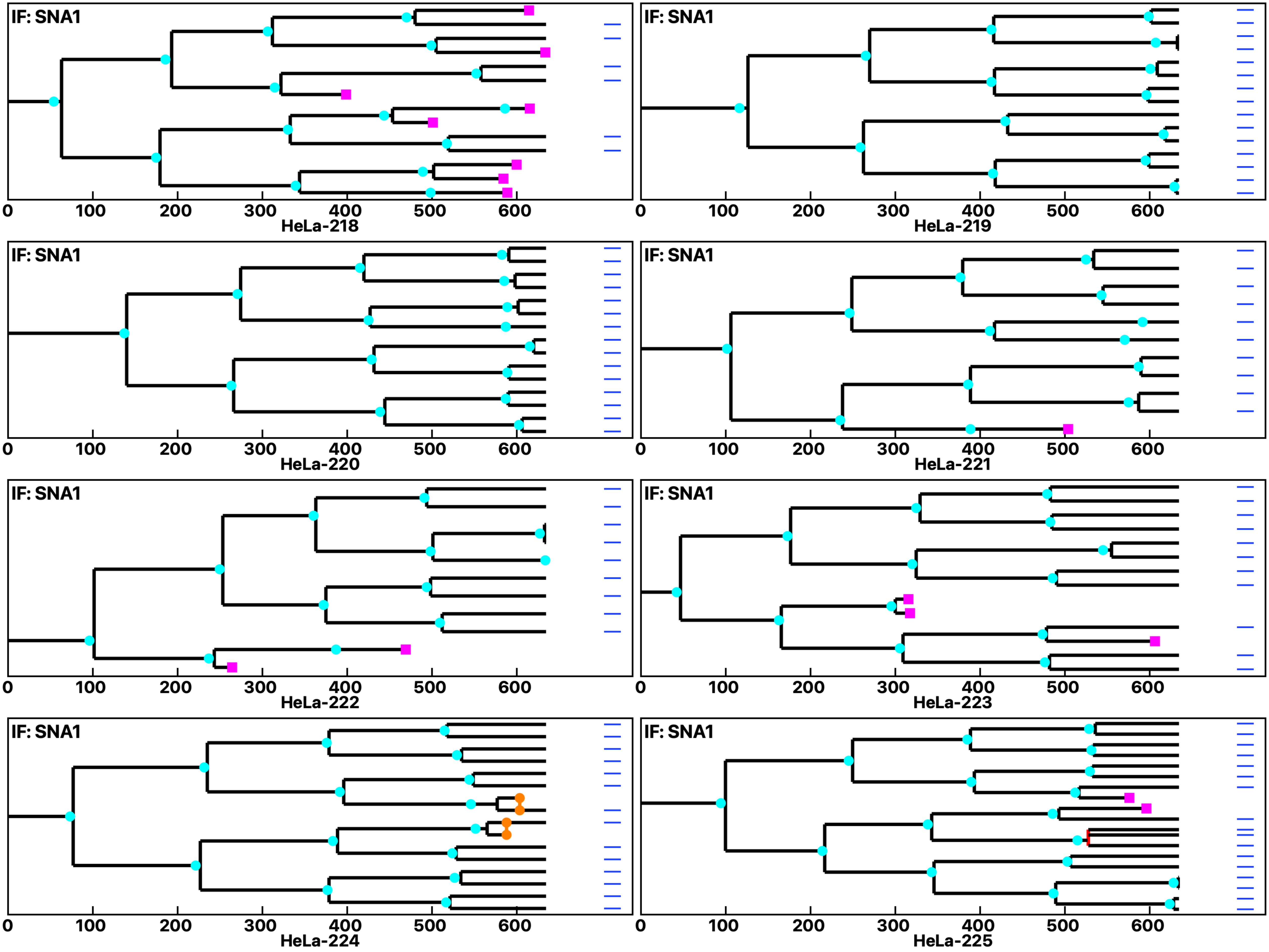

Analysis: HeLa, Treat.: HeLa, Cell: HeLa

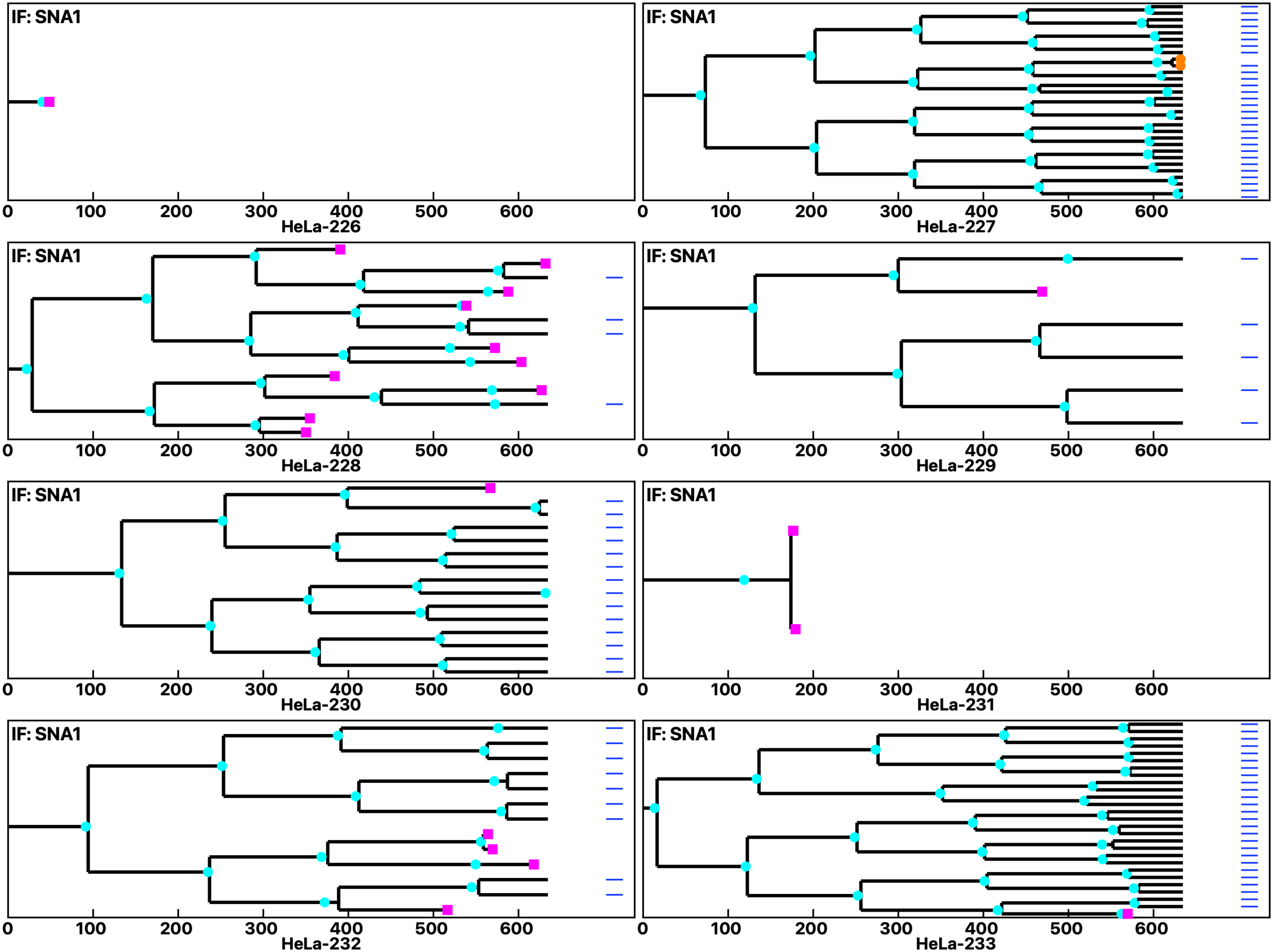

Analysis: HeLa, Treat.: HeLa, Cell: HeLa

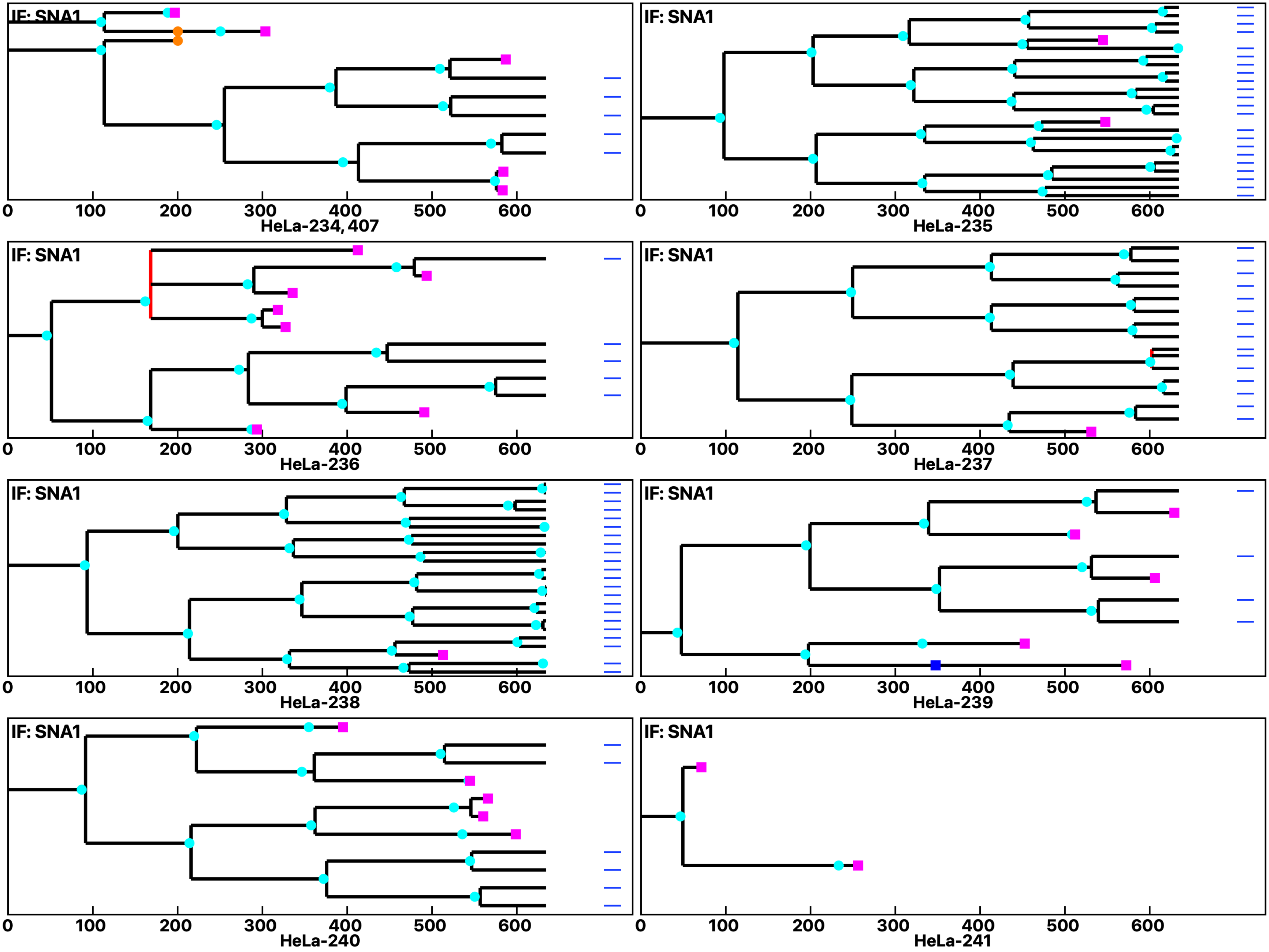

Analysis: HeLa, Treat.: HeLa, Cell: HeLa

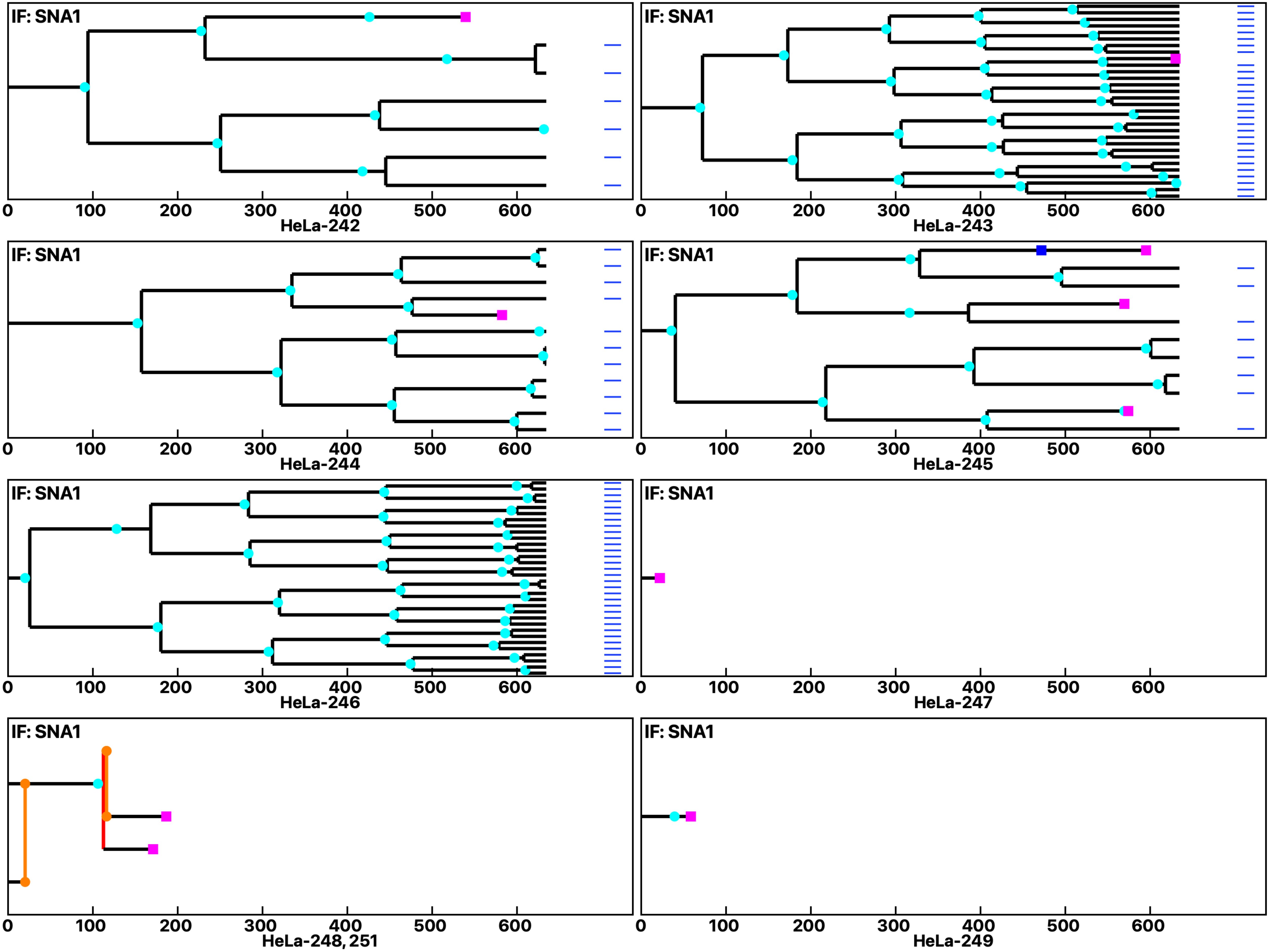

Analysis: HeLa, Treat.: HeLa, Cell: HeLa

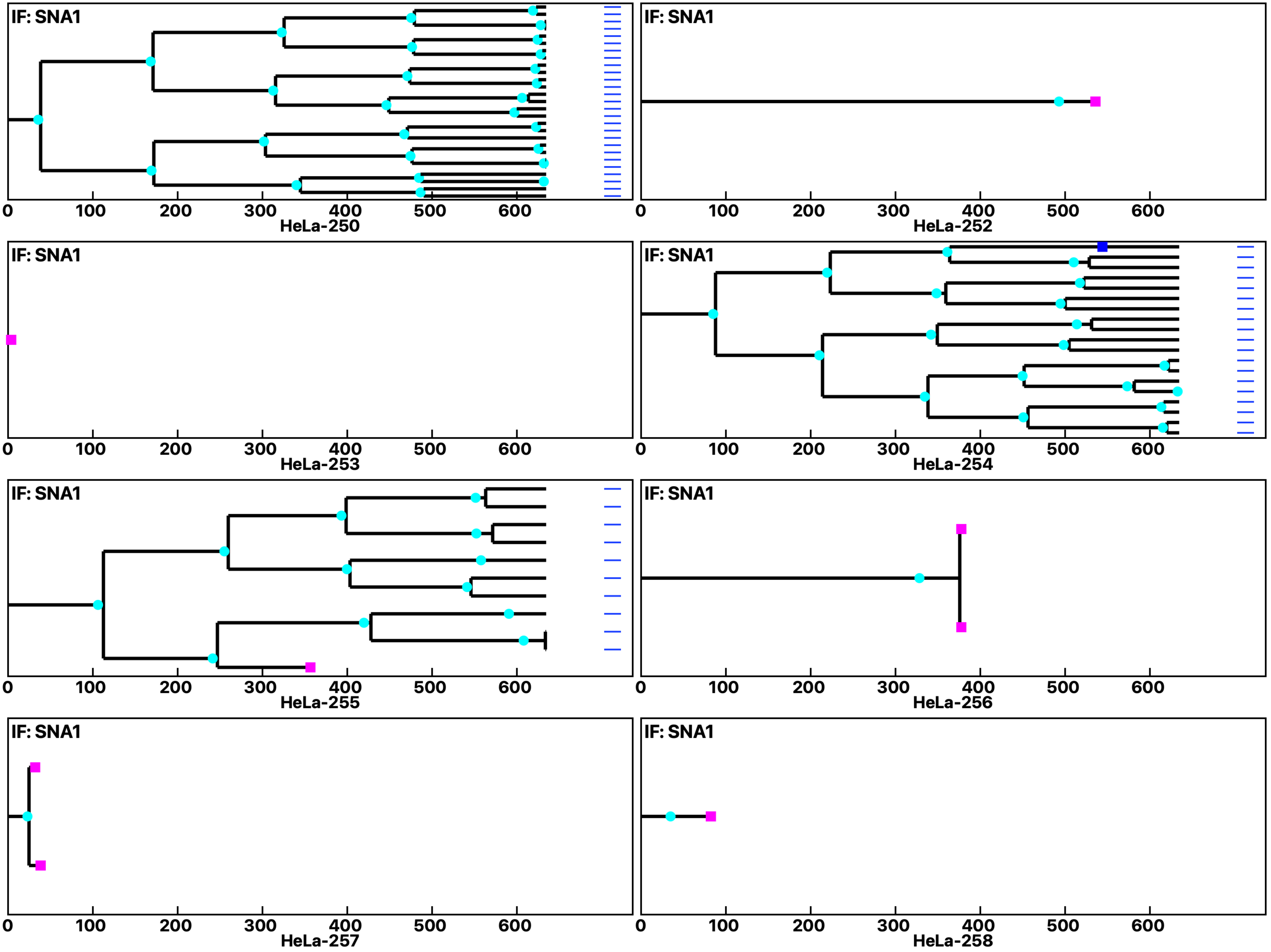

Analysis: HeLa, Treat.: HeLa, Cell: HeLa

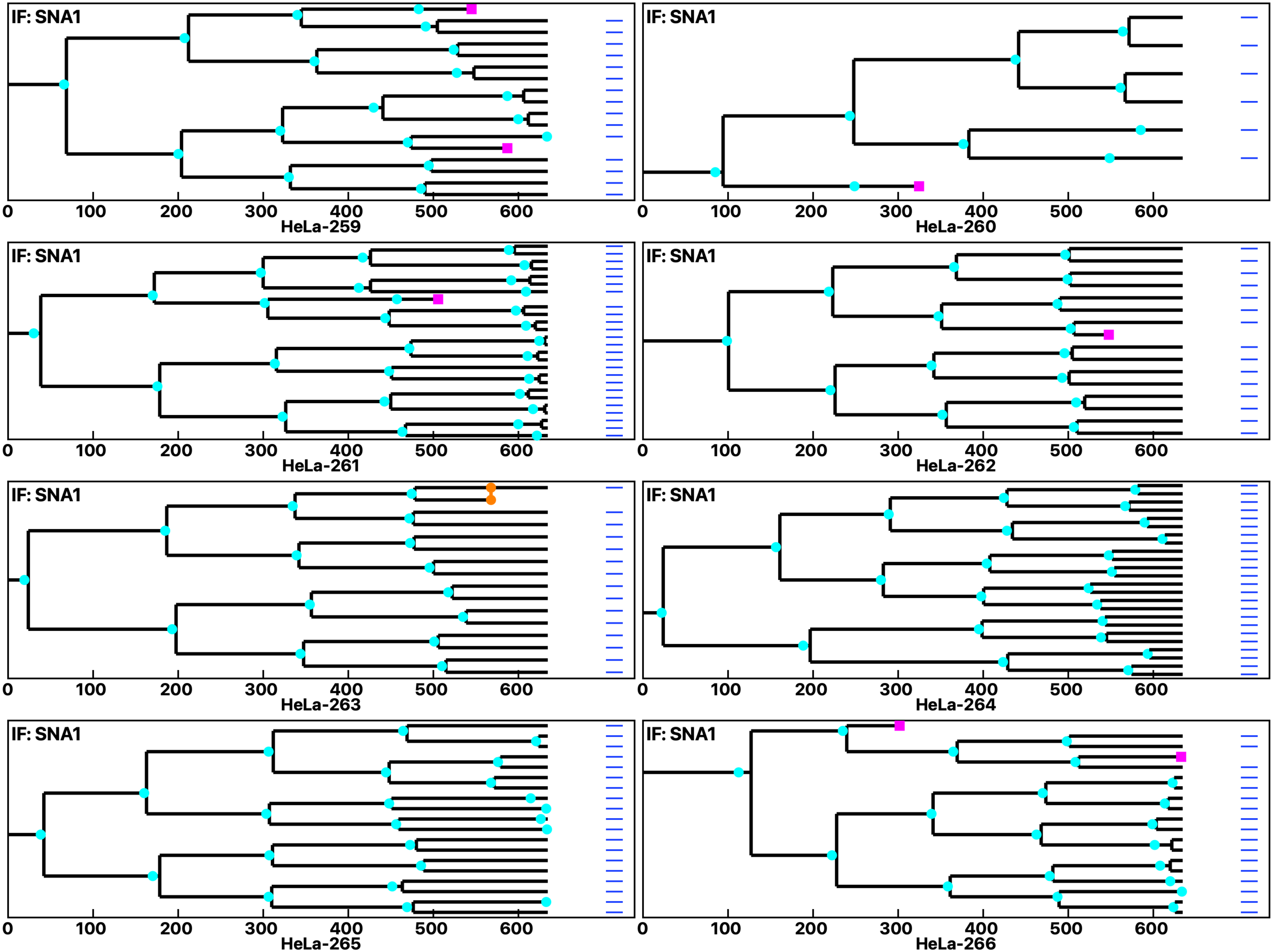

Analysis: HeLa, Treat.: HeLa, Cell: HeLa

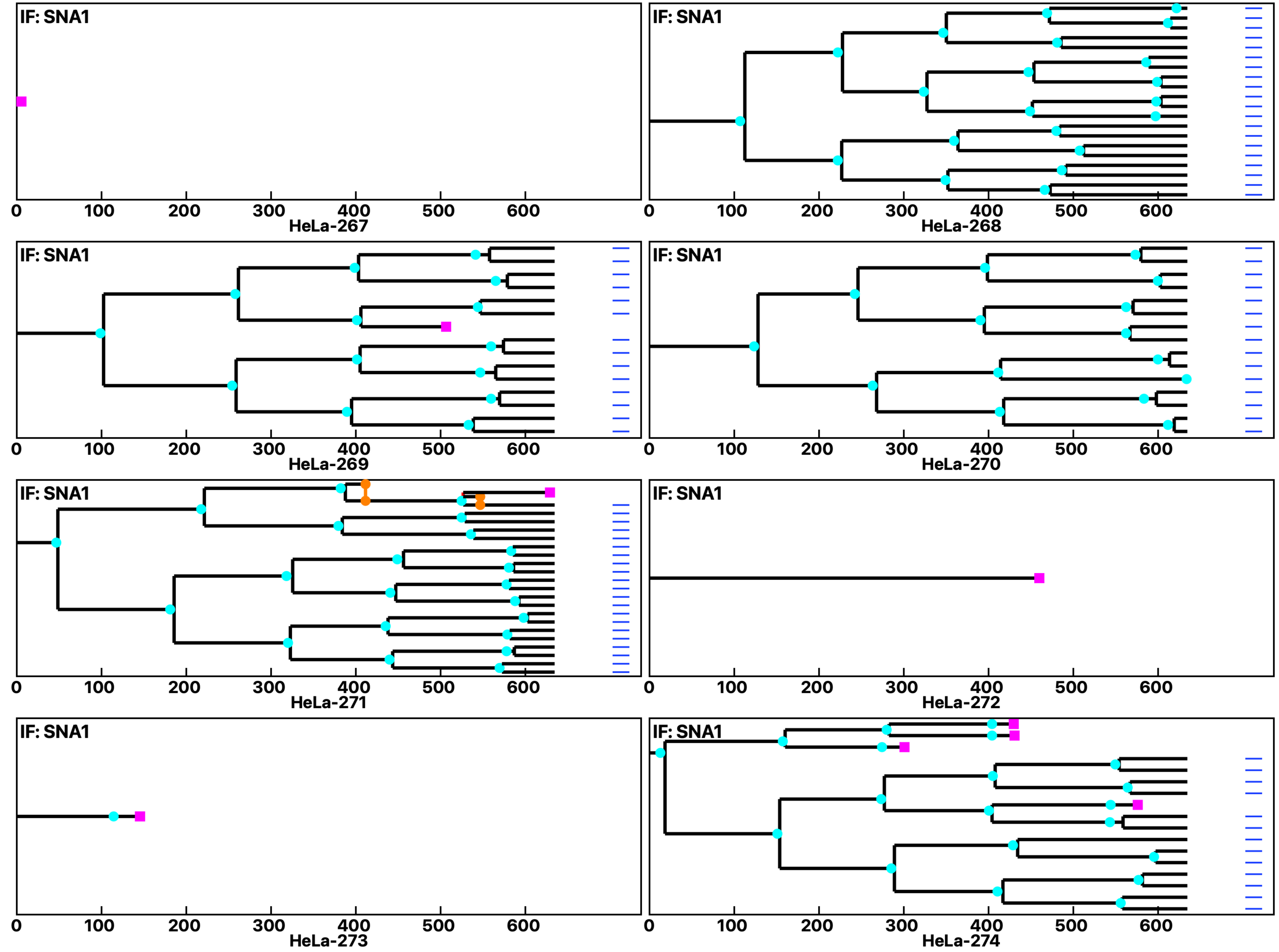

Analysis: HeLa, Treat.: HeLa, Cell: HeLa

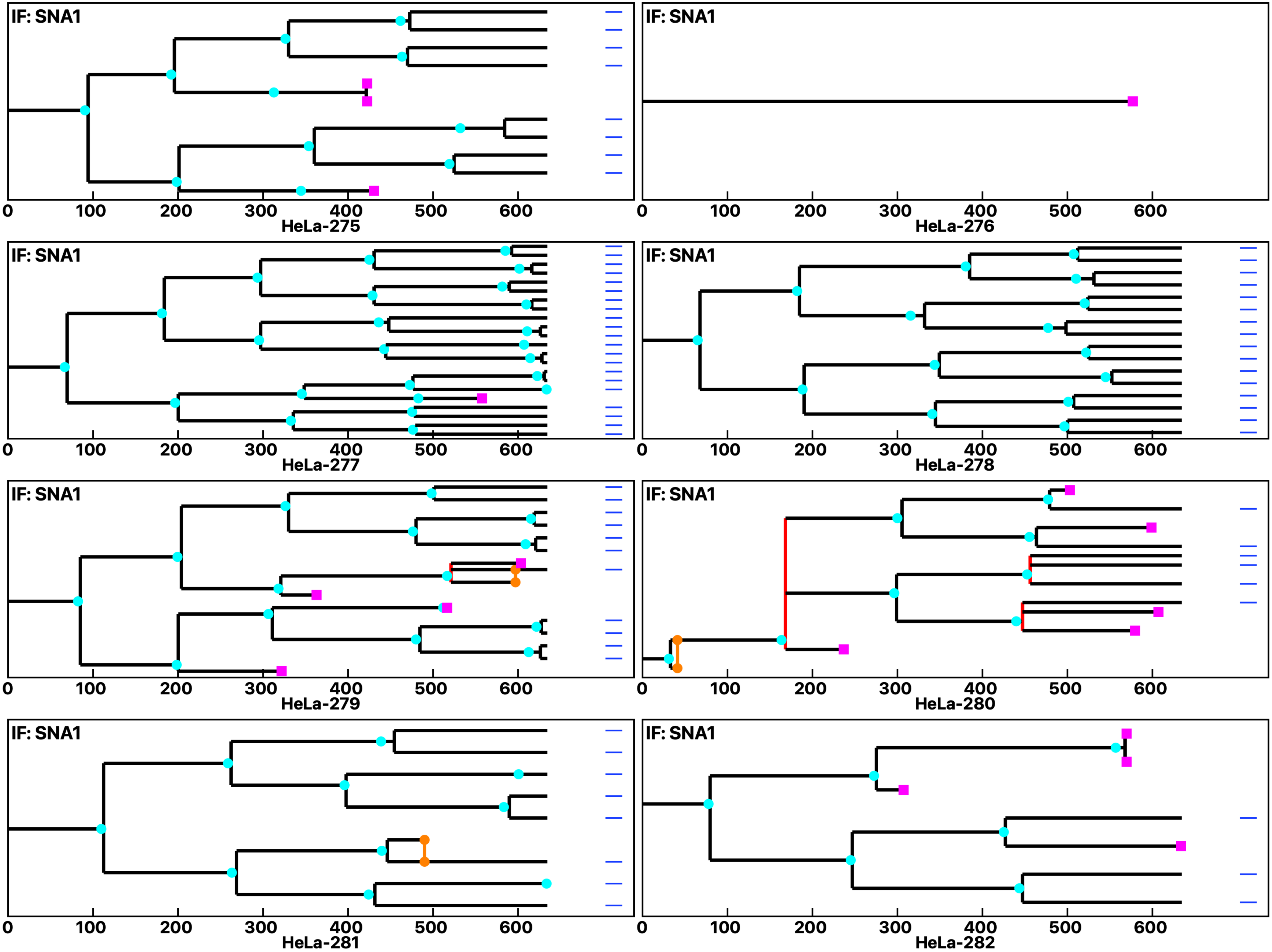

Analysis: HeLa, Treat.: HeLa, Cell: HeLa

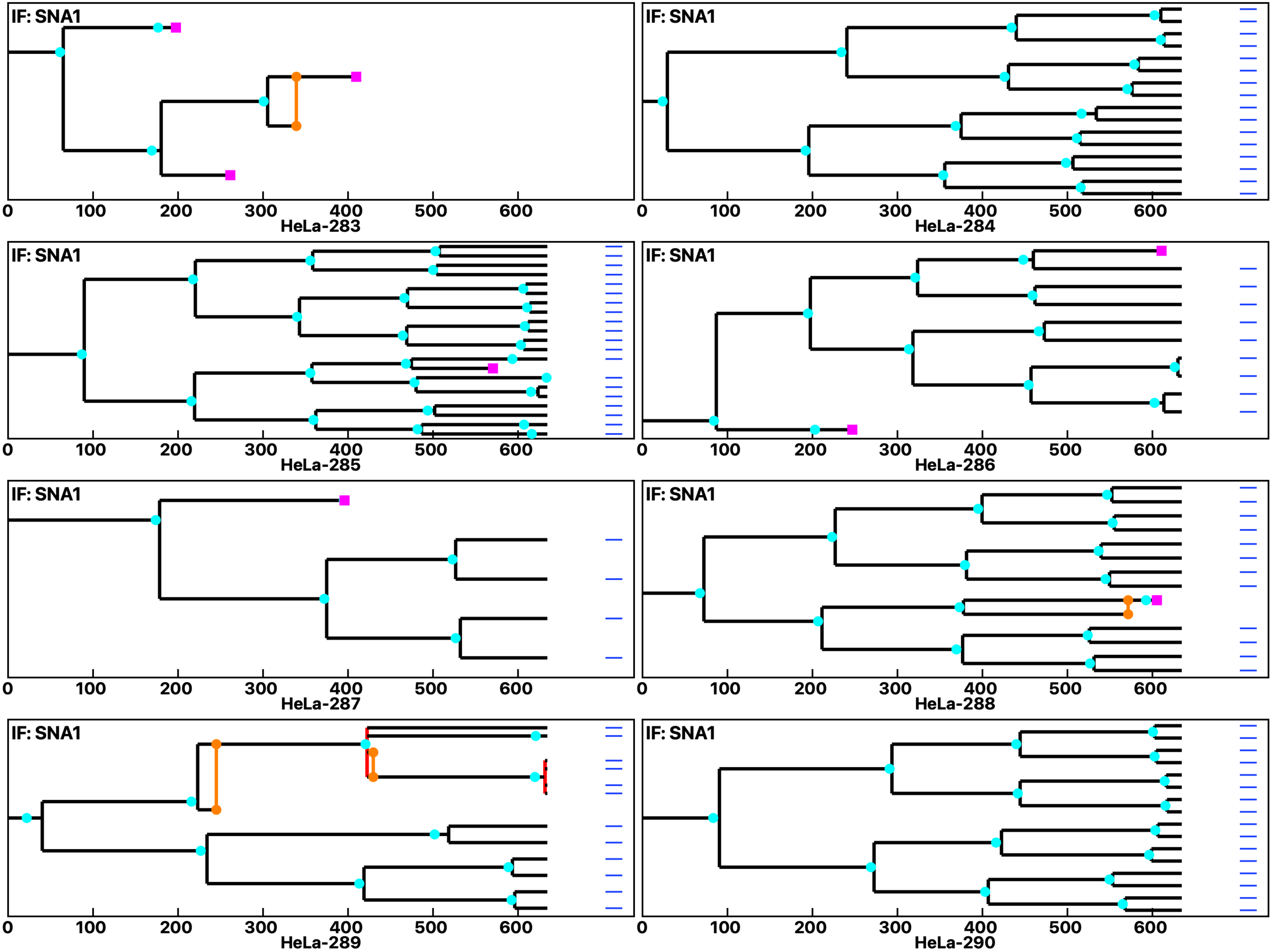

Analysis: HeLa, Treat.: HeLa, Cell: HeLa

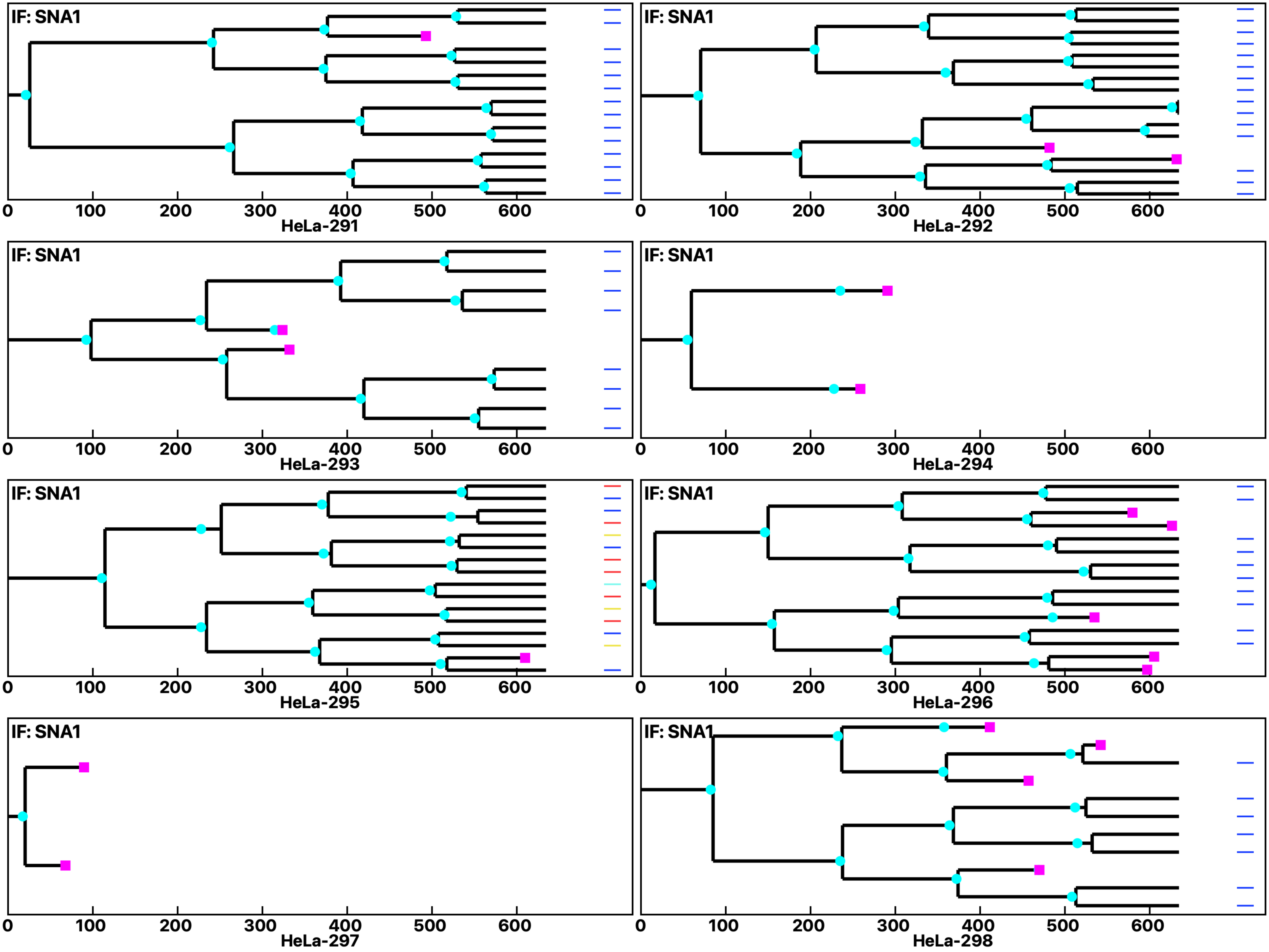

Analysis: HeLa, Treat.: HeLa, Cell: HeLa

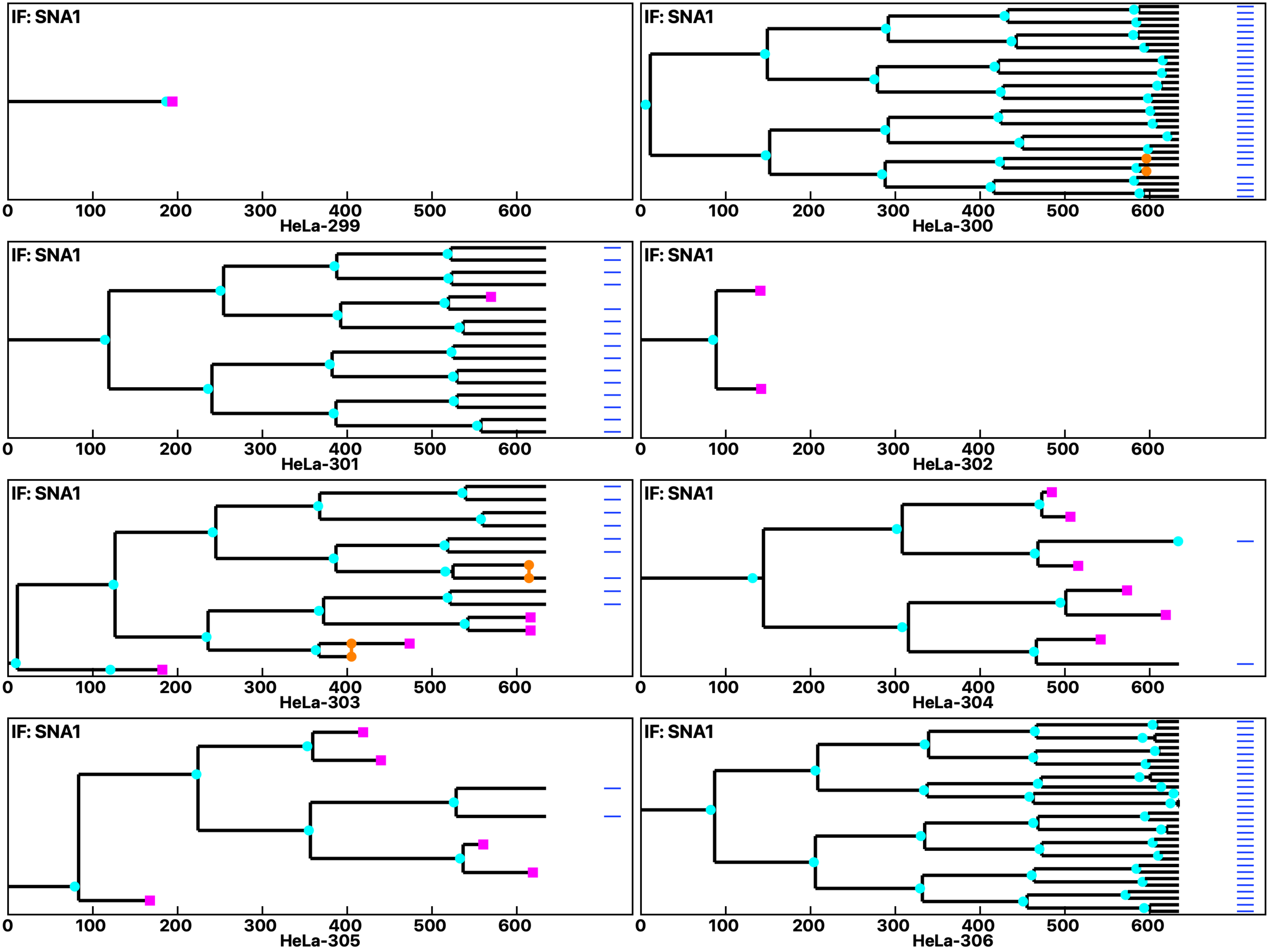

Analysis: HeLa, Treat.: HeLa, Cell: HeLa

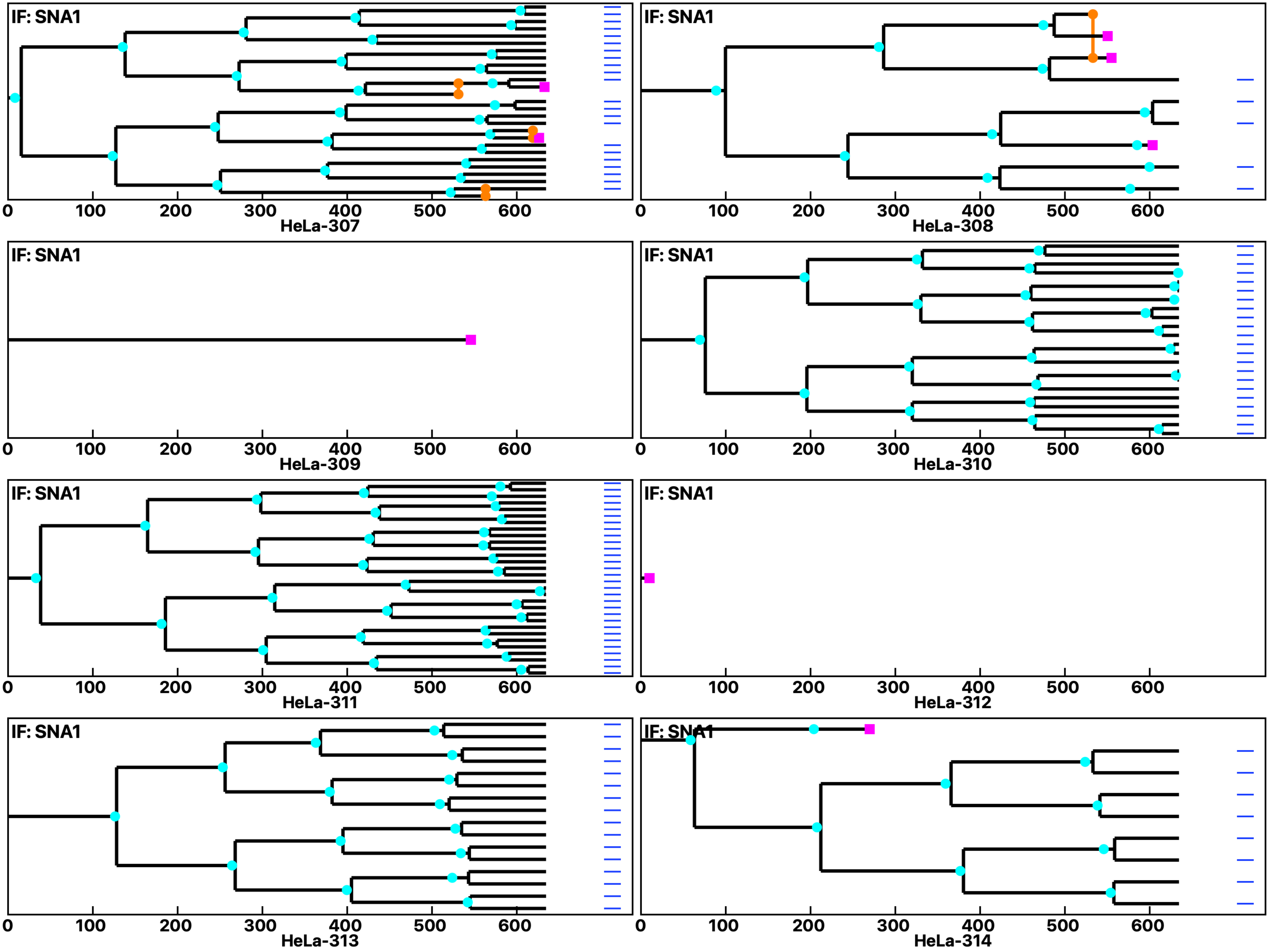

Analysis: HeLa, Treat.: HeLa, Cell: HeLa

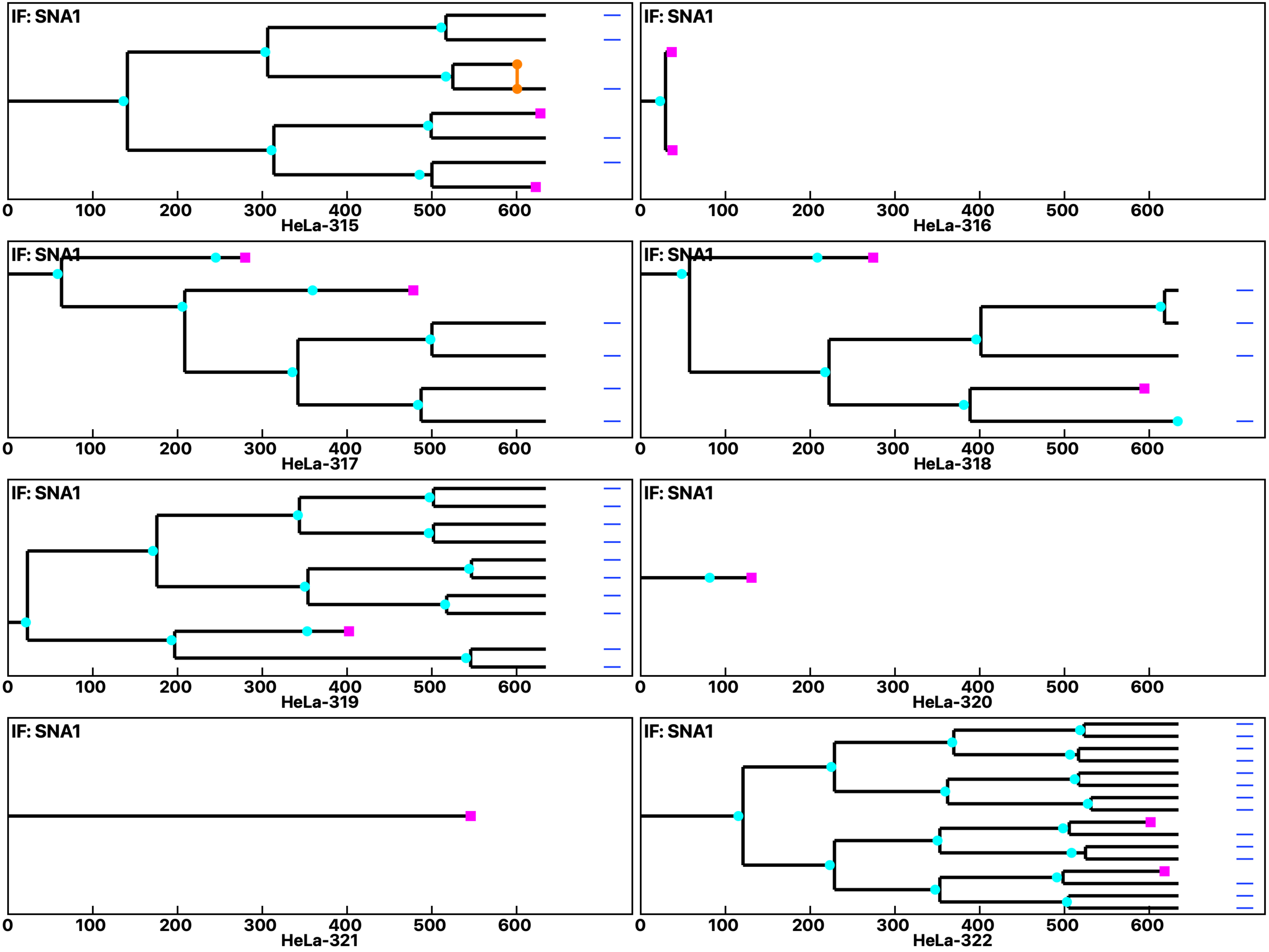

Analysis: HeLa, Treat.: HeLa, Cell: HeLa

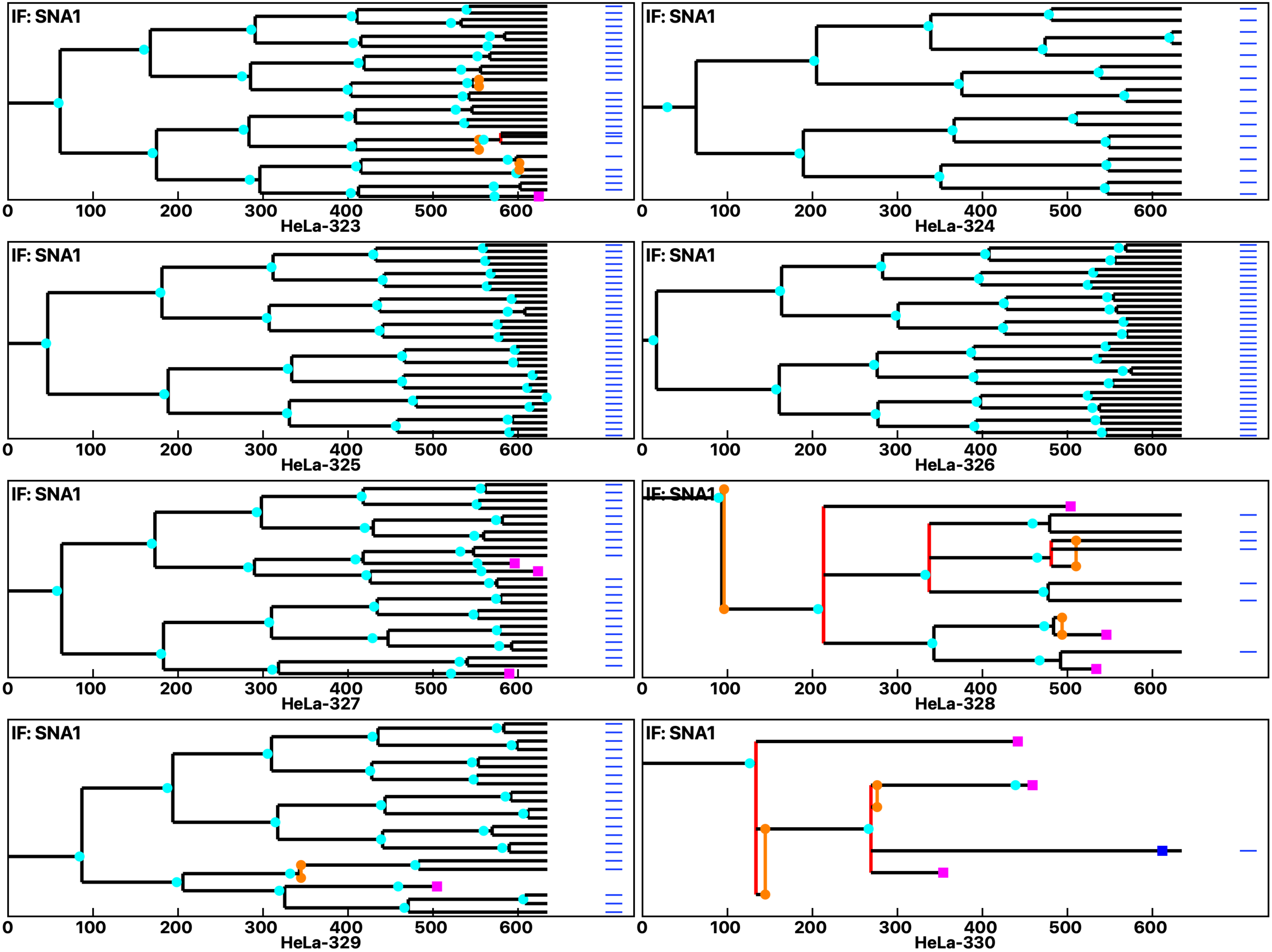

Analysis: HeLa, Treat.: HeLa, Cell: HeLa

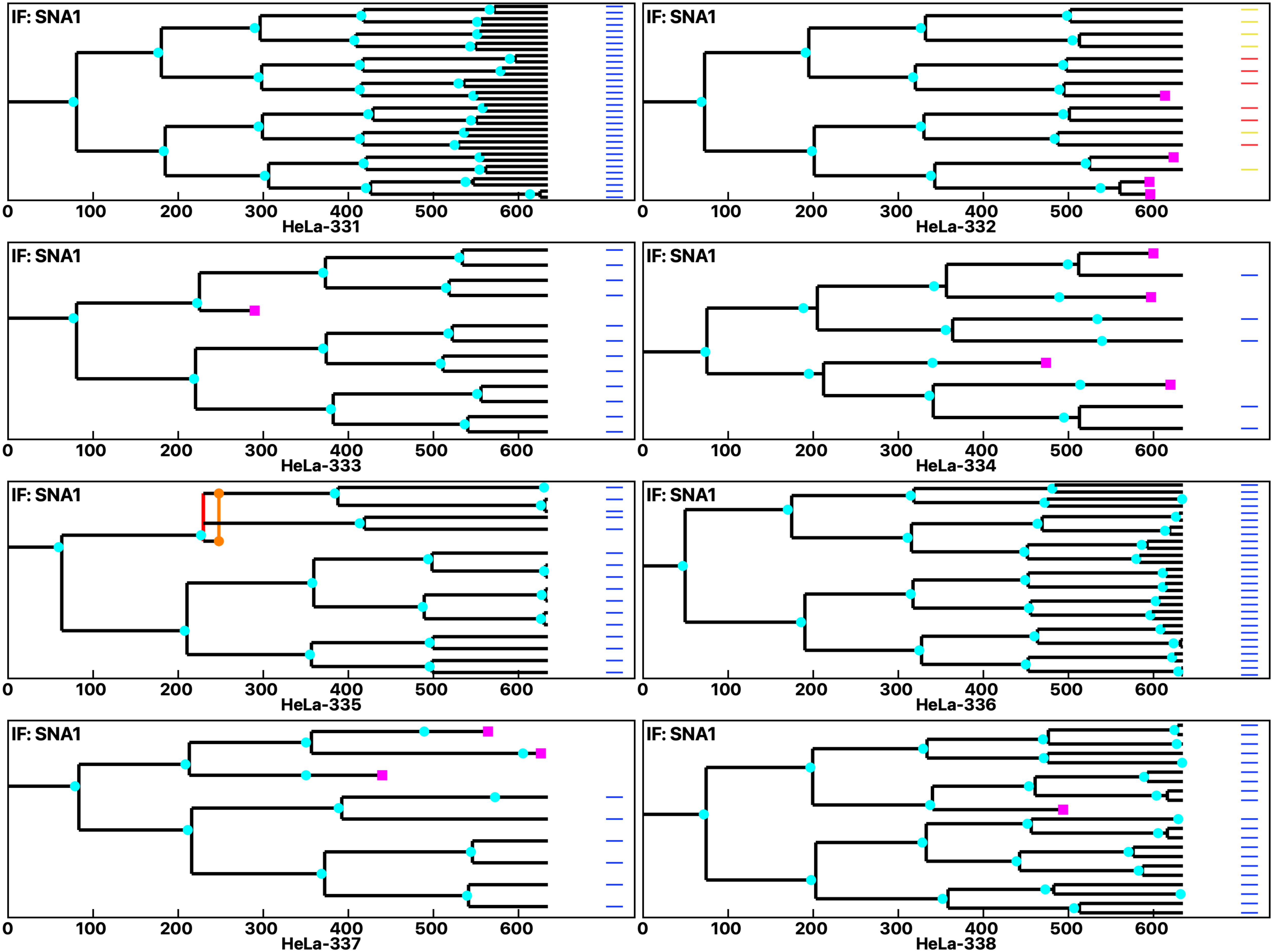

**Analysis: HeLa, Treat.: HeLa, Cell: HeLa**

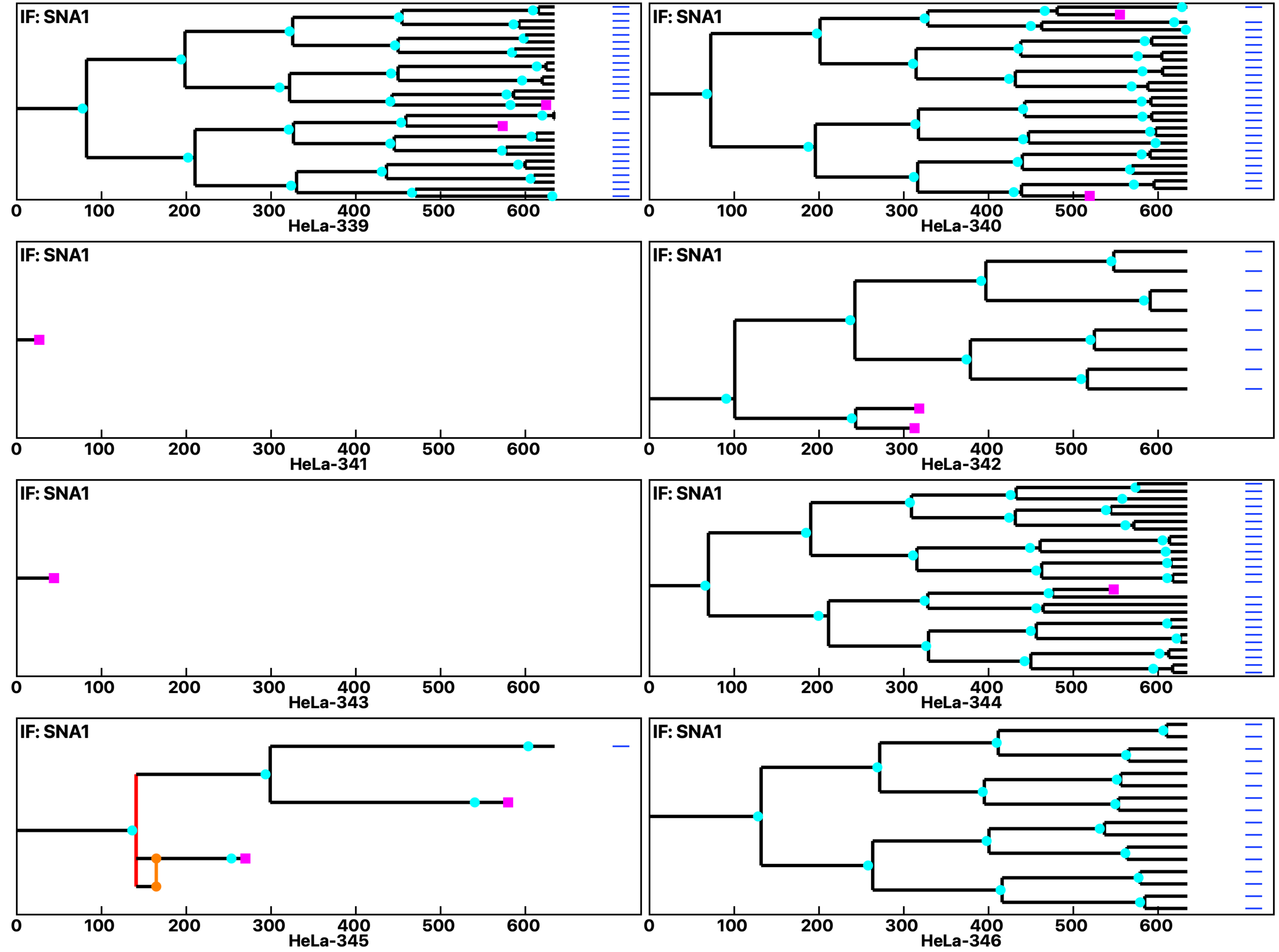

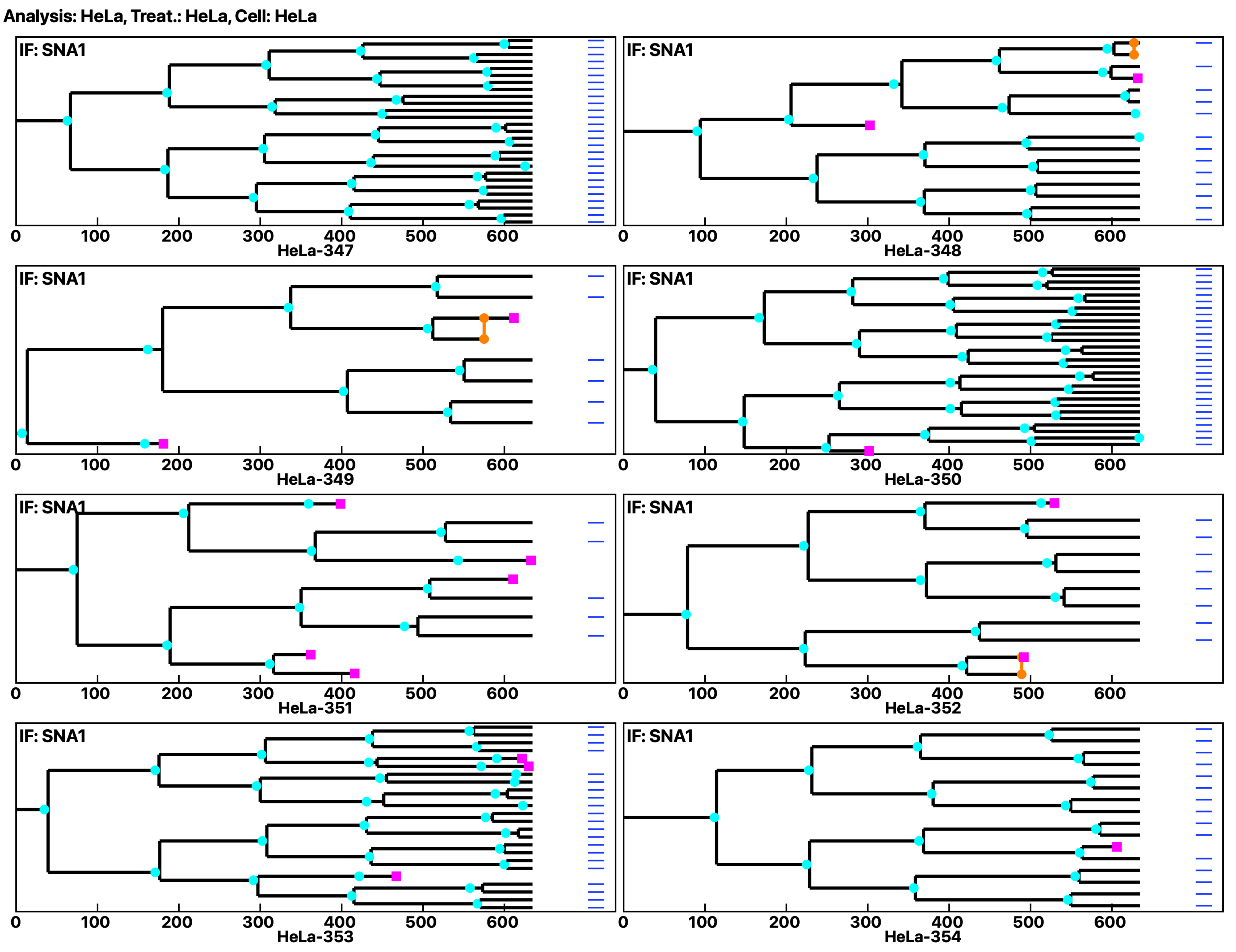

Analysis: HeLa, Treat.: HeLa, Cell: HeLa
IF: SNA1
HeLa-347
IF: SNA1
HeLa-348
IF: SNA1
HeLa-349
IF: SNA1
HeLa-350
IF: SNA1
HeLa-351
IF: SNA1
HeLa-352
IF: SNA1
HeLa-353
IF: SNA1
HeLa-354
0
100
200
300
400
500
600

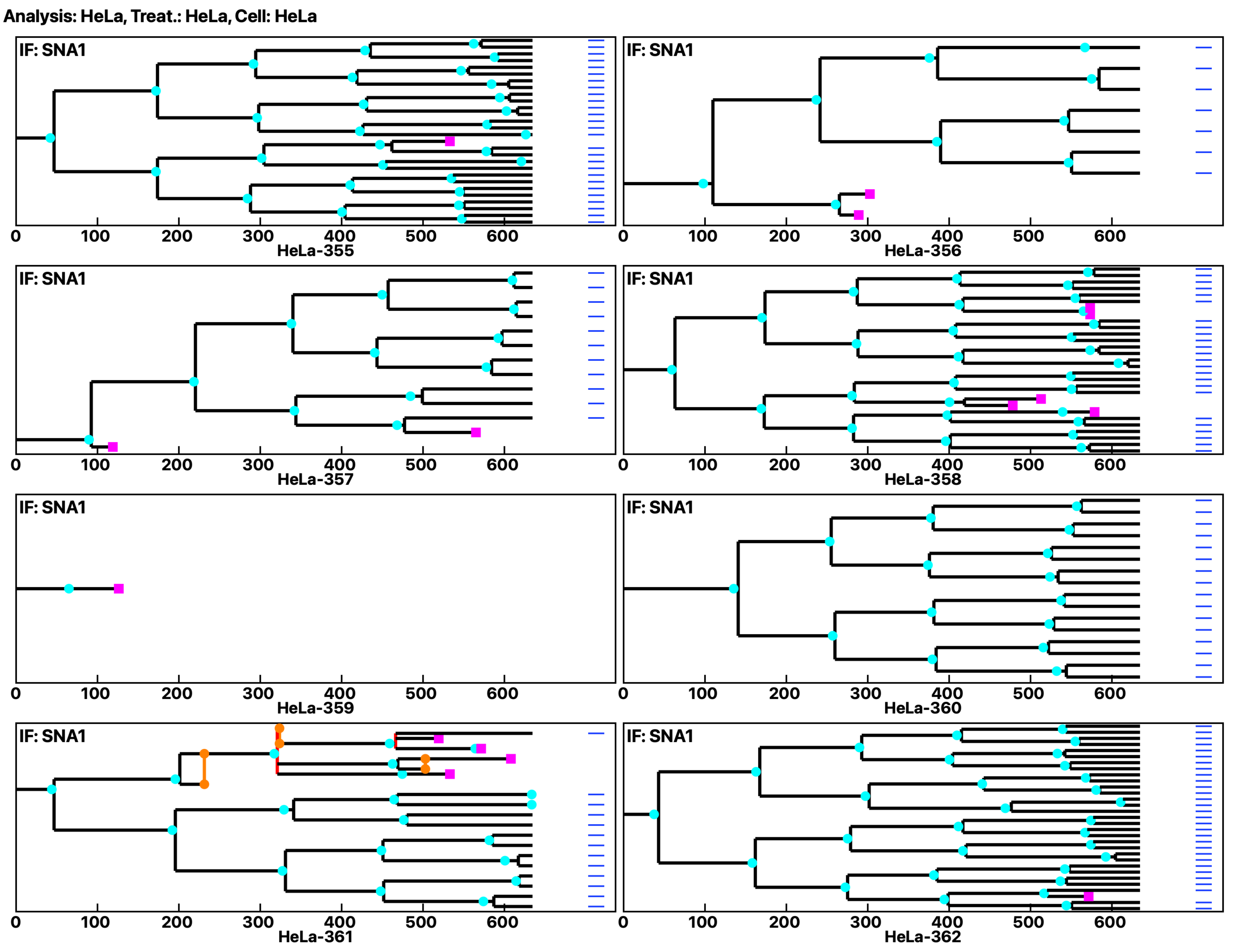
Analysis: HeLa, Treat.: HeLa, Cell: HeLa
IF: SNA1
0 100 200 300 400 500 600
HeLa-355
HeLa-356
HeLa-357
HeLa-358
HeLa-359
HeLa-360
HeLa-361
HeLa-362

Analysis: HeLa, Treat.: HeLa, Cell: HeLa

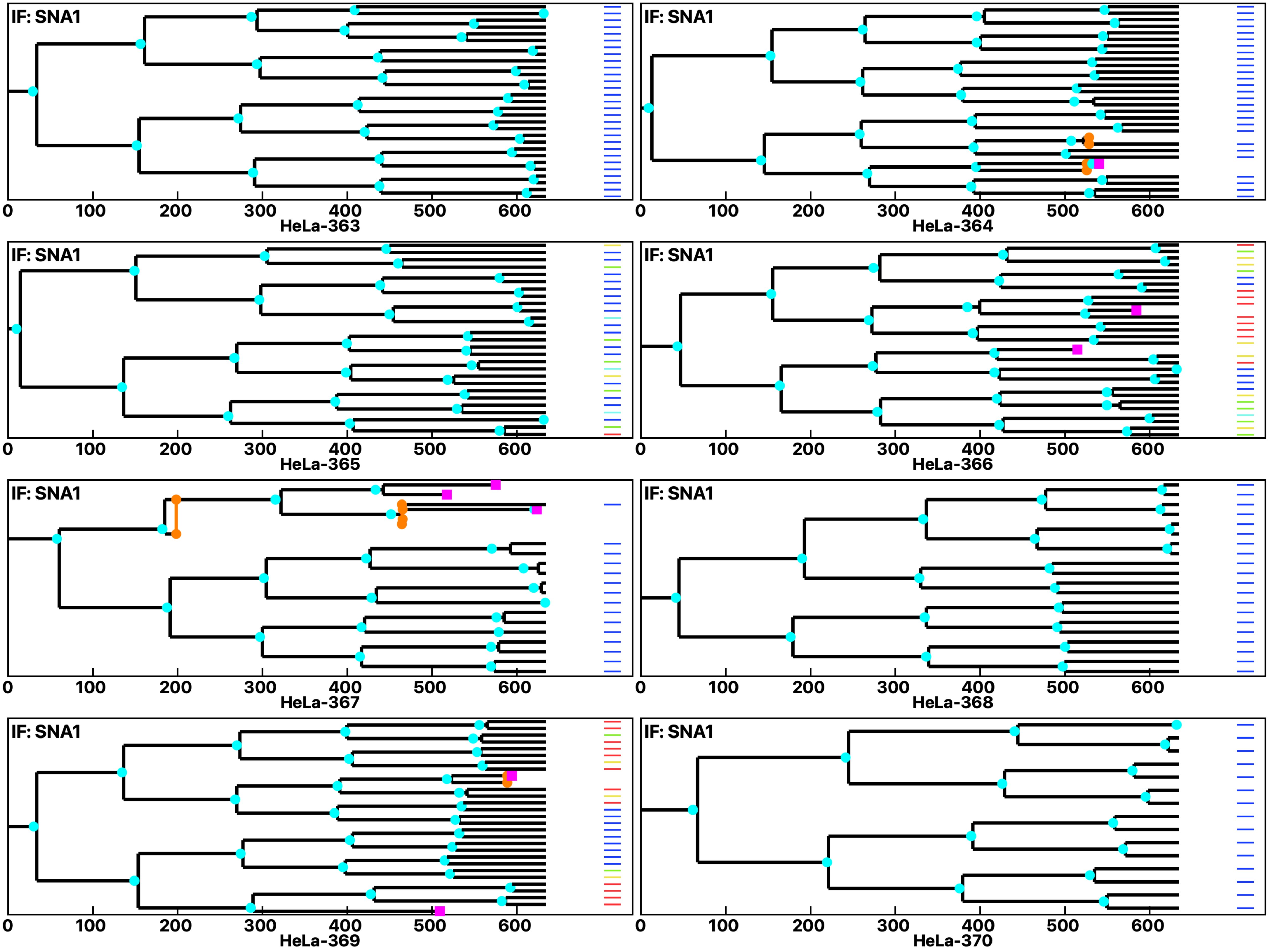

Analysis: HeLa, Treat.: HeLa, Cell: HeLa

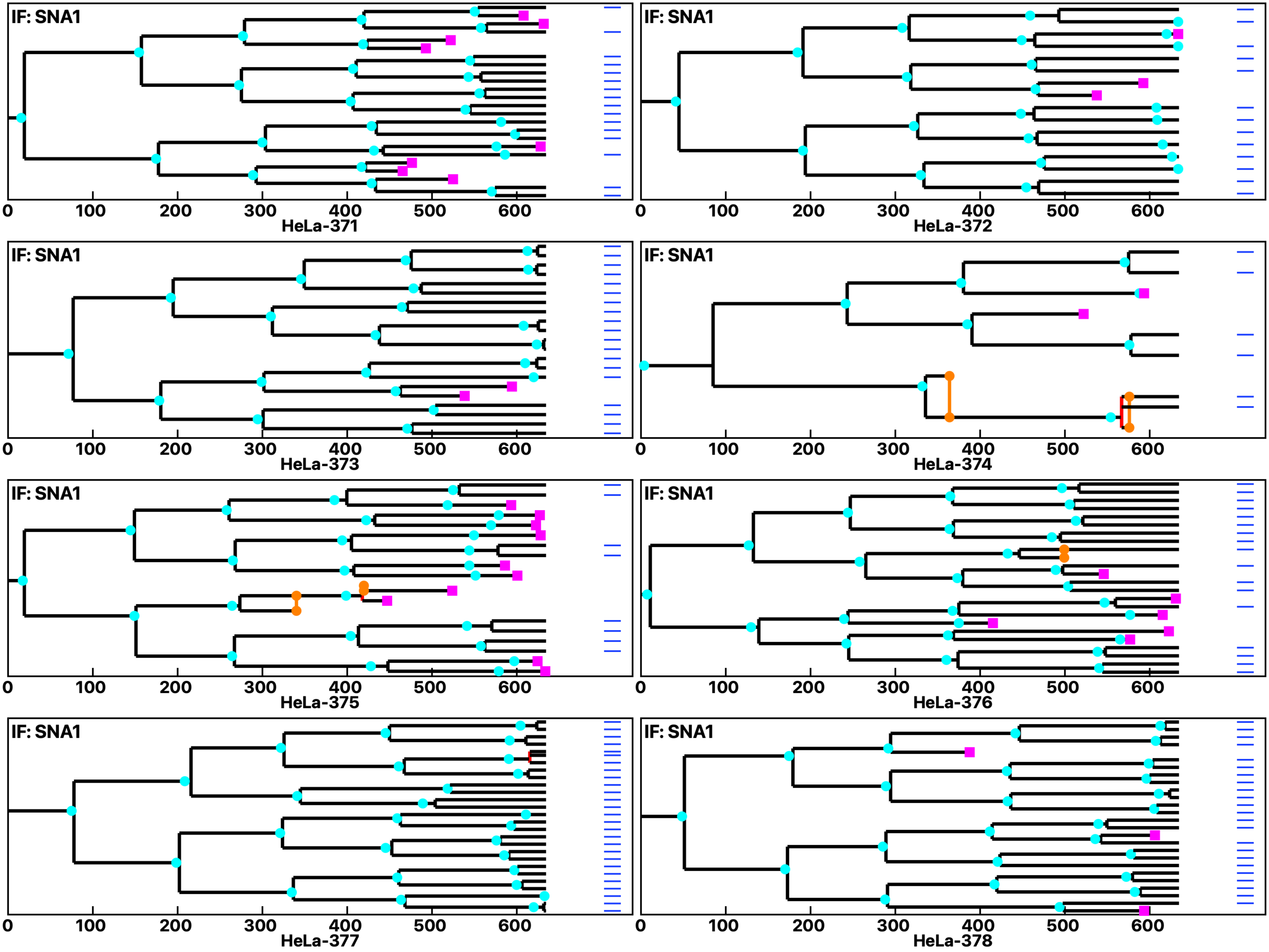

**Analysis: HeLa, Treat.: HeLa, Cell: HeLa**

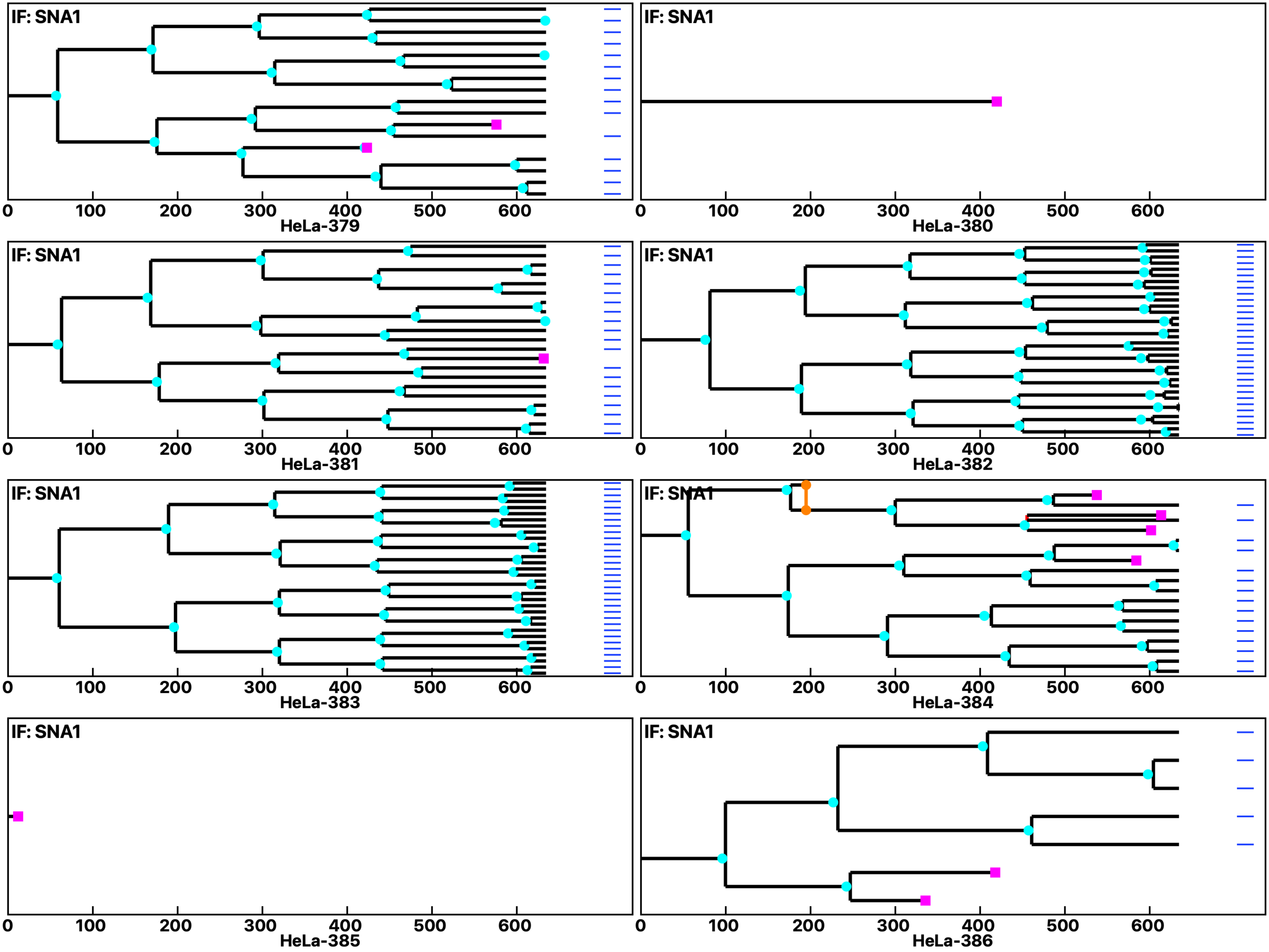

Analysis: HeLa, Treat.: HeLa, Cell: HeLa

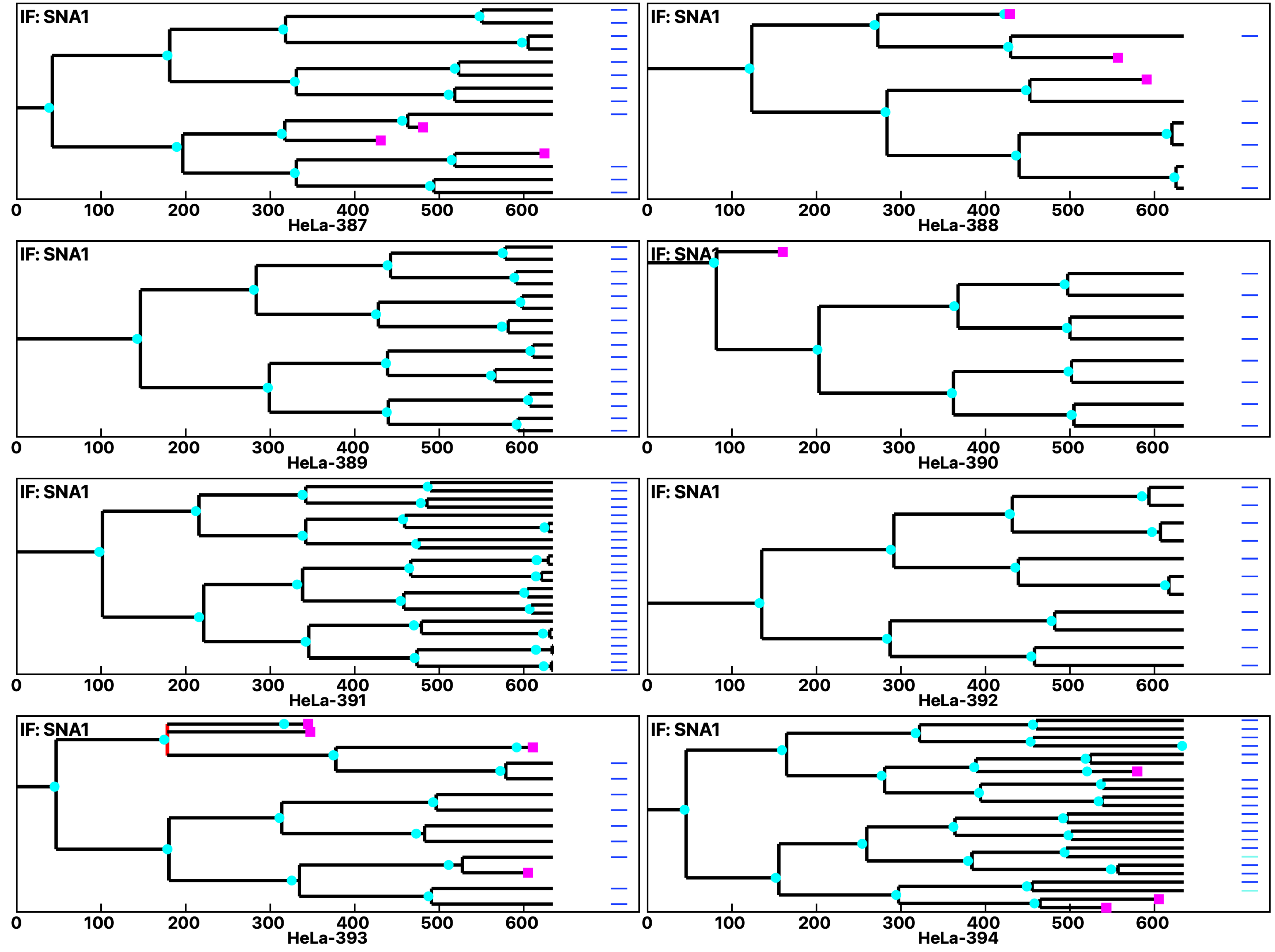

**Analysis: HeLa, Treat.: HeLa, Cell: HeLa**

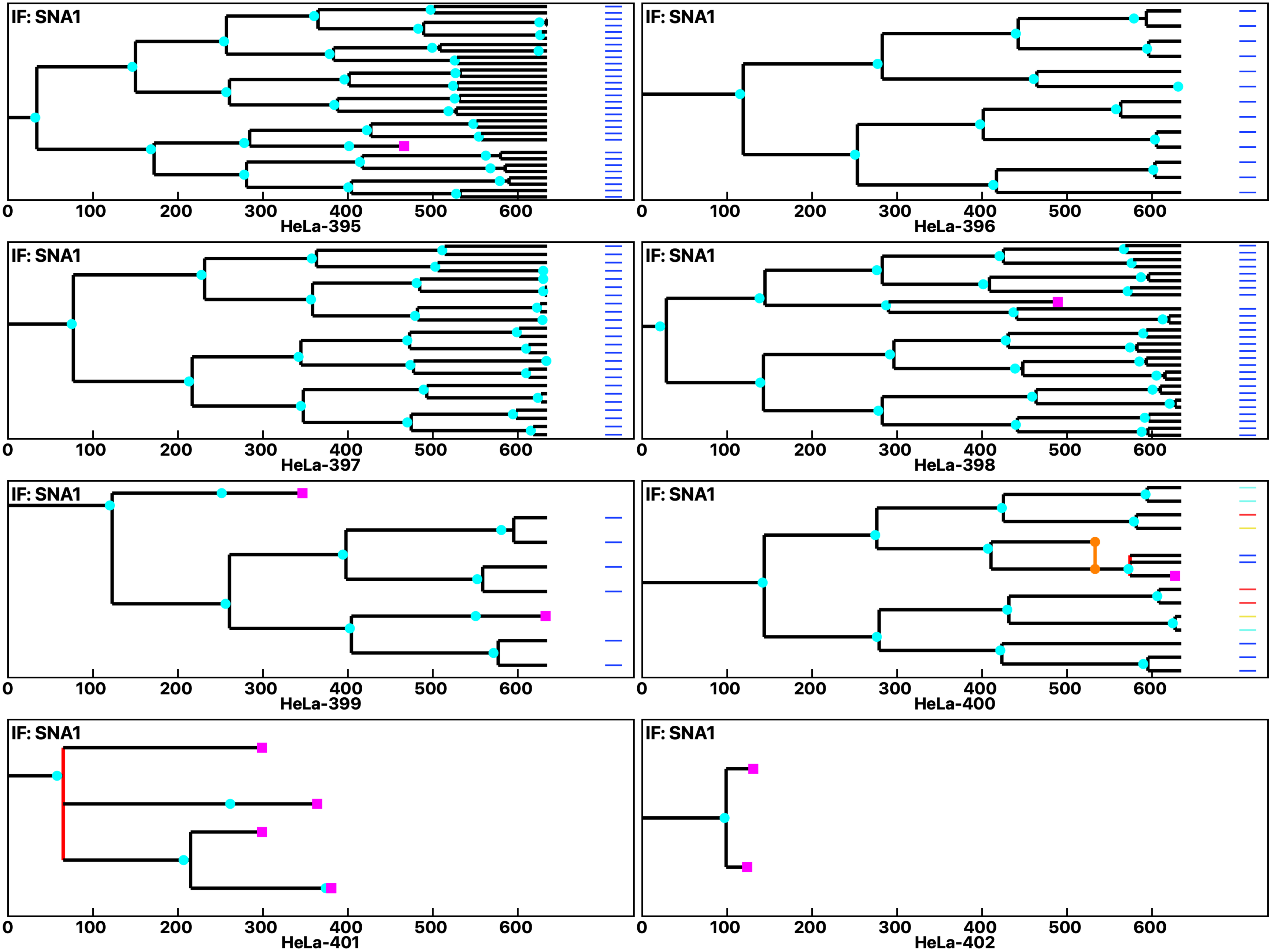

Analysis: HeLa, Treat.: HeLa, Cell: HeLa

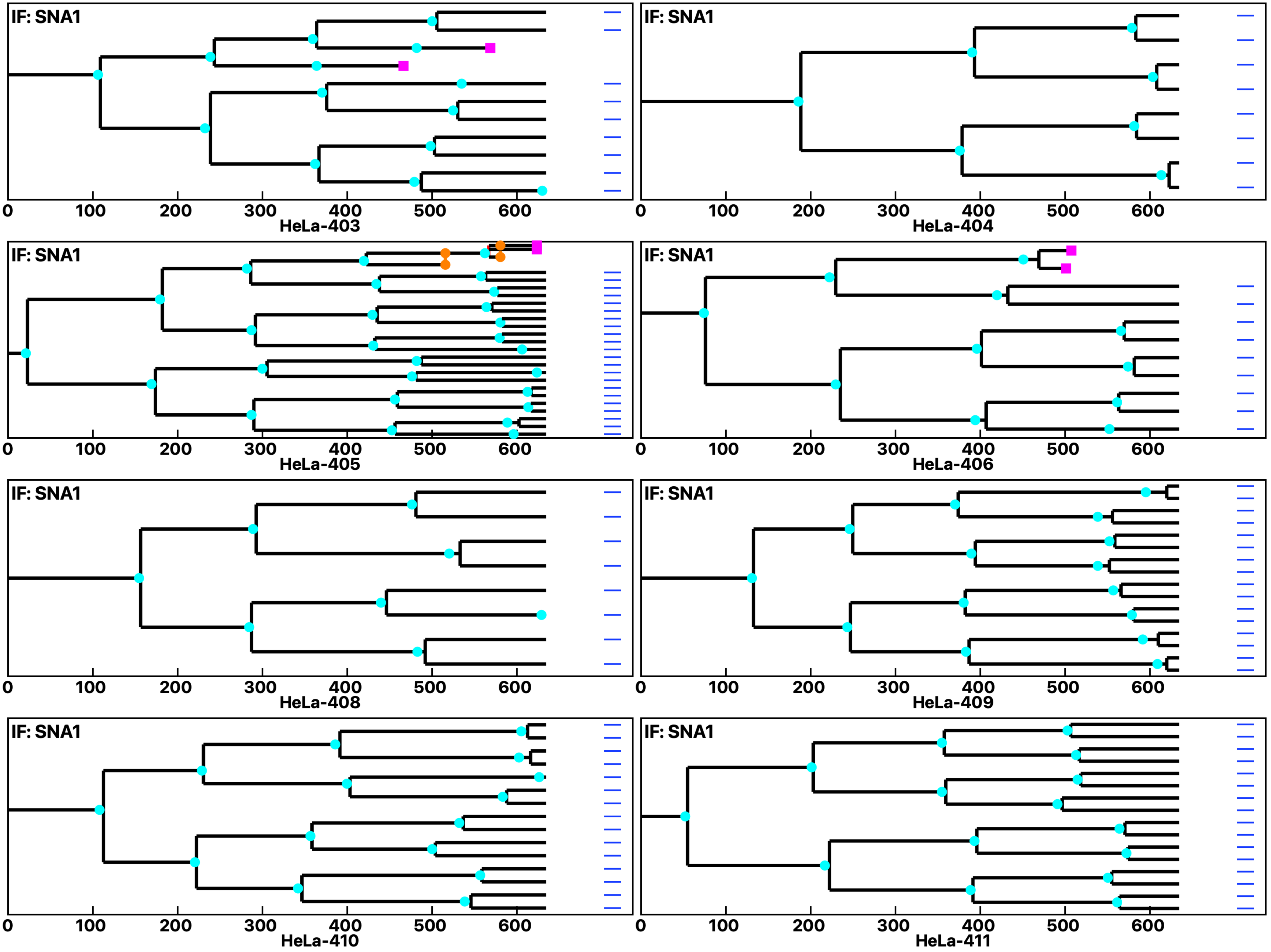

Analysis: HeLa, Treat.: HeLa, Cell: HeLa

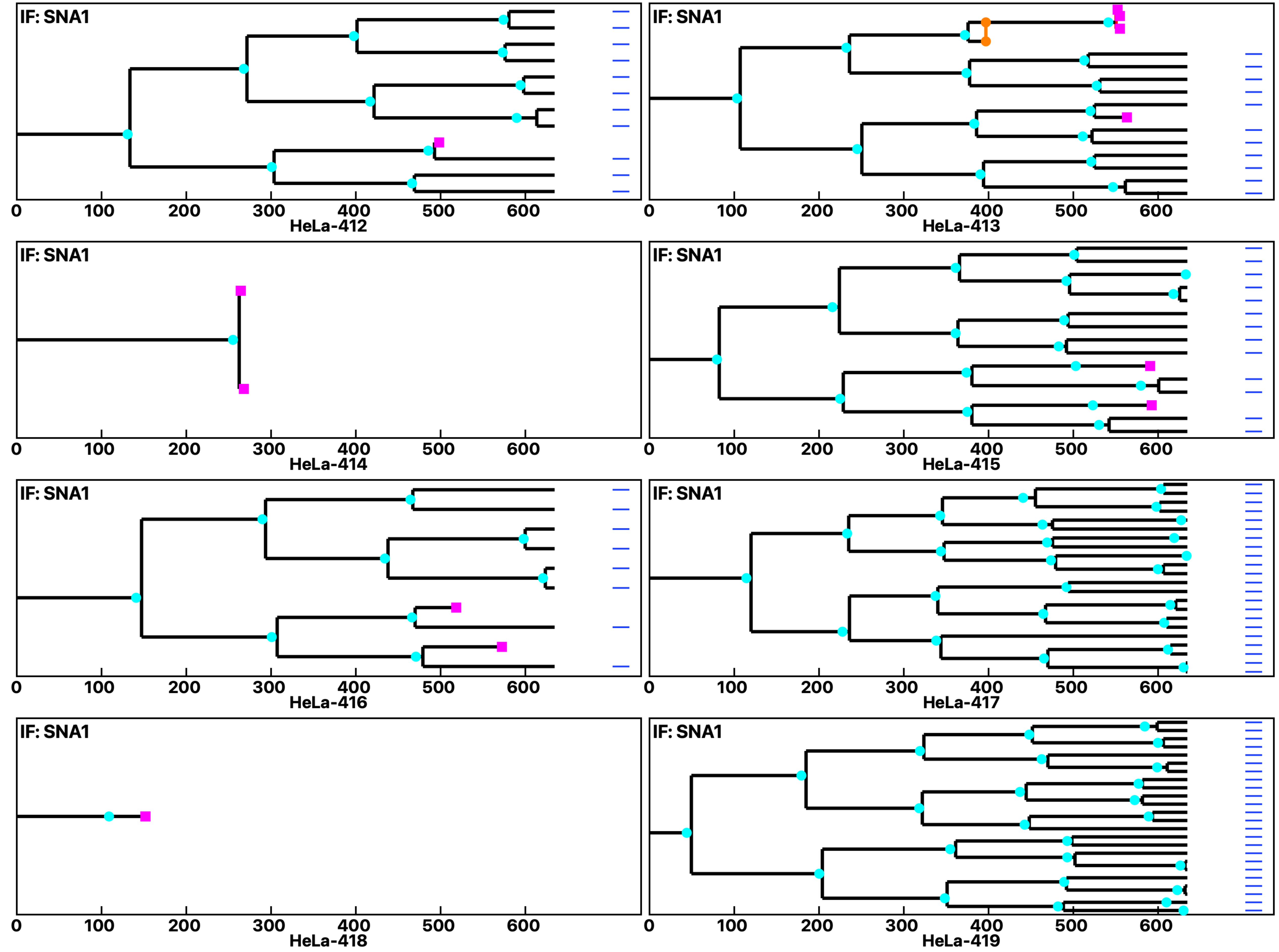

Analysis: HeLa, Treat.: HeLa, Cell: HeLa

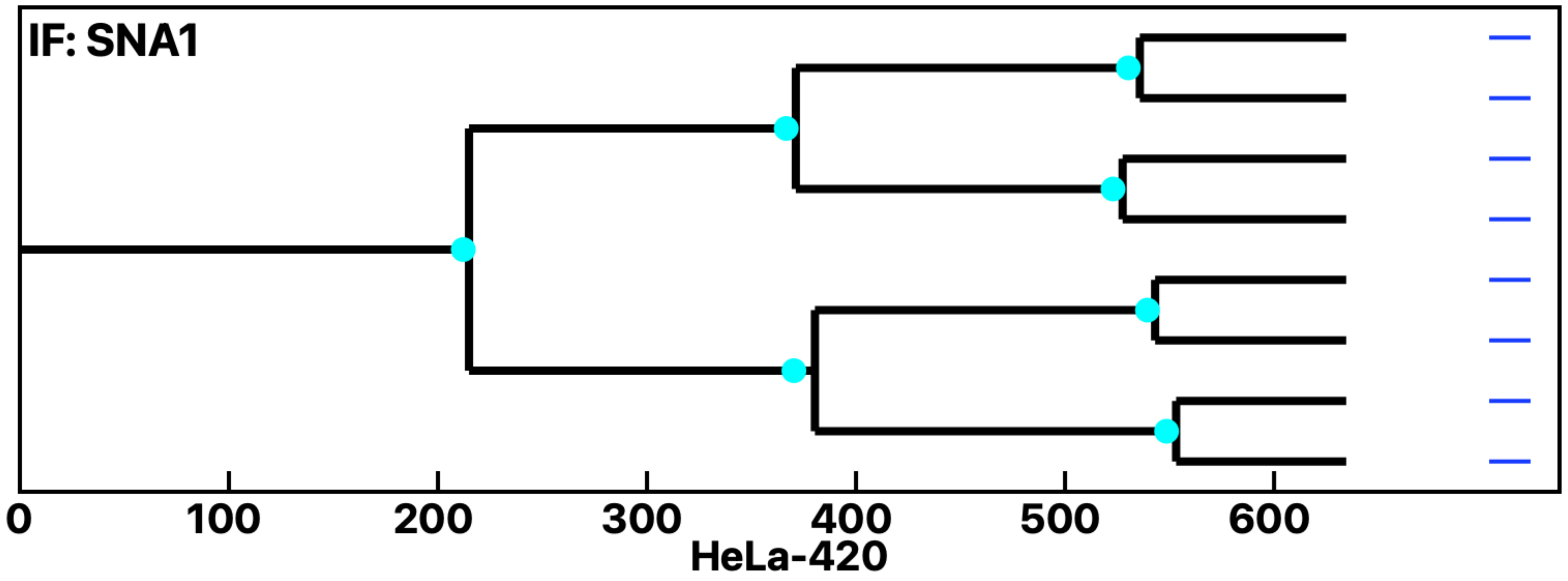
