## Supplementary Data 2 for "Cell Fate Simulation Reveals Cancer Cell Features in the Tumor Microenvironment"

Analysis: MiaPaCa2, Treat.: MiaPaCa2, Cell: MiaPaCa2

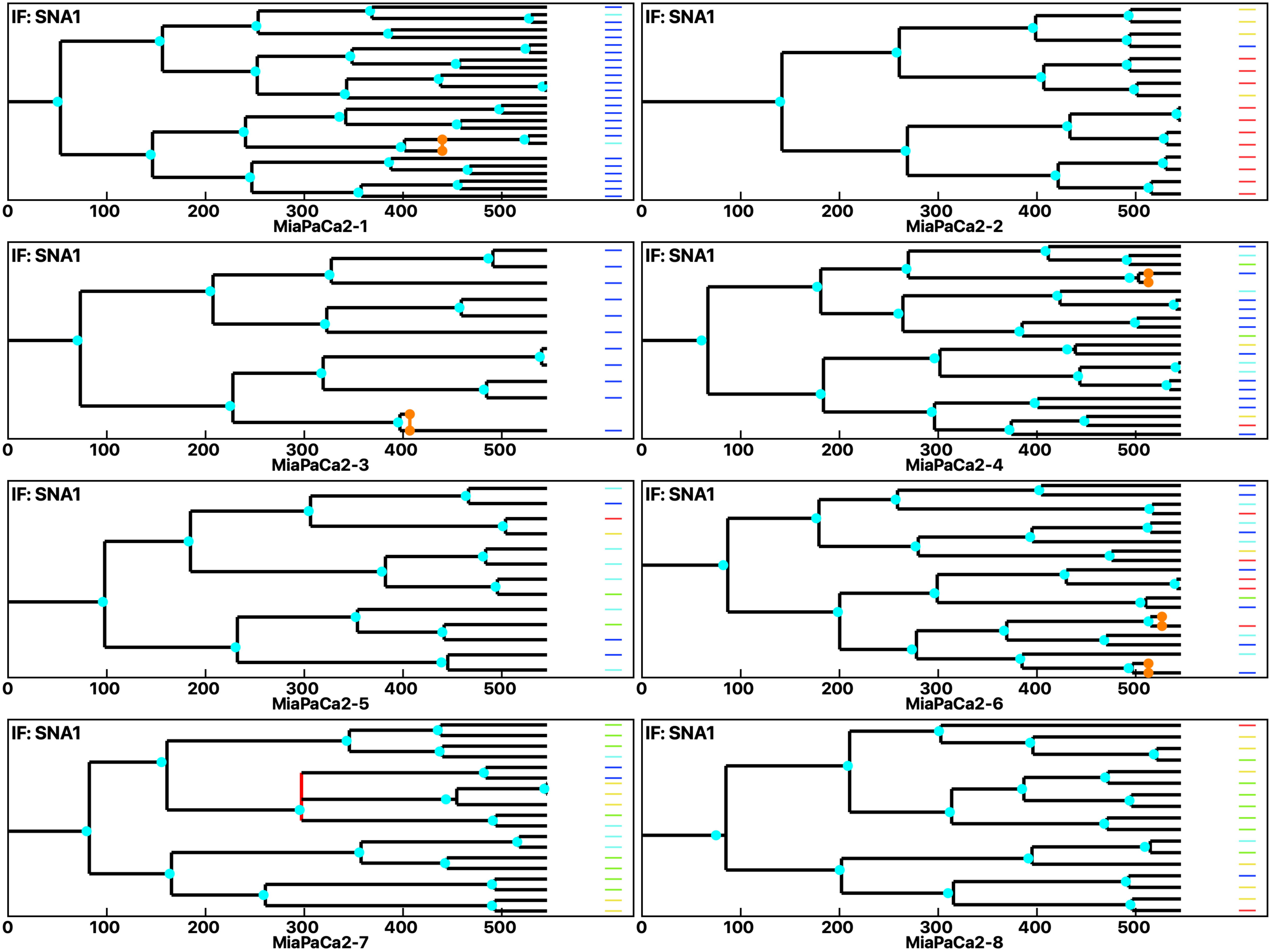

Analysis: MiaPaCa2, Treat.: MiaPaCa2, Cell: MiaPaCa2

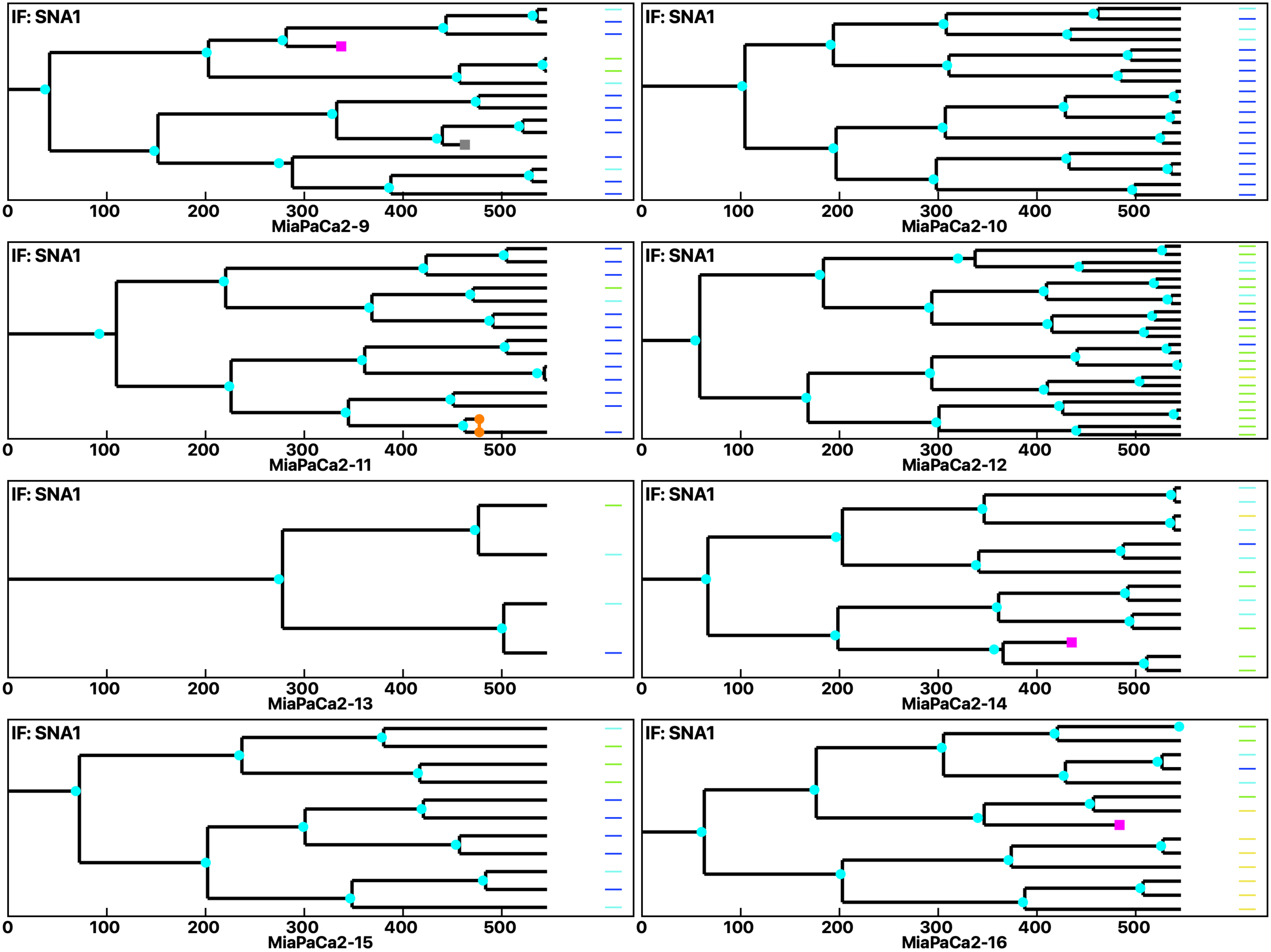

Analysis: MiaPaCa2, Treat.: MiaPaCa2, Cell: MiaPaCa2

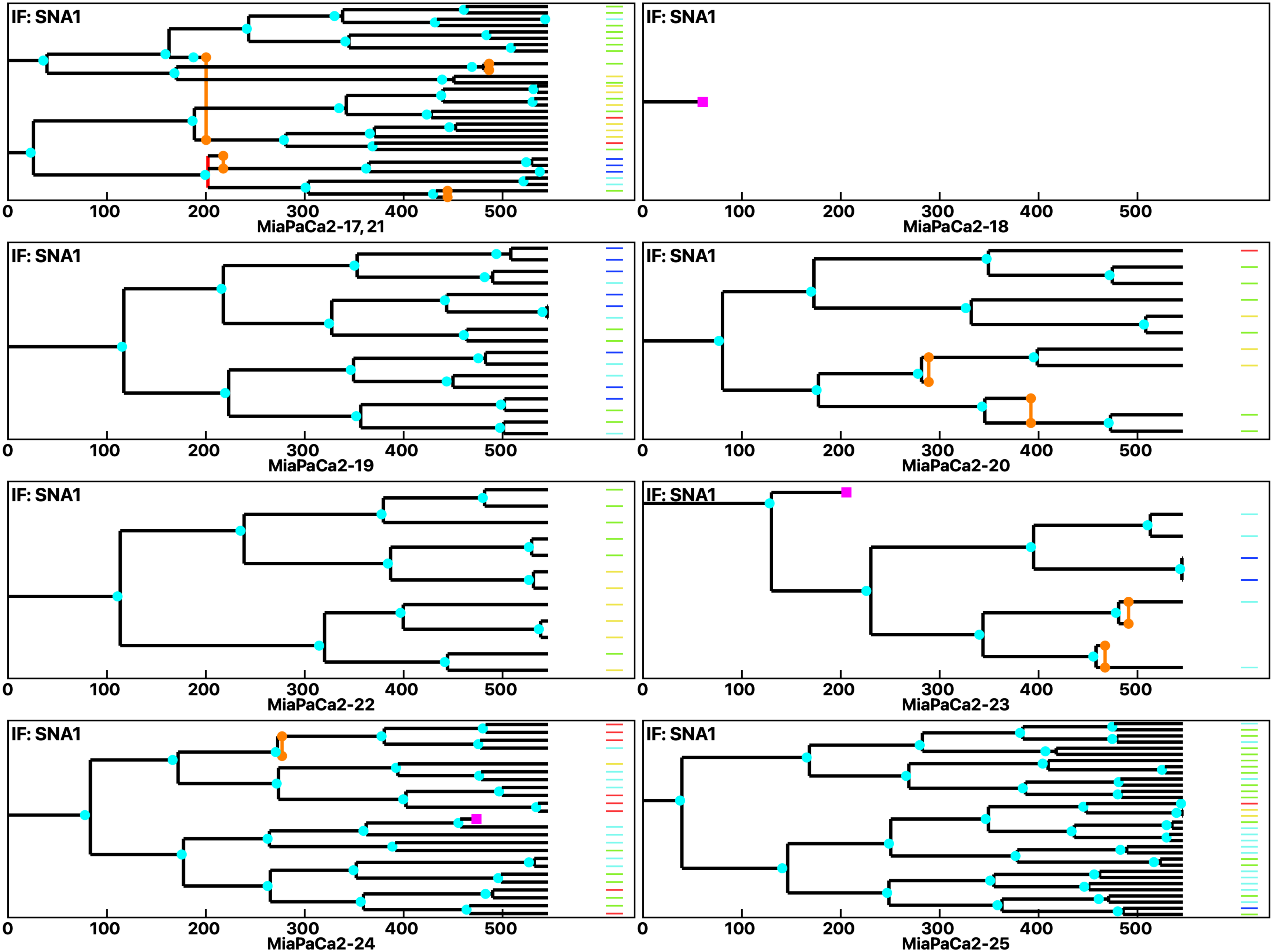

**Analysis: MiaPaCa2, Treat.: MiaPaCa2, Cell: MiaPaCa2**

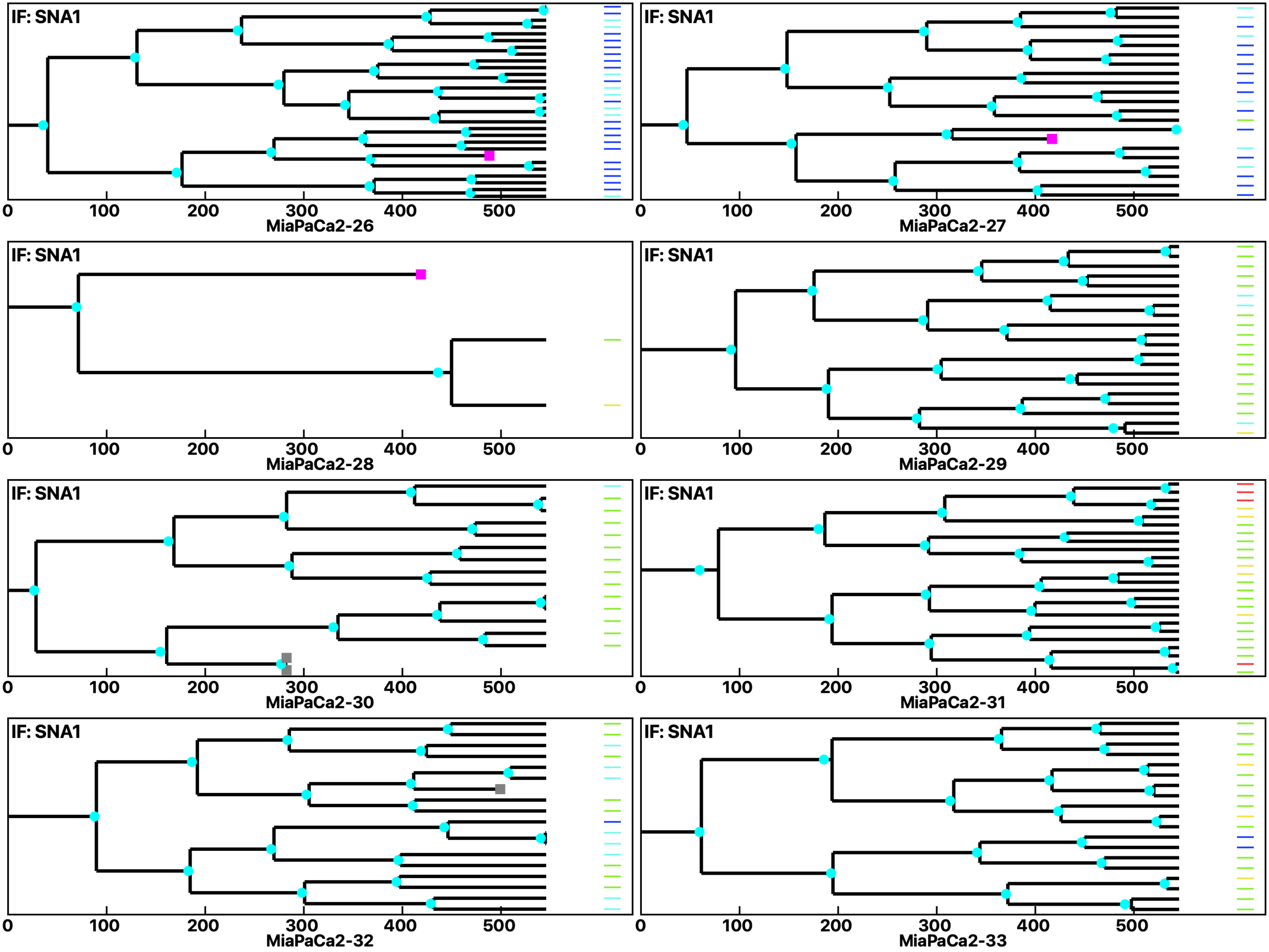

Analysis: MiaPaCa2, Treat.: MiaPaCa2, Cell: MiaPaCa2

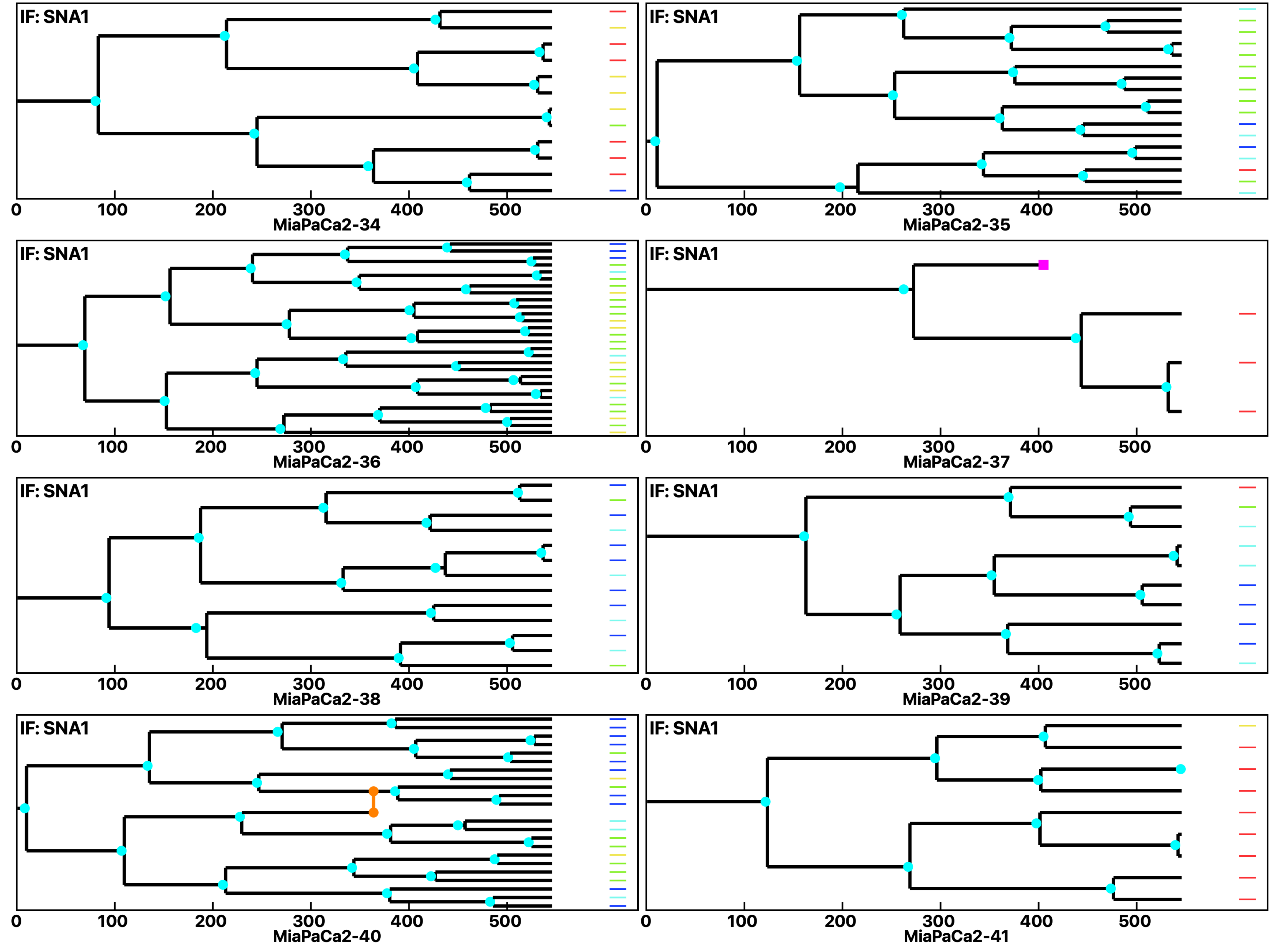

Analysis: MiaPaCa2, Treat.: MiaPaCa2, Cell: MiaPaCa2

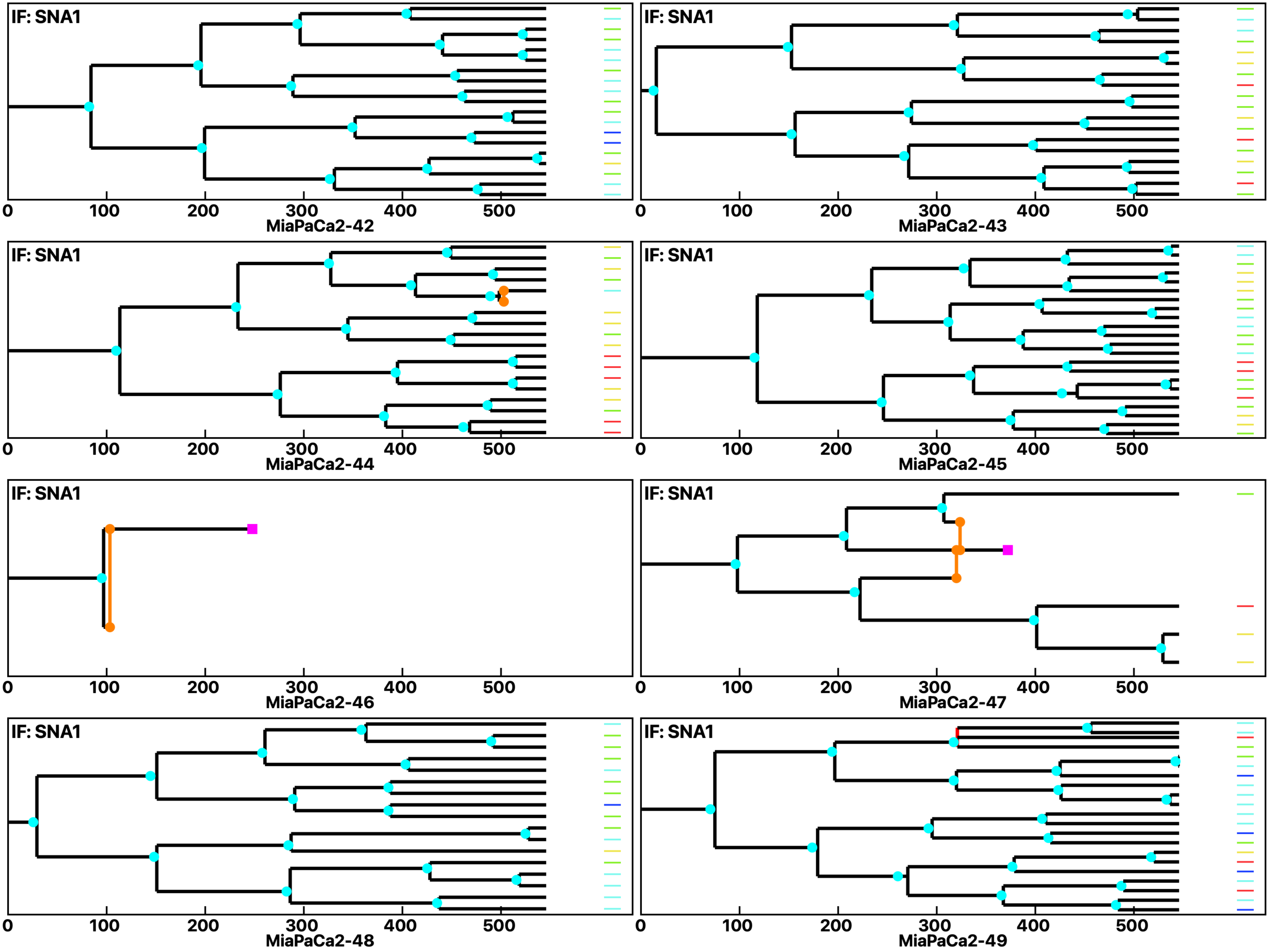

Analysis: MiaPaCa2, Treat.: MiaPaCa2, Cell: MiaPaCa2

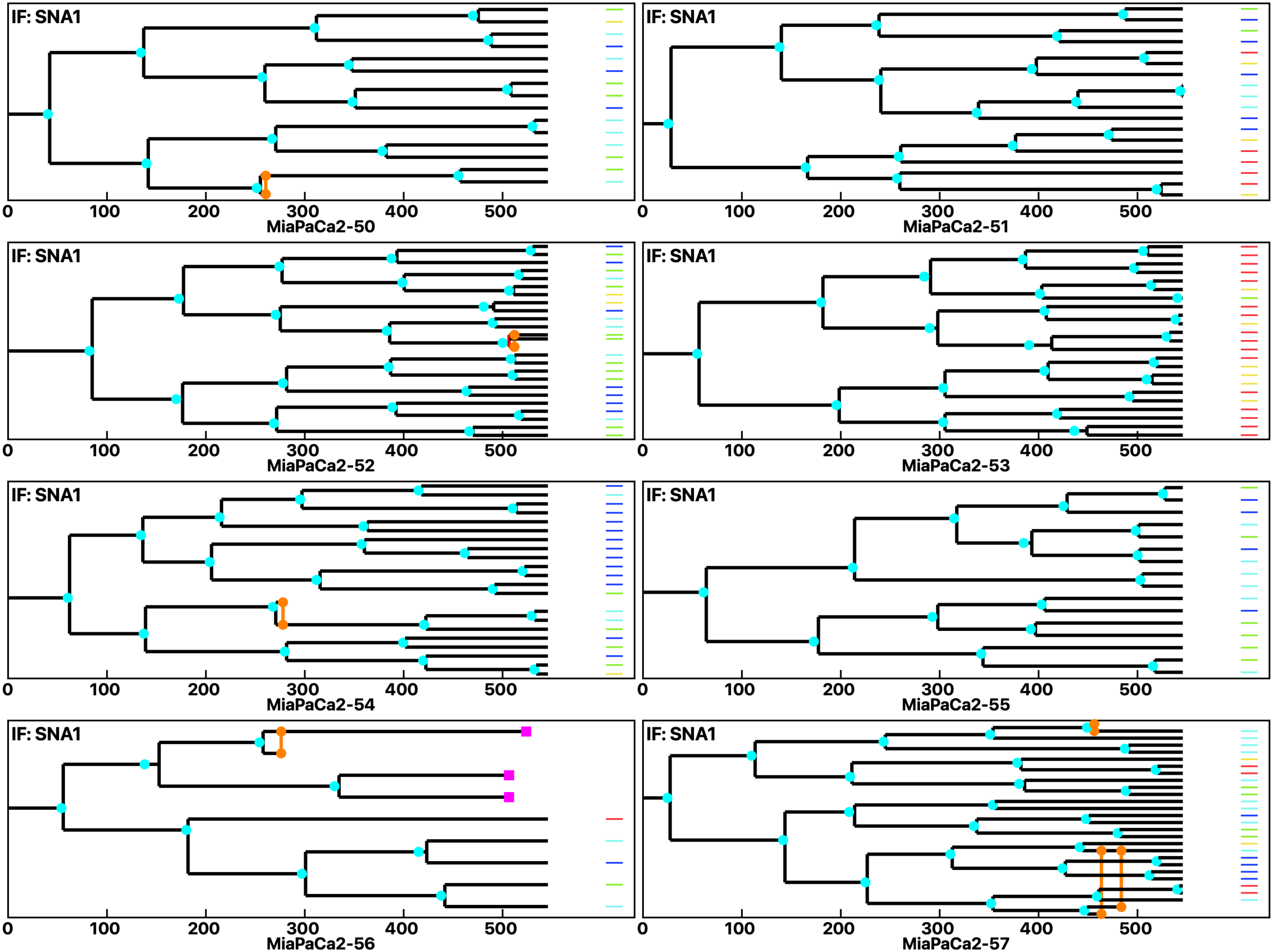

Analysis: MiaPaCa2, Treat.: MiaPaCa2, Cell: MiaPaCa2

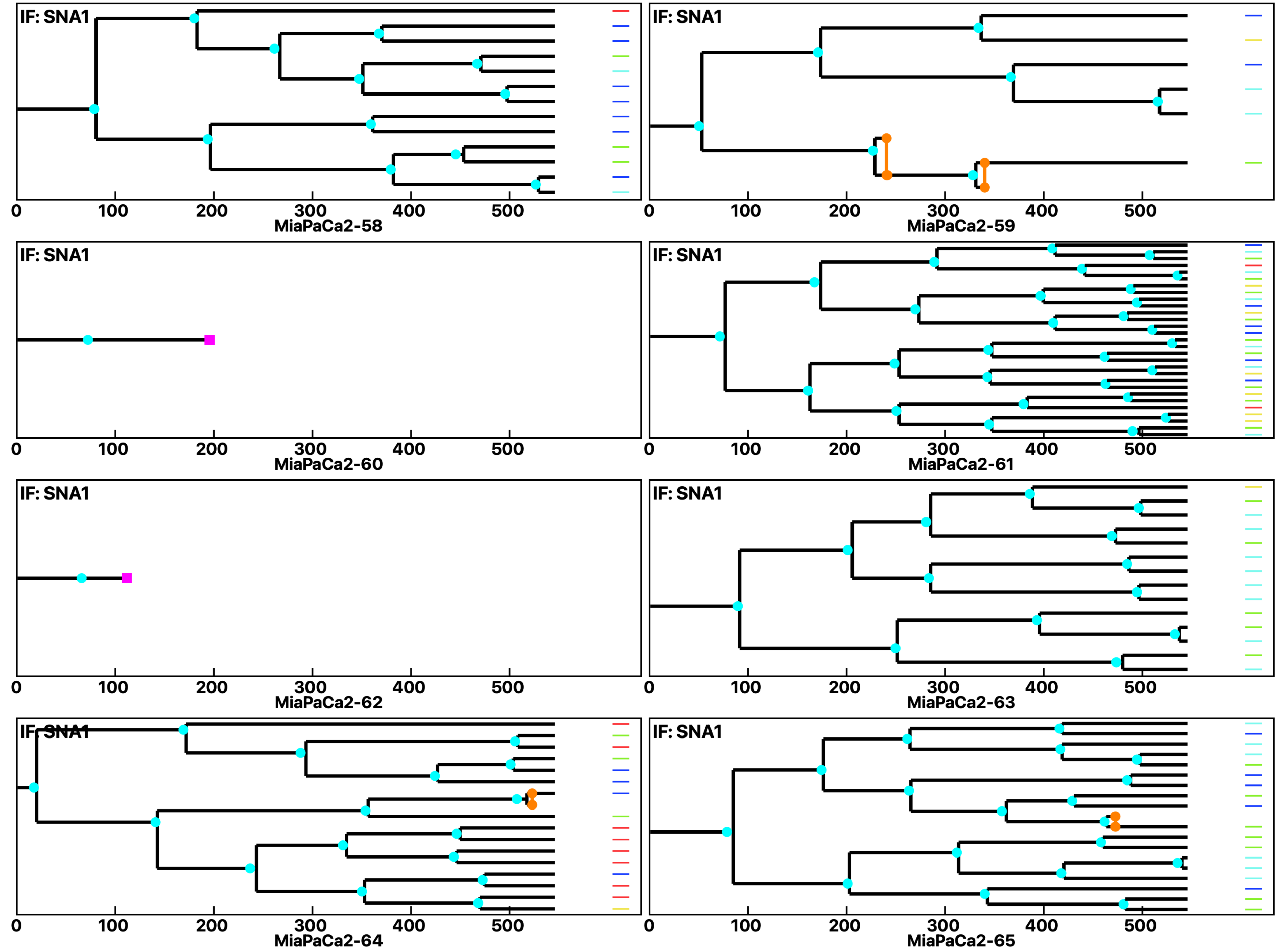

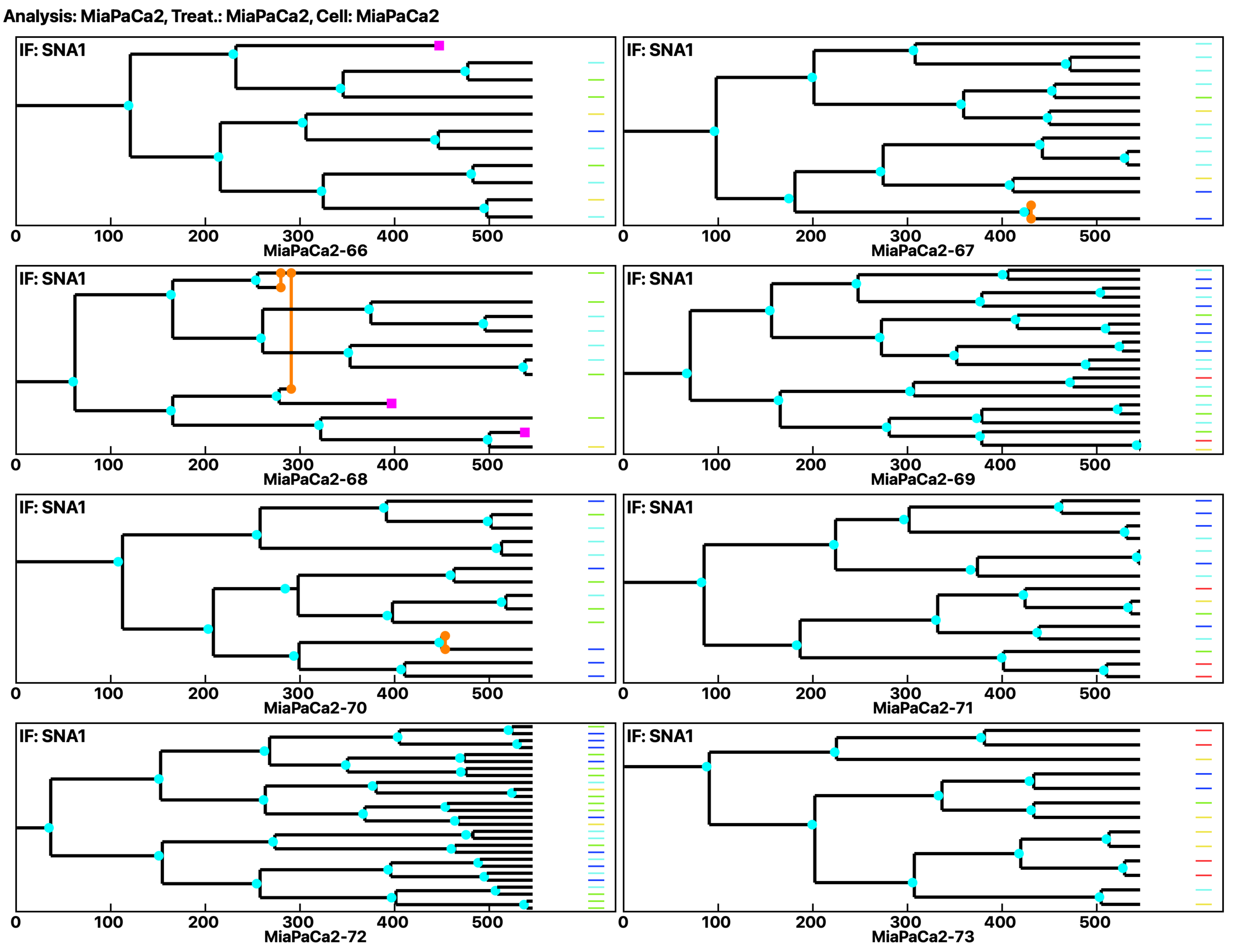
Analysis: MiaPaCa2, Treat.: MiaPaCa2, Cell: MiaPaCa2
IF: SNA1
0
100
200
300
400
500
MiaPaCa2-66
MiaPaCa2-67
MiaPaCa2-68
MiaPaCa2-69
MiaPaCa2-70
MiaPaCa2-71
MiaPaCa2-72
MiaPaCa2-73

Analysis: MiaPaCa2, Treat.: MiaPaCa2, Cell: MiaPaCa2

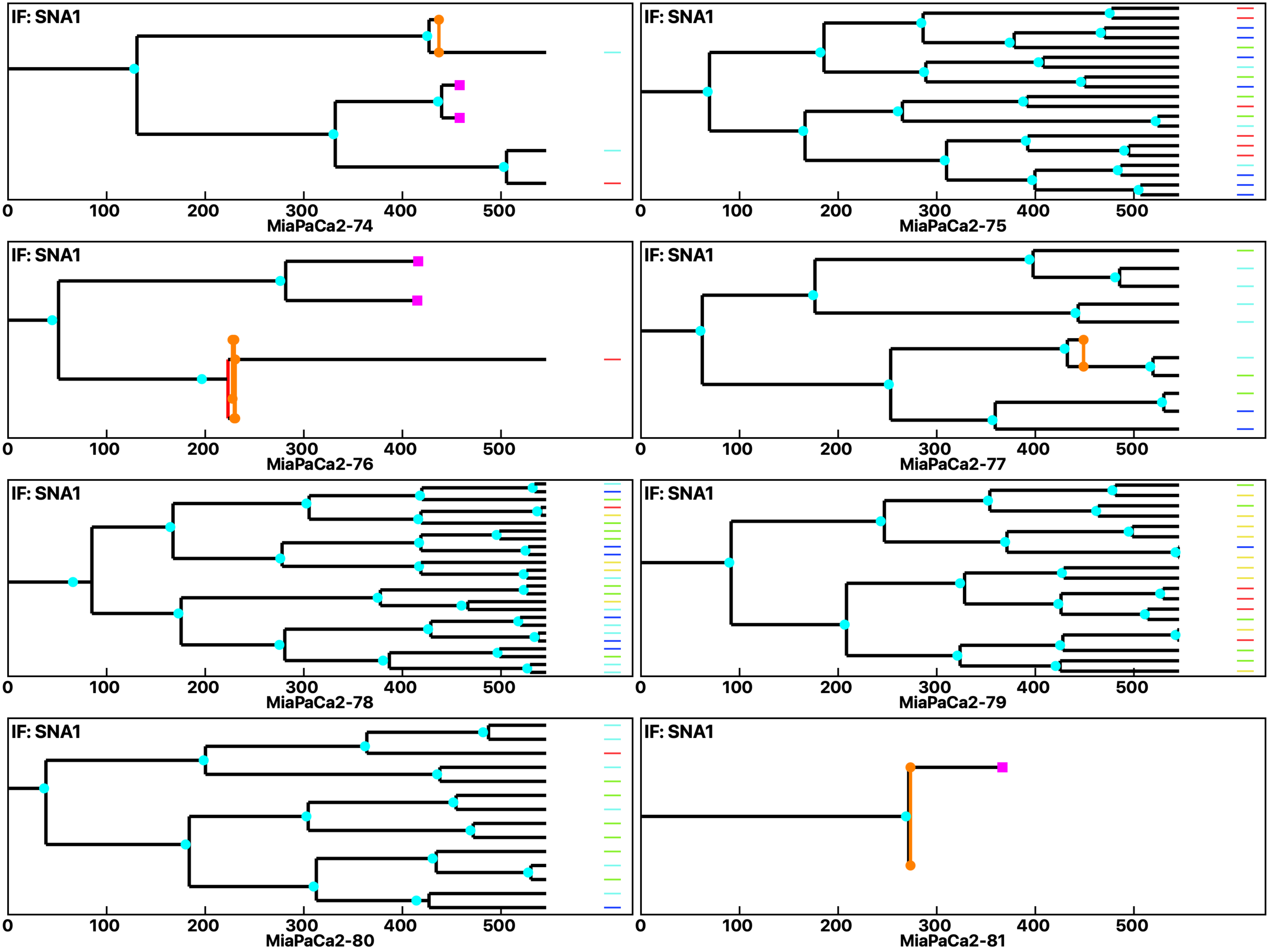

Analysis: MiaPaCa2, Treat.: MiaPaCa2, Cell: MiaPaCa2

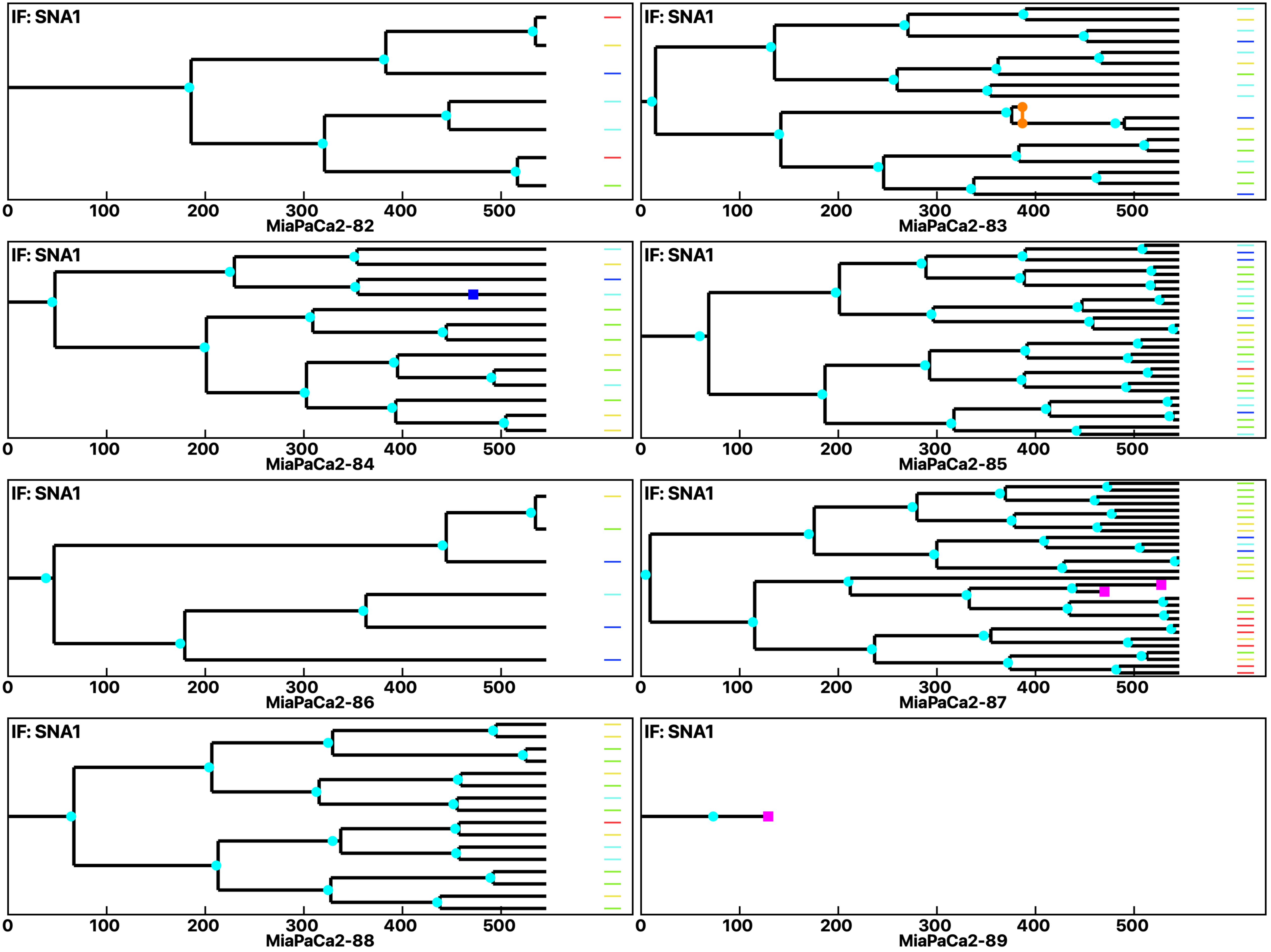

Analysis: MiaPaCa2, Treat.: MiaPaCa2, Cell: MiaPaCa2

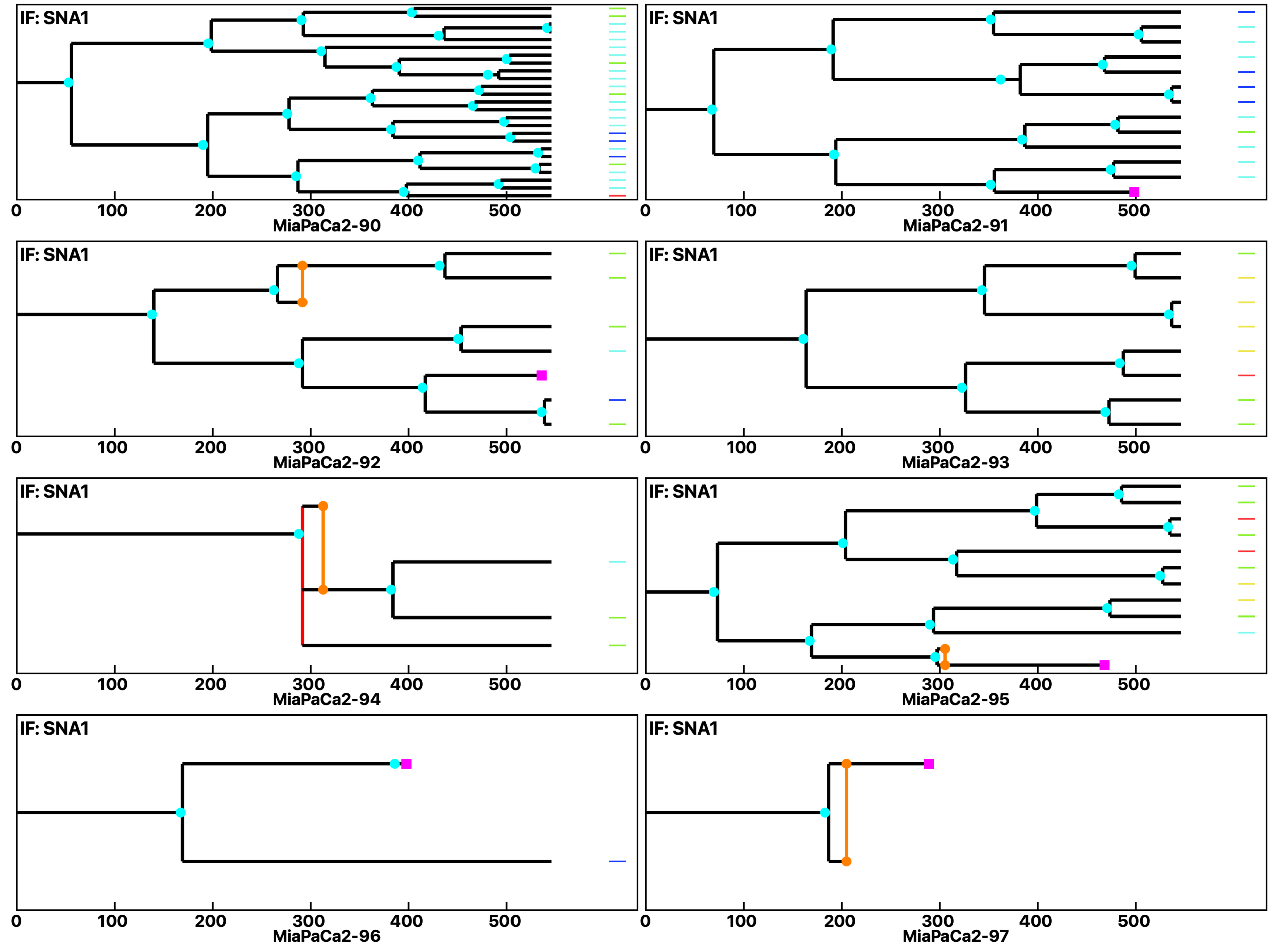

Analysis: MiaPaCa2, Treat.: MiaPaCa2, Cell: MiaPaCa2

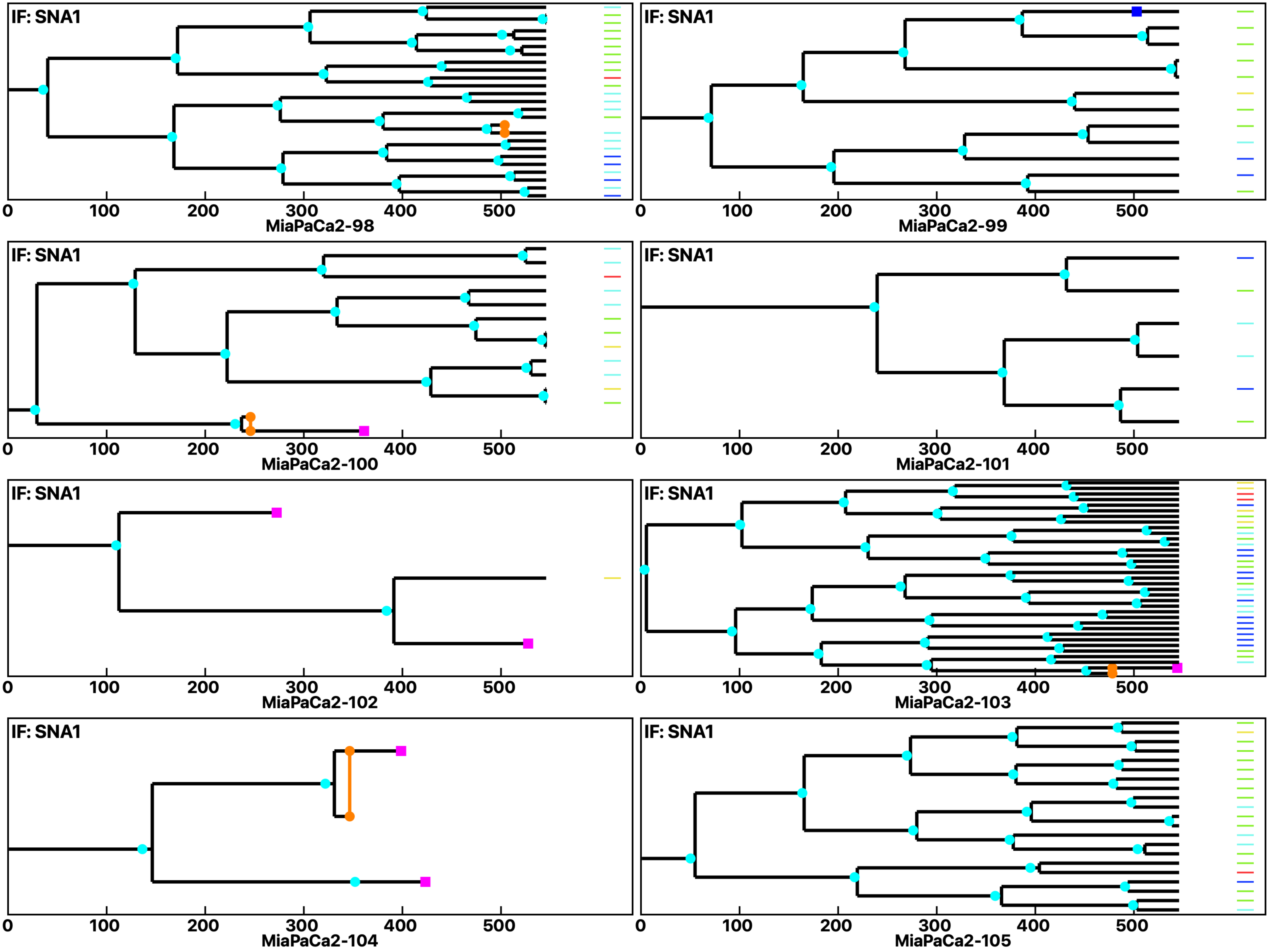

Analysis: MiaPaCa2, Treat.: MiaPaCa2, Cell: MiaPaCa2

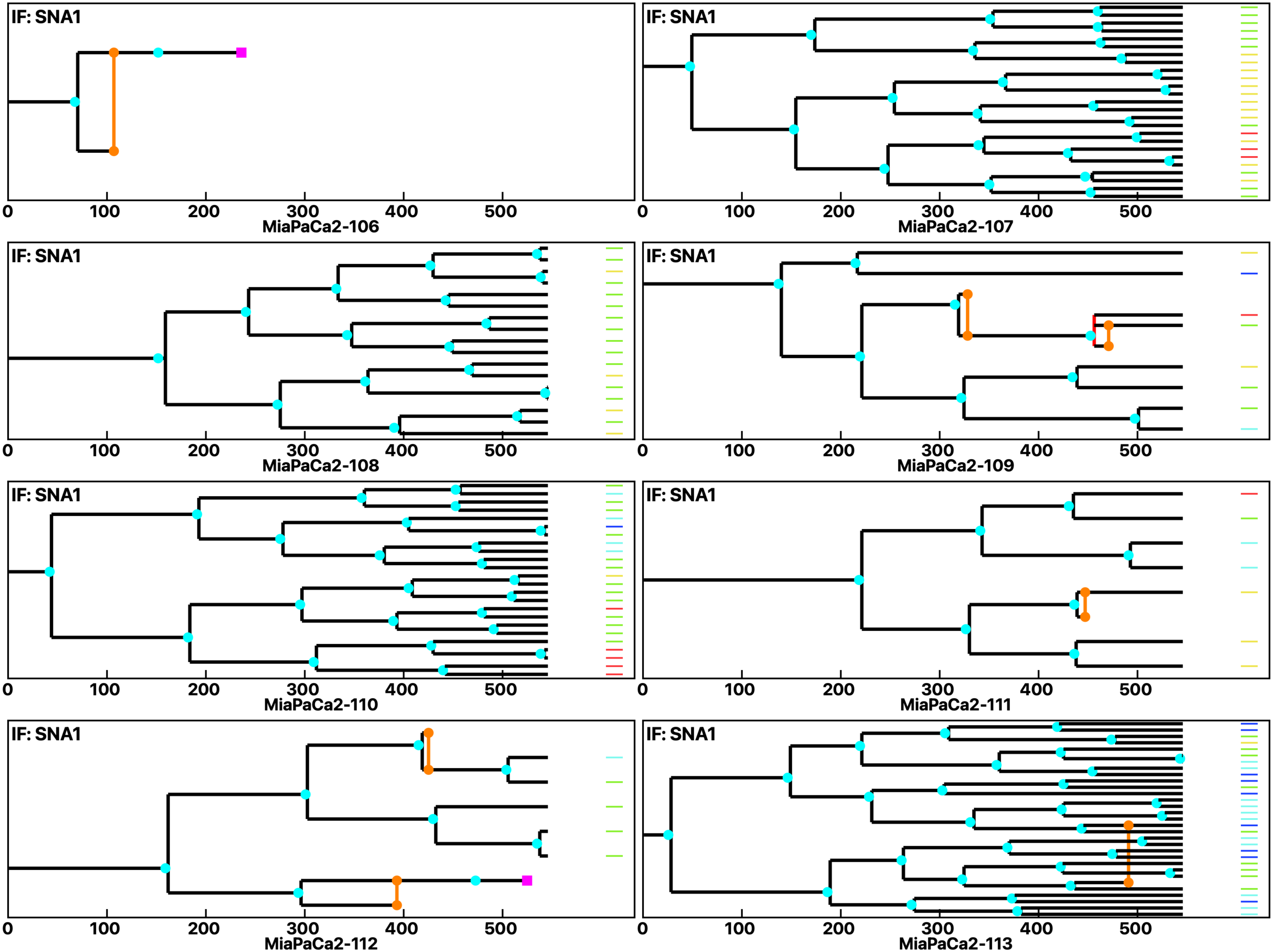

Analysis: MiaPaCa2, Treat.: MiaPaCa2, Cell: MiaPaCa2

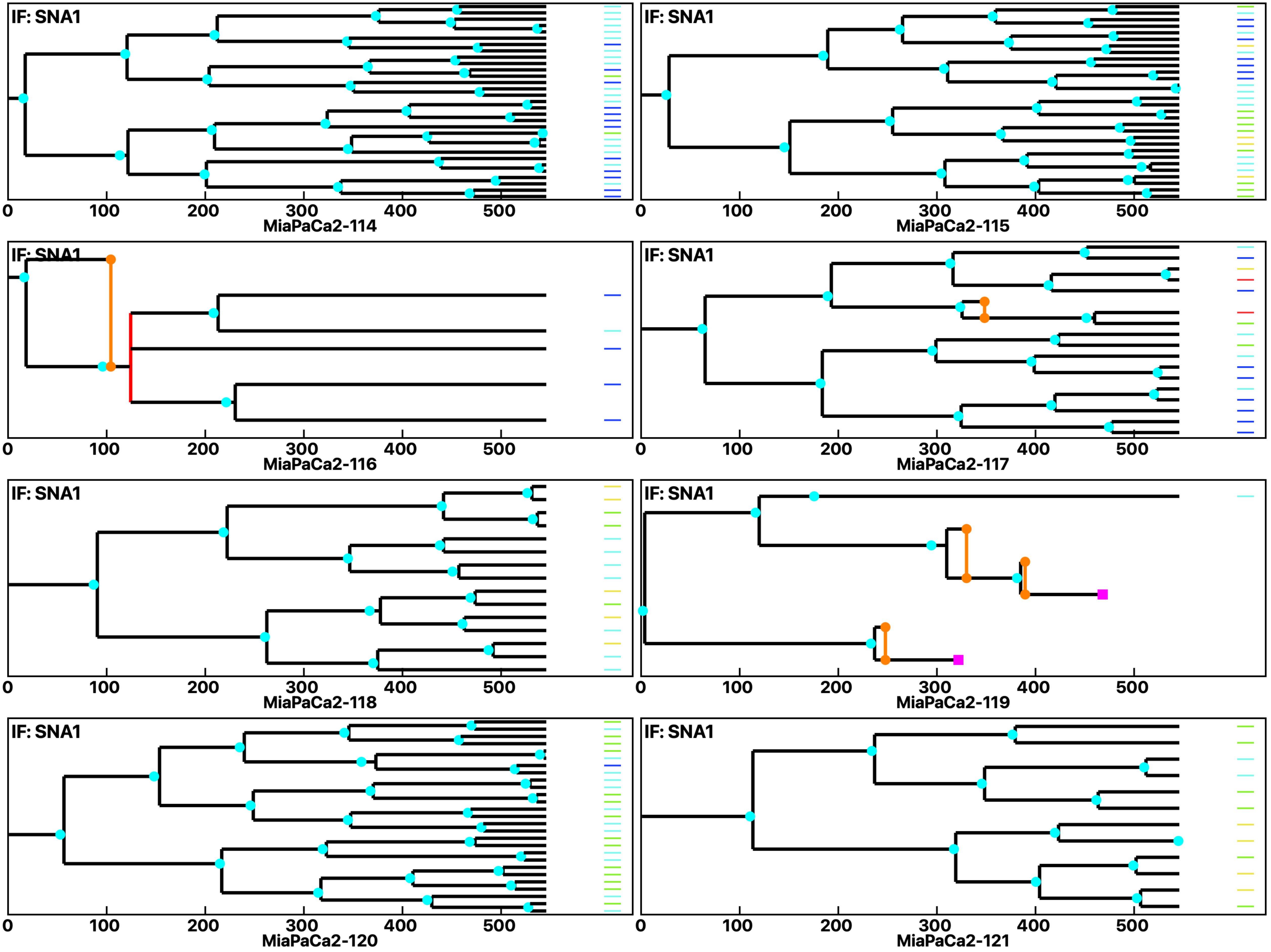

Analysis: MiaPaCa2, Treat.: MiaPaCa2, Cell: MiaPaCa2

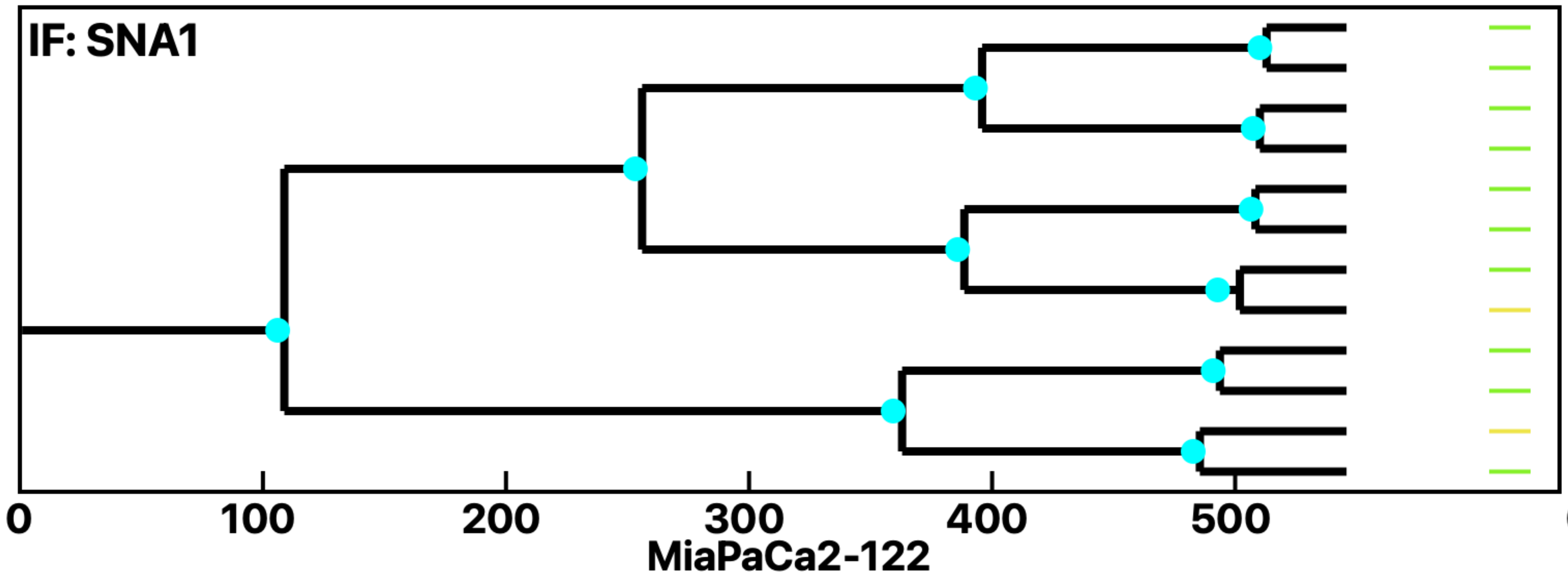


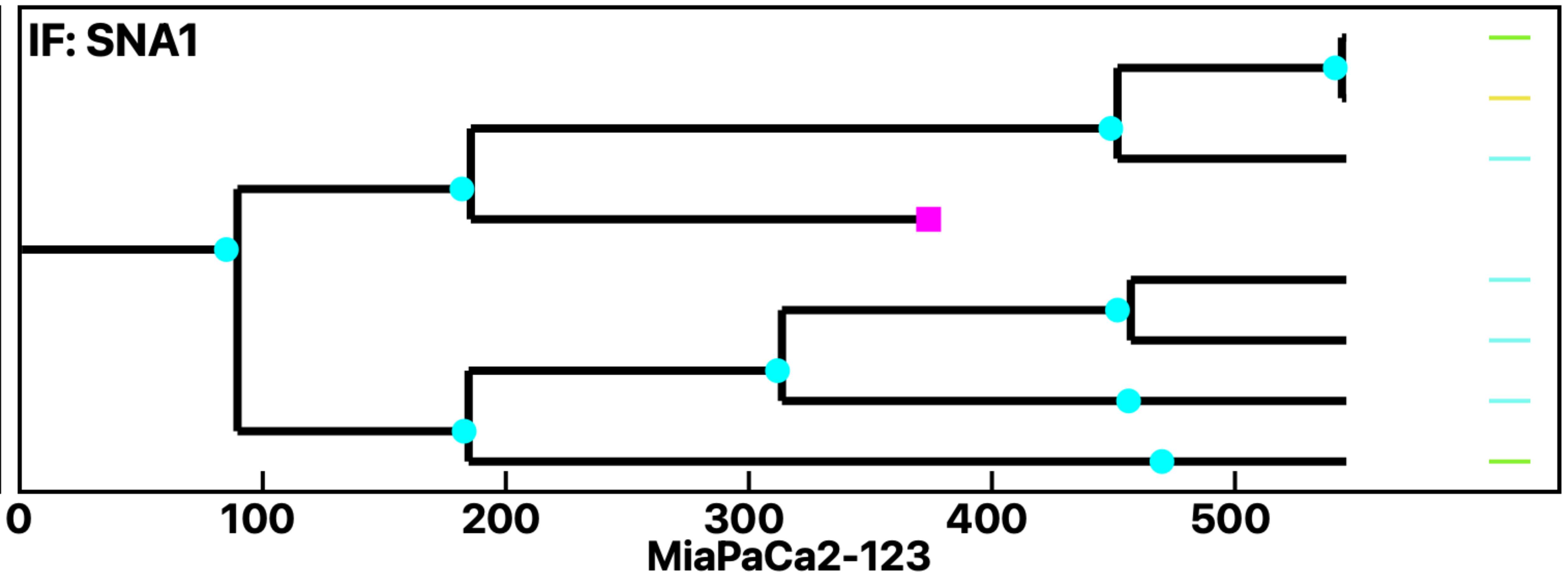


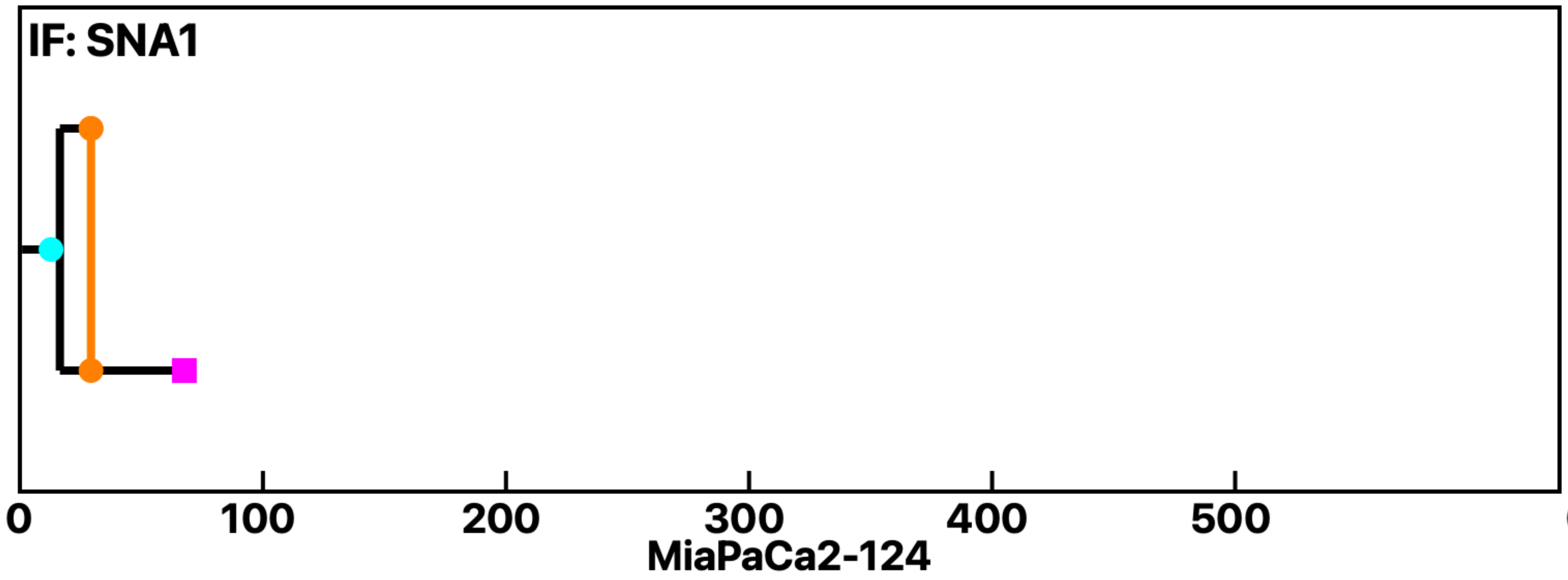


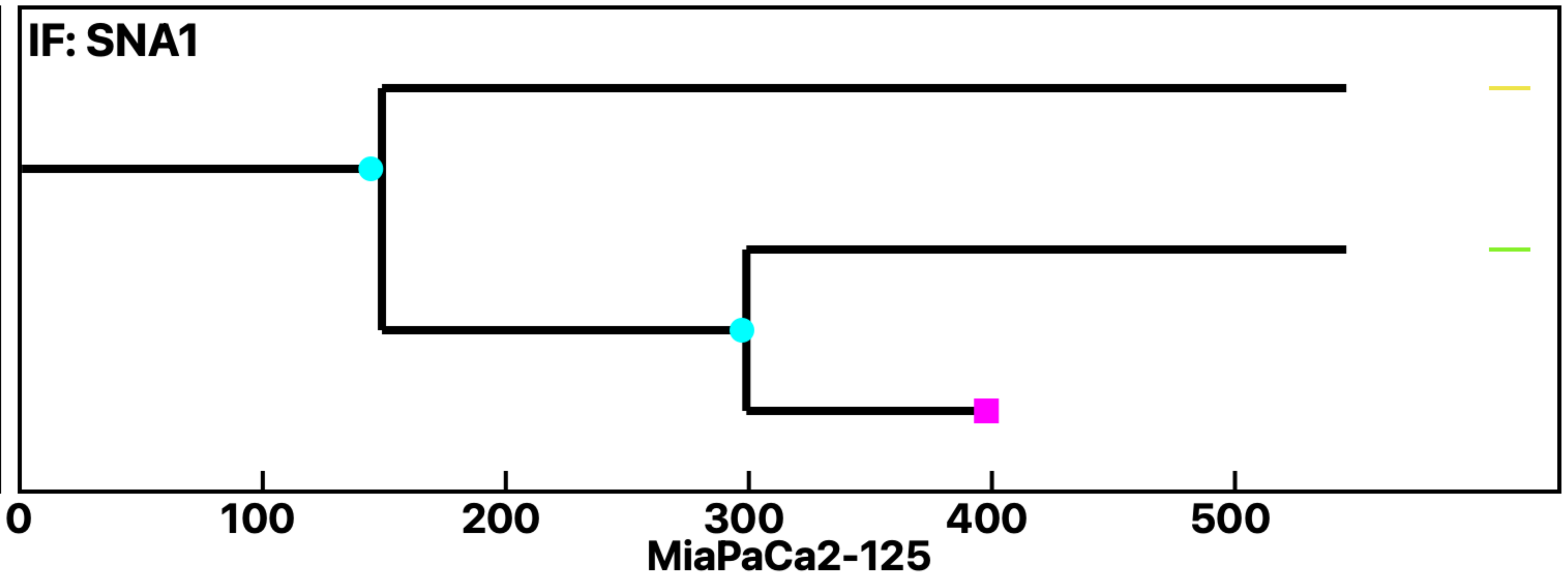


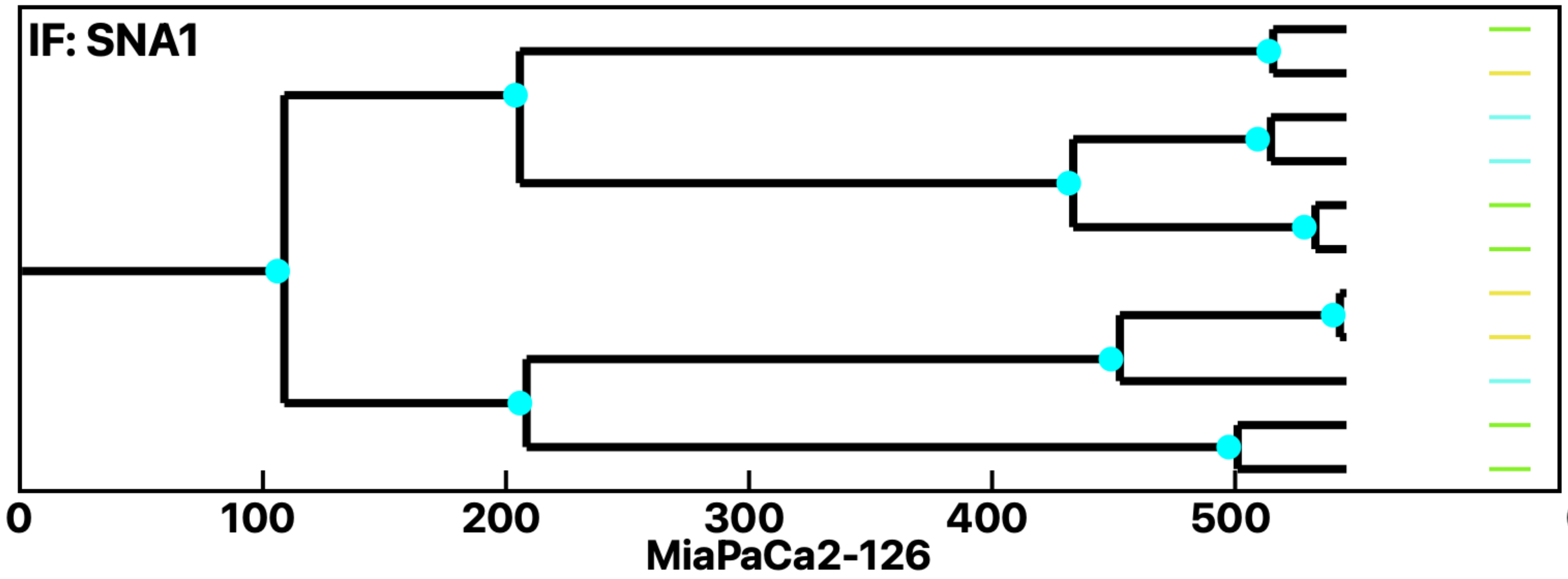


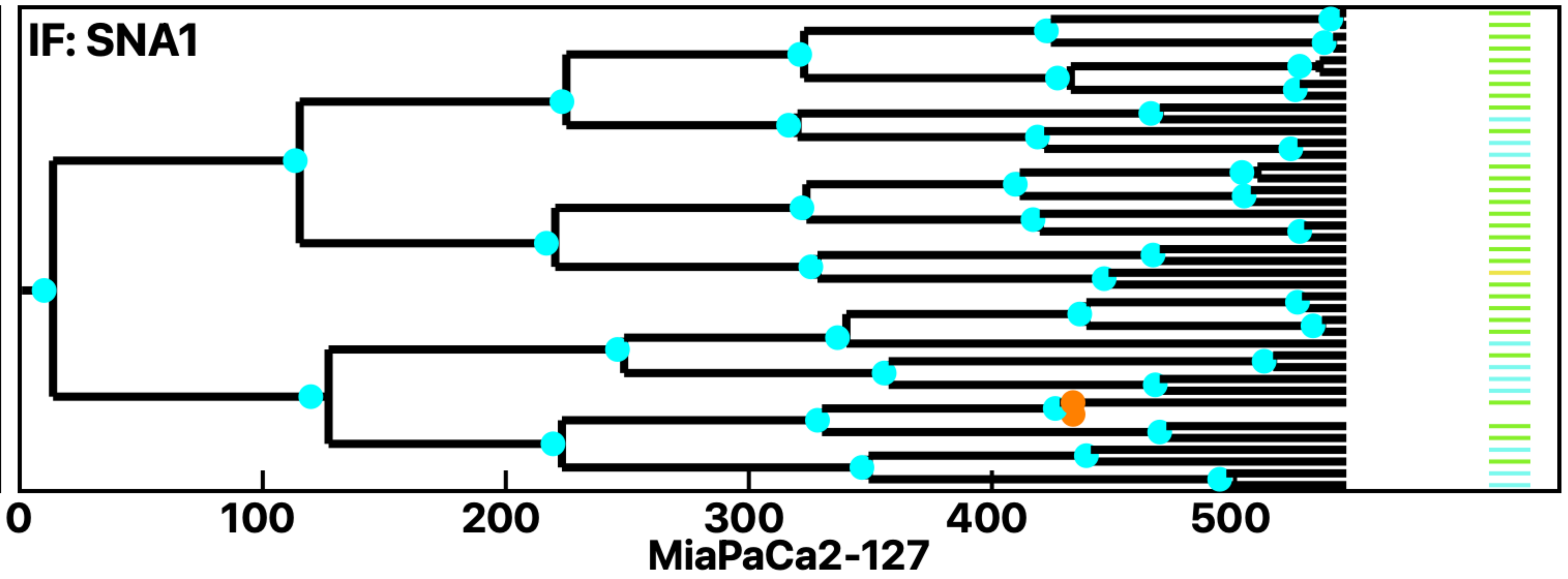


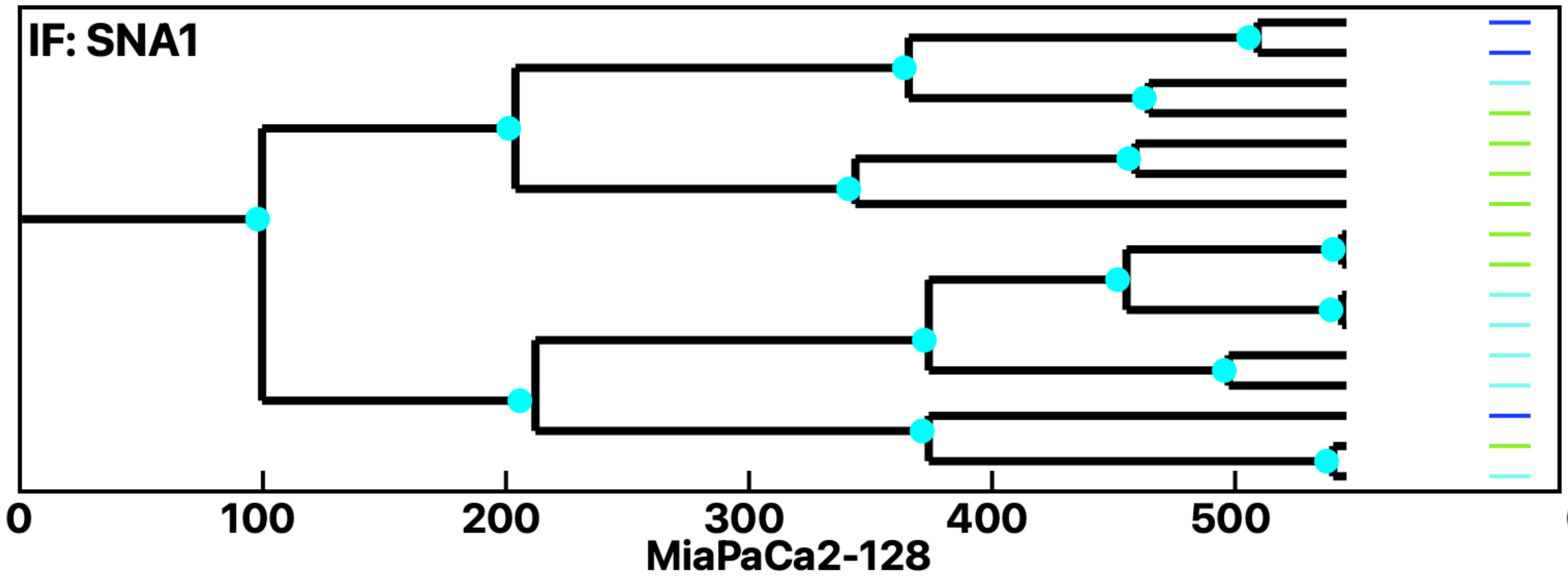


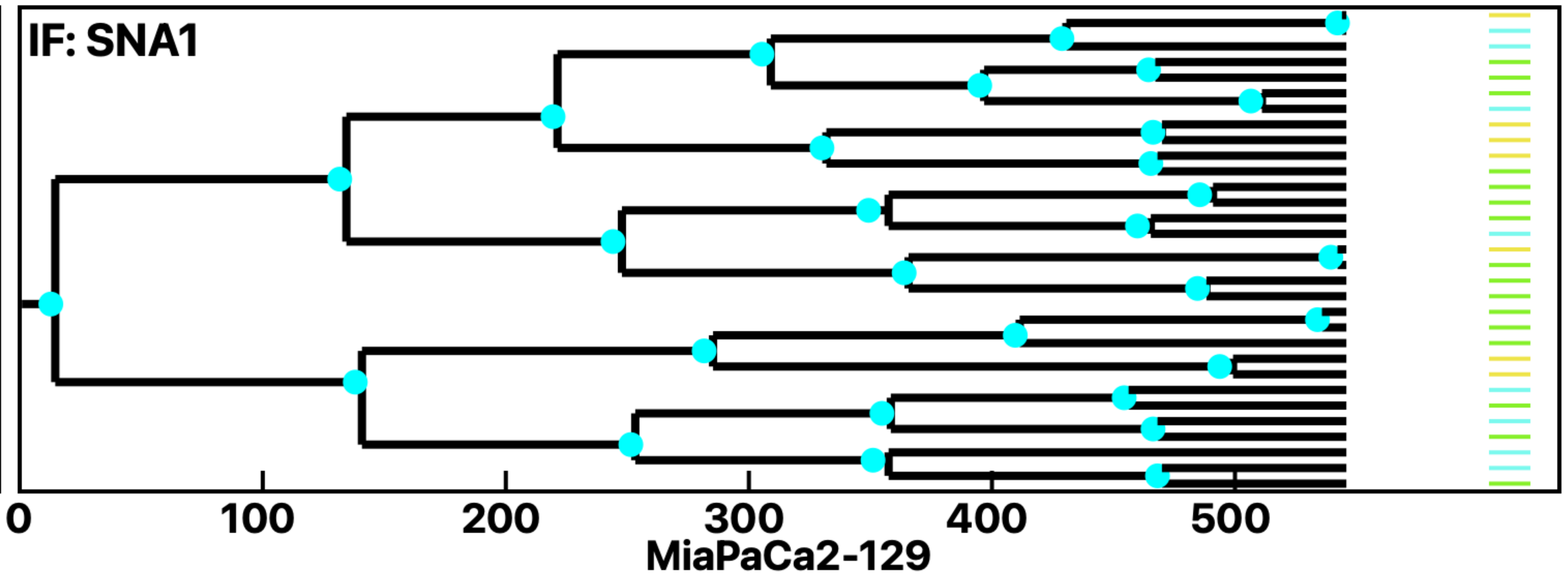

Analysis: MiaPaCa2, Treat.: MiaPaCa2, Cell: MiaPaCa2

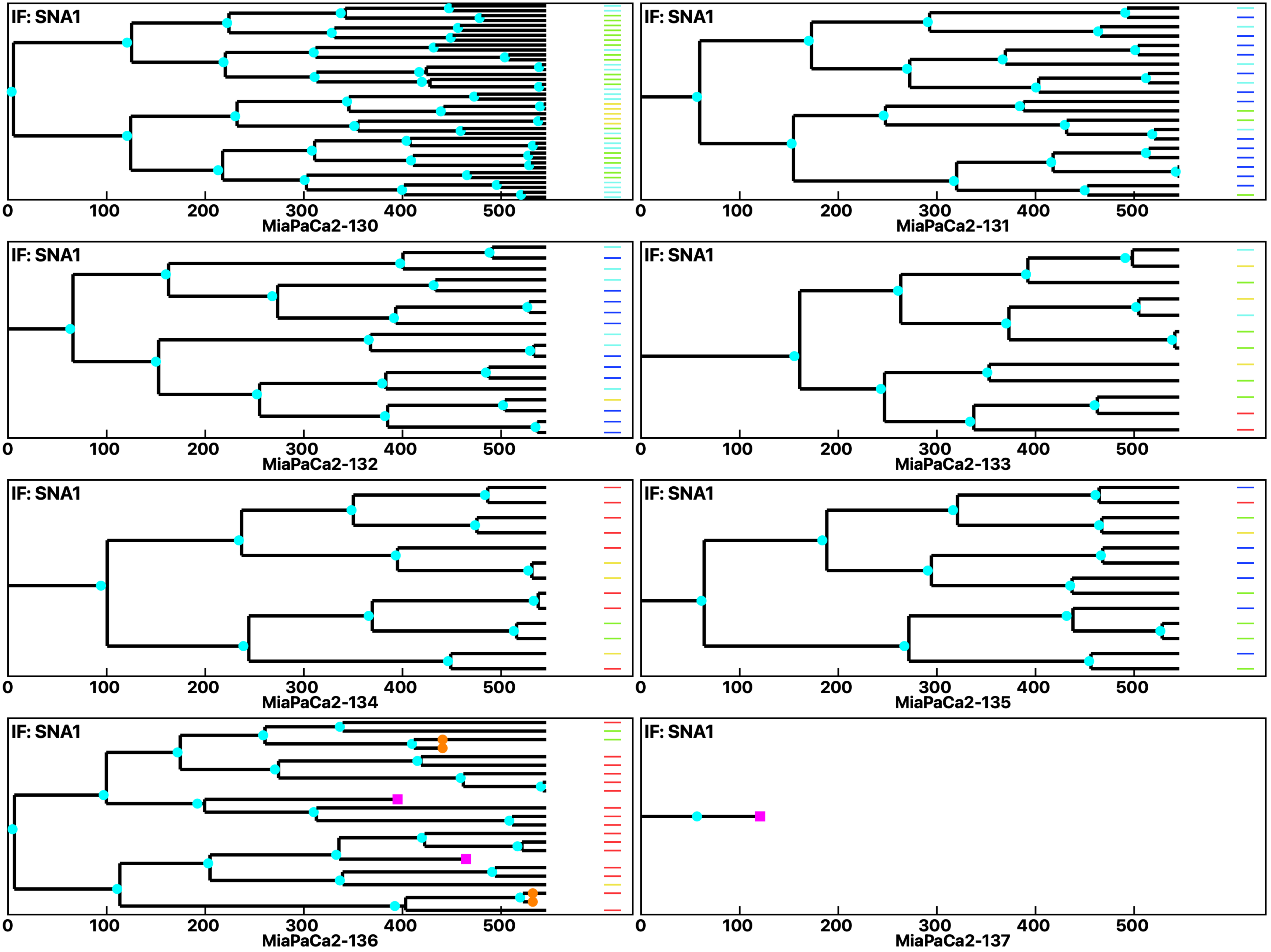

Analysis: MiaPaCa2, Treat.: MiaPaCa2, Cell: MiaPaCa2

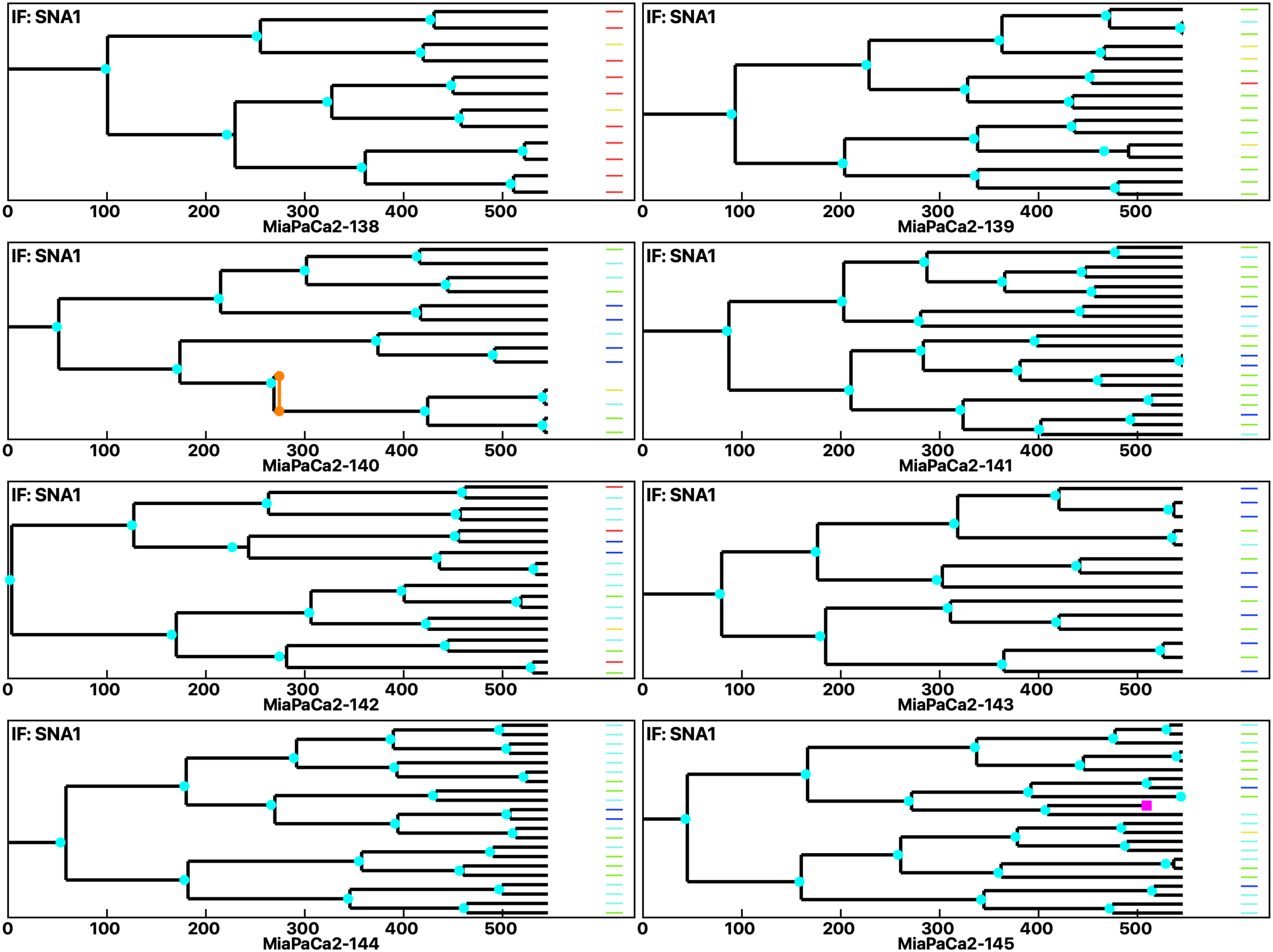

Analysis: MiaPaCa2, Treat.: MiaPaCa2, Cell: MiaPaCa2

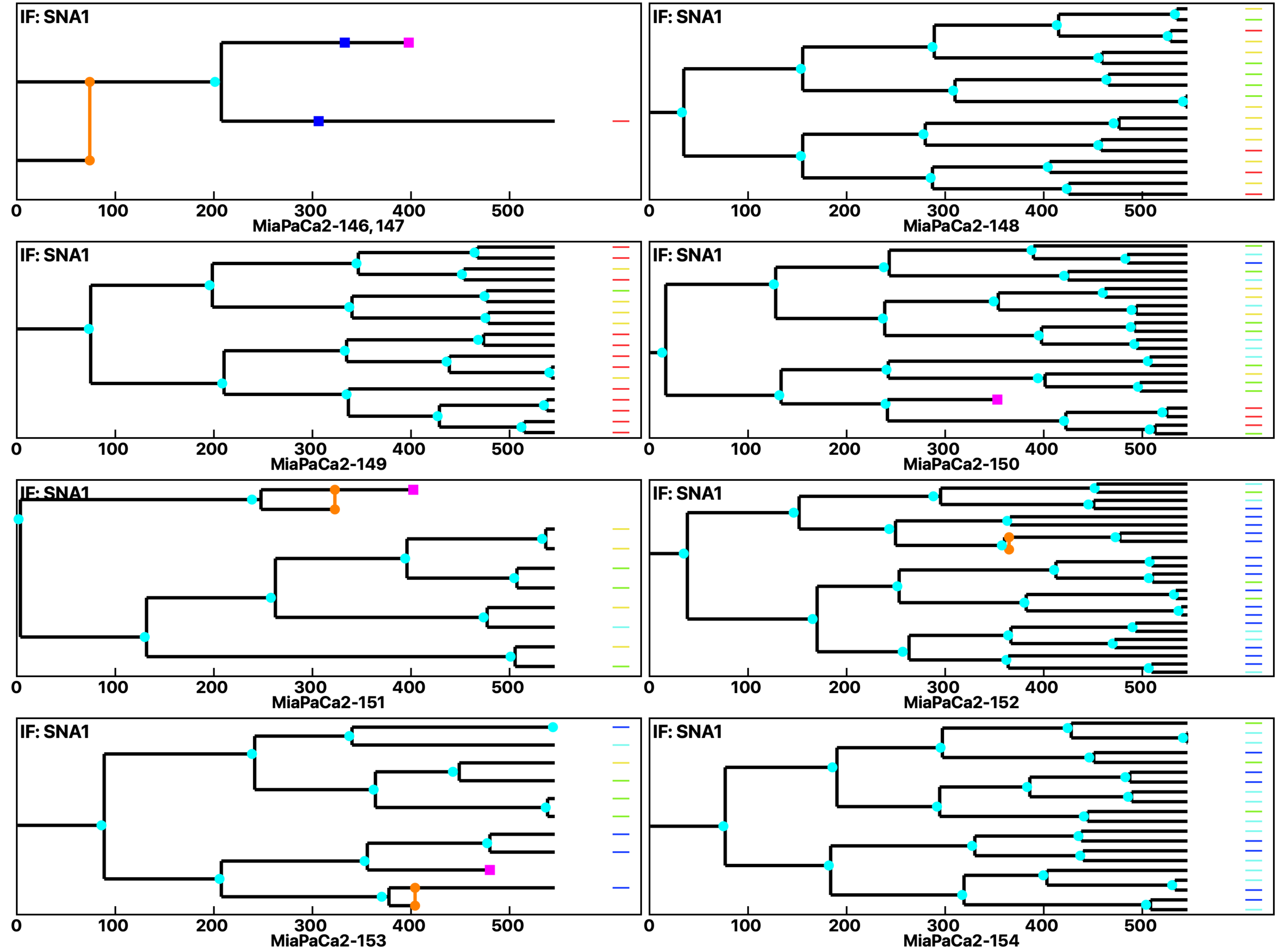

Analysis: MiaPaCa2, Treat.: MiaPaCa2, Cell: MiaPaCa2

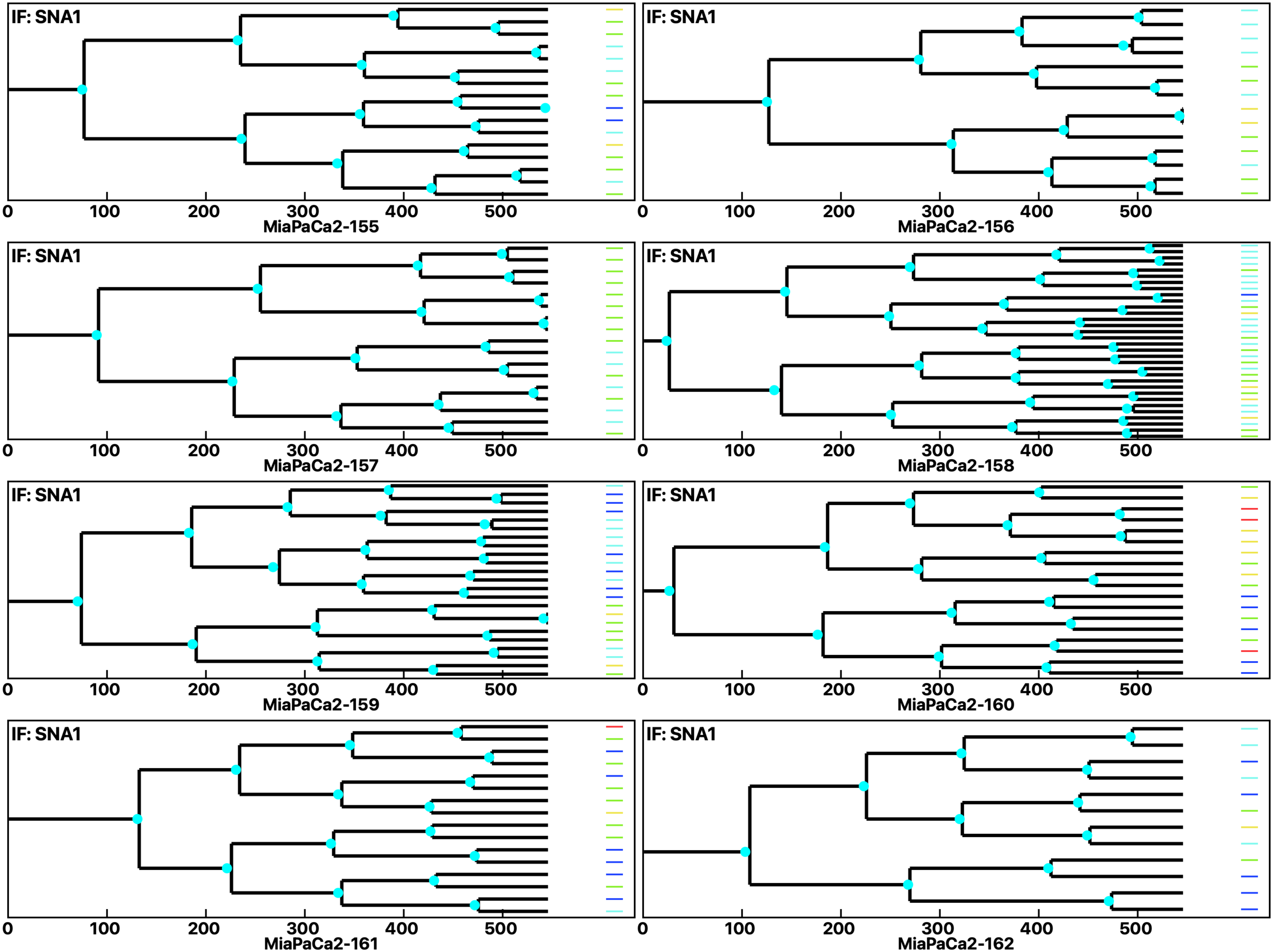

Analysis: MiaPaCa2, Treat.: MiaPaCa2, Cell: MiaPaCa2

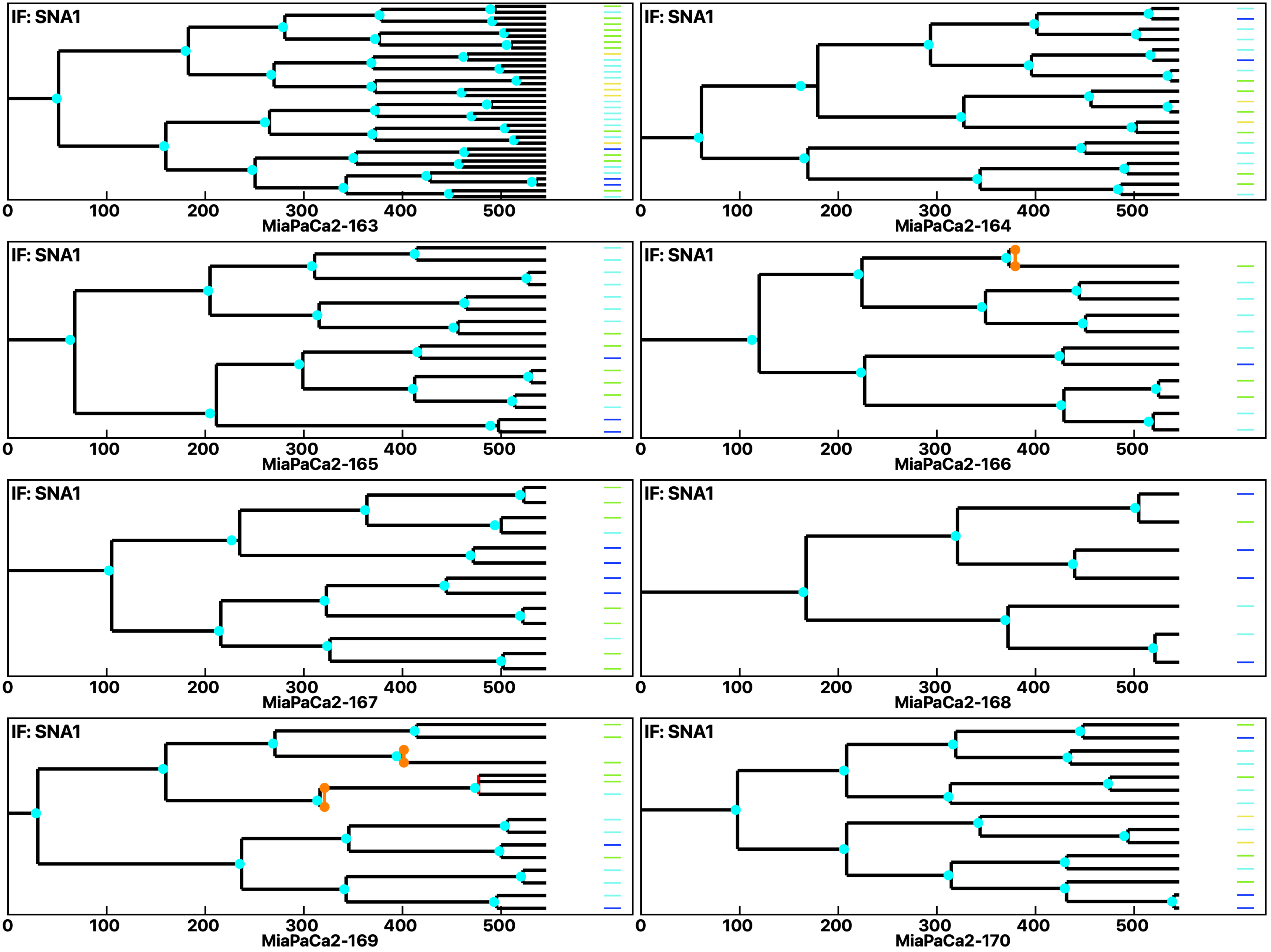

Analysis: MiaPaCa2, Treat.: MiaPaCa2, Cell: MiaPaCa2

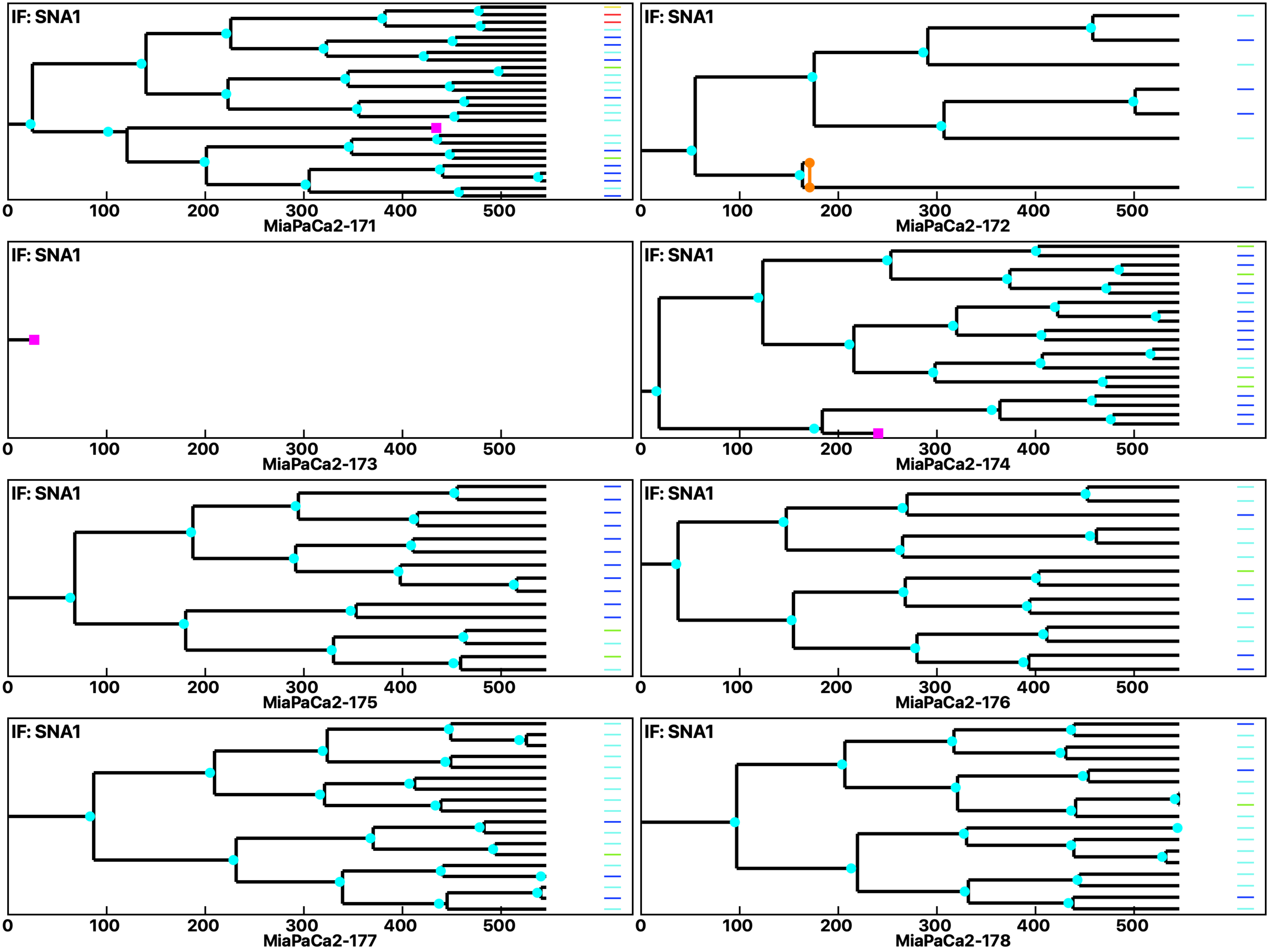

Analysis: MiaPaCa2, Treat.: MiaPaCa2, Cell: MiaPaCa2

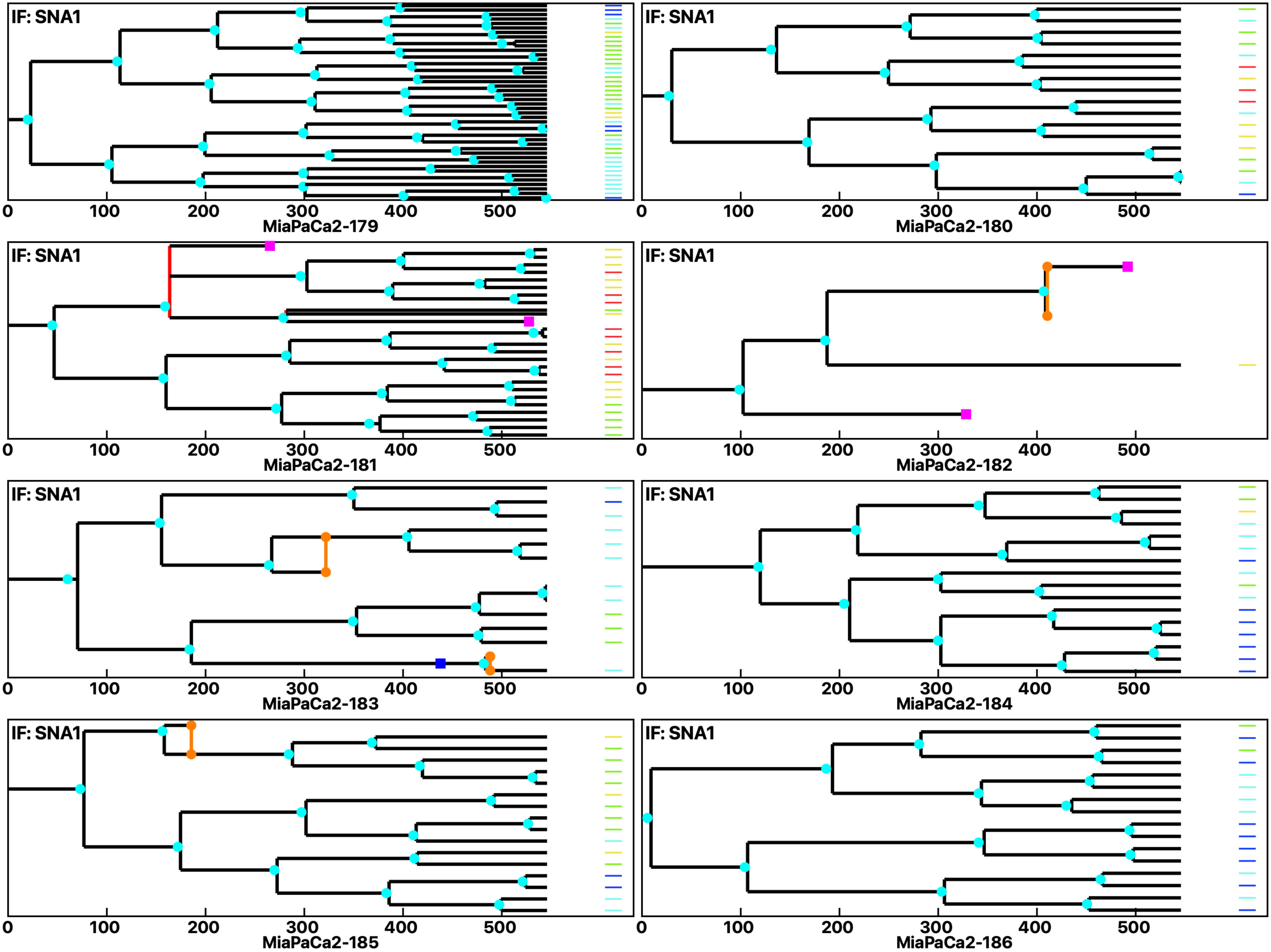

Analysis: MiaPaCa2, Treat.: MiaPaCa2, Cell: MiaPaCa2

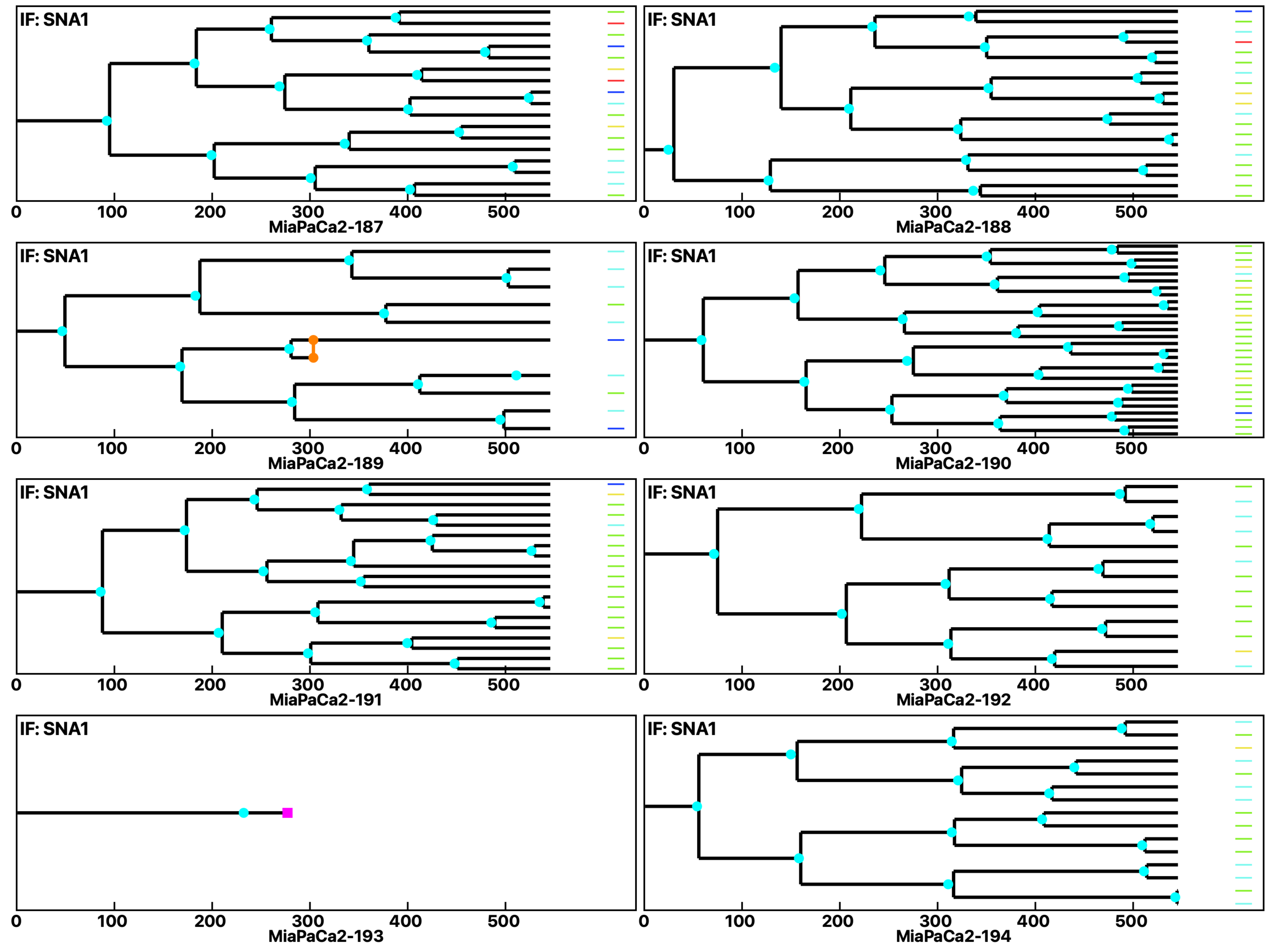

Analysis: MiaPaCa2, Treat.: MiaPaCa2, Cell: MiaPaCa2

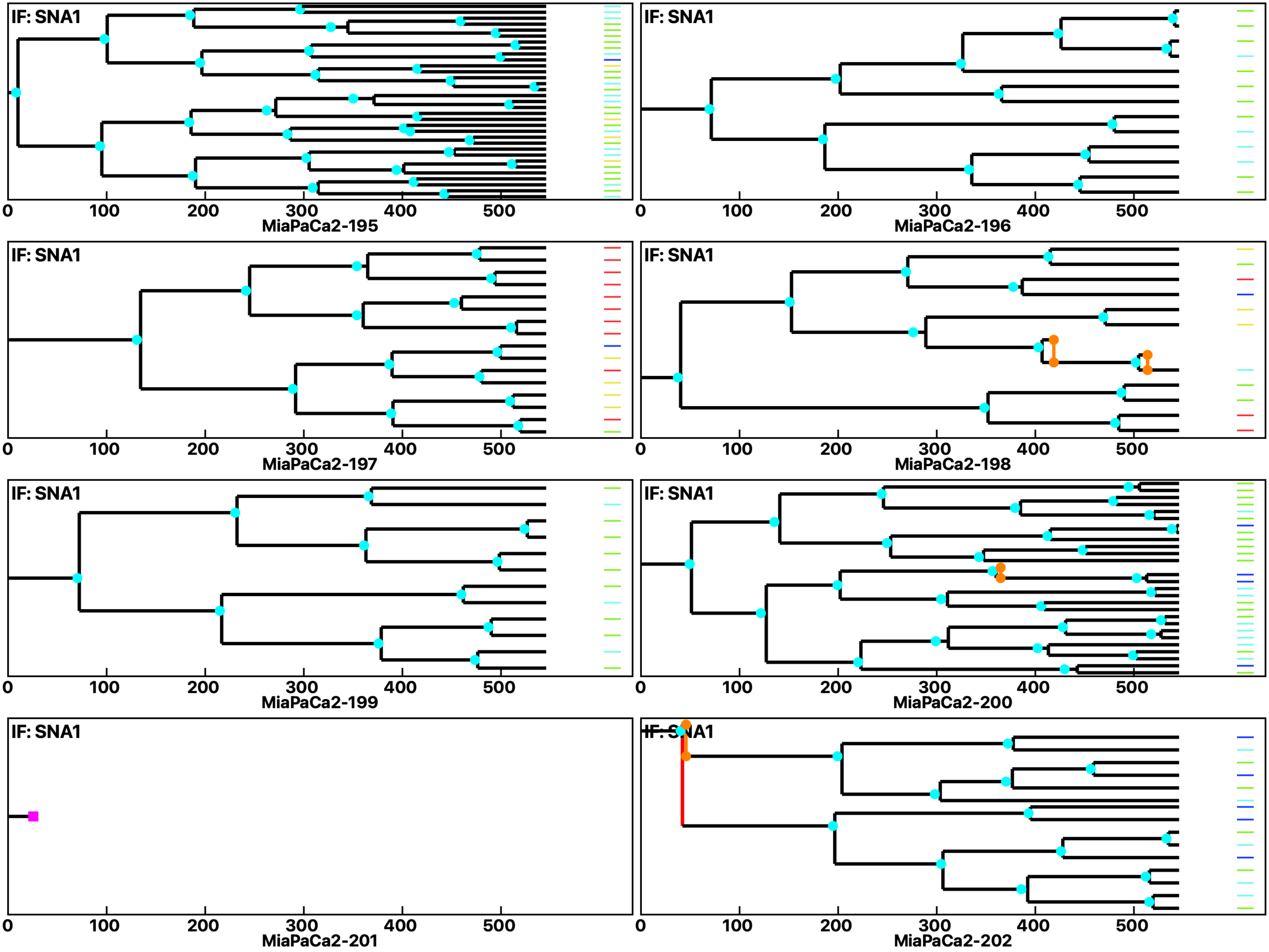

Analysis: MiaPaCa2, Treat.: MiaPaCa2, Cell: MiaPaCa2

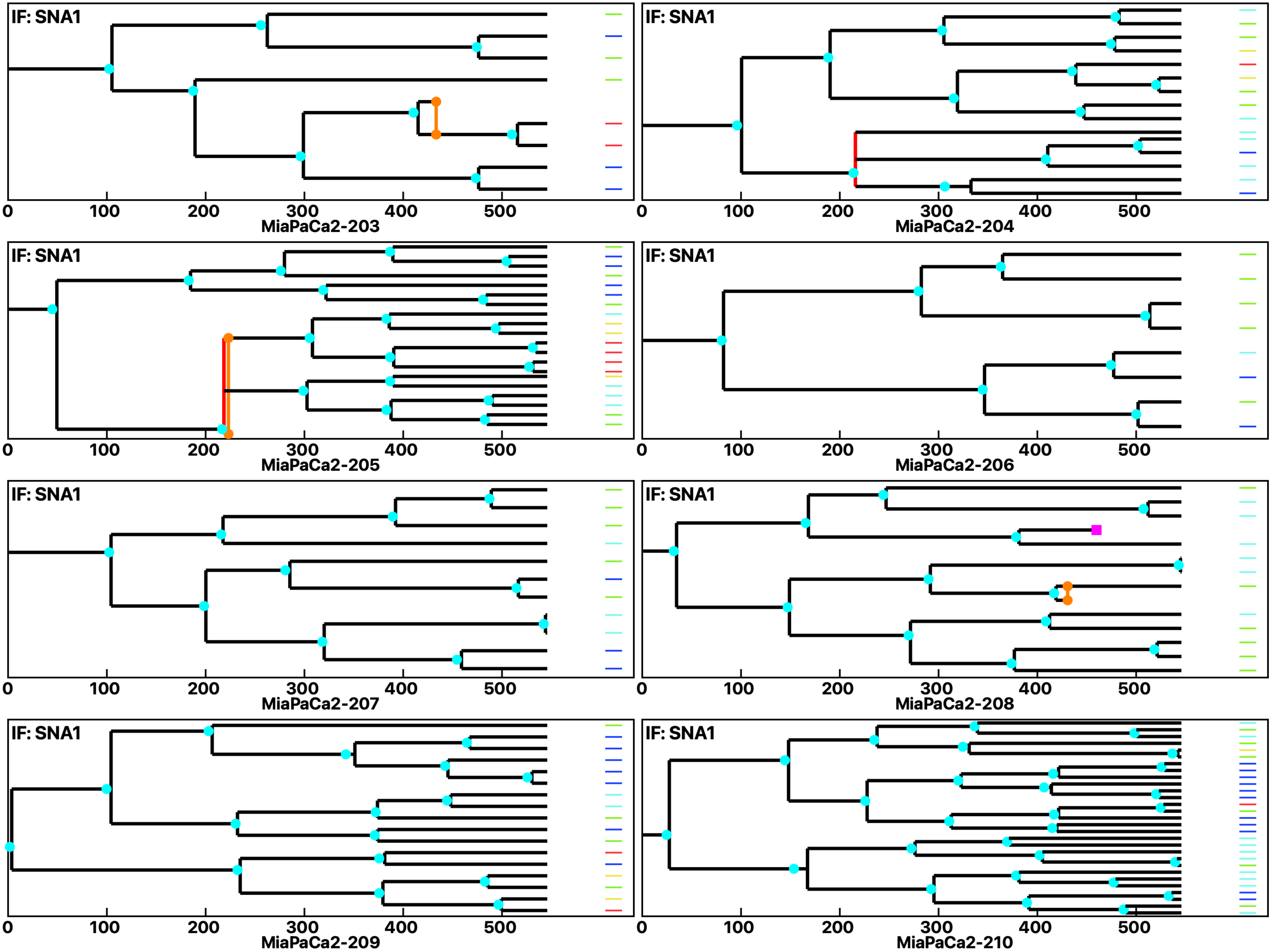

Analysis: MiaPaCa2, Treat.: MiaPaCa2, Cell: MiaPaCa2

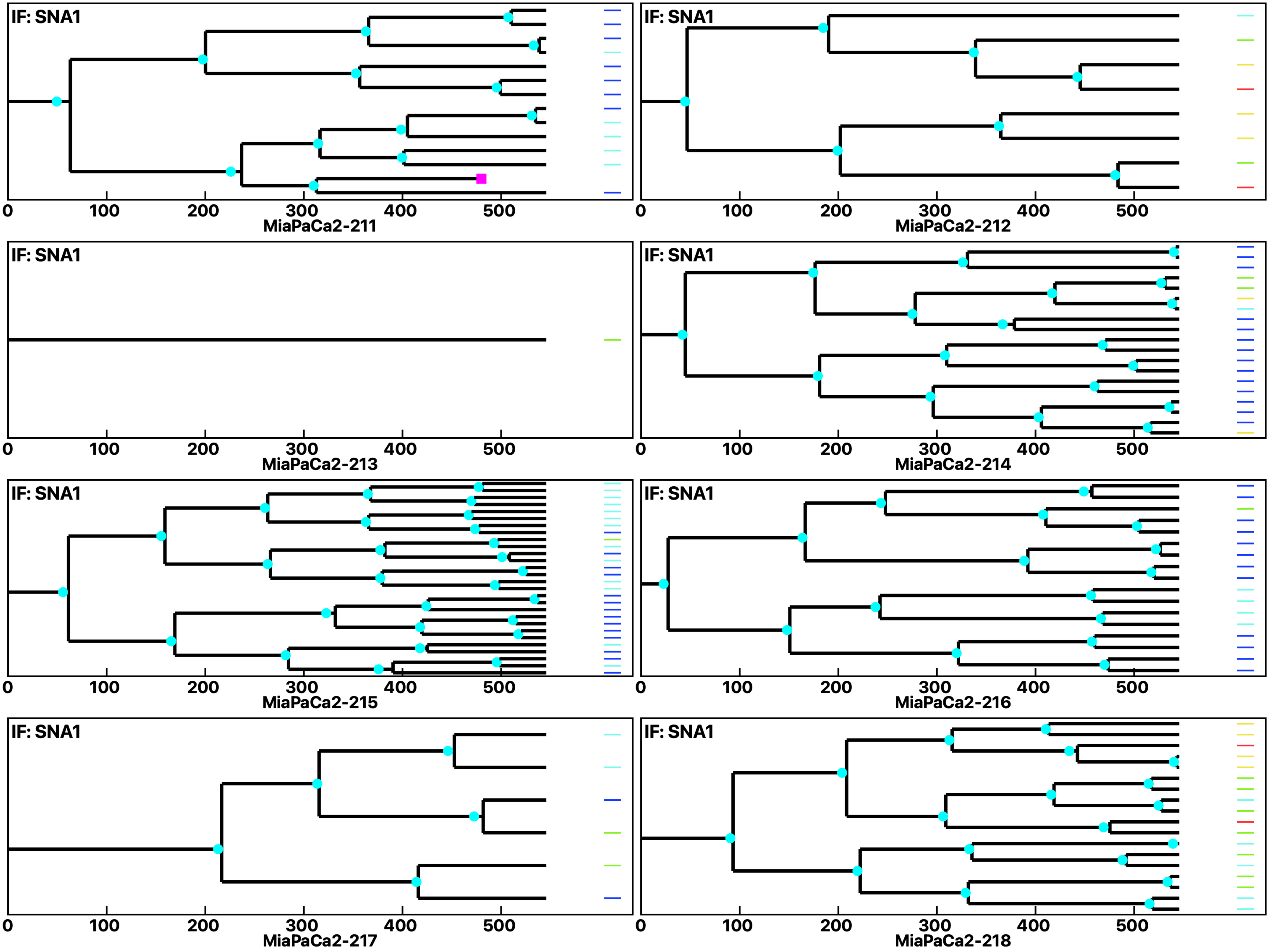

Analysis: MiaPaCa2, Treat.: MiaPaCa2, Cell: MiaPaCa2

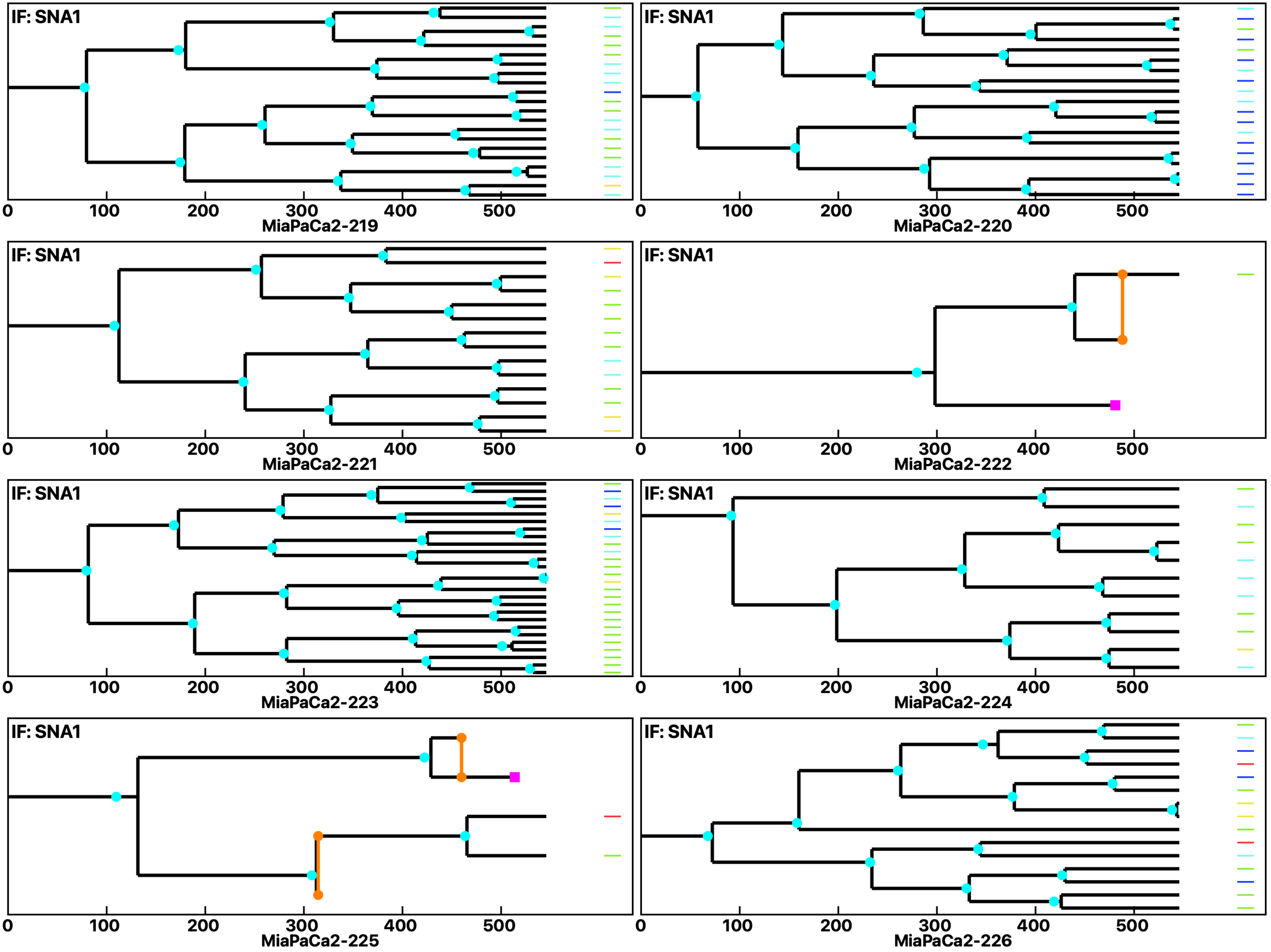

Analysis: MiaPaCa2, Treat.: MiaPaCa2, Cell: MiaPaCa2

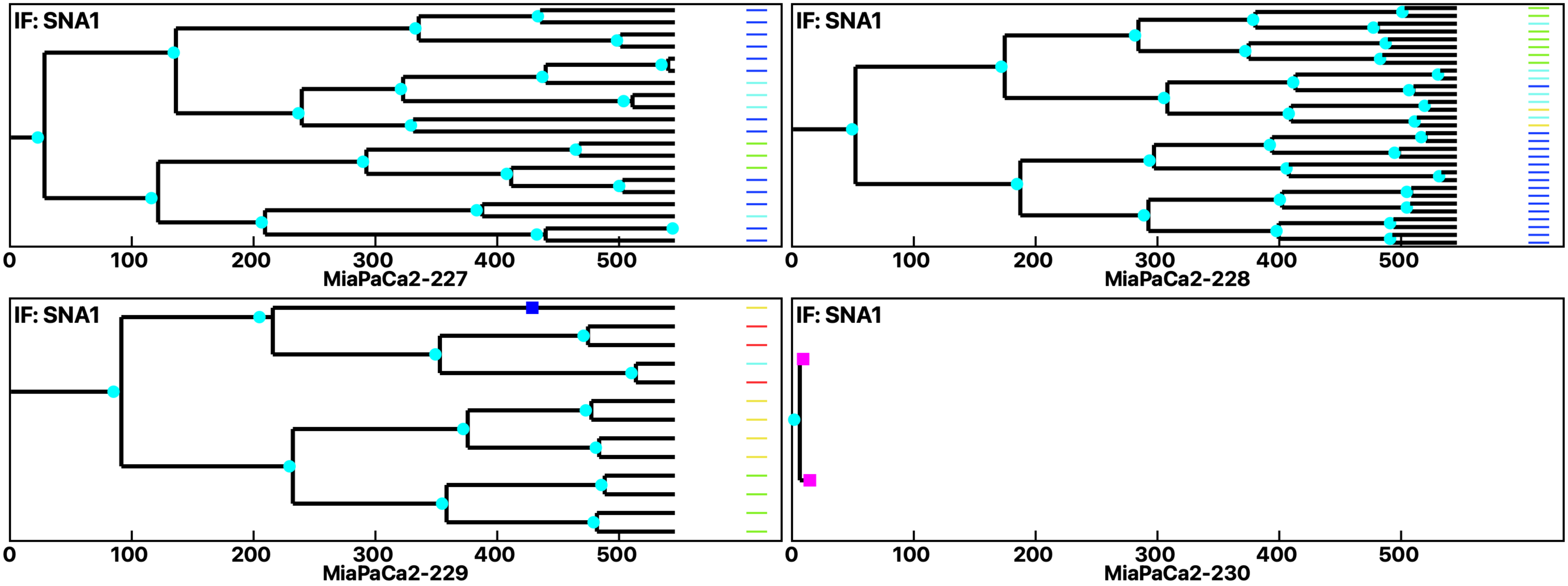
