## Supplementary Data 4 for "Cell Fate Simulation Reveals Cancer Cell Features in the Tumor Microenvironment"

Analysis: Simulation, Treat.: Pancreatic 2-6Sia 1.5

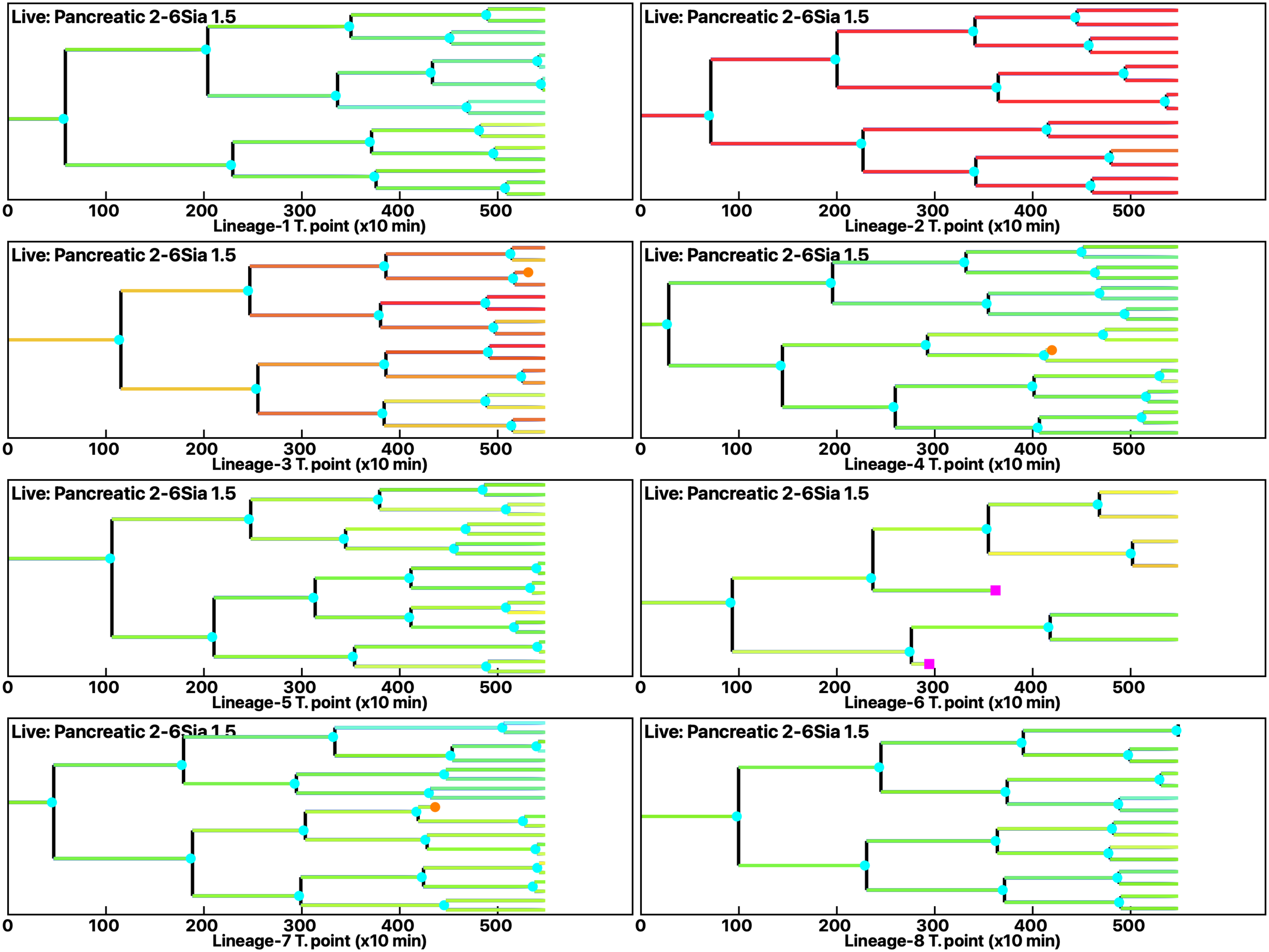

**Analysis: Simulation, Treat.: Pancreatic 2-6Sia 1.5**

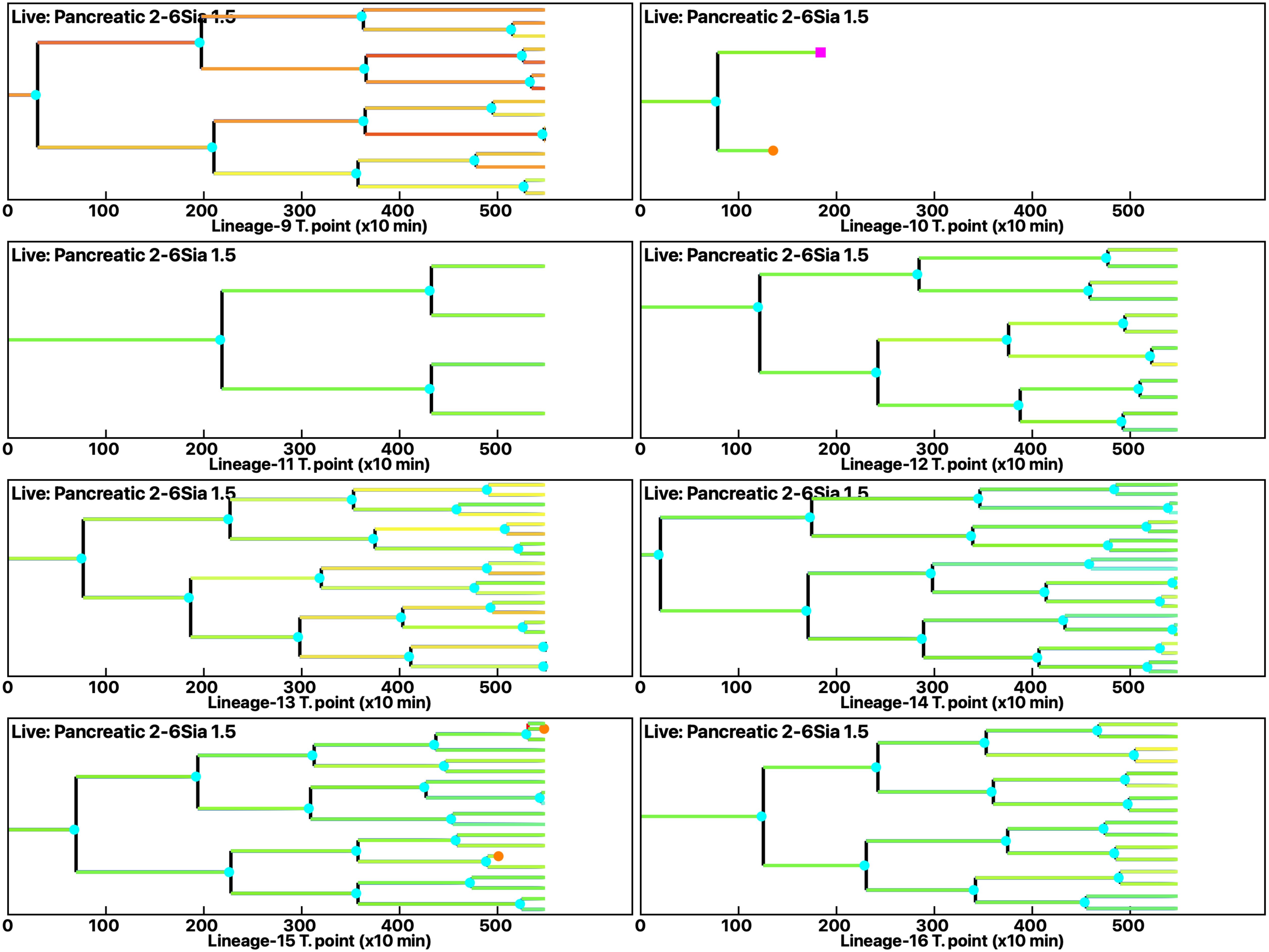

Analysis: Simulation, Treat.: Pancreatic 2-6Sia 1.5

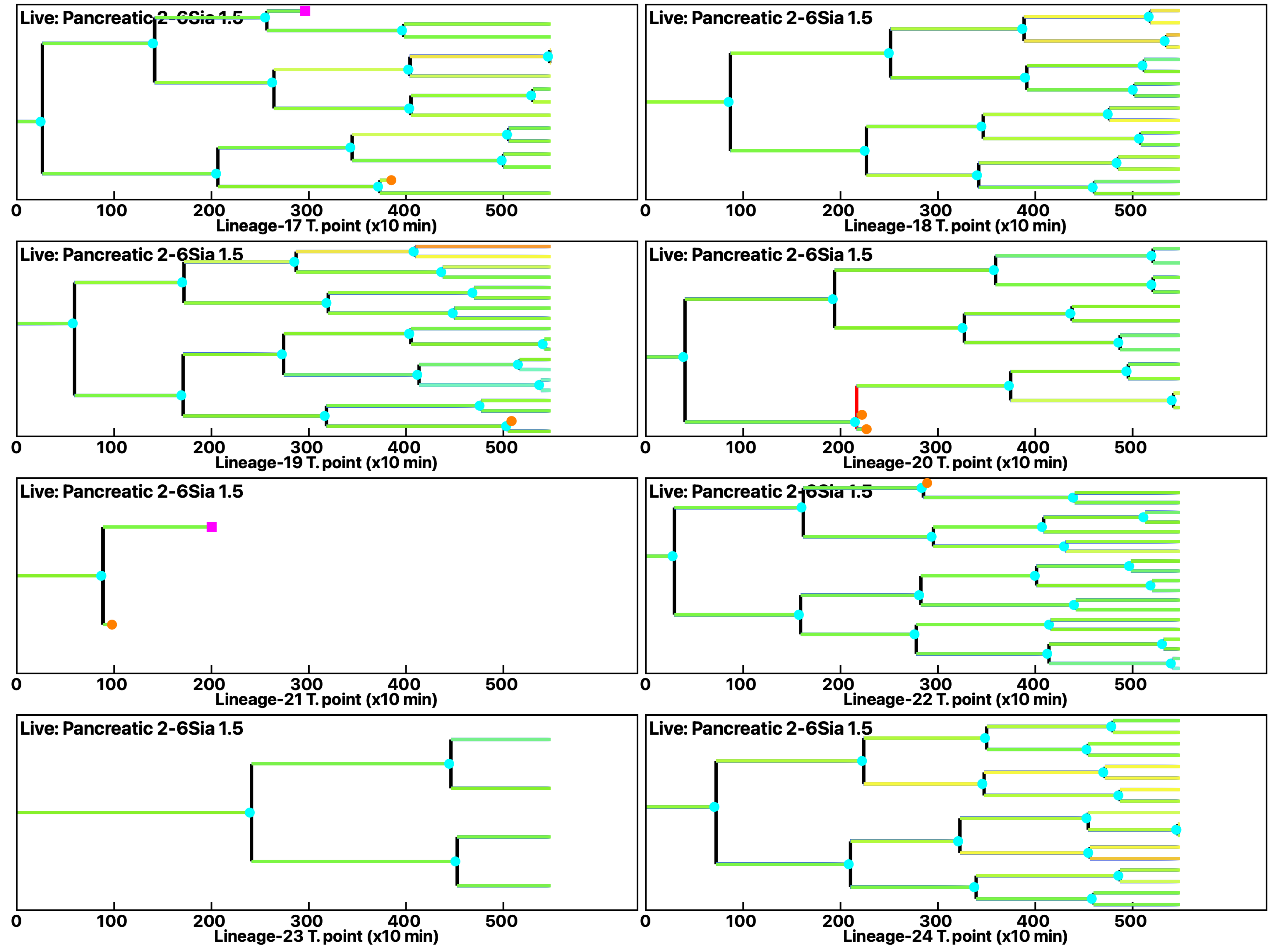

Analysis: Simulation, Treat.: Pancreatic 2-6Sia 1.5

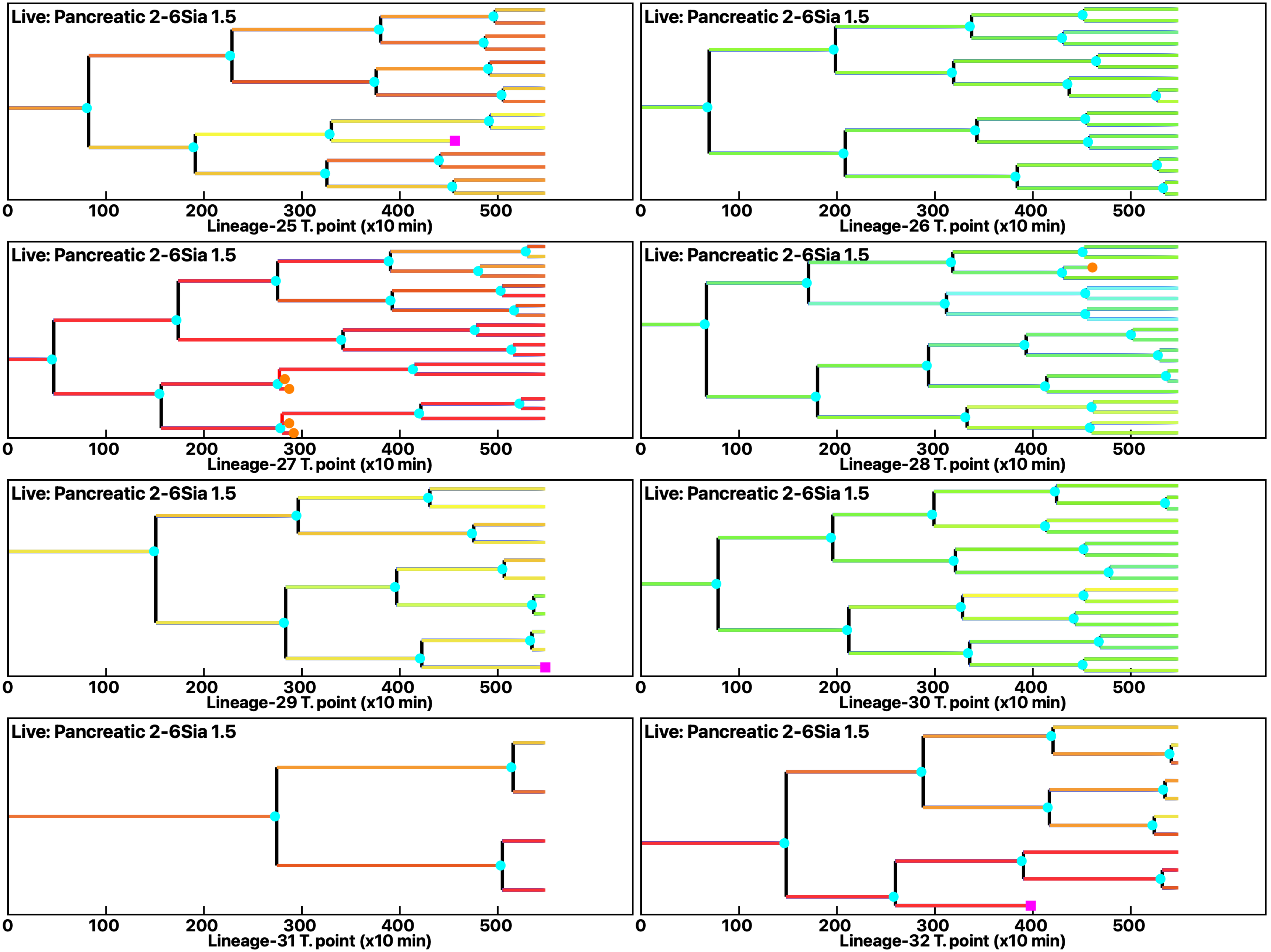

Analysis: Simulation, Treat.: Pancreatic 2-6Sia 1.5

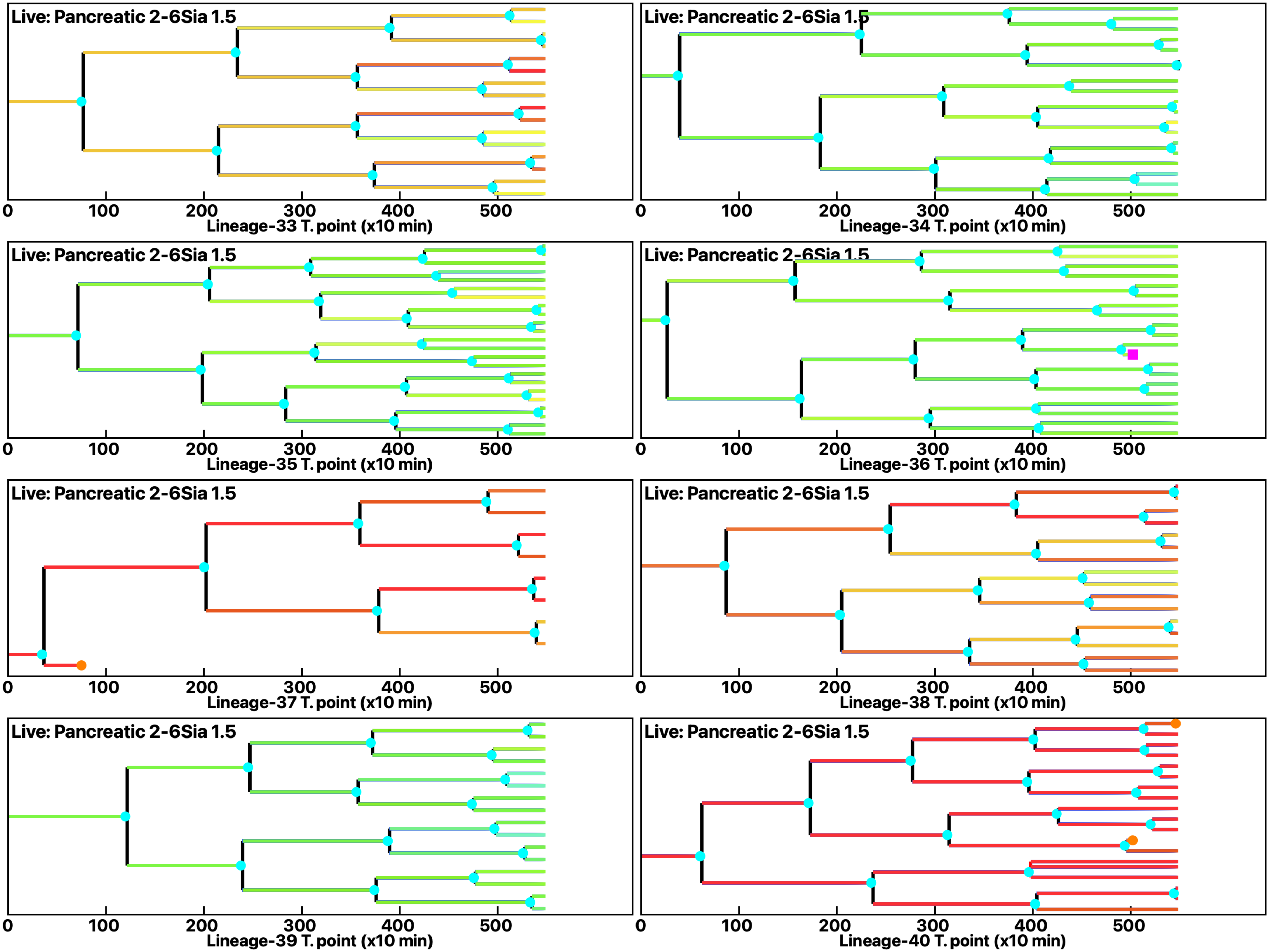

**Analysis: Simulation, Treat.: Pancreatic 2-6Sia 1.5**

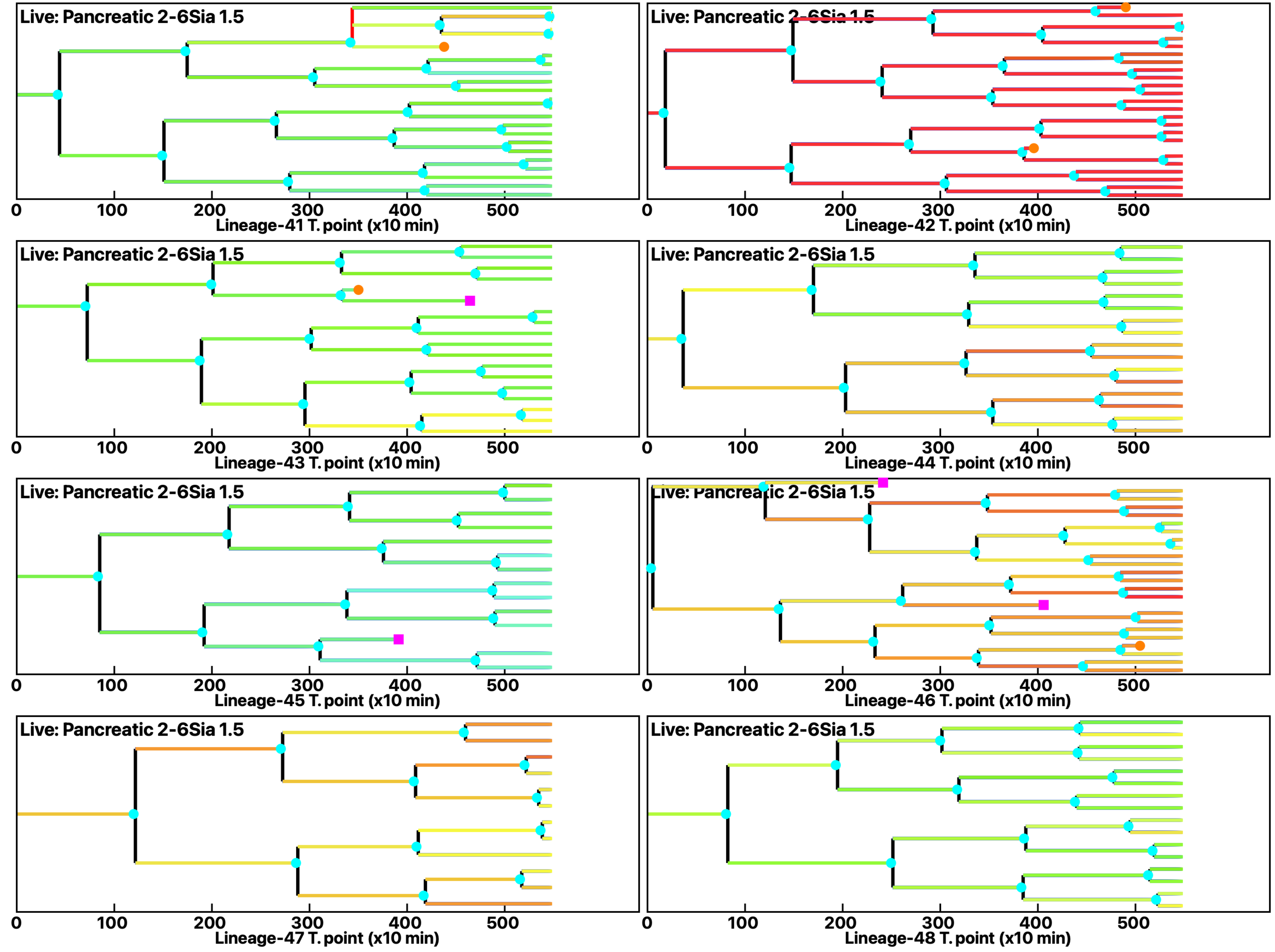

Analysis: Simulation, Treat.: Pancreatic 2-6Sia 1.5

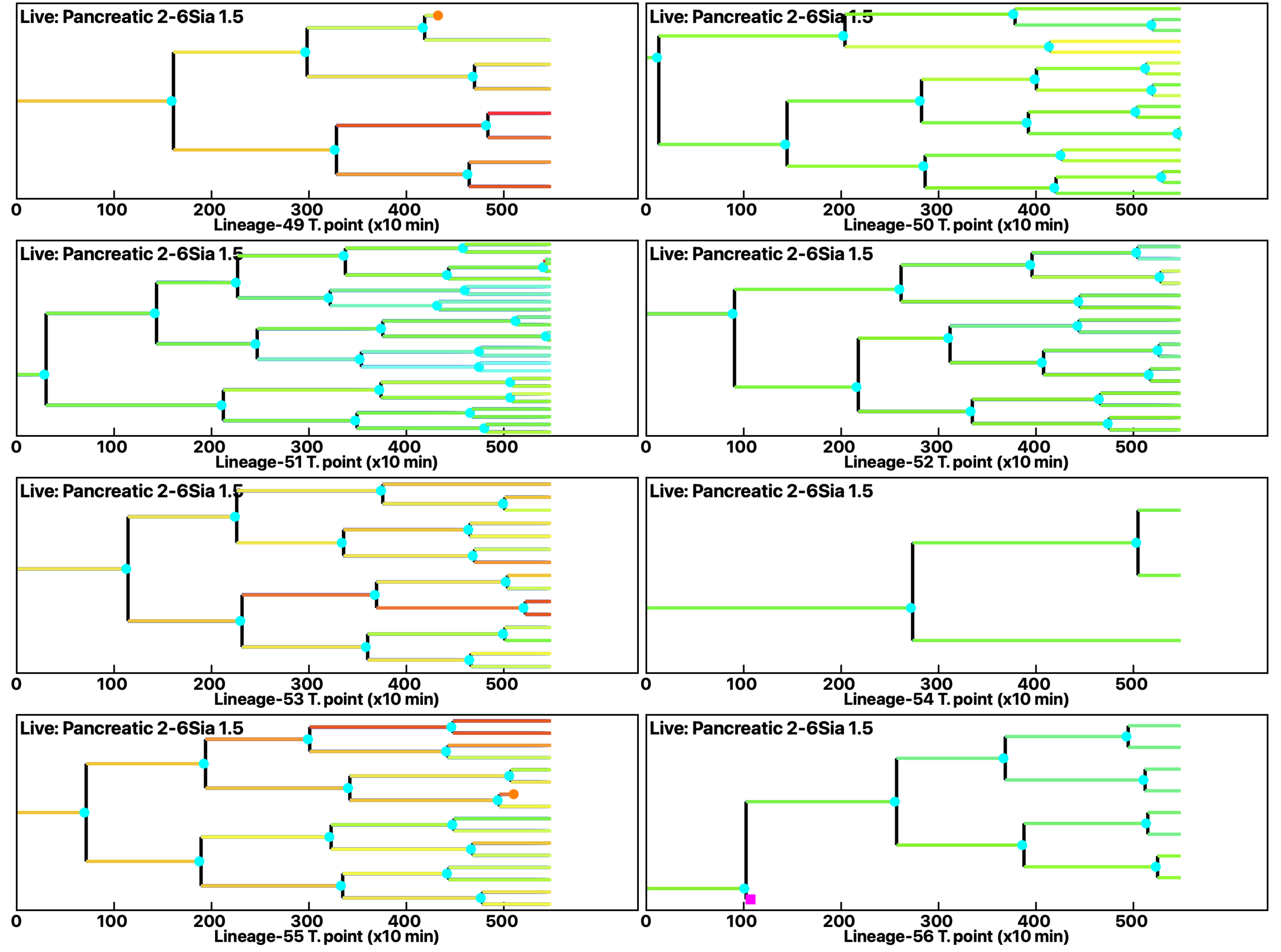

Analysis: Simulation, Treat.: Pancreatic 2-6Sia 1.5

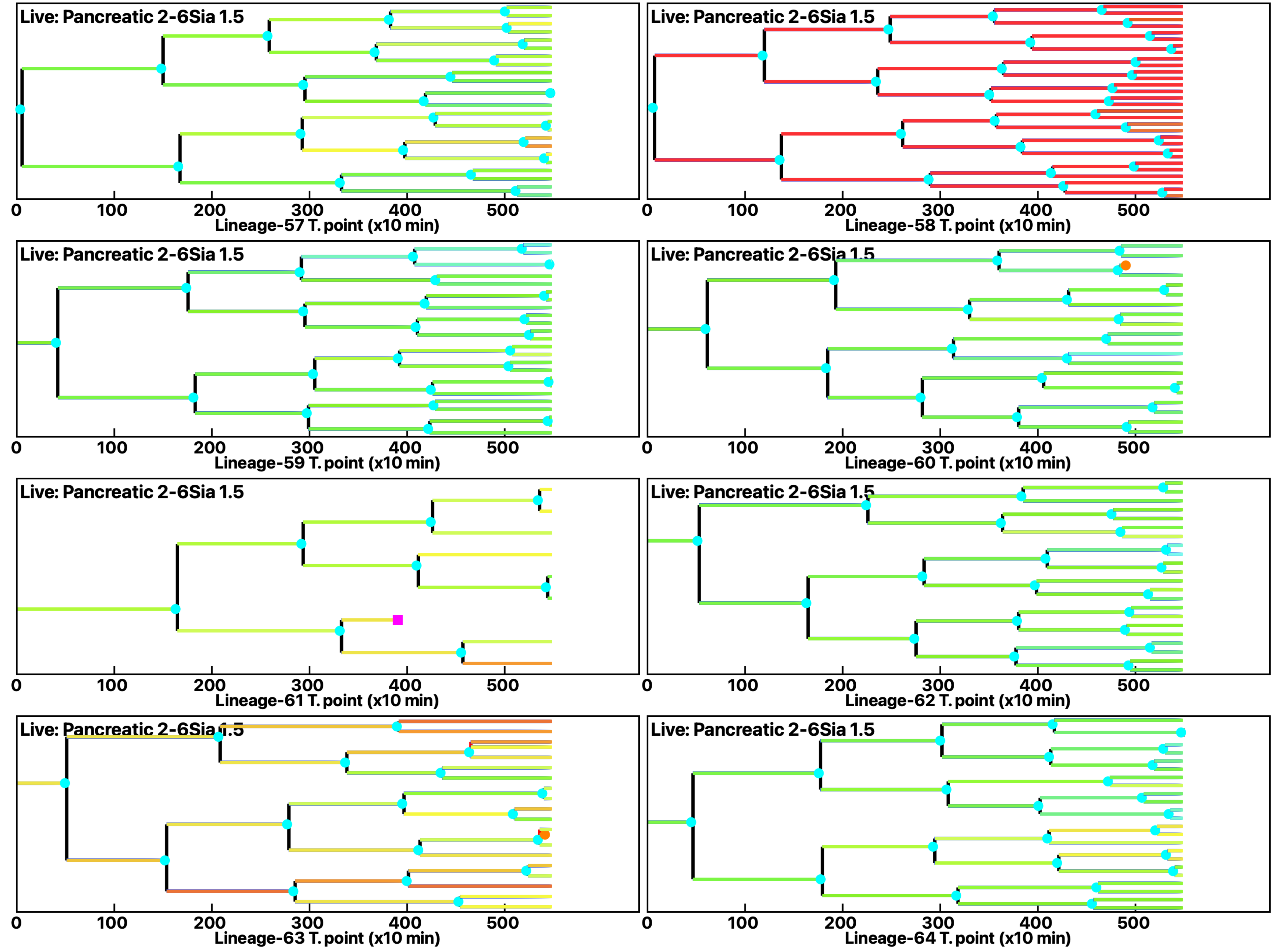

Analysis: Simulation, Treat.: Pancreatic 2-6Sia 1.5

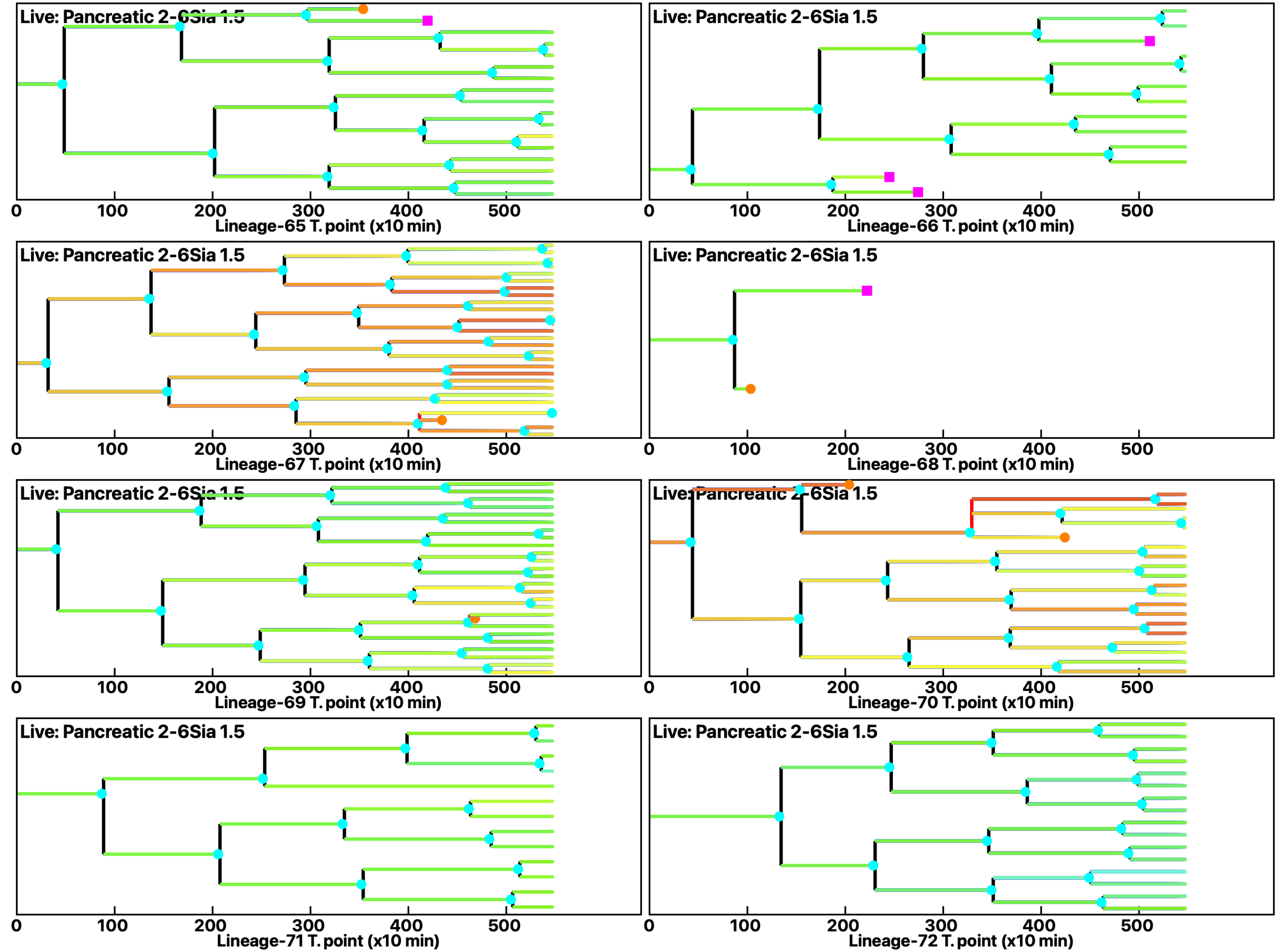

Analysis: Simulation, Treat.: Pancreatic 2-6Sia 1.5

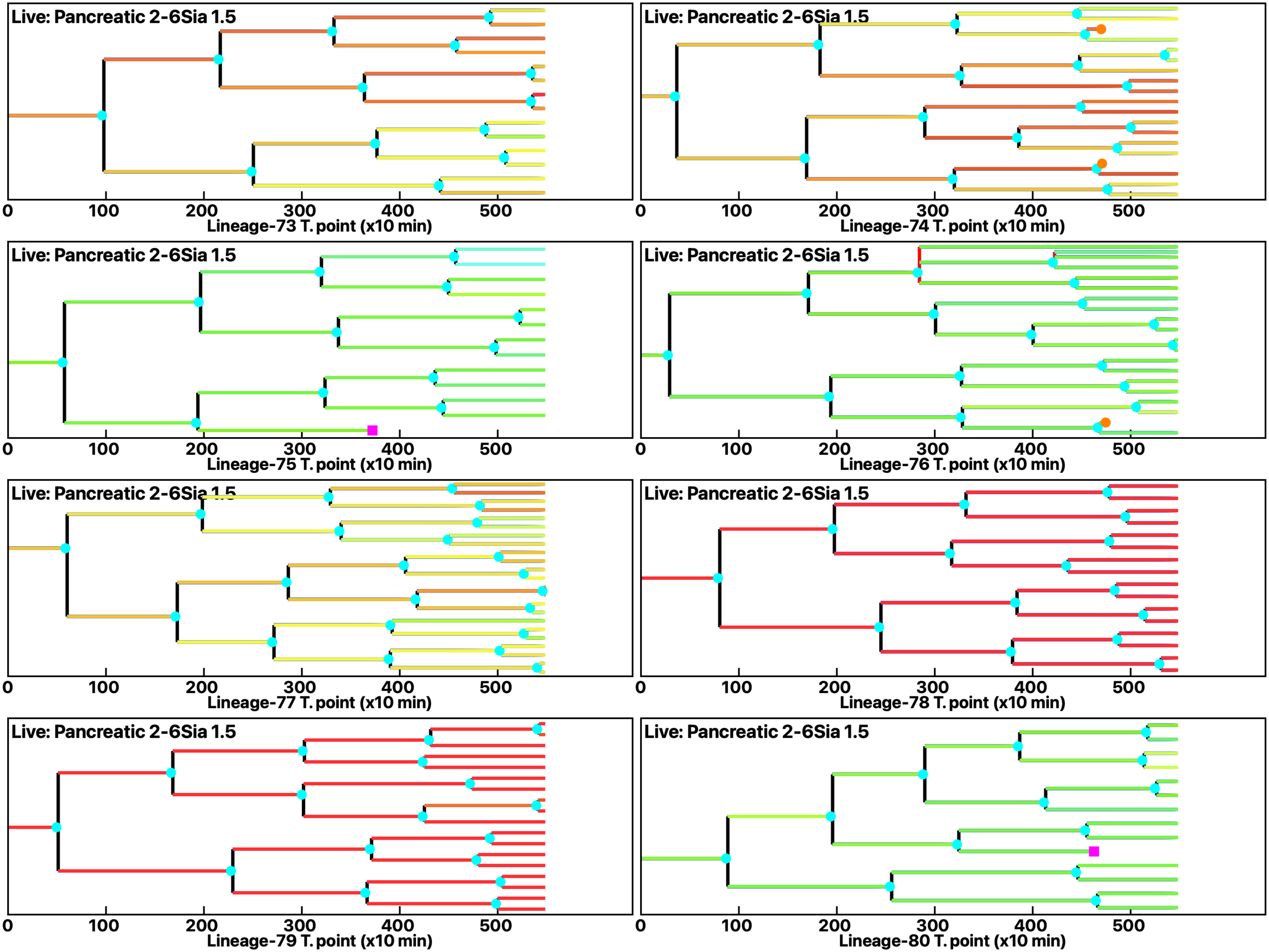

Analysis: Simulation, Treat.: Pancreatic 2-6Sia 1.5

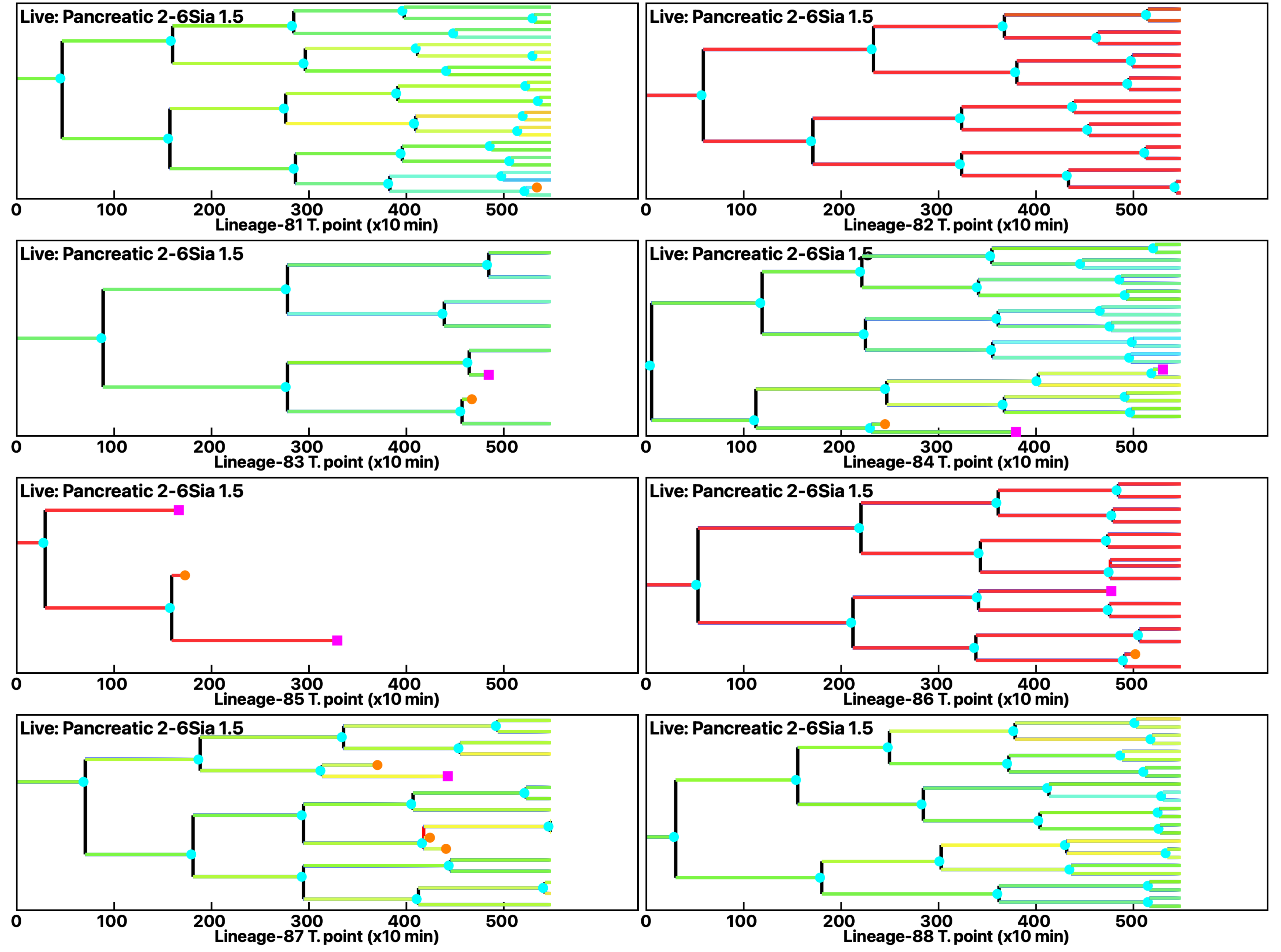

Analysis: Simulation, Treat.: Pancreatic 2-6Sia 1.5

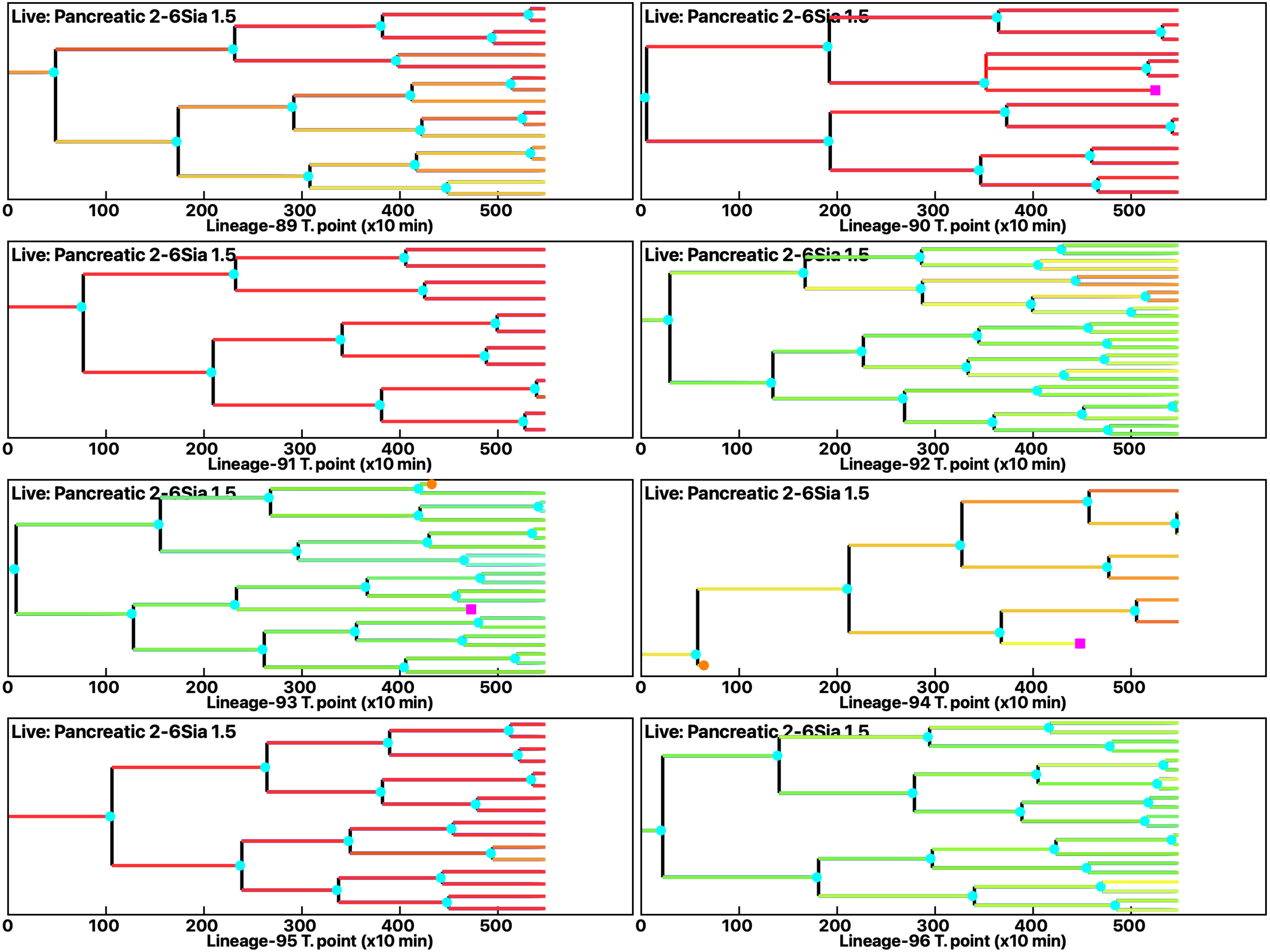

Analysis: Simulation, Treat.: Pancreatic 2-6Sia 1.5

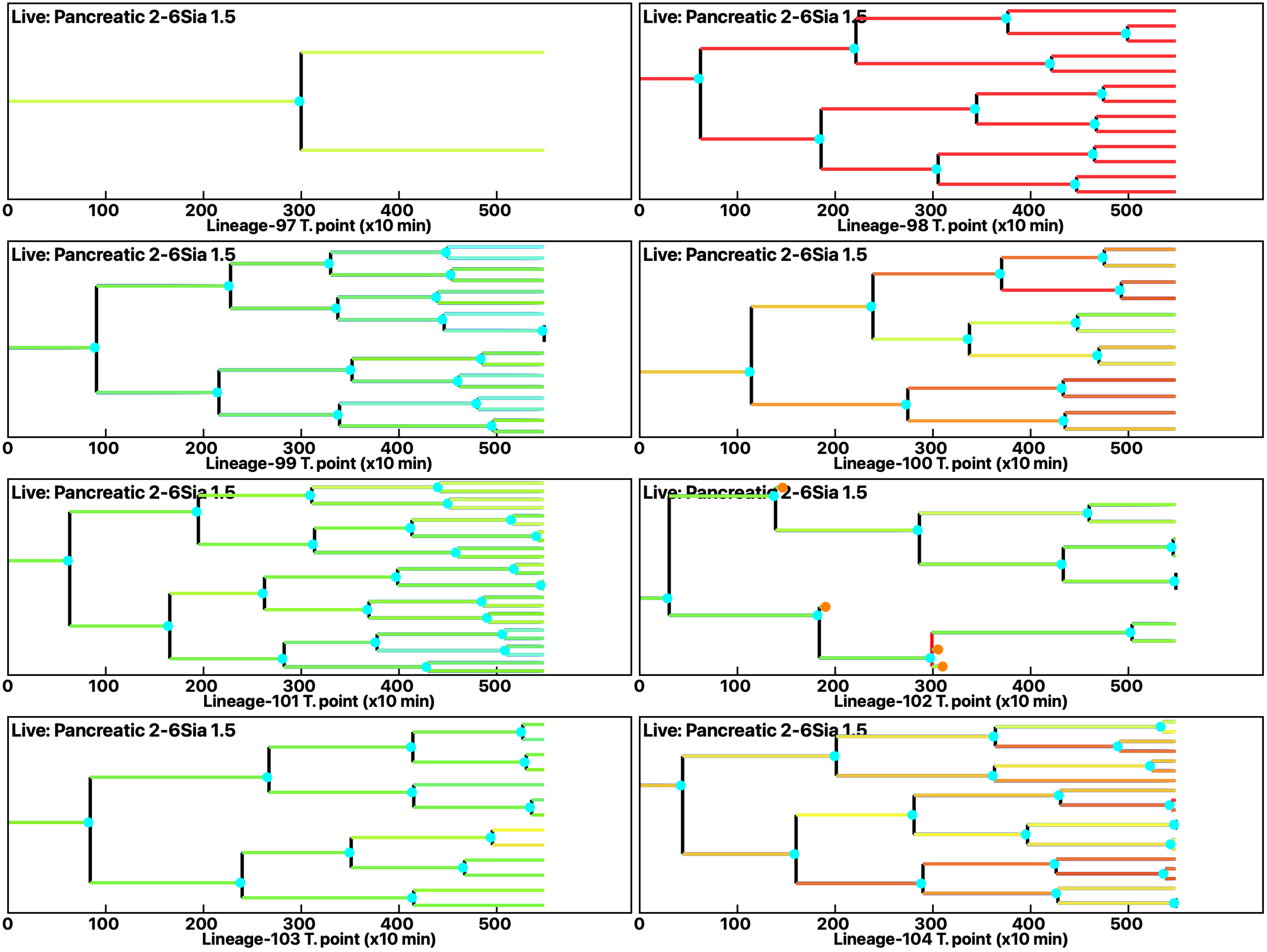

Analysis: Simulation, Treat.: Pancreatic 2-6Sia 1.5

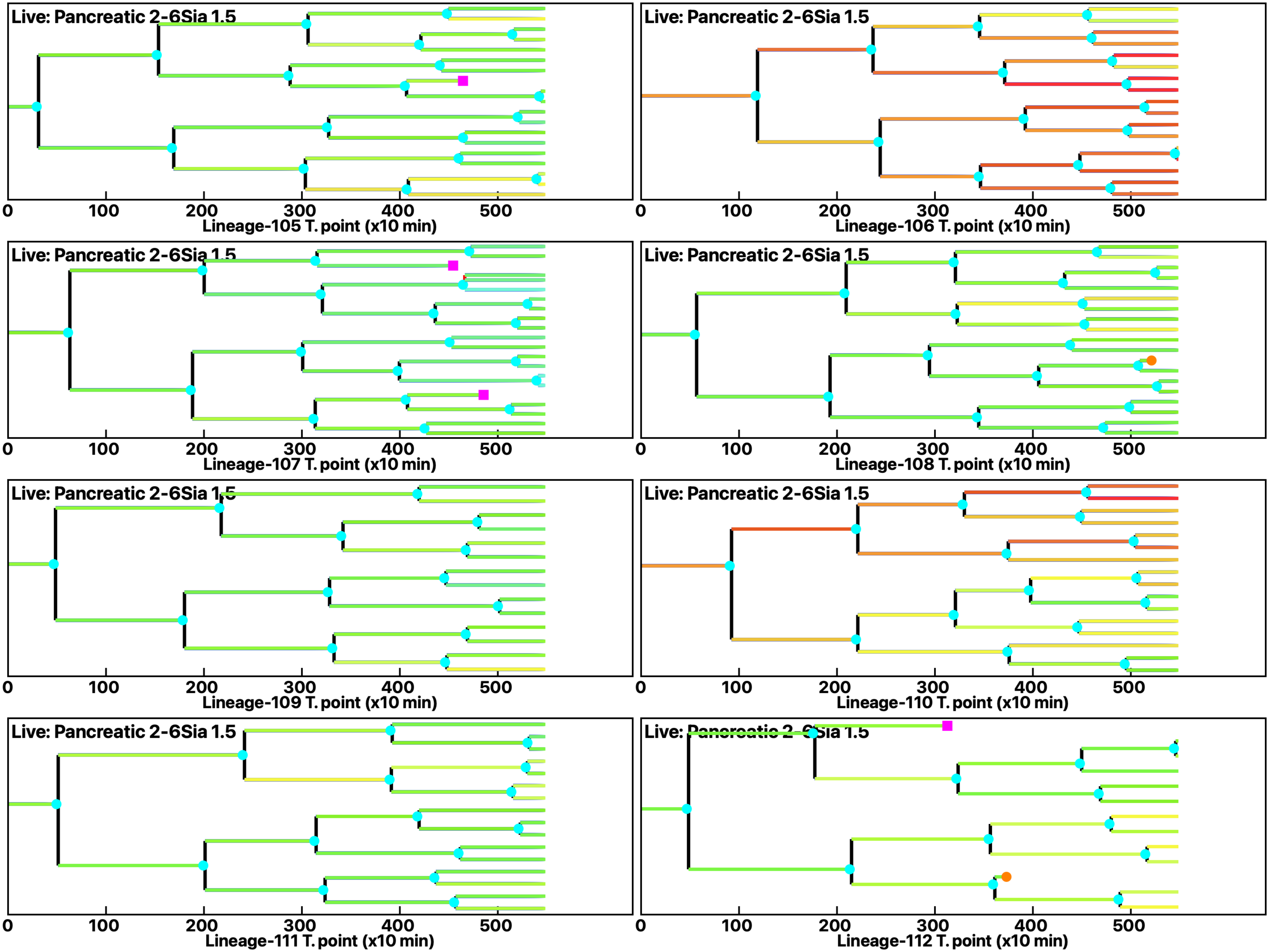

Analysis: Simulation, Treat.: Pancreatic 2-6Sia 1.5

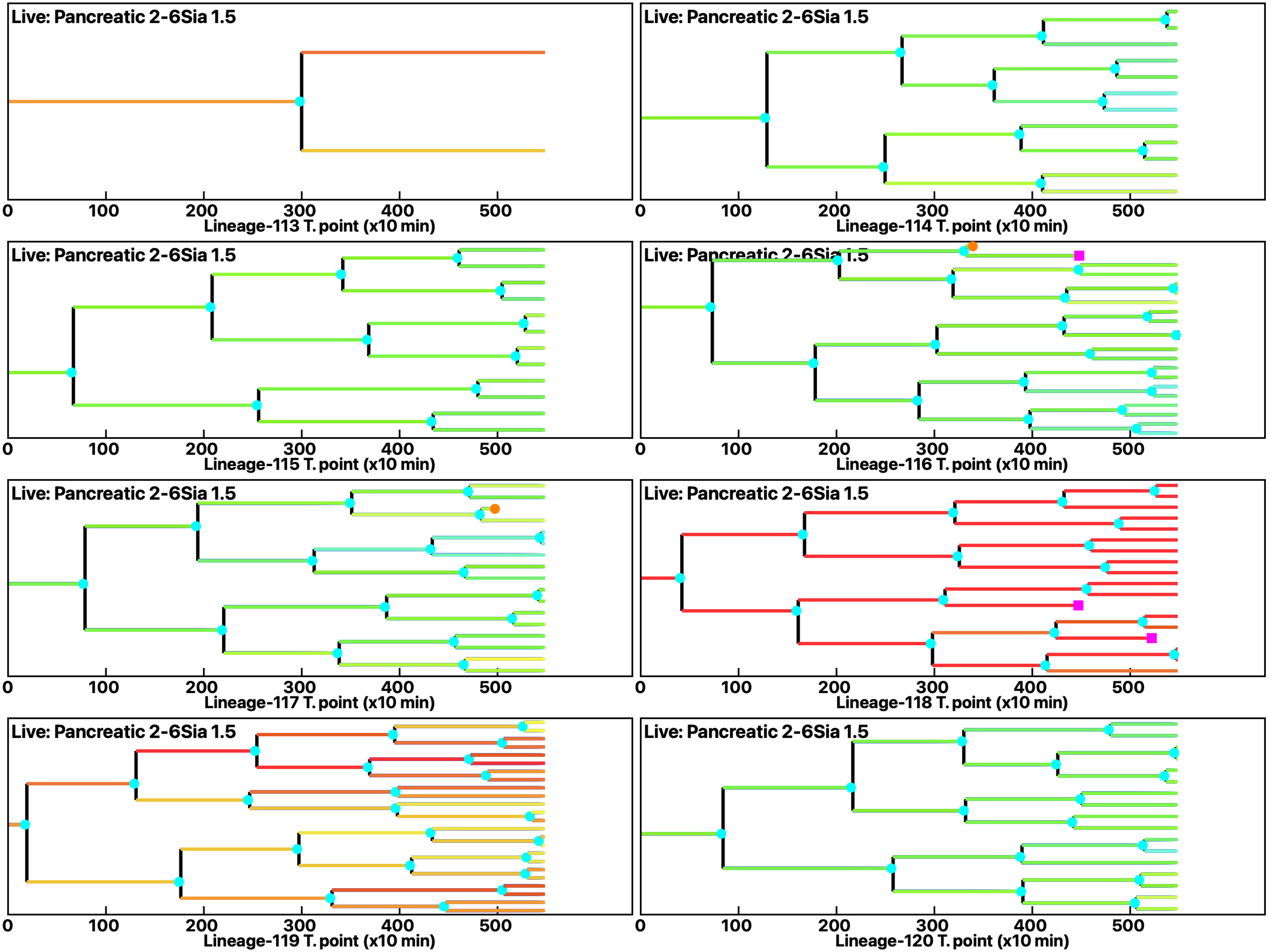

Analysis: Simulation, Treat.: Pancreatic 2-6Sia 1.5

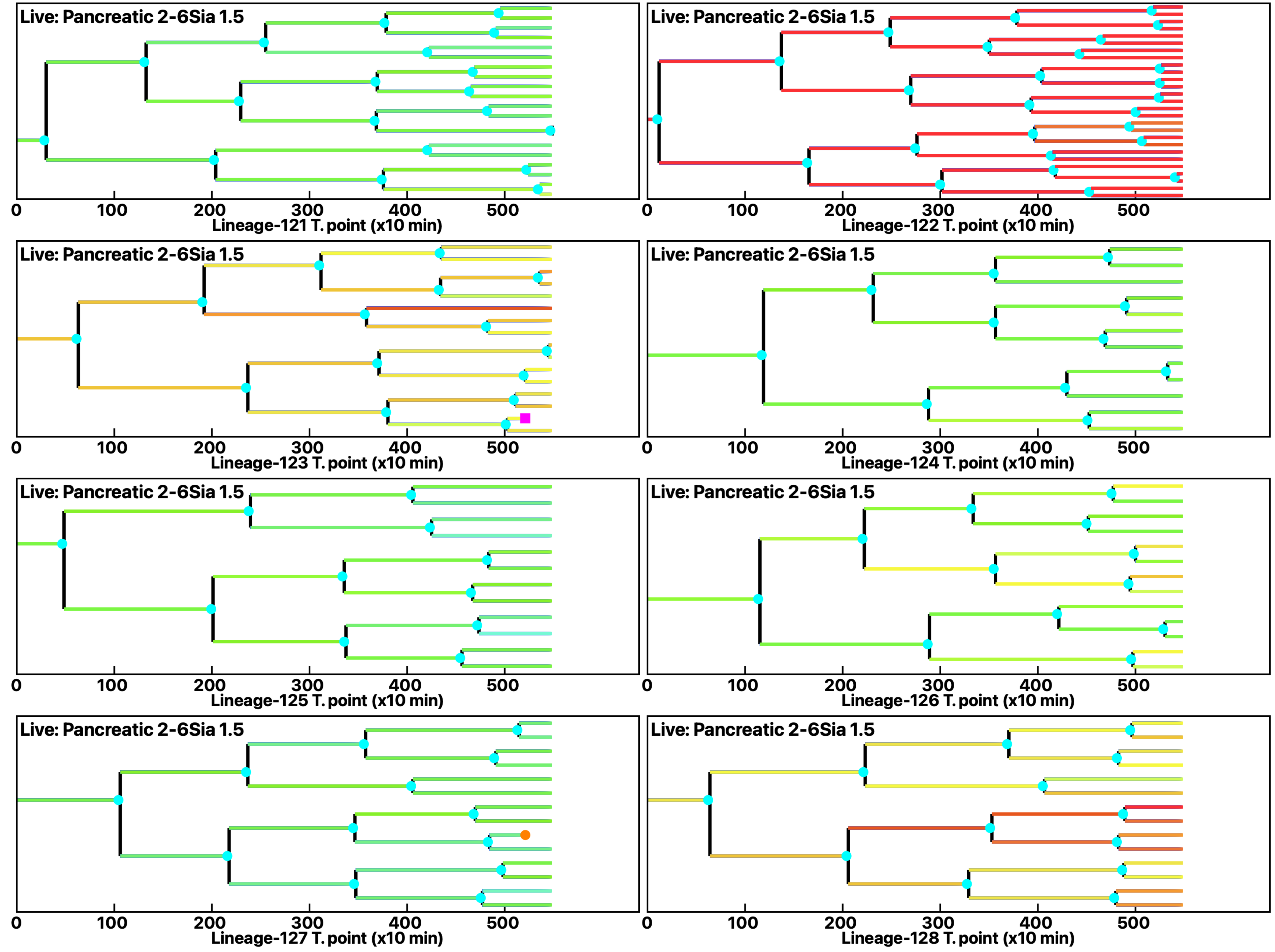

Analysis: Simulation, Treat.: Pancreatic 2-6Sia 1.5

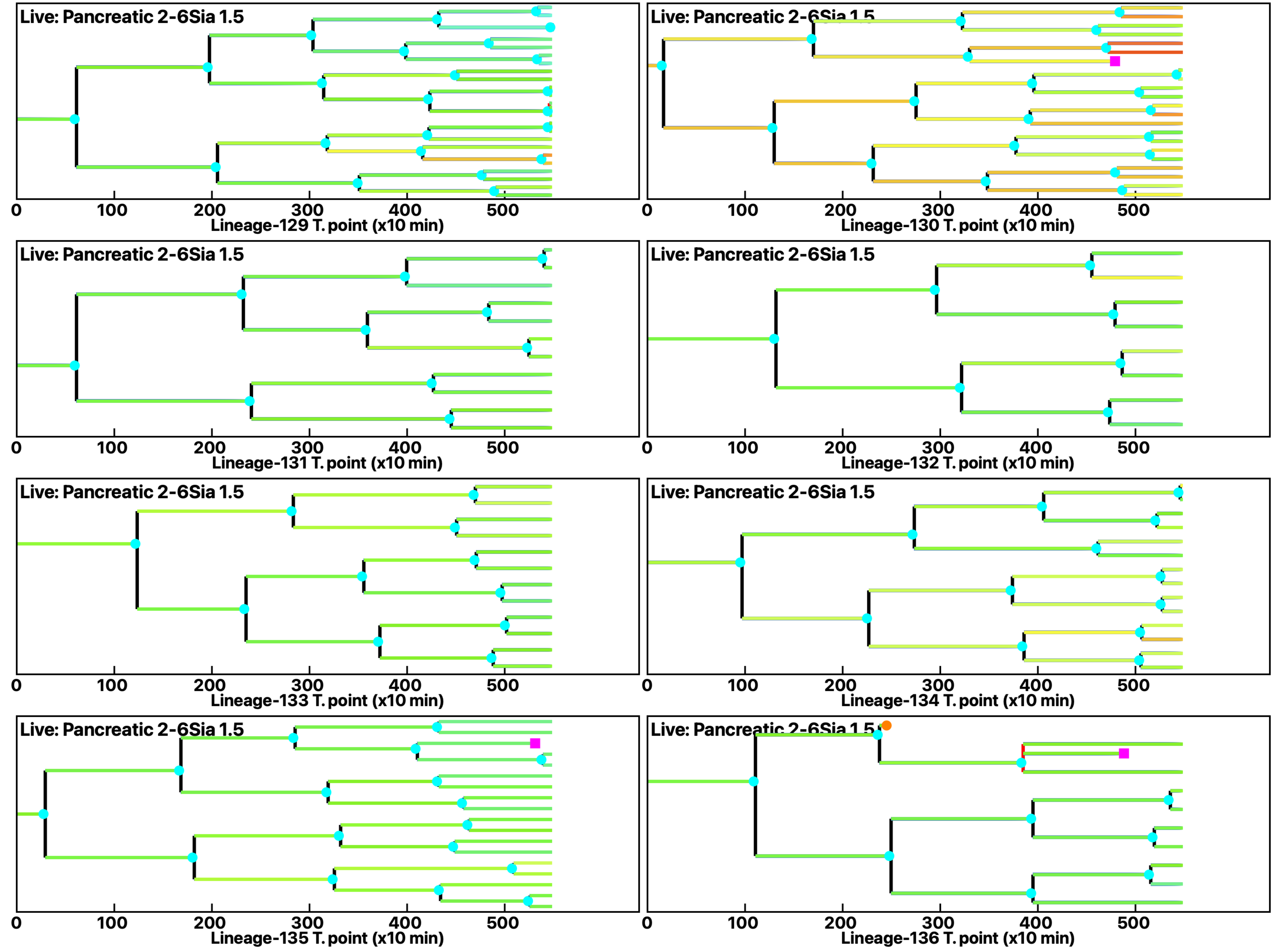

**Analysis: Simulation, Treat.: Pancreatic 2-6Sia 1.5**

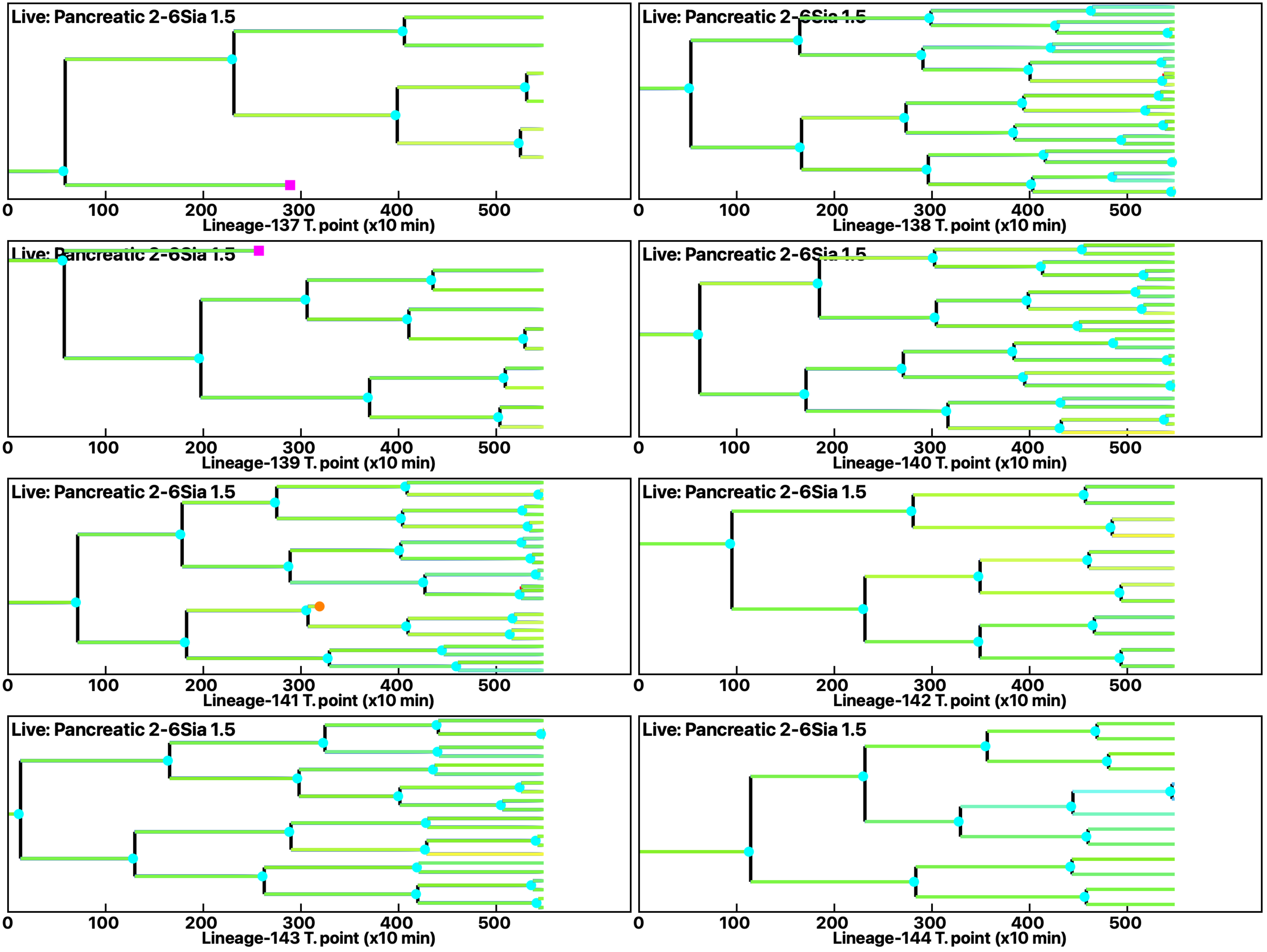

**Analysis: Simulation, Treat.: Pancreatic 2-6Sia 1.5**

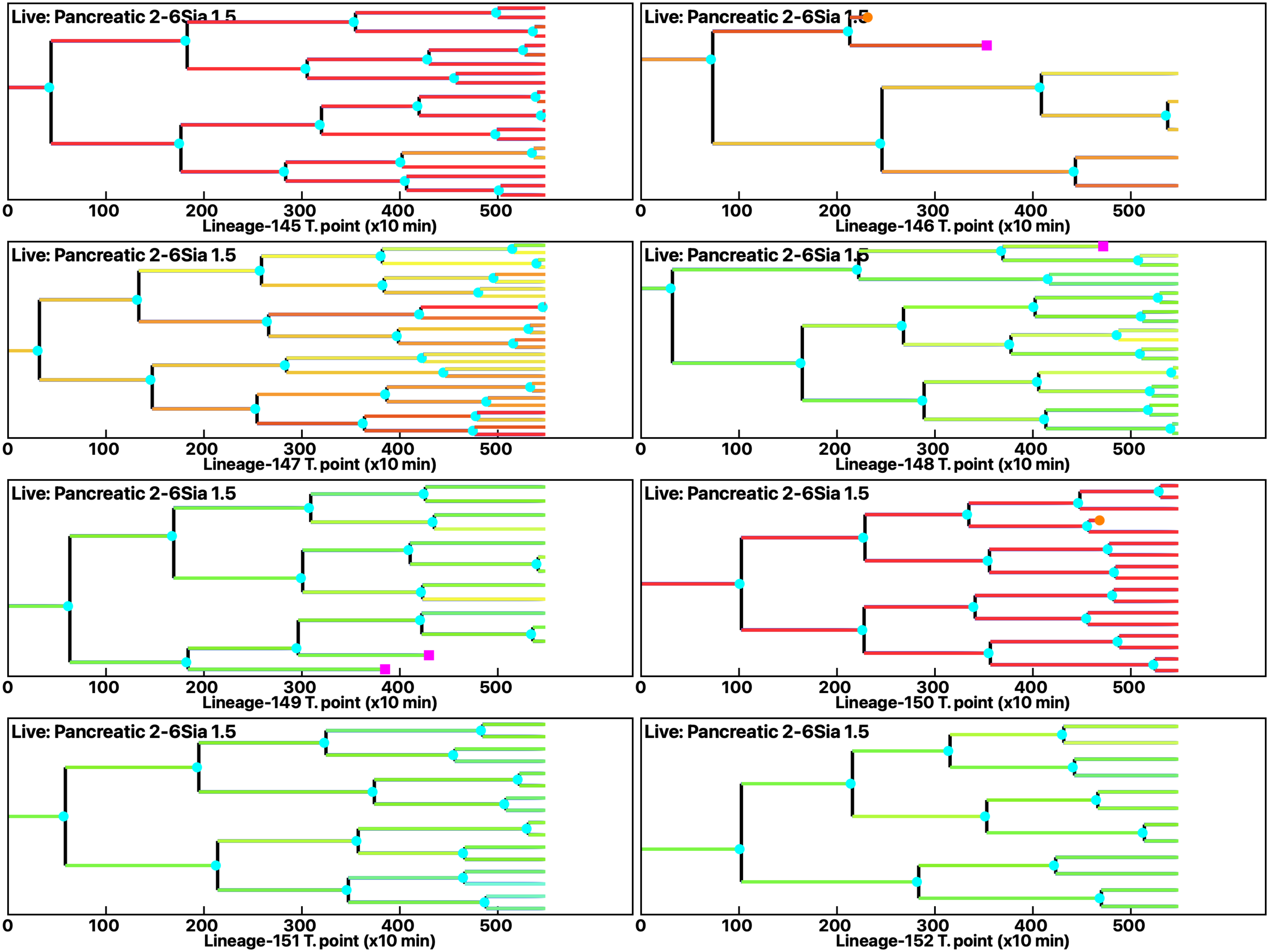

Analysis: Simulation, Treat.: Pancreatic 2-6Sia 1.5

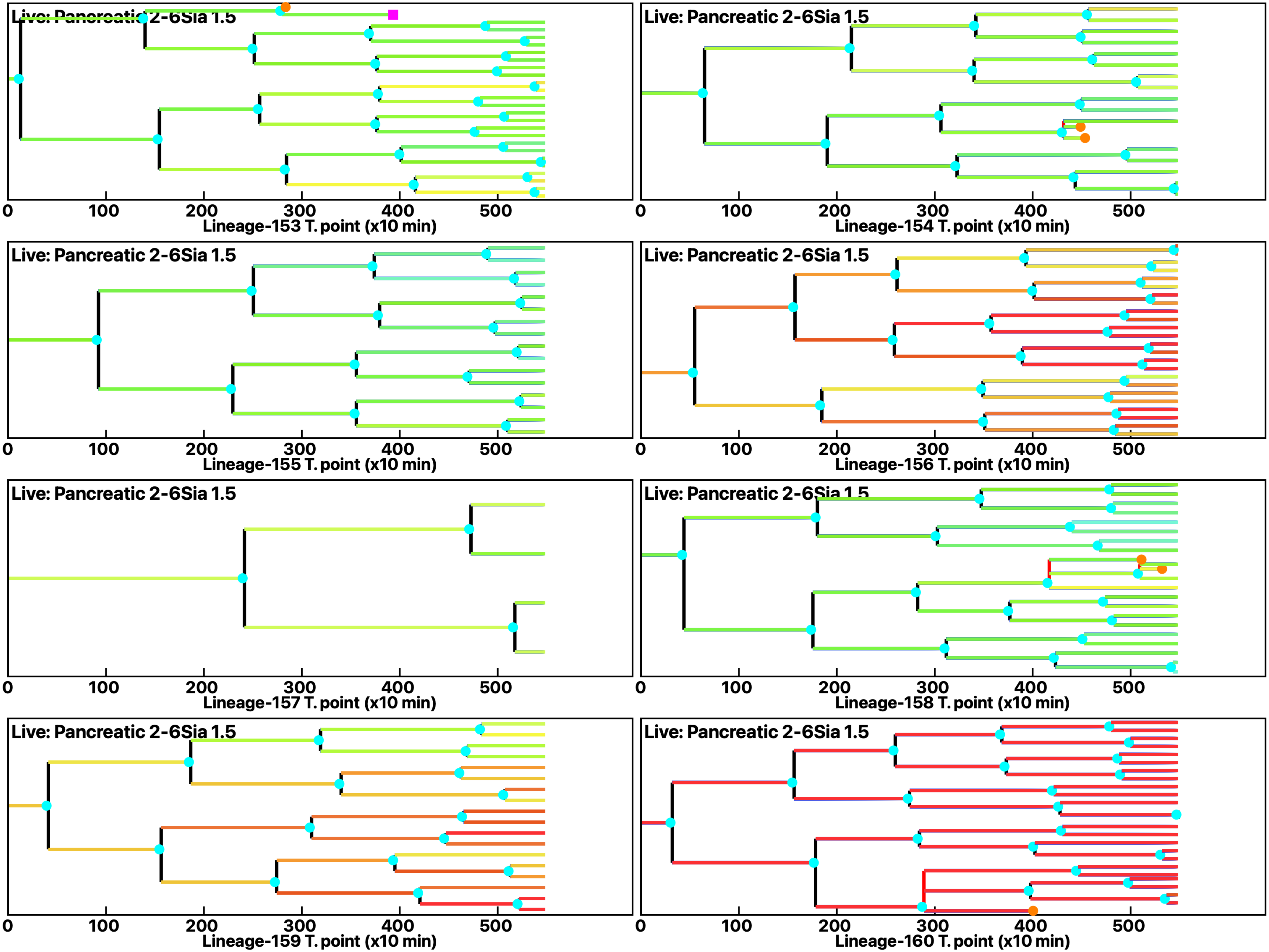

Analysis: Simulation, Treat.: Pancreatic 2-6Sia 1.5

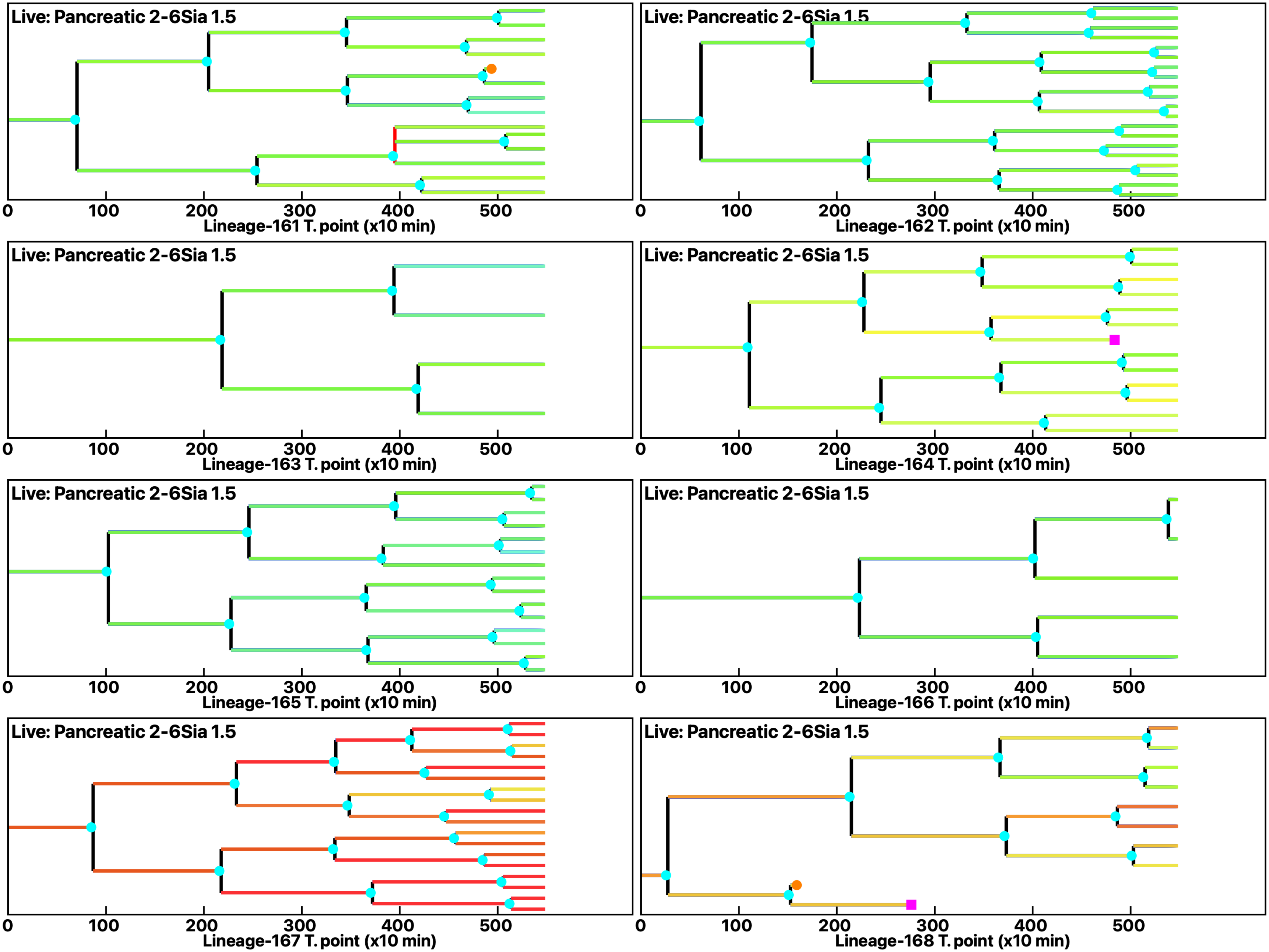

Analysis: Simulation, Treat.: Pancreatic 2-6Sia 1.5

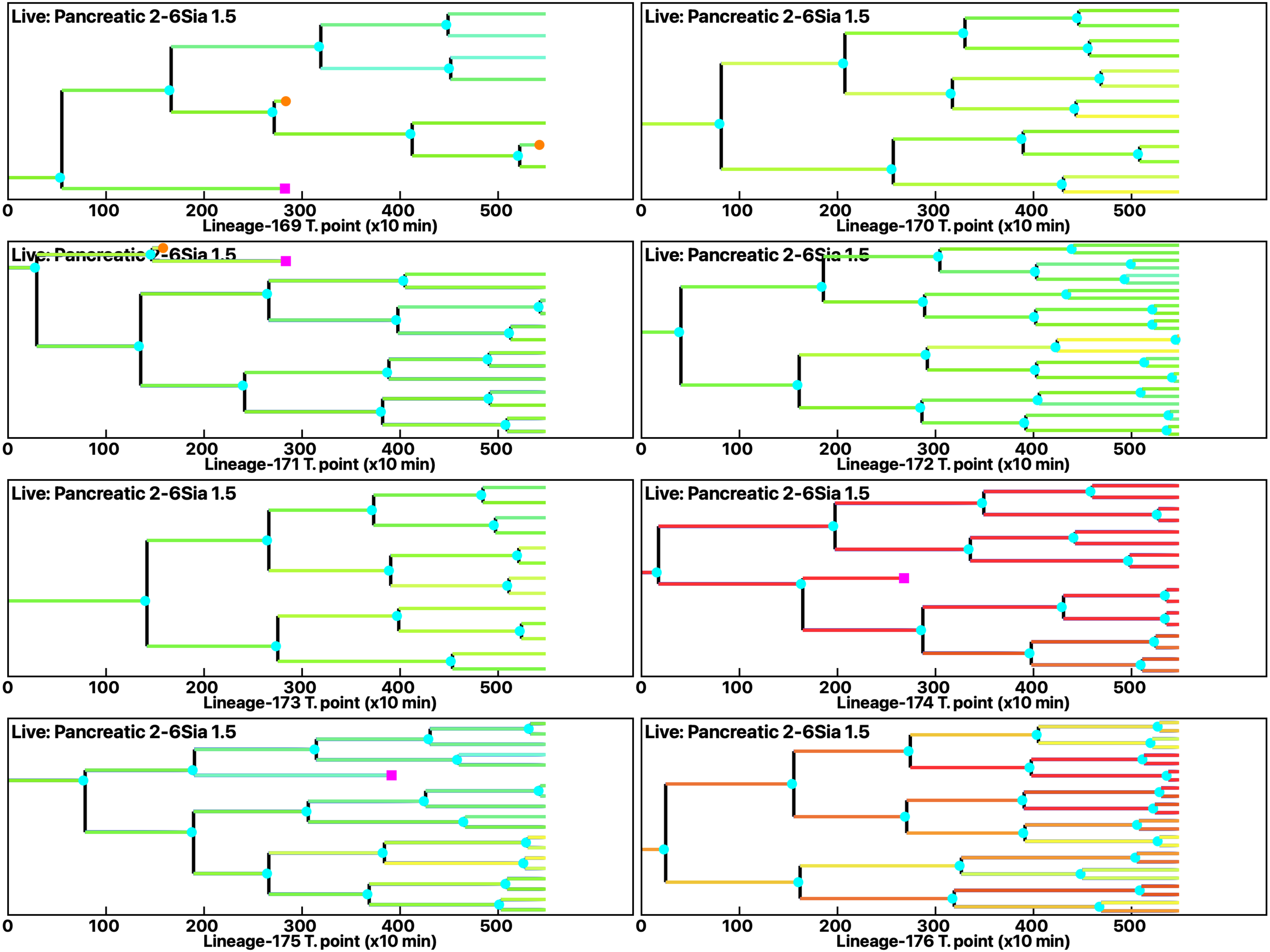

Analysis: Simulation, Treat.: Pancreatic 2-6Sia 1.5

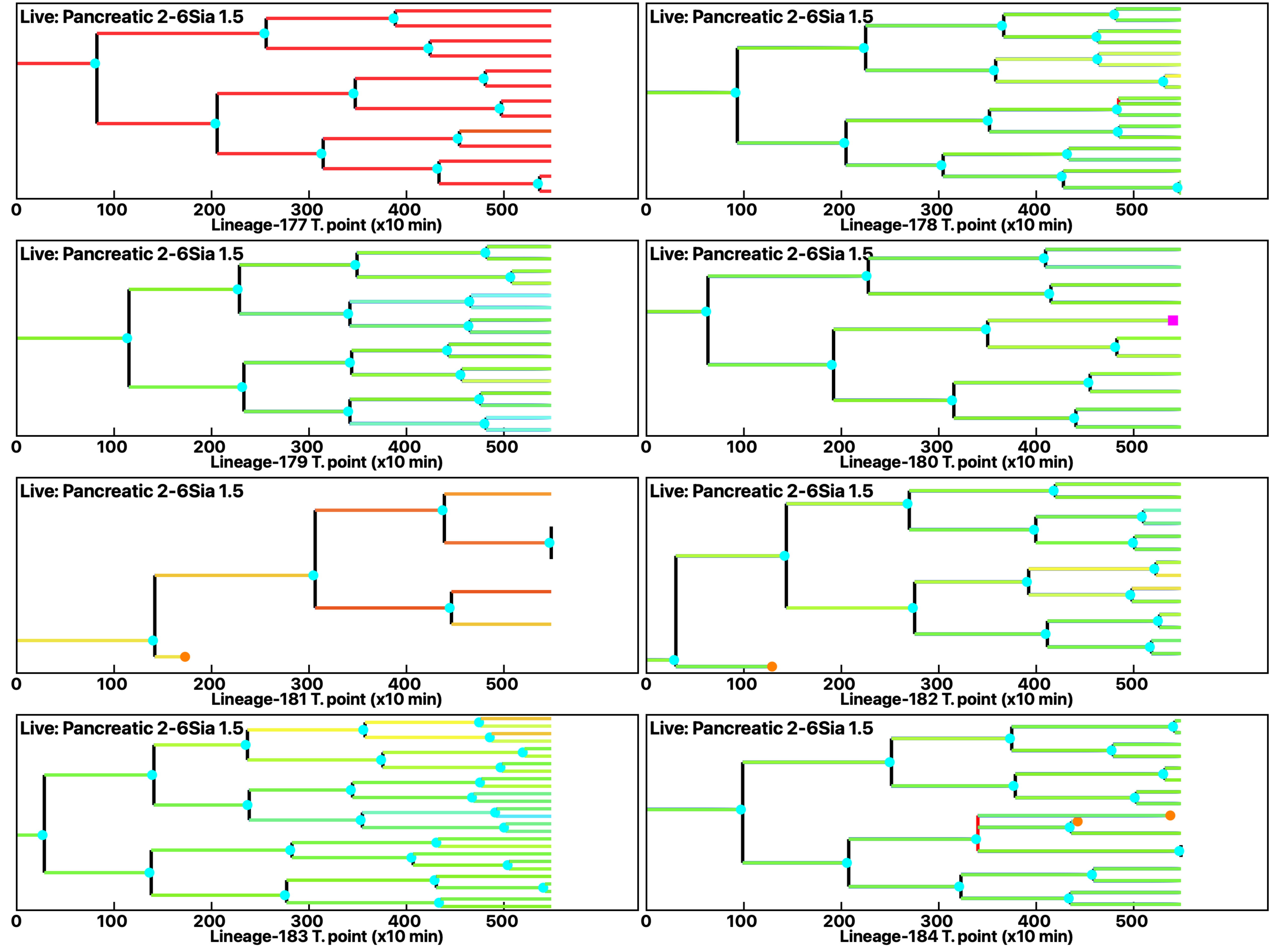

Analysis: Simulation, Treat.: Pancreatic 2-6Sia 1.5

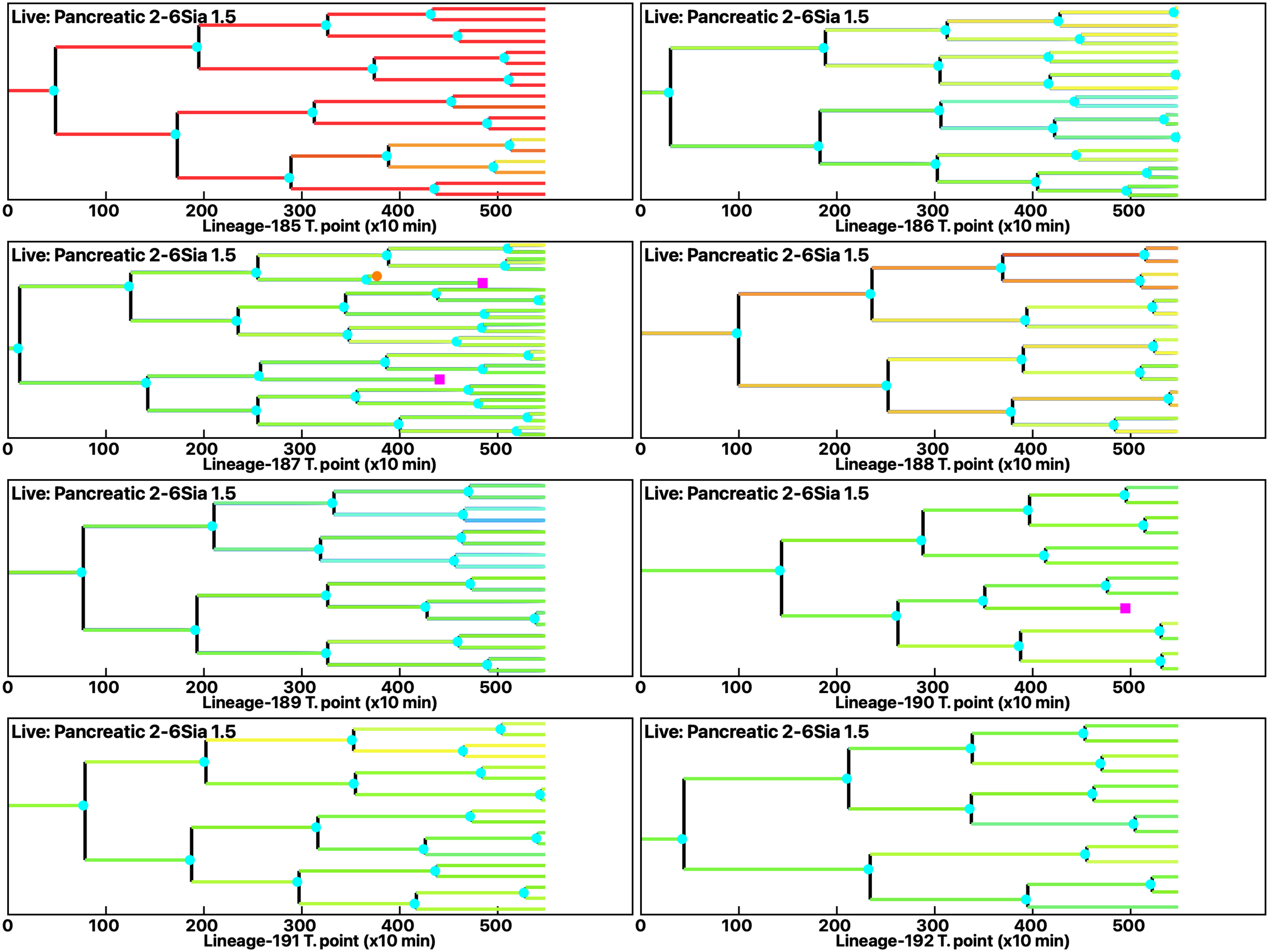

Analysis: Simulation, Treat.: Pancreatic 2-6Sia 1.5

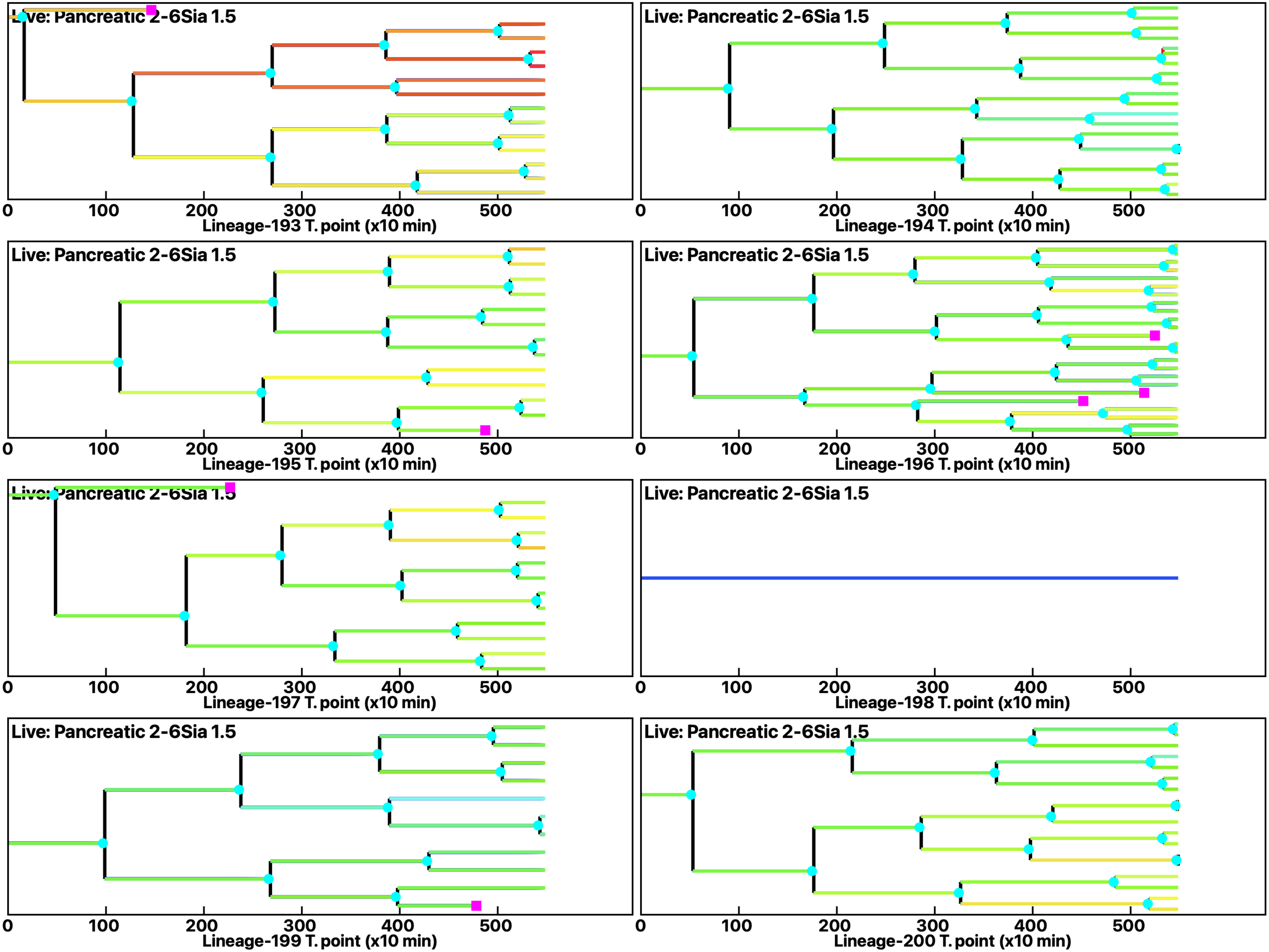

Analysis: Simulation, Treat.: Pancreatic 2-6Sia 1.5

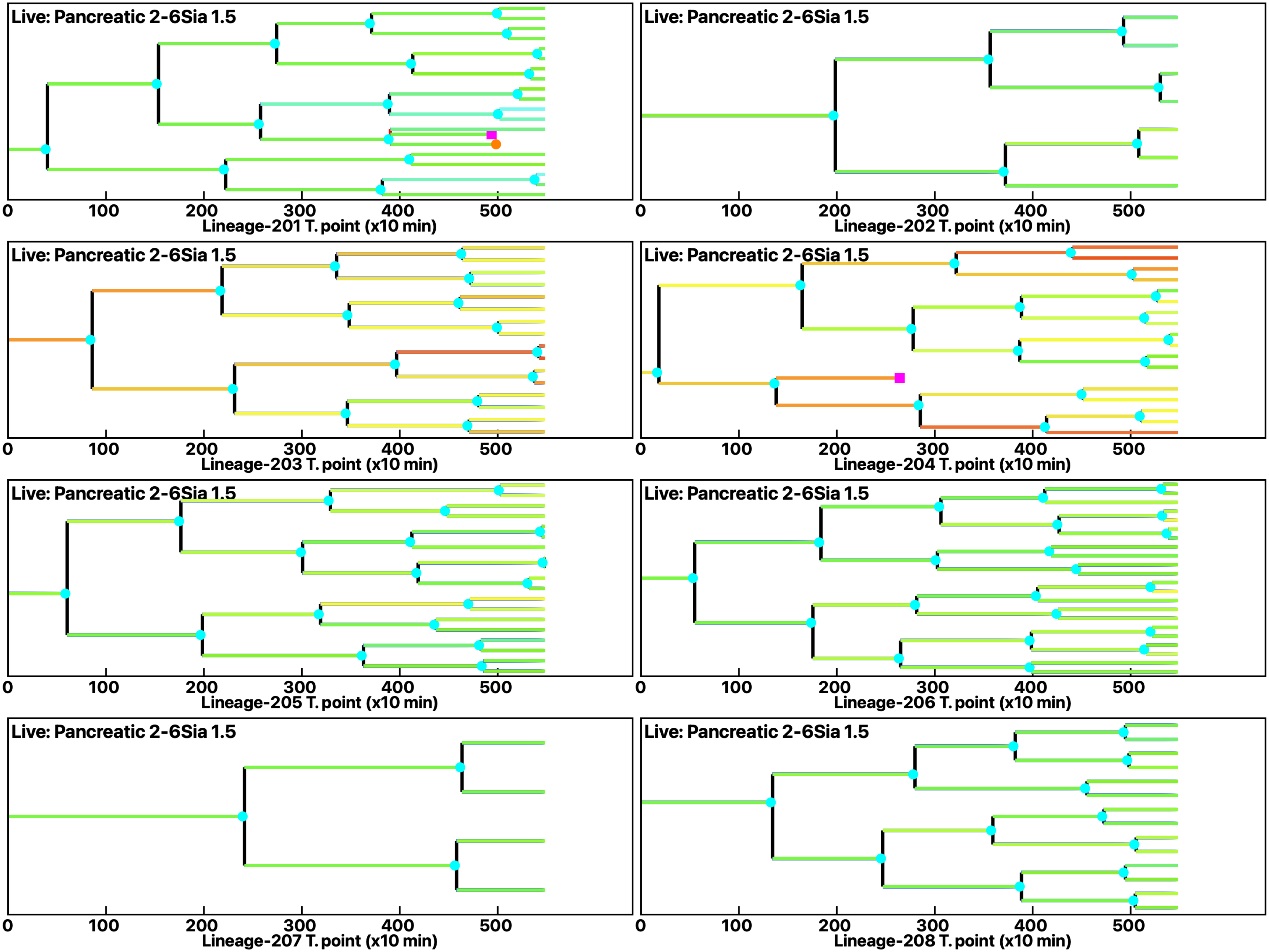

Analysis: Simulation, Treat.: Pancreatic 2-6Sia 1.5

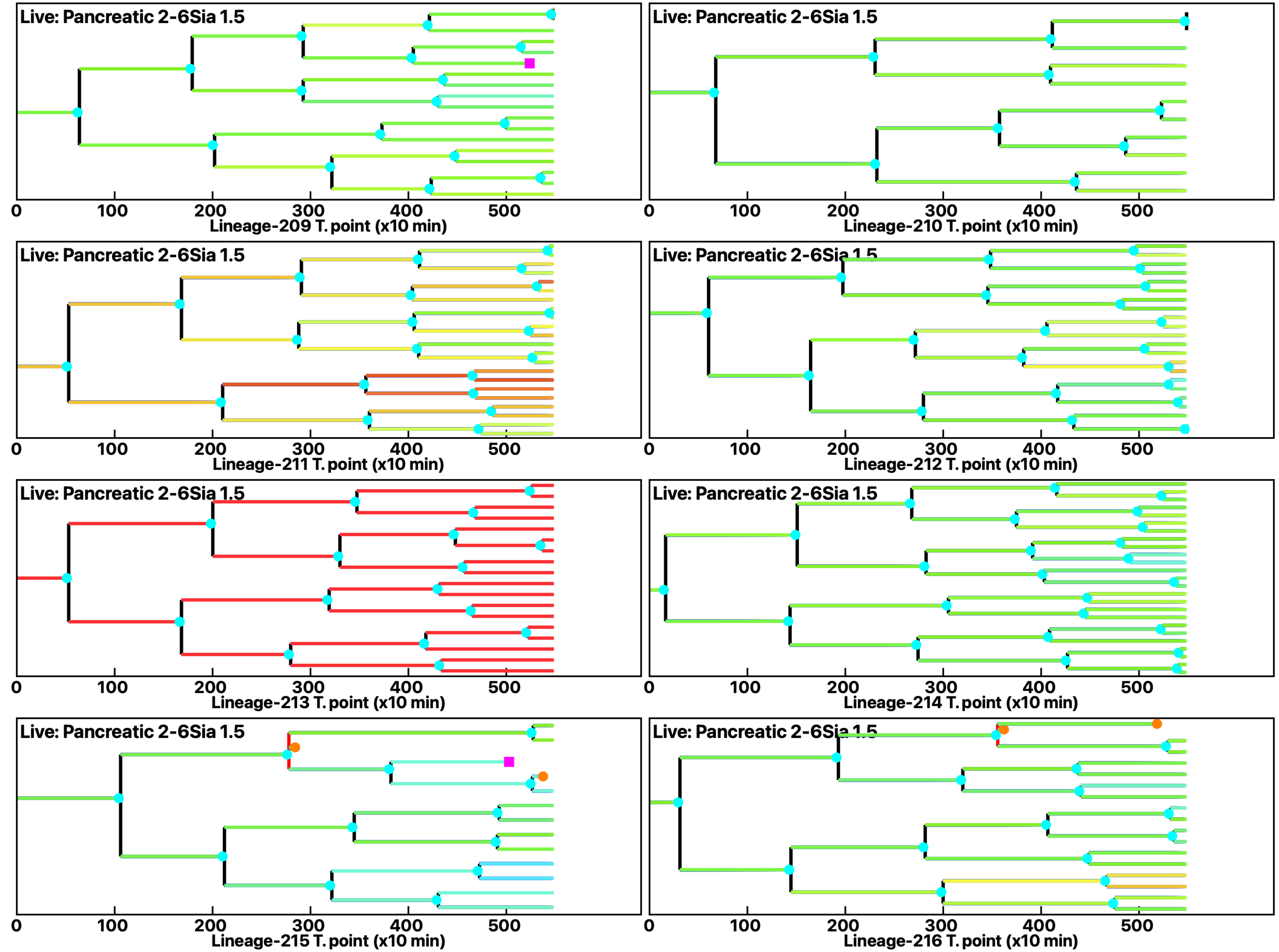

Analysis: Simulation, Treat.: Pancreatic 2-6Sia 1.5

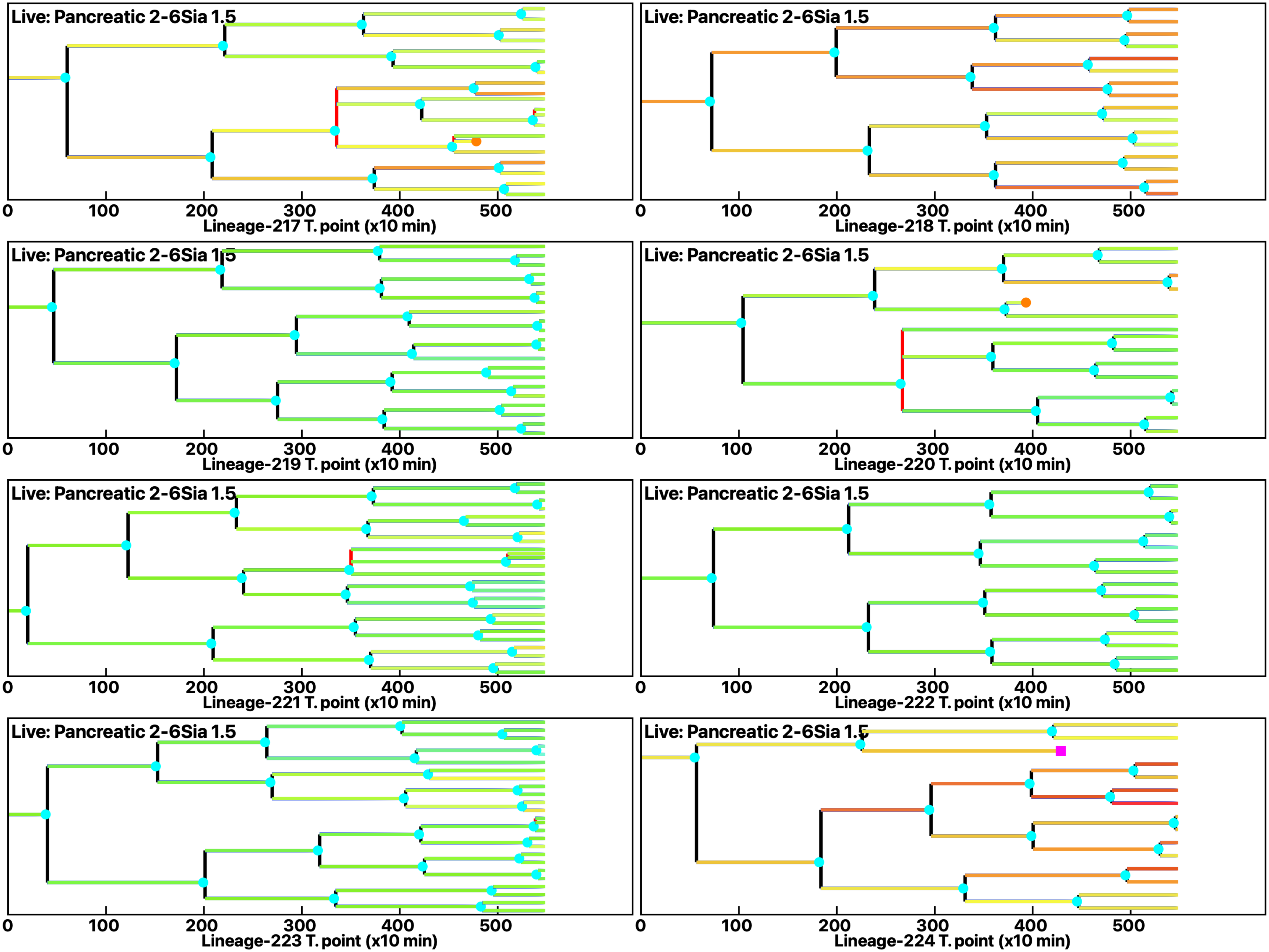

Analysis: Simulation, Treat.: Pancreatic 2-6Sia 1.5

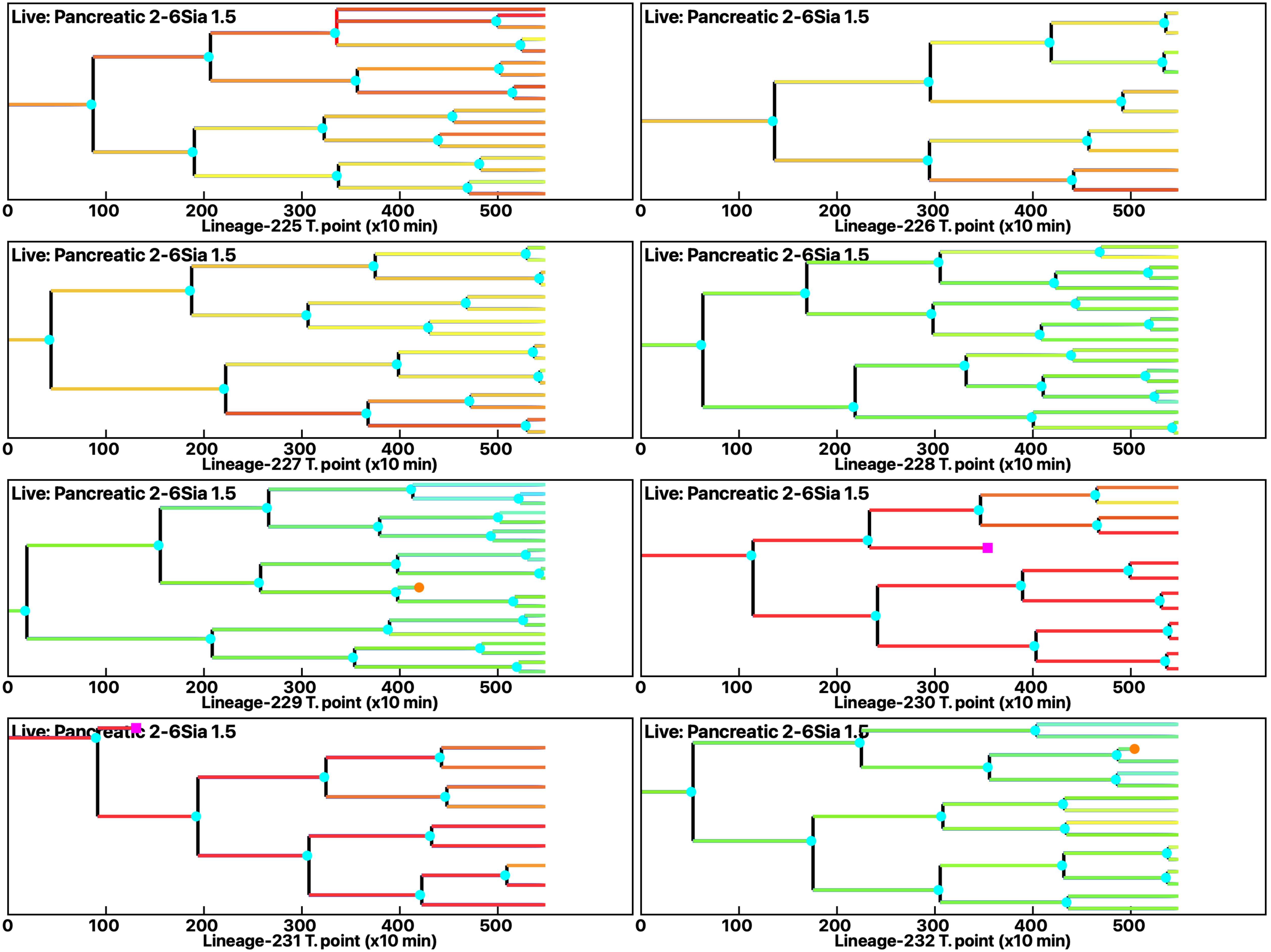

Analysis: Simulation, Treat.: Pancreatic 2-6Sia 1.5

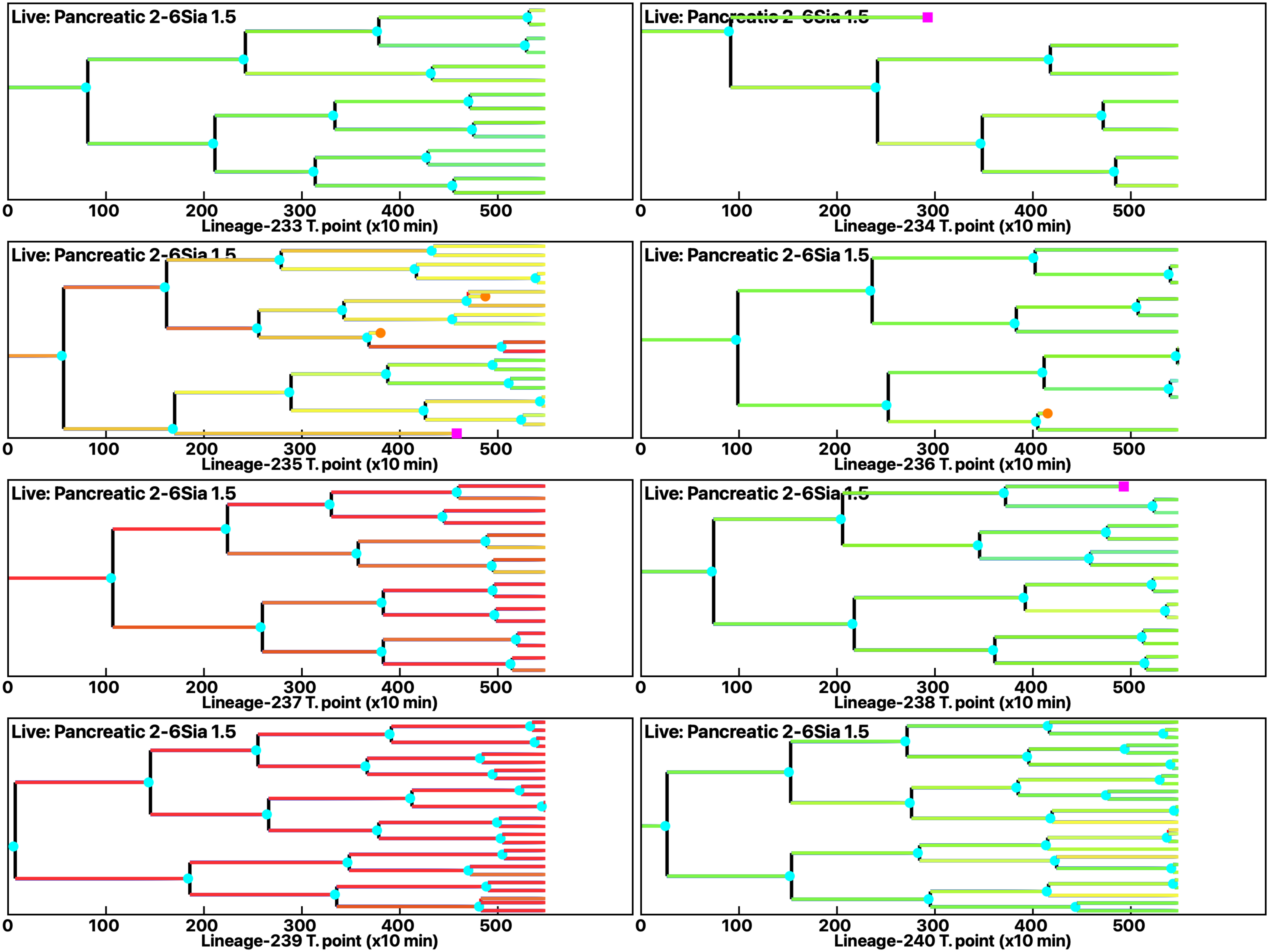

Analysis: Simulation, Treat.: Pancreatic 2-6Sia 1.5

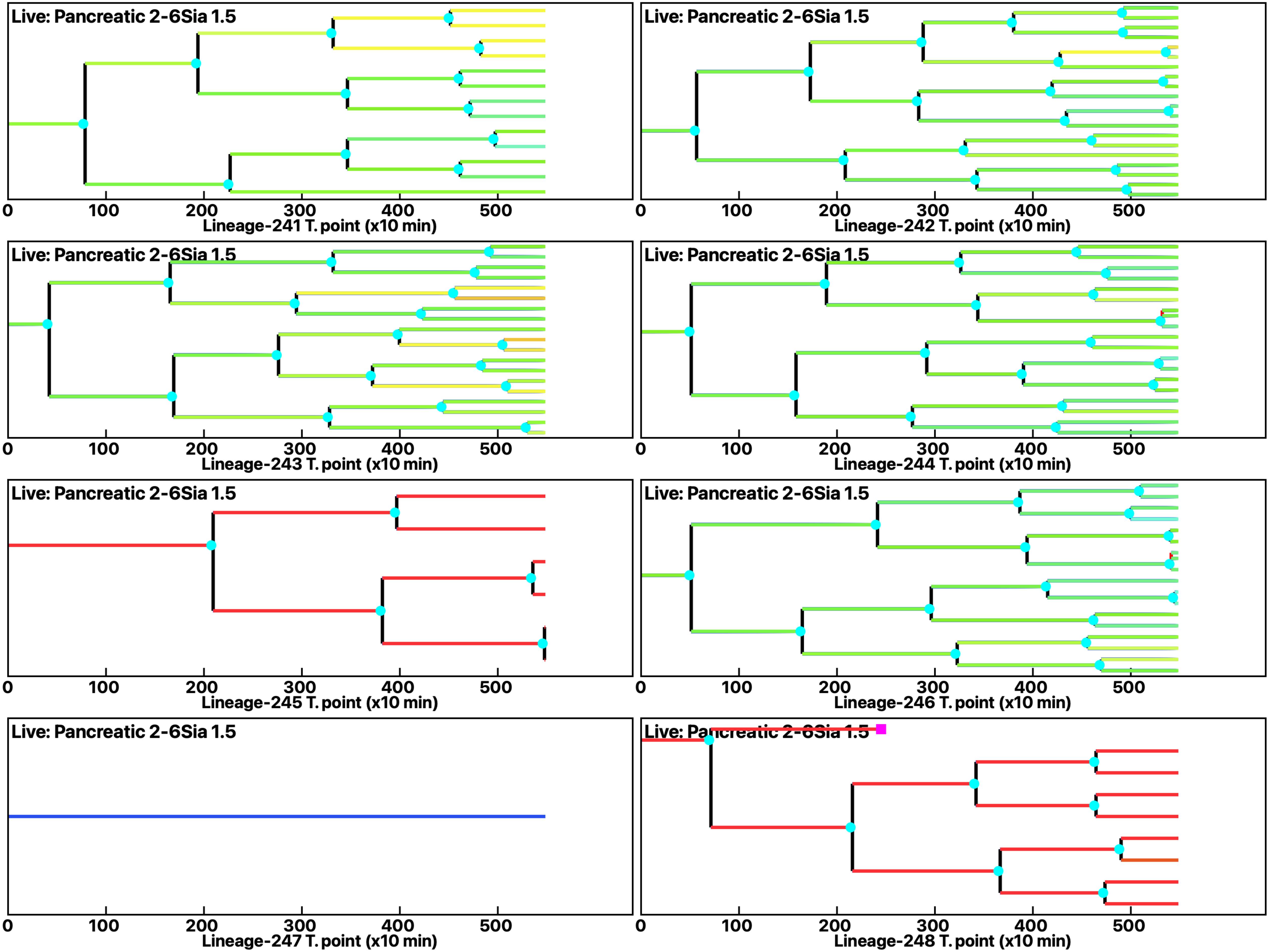

Analysis: Simulation, Treat.: Pancreatic 2-6Sia 1.5

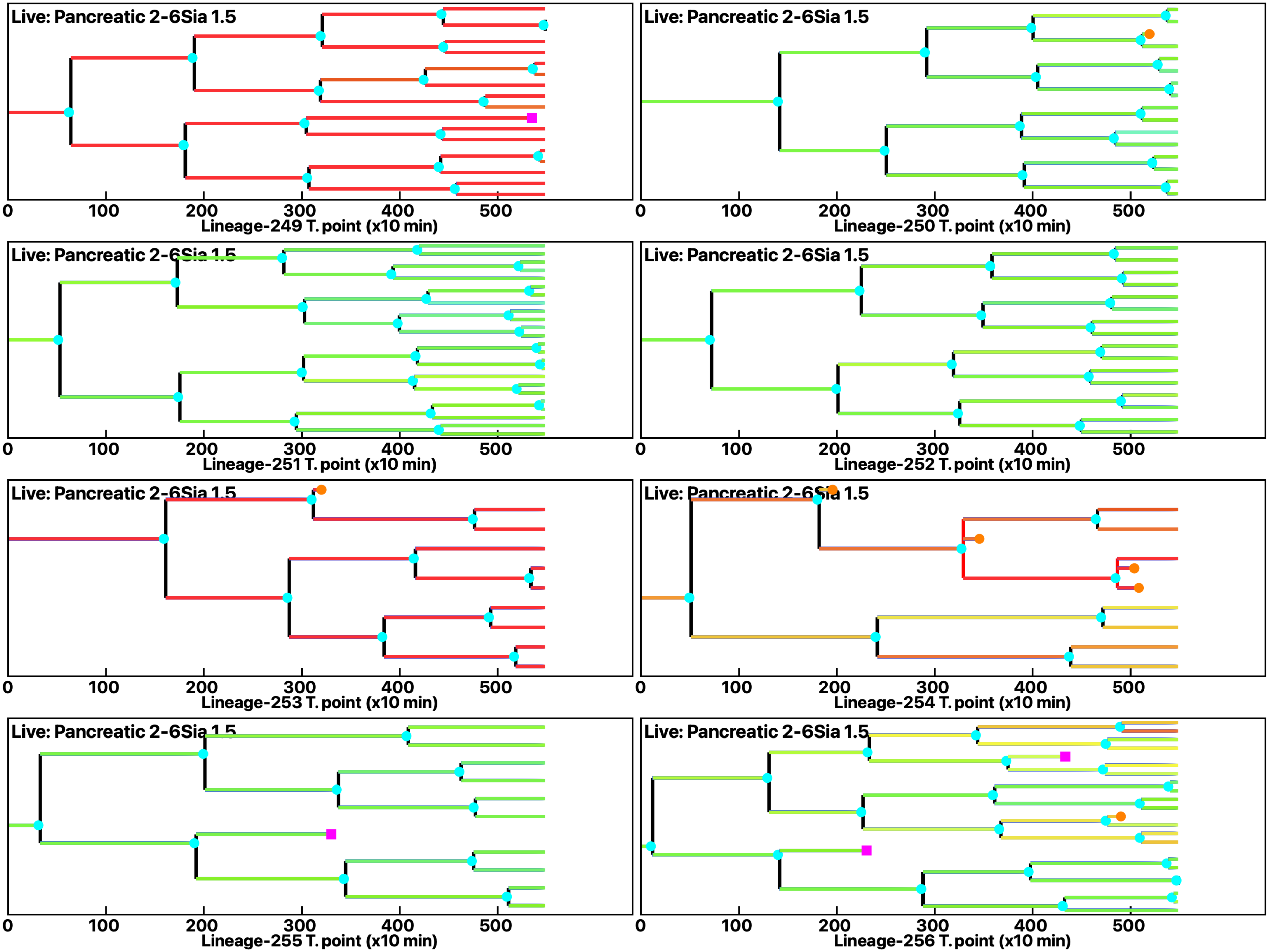

Analysis: Simulation, Treat.: Pancreatic 2-6Sia 1.5

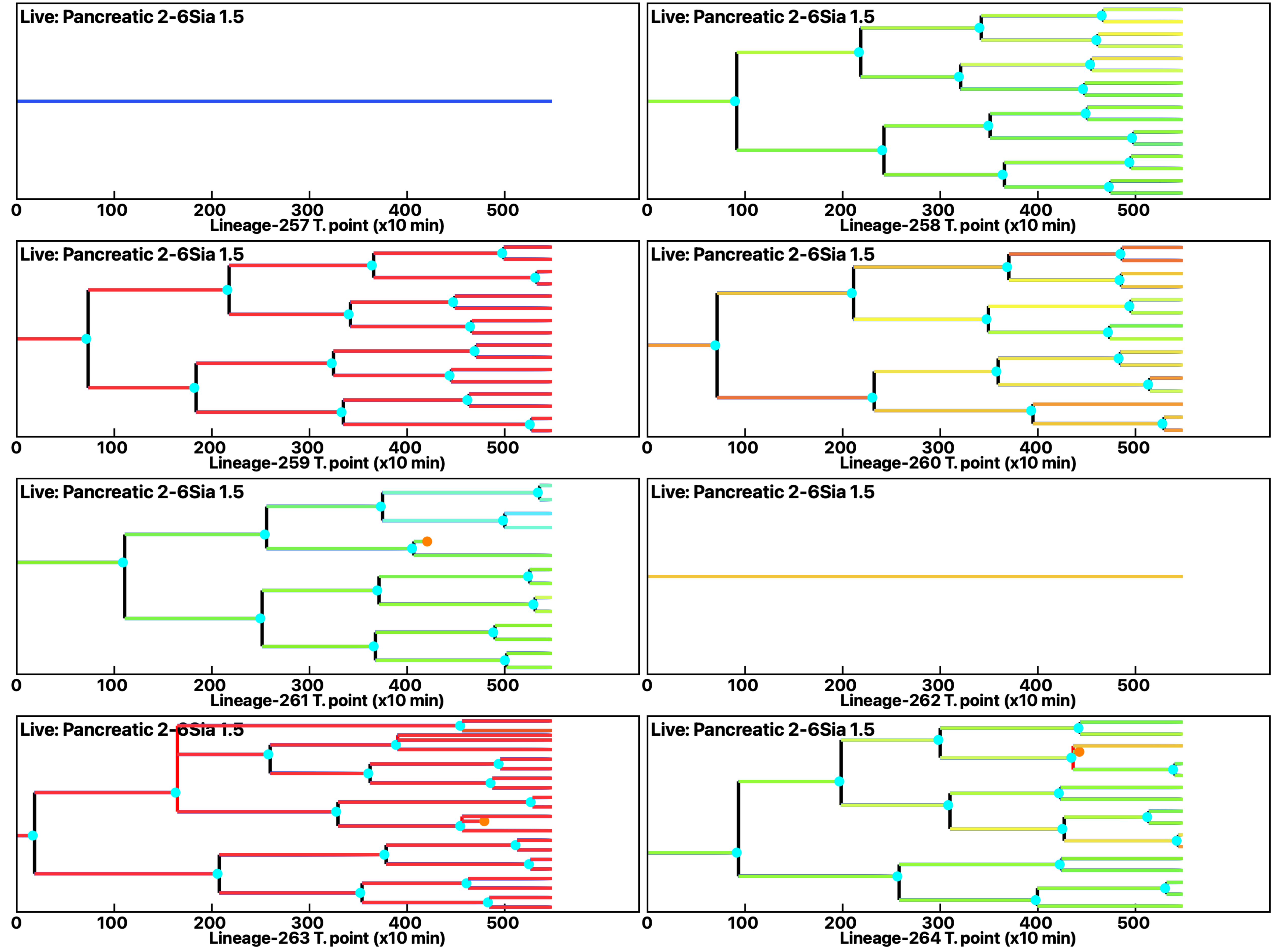

Analysis: Simulation, Treat.: Pancreatic 2-6Sia 1.5

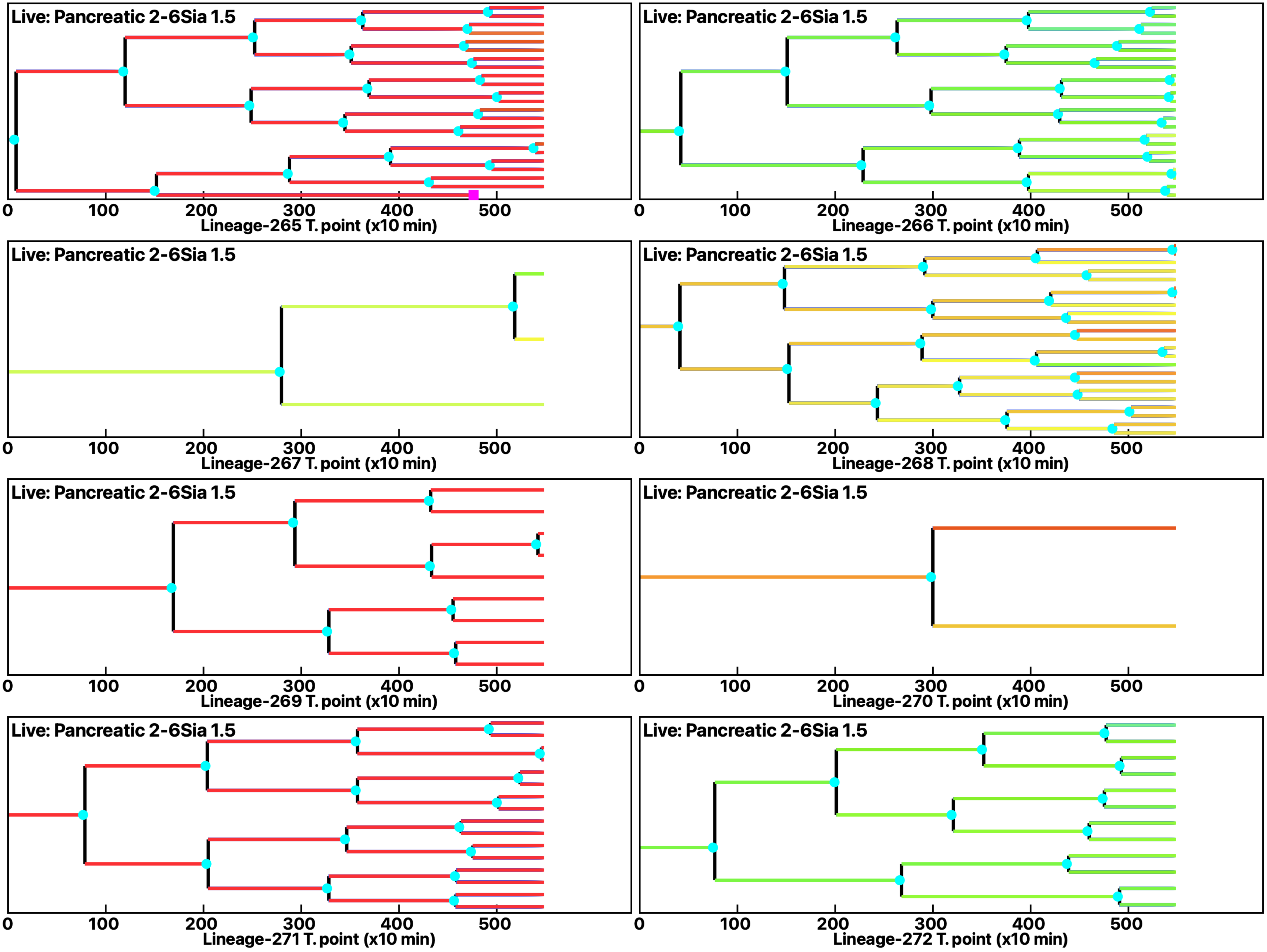

Analysis: Simulation, Treat.: Pancreatic 2-6Sia 1.5

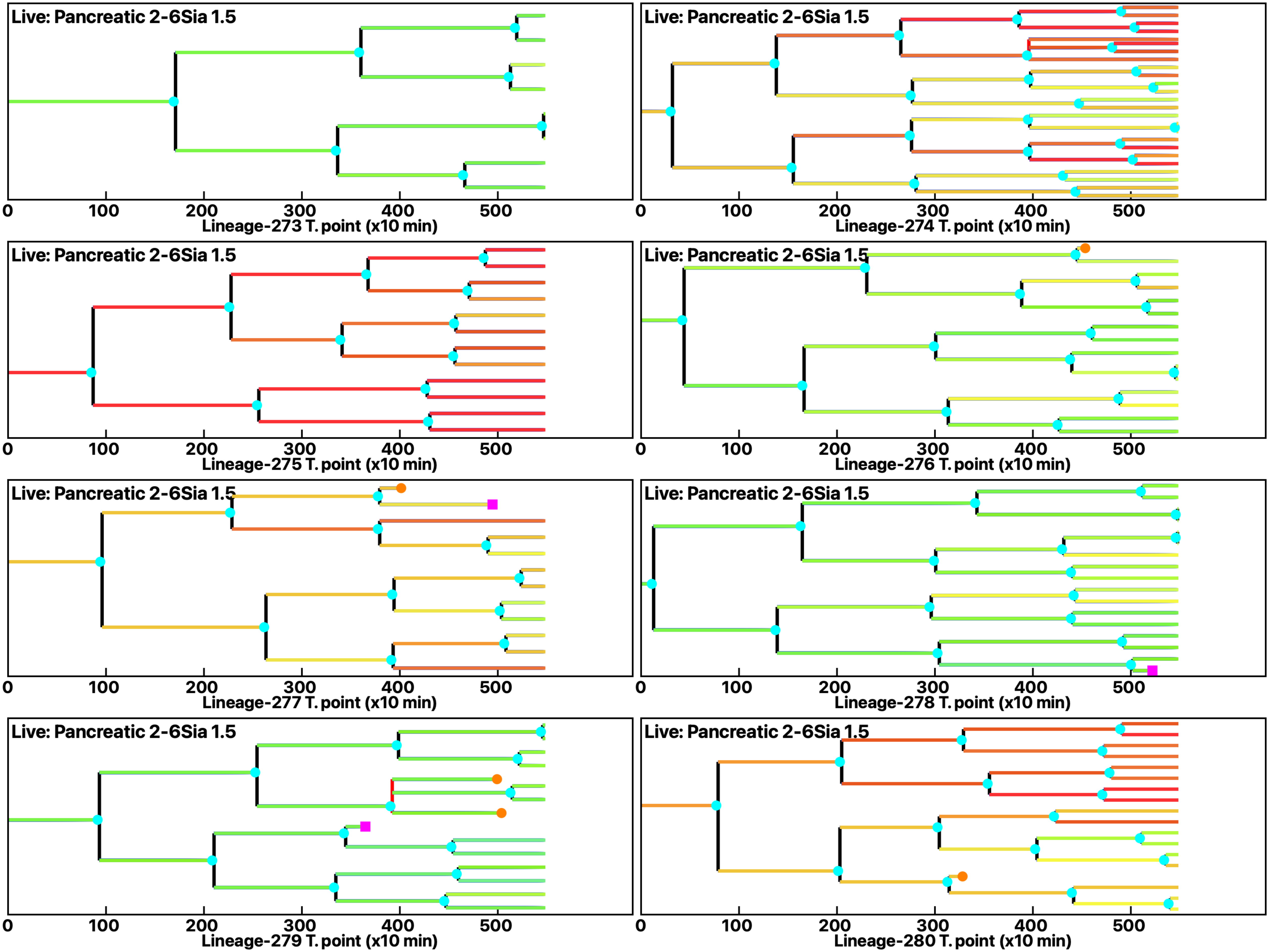

Analysis: Simulation, Treat.: Pancreatic 2-6Sia 1.5

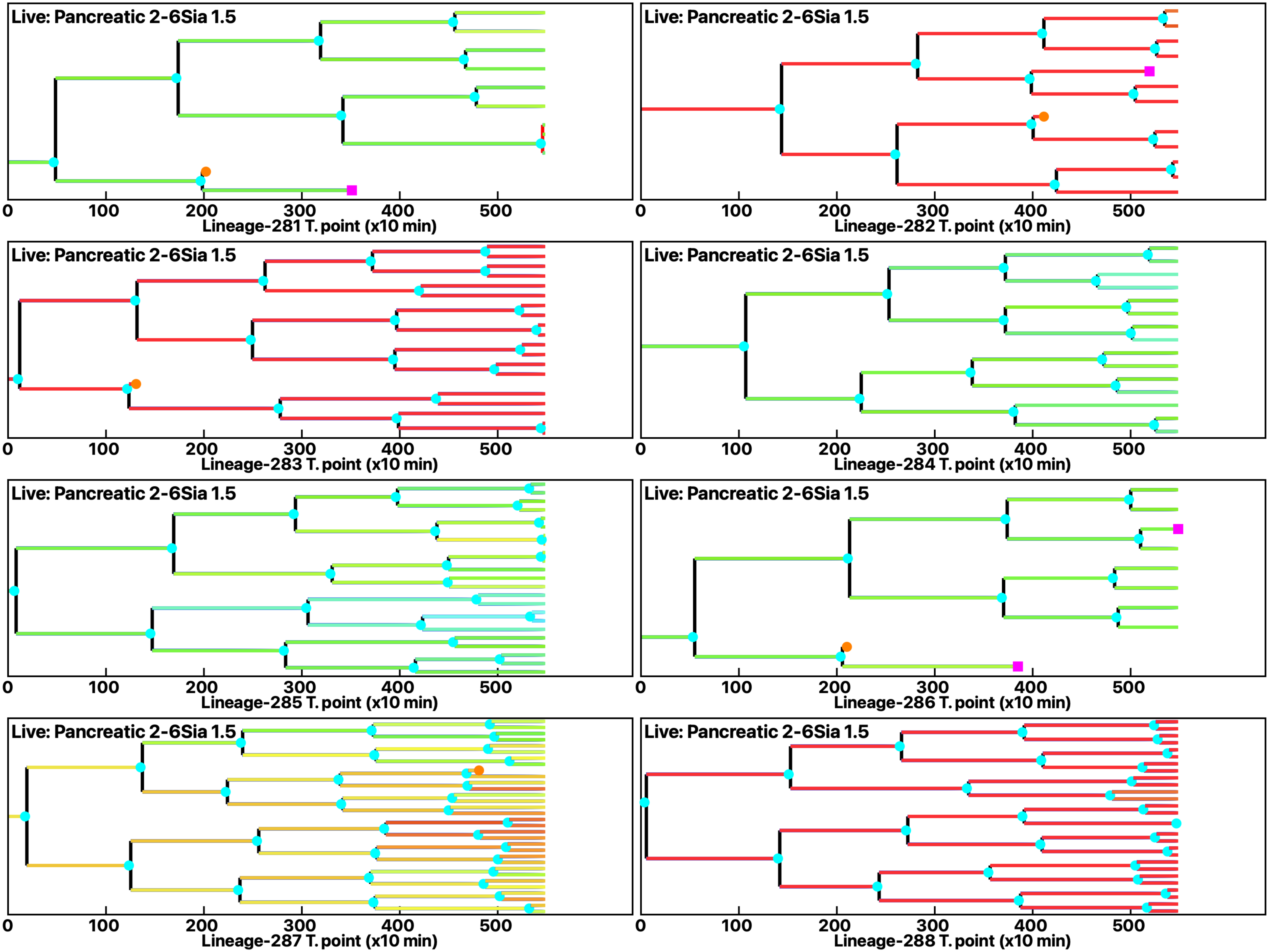

**Analysis: Simulation, Treat.: Pancreatic 2-6Sia 1.5**

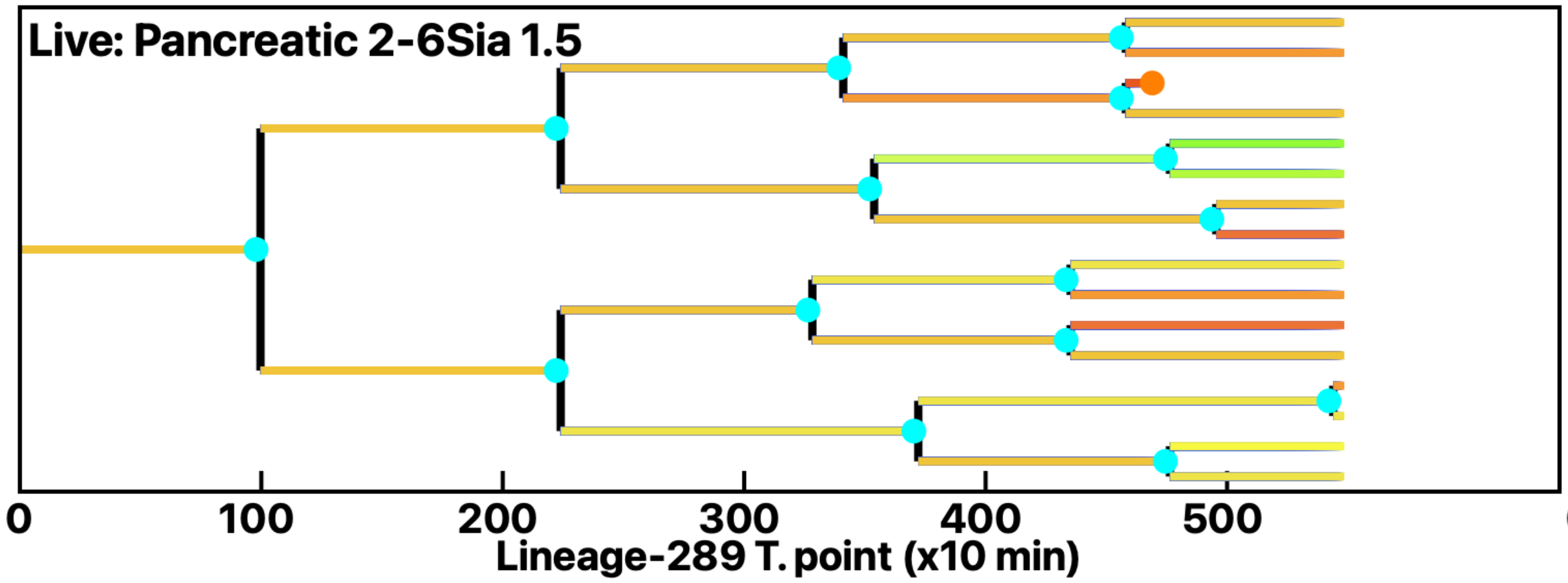


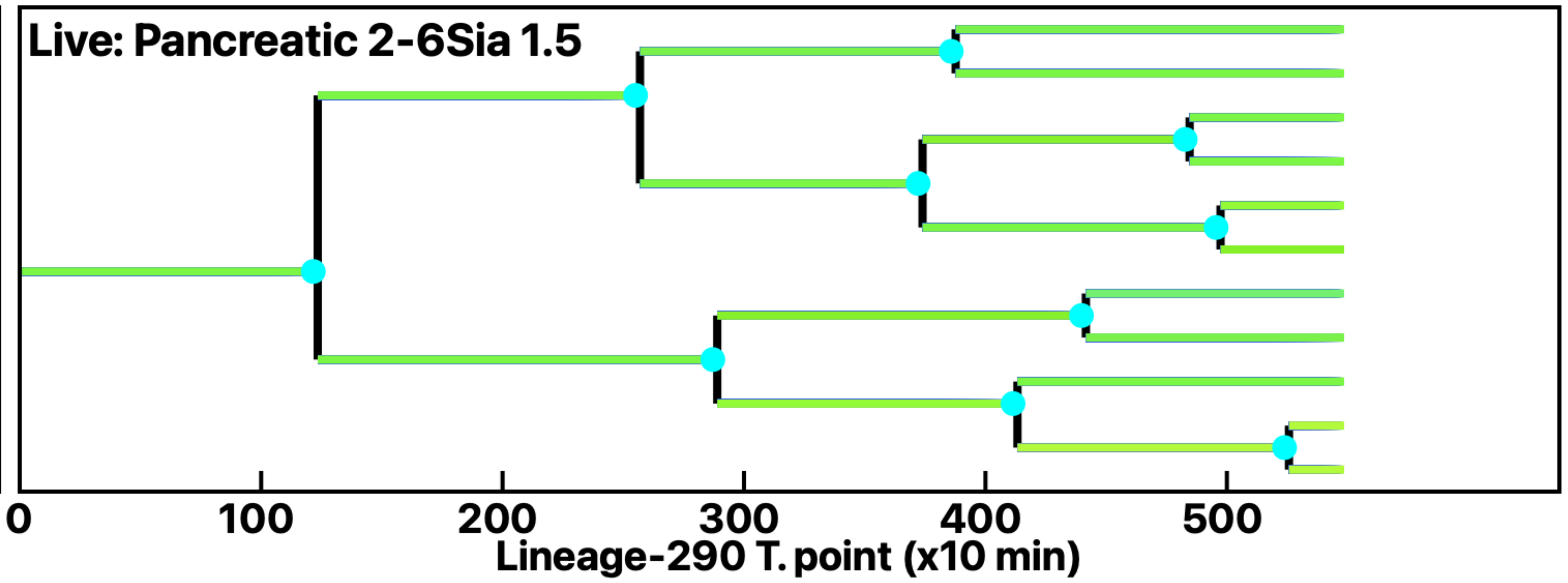


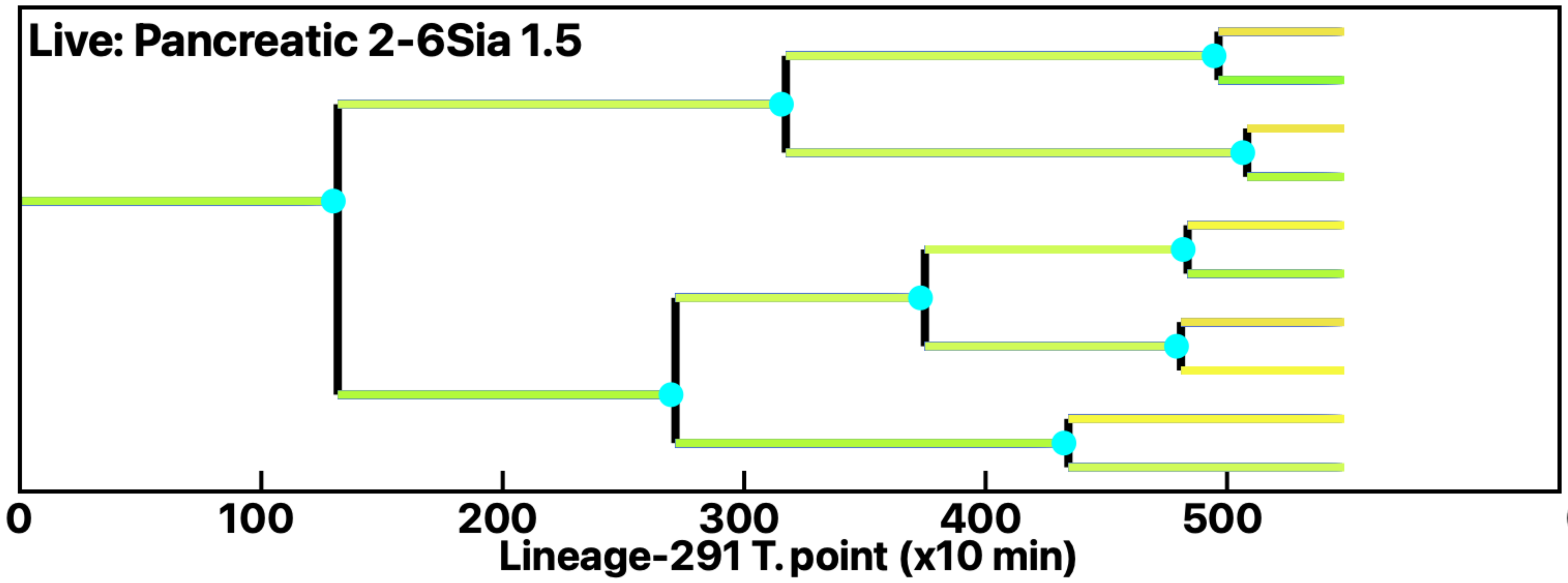


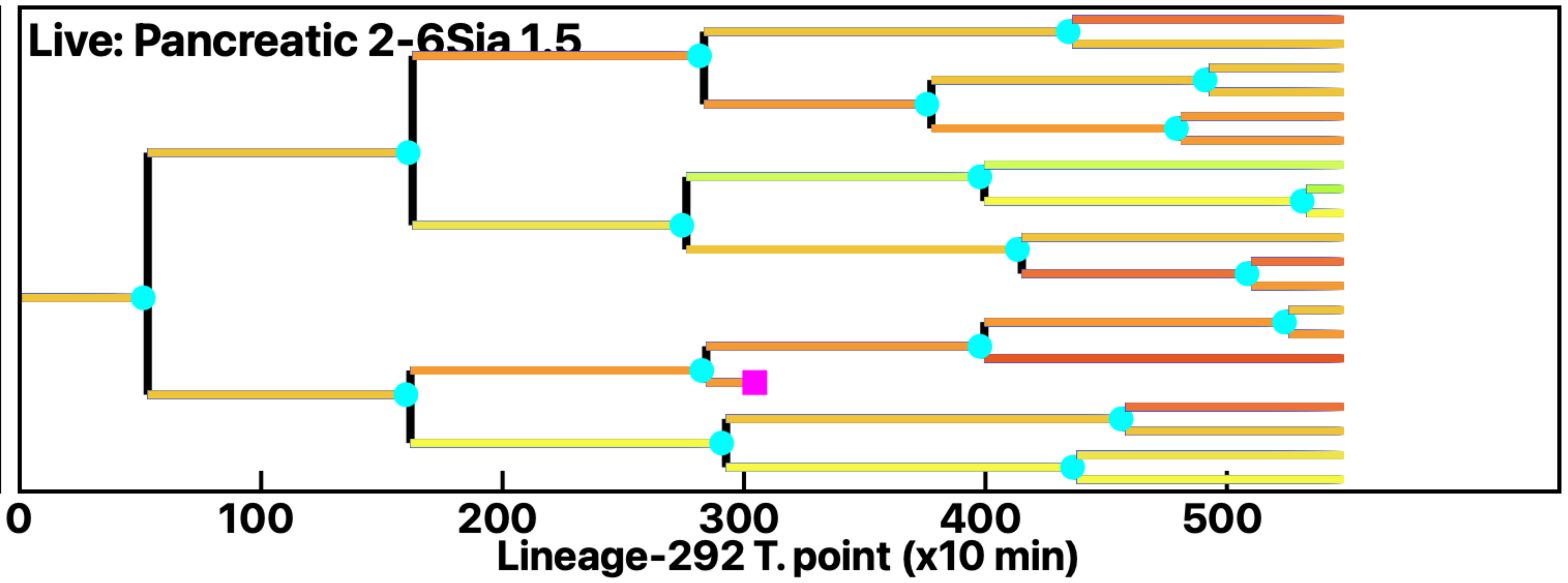


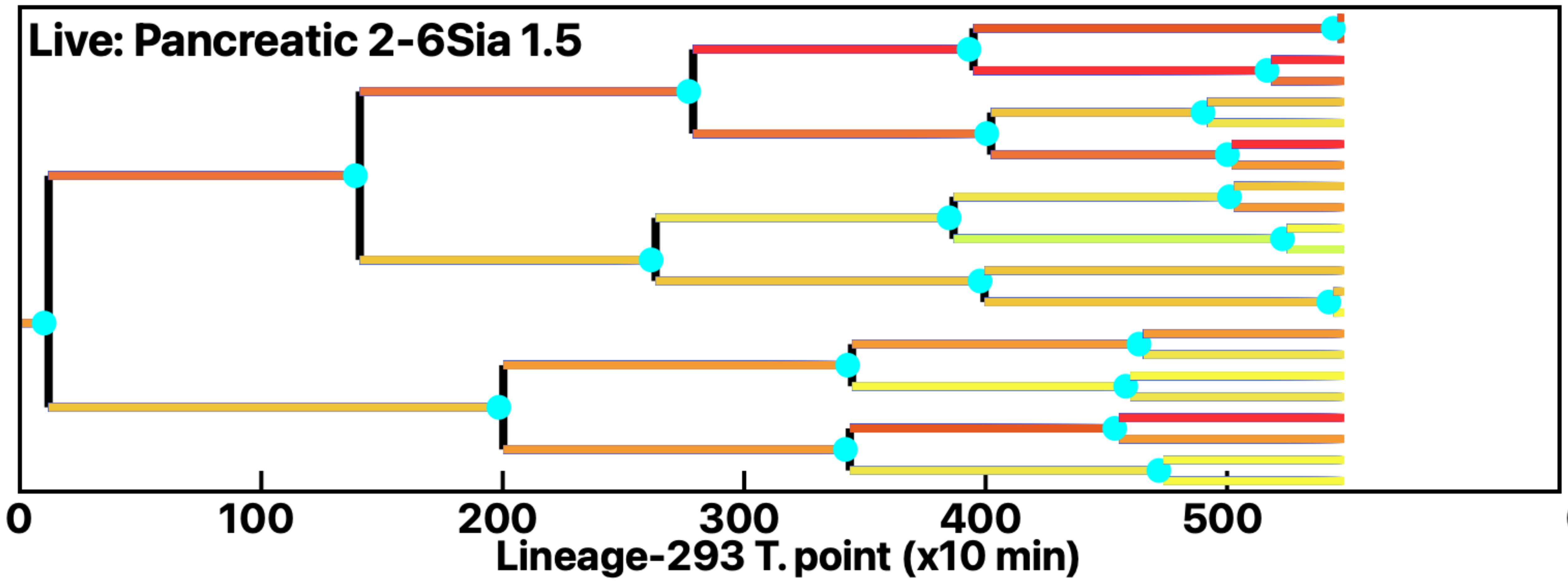


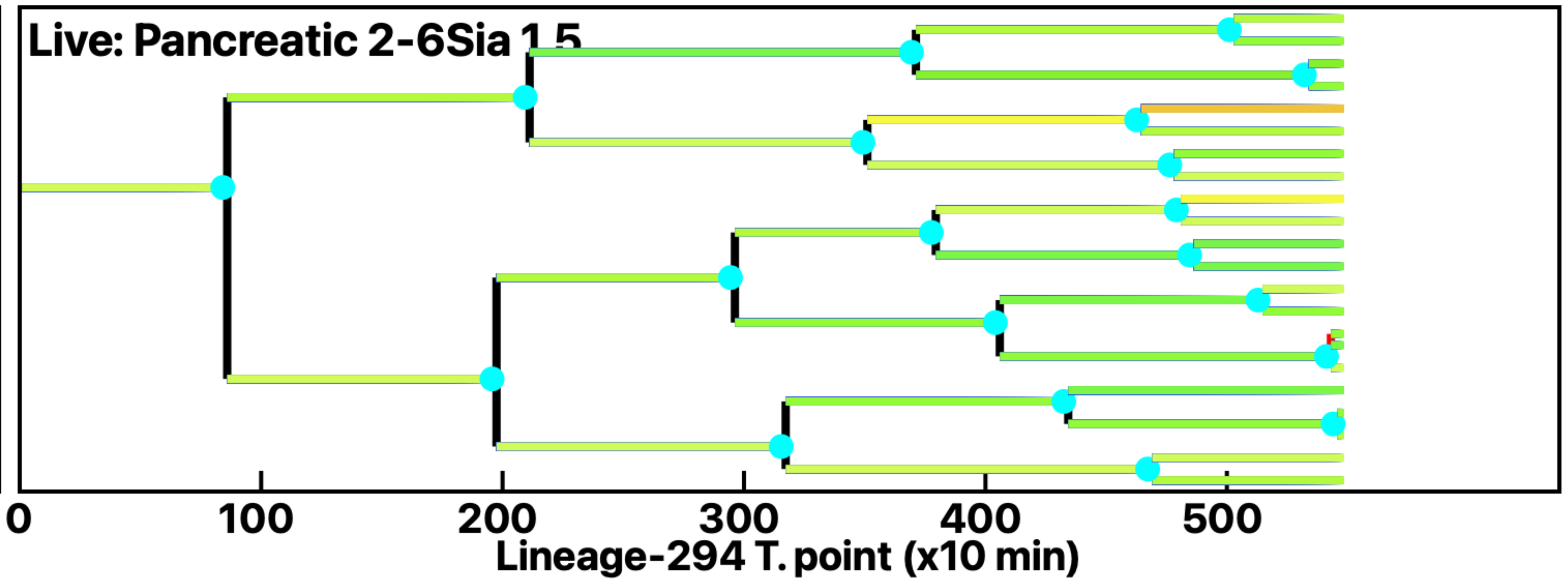


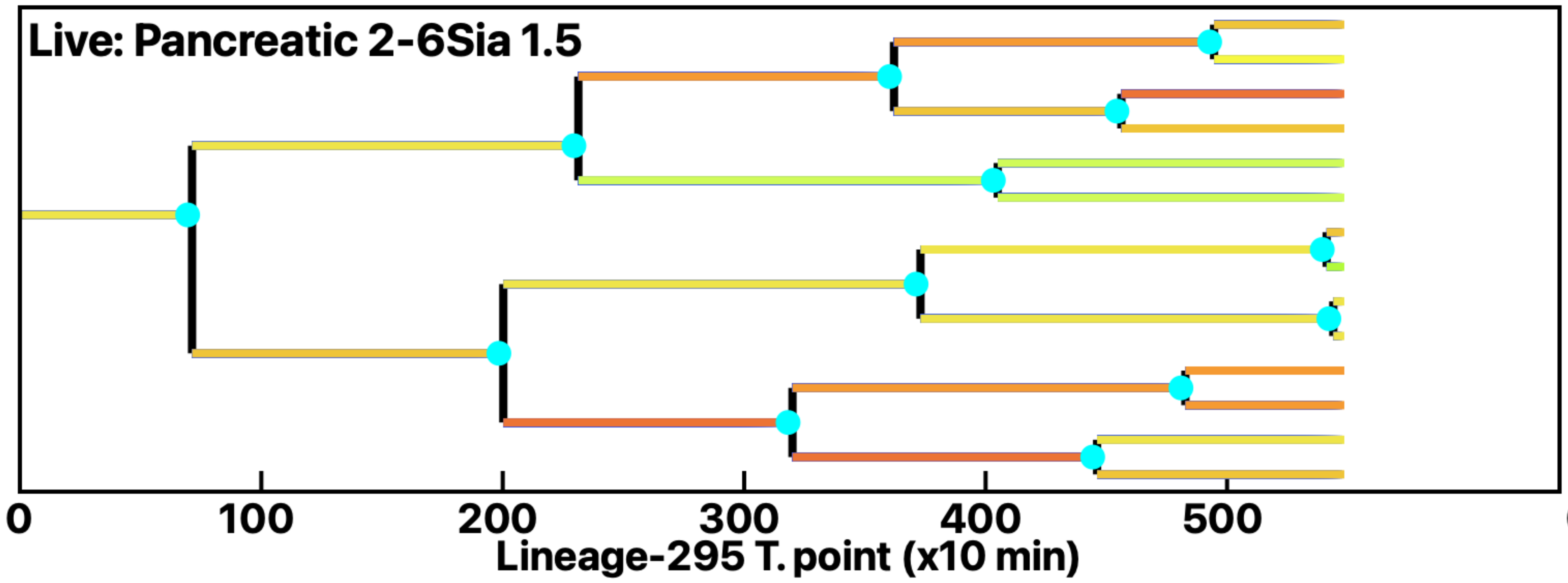


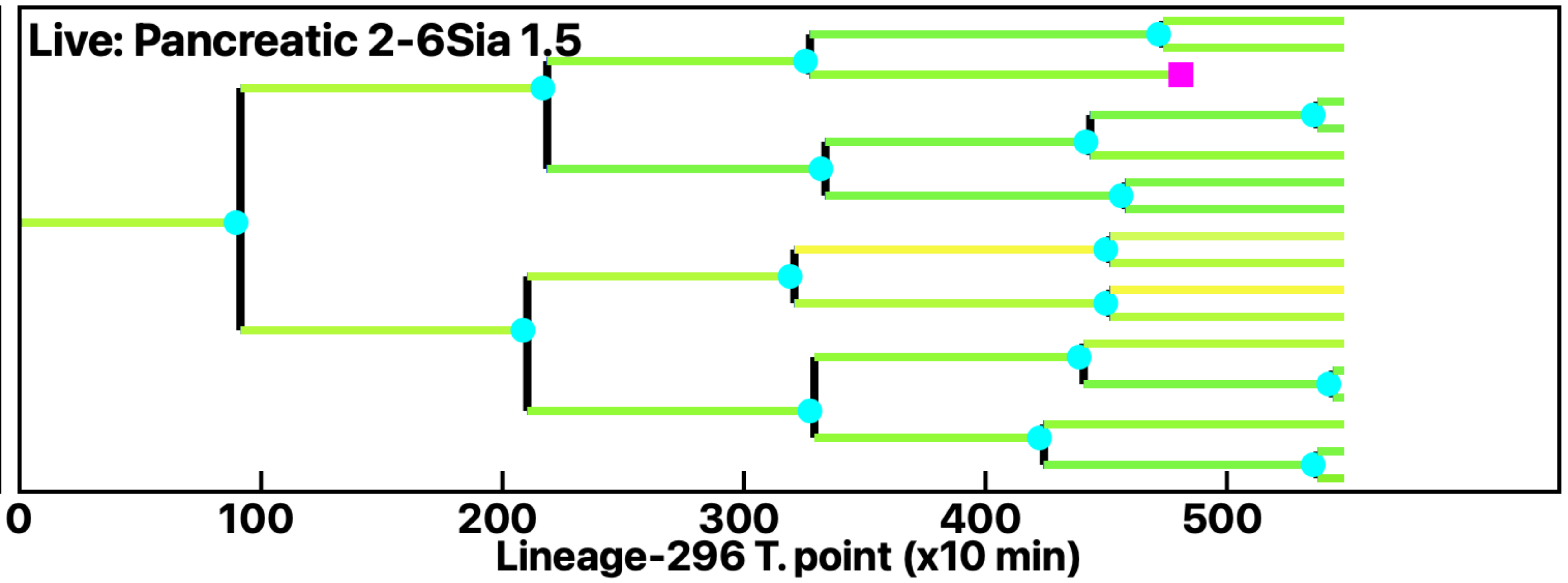

Analysis: Simulation, Treat.: Pancreatic 2-6Sia 1.5

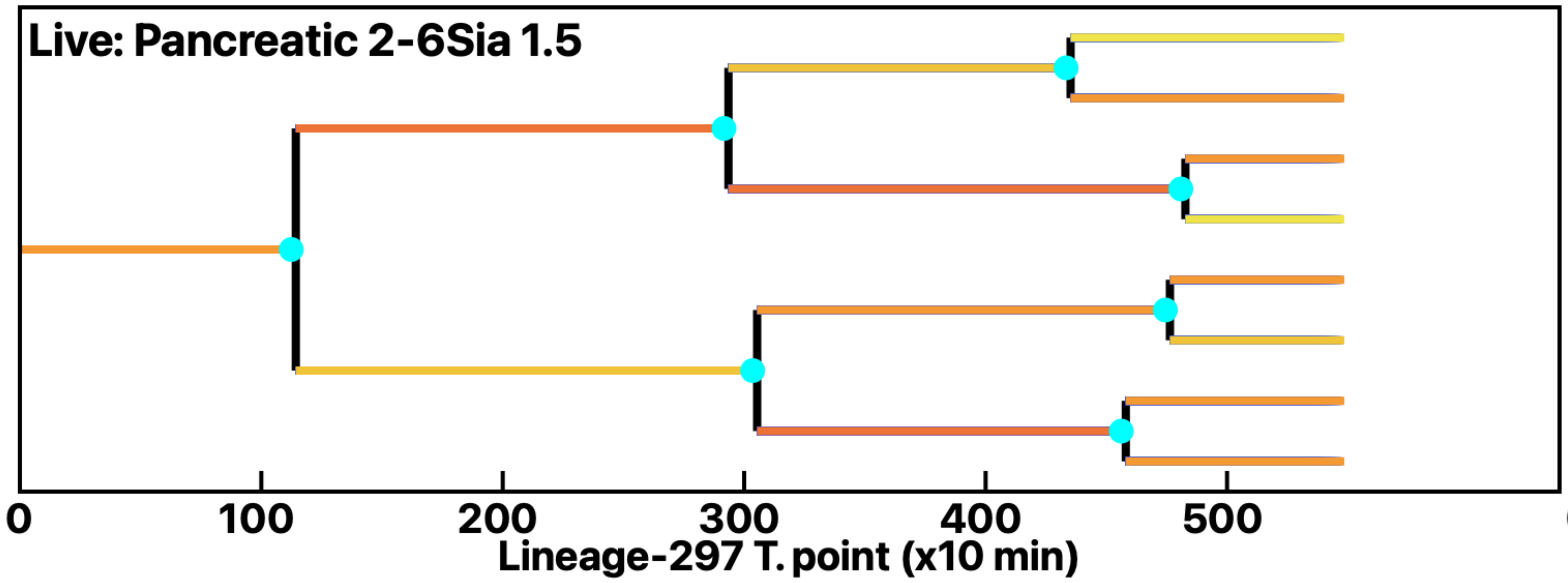


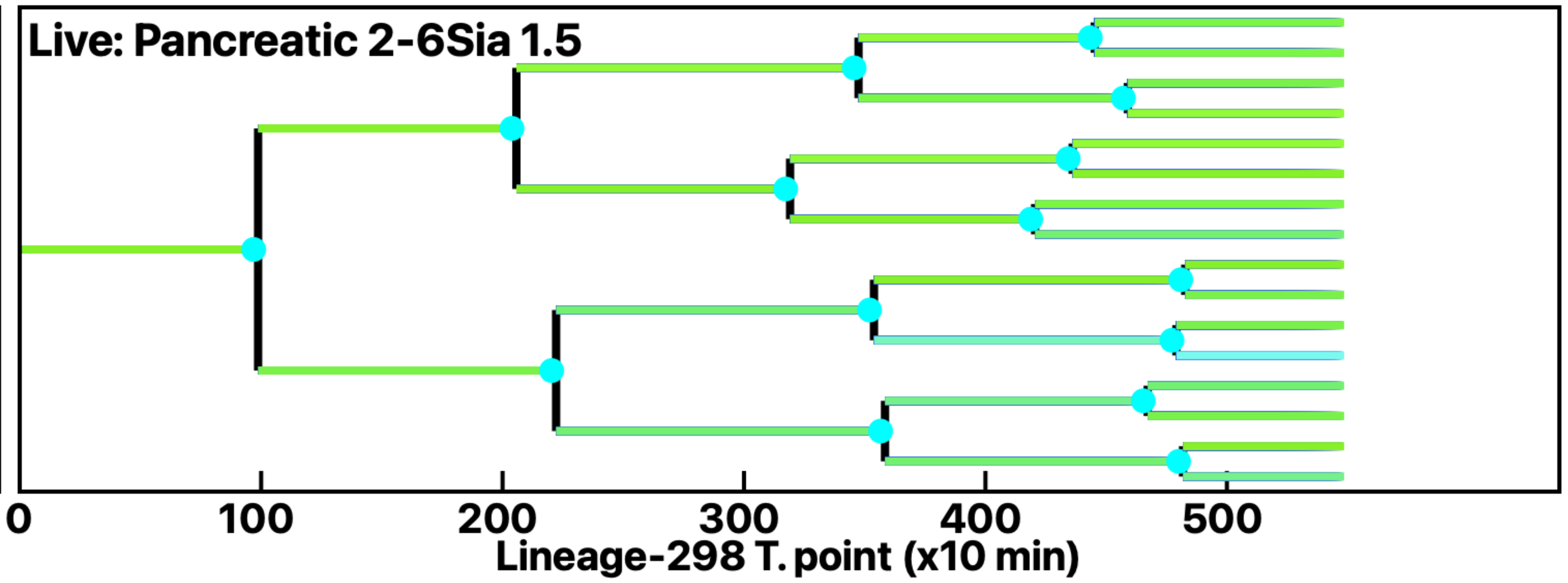


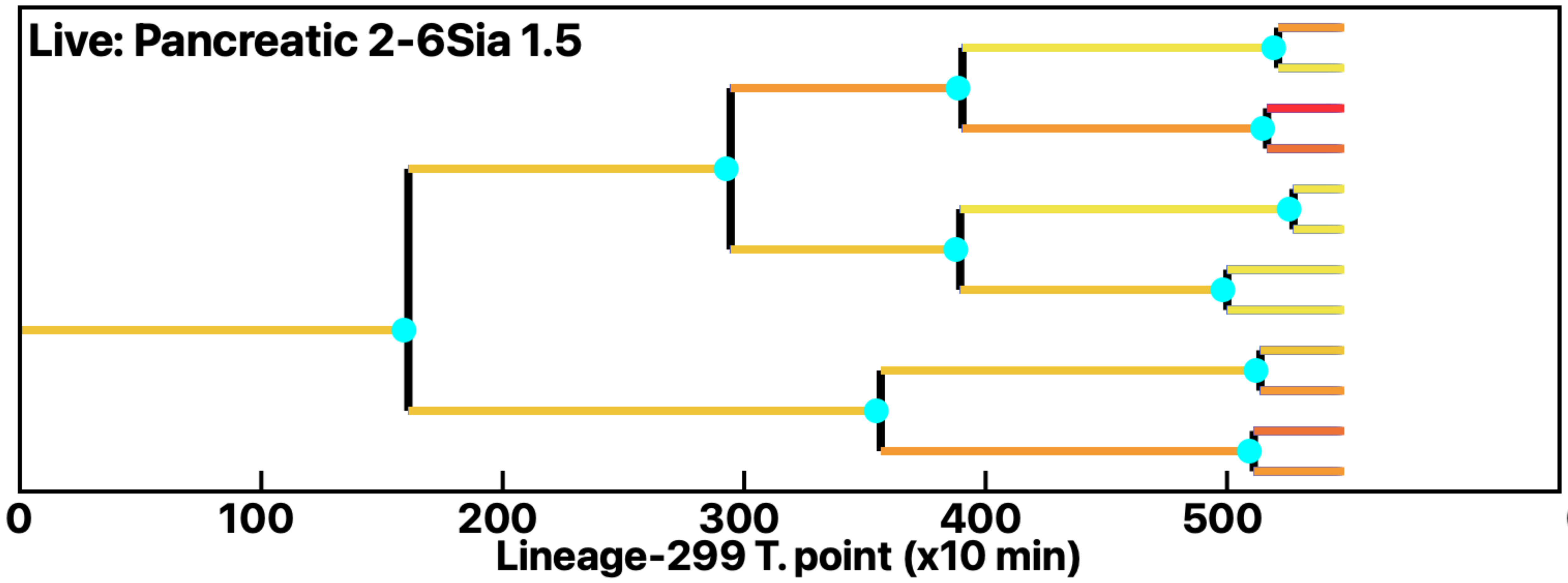


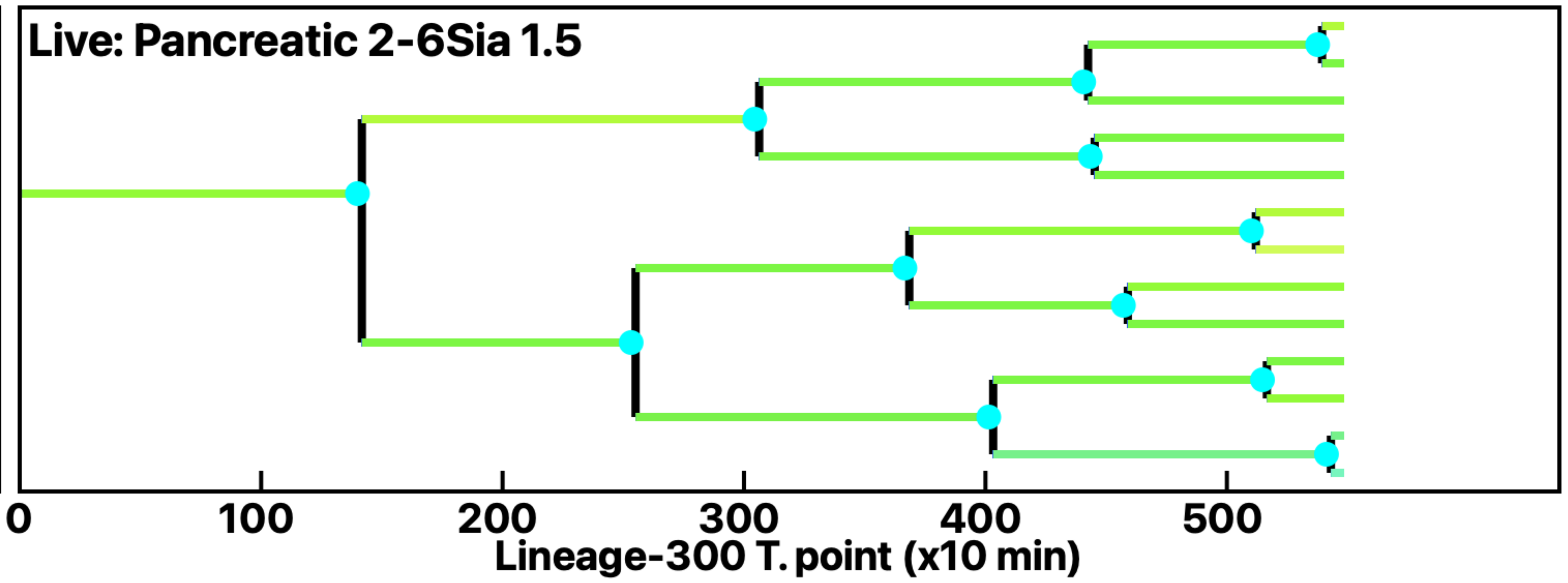


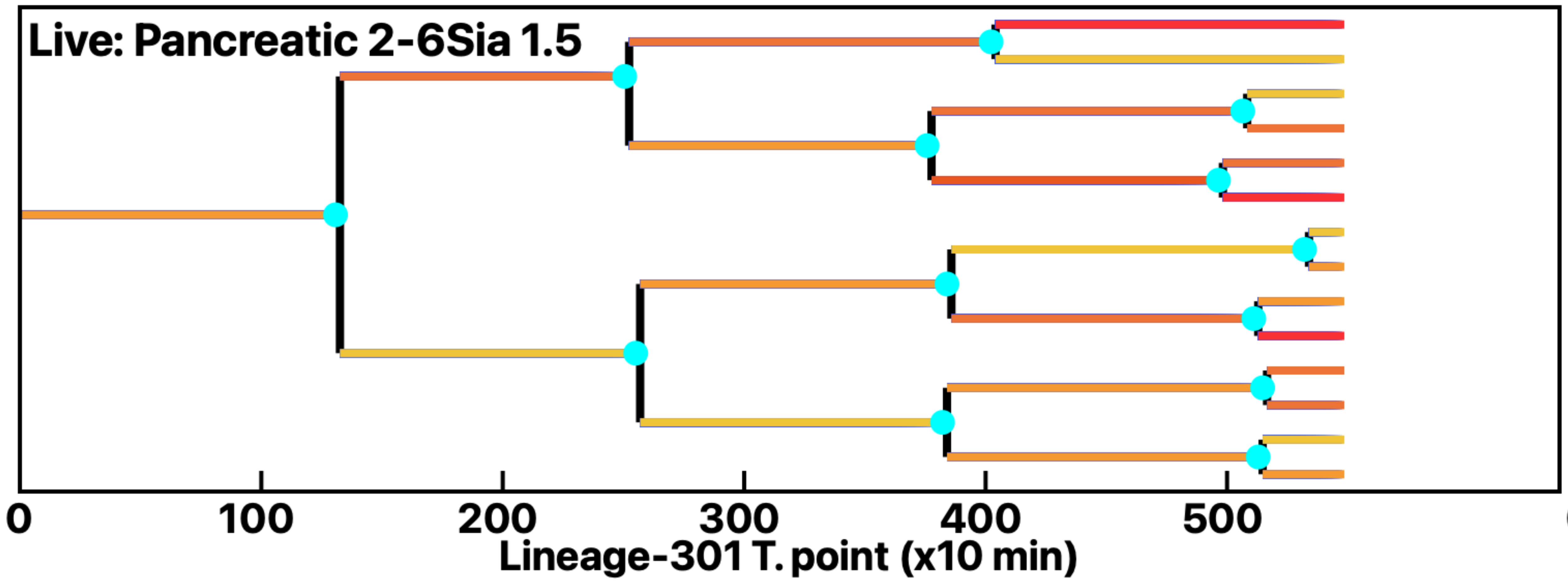


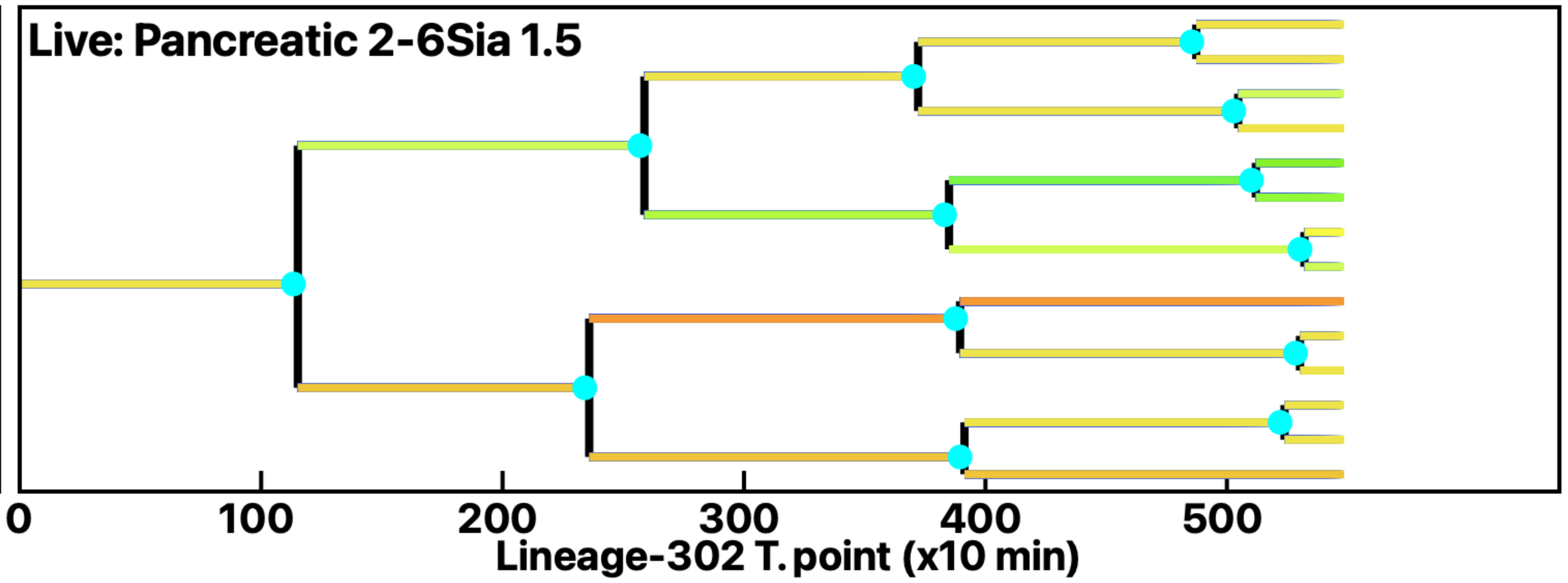


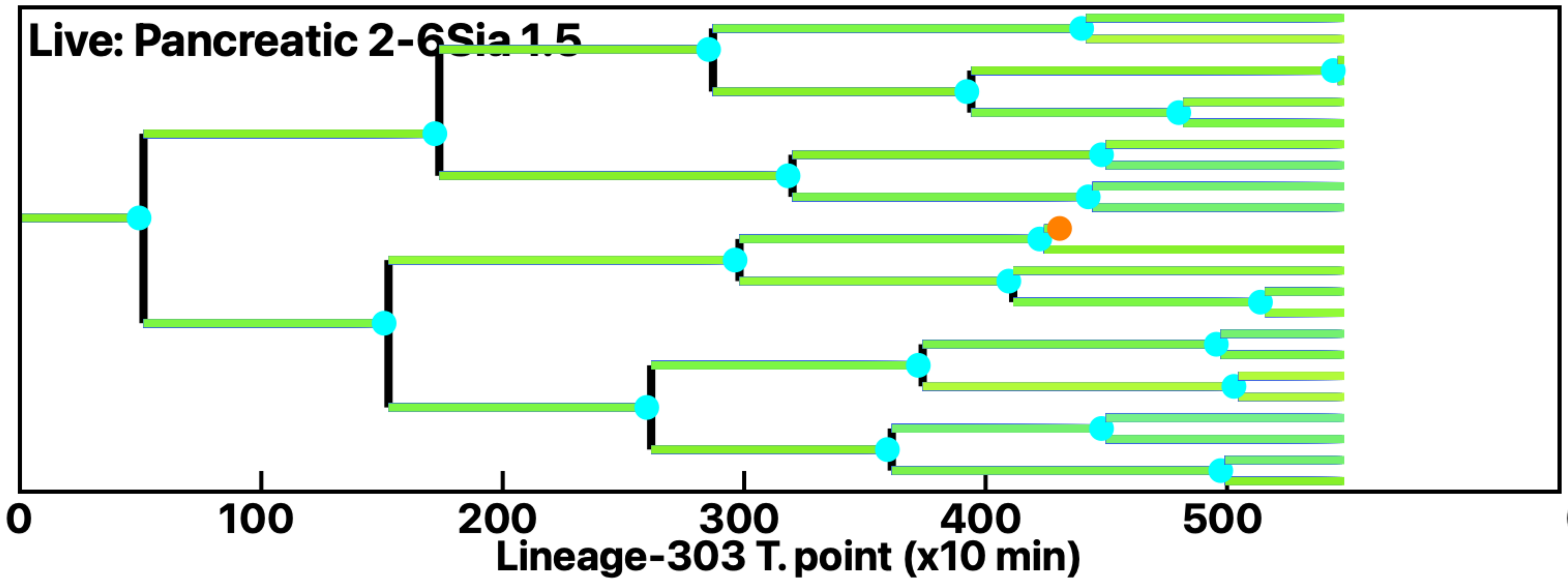


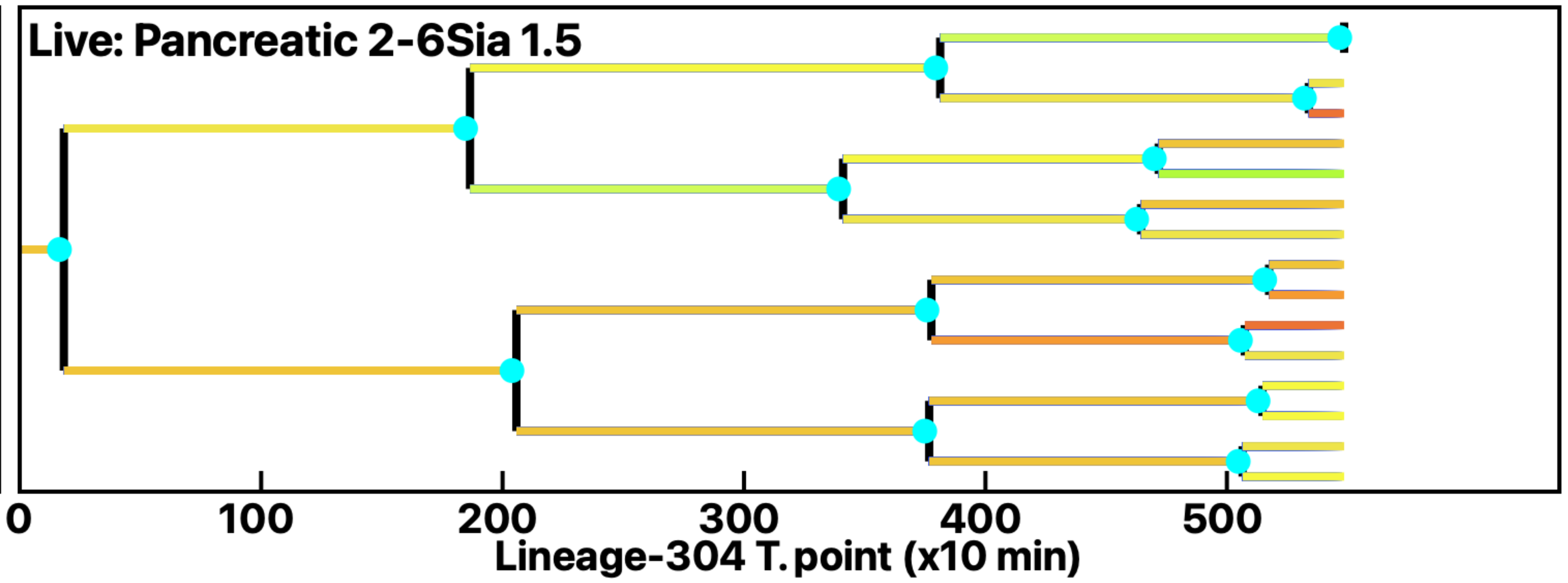

Analysis: Simulation, Treat.: Pancreatic 2-6Sia 1.5

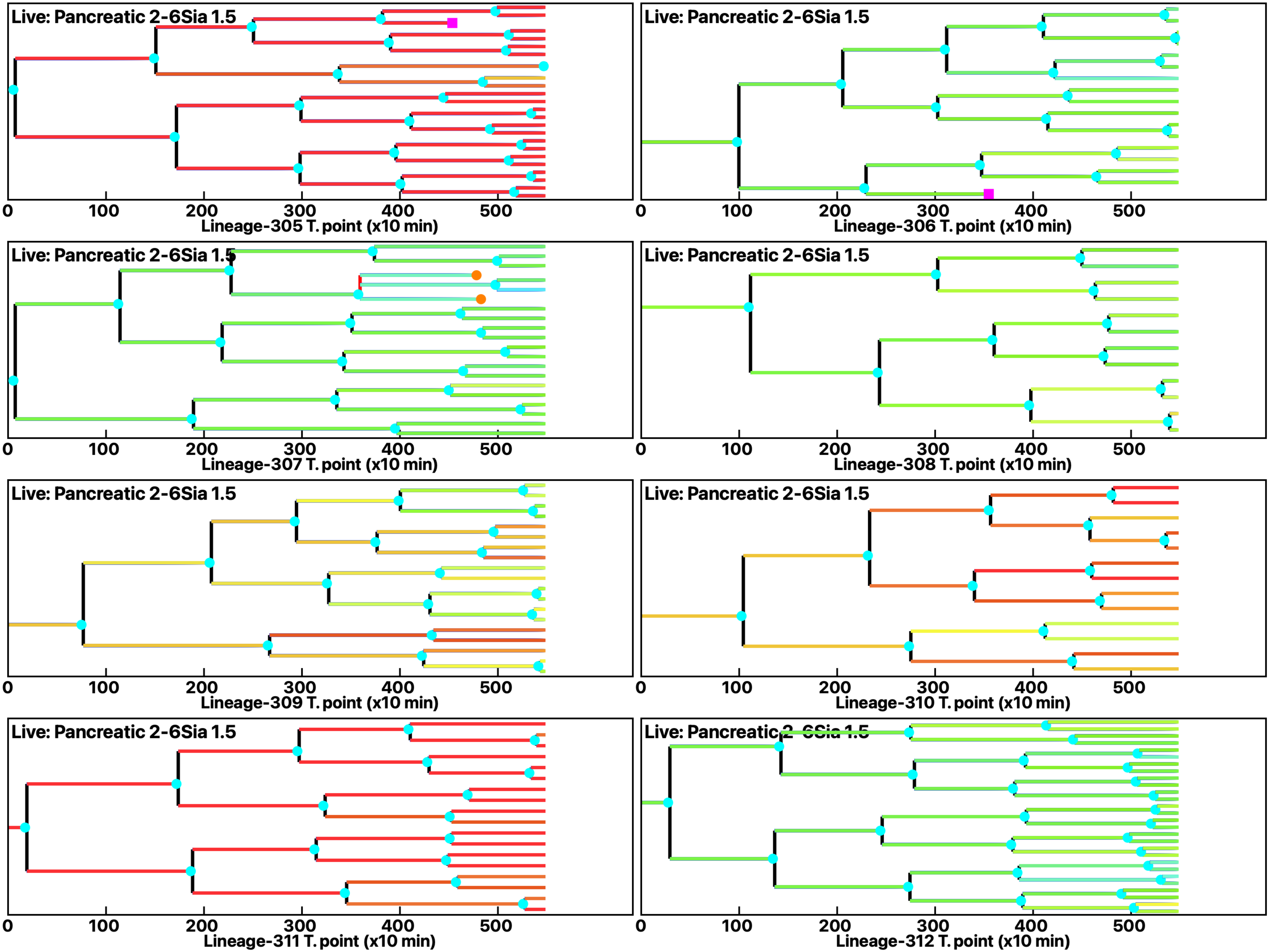

Analysis: Simulation, Treat.: Pancreatic 2-6Sia 1.5

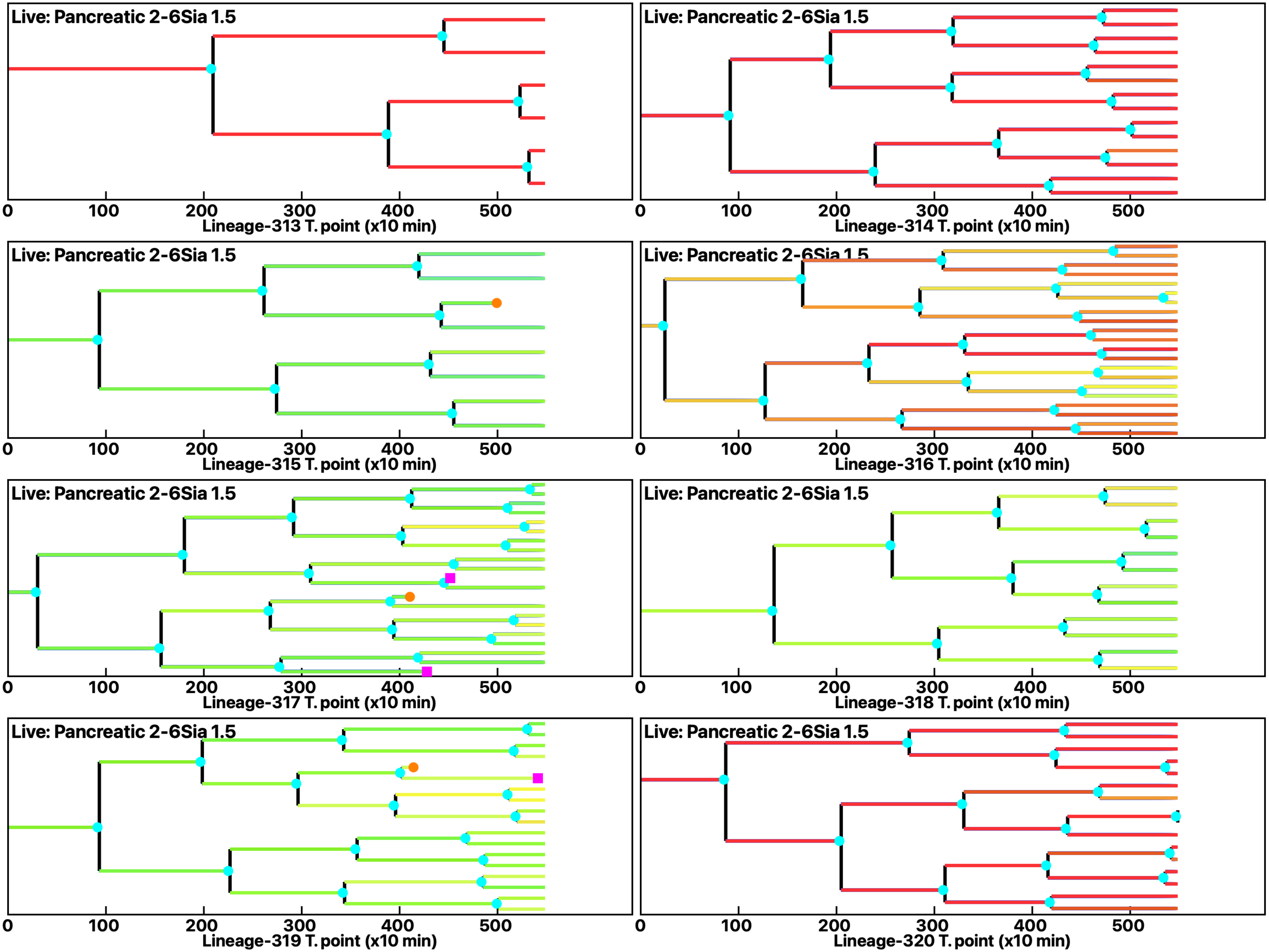

Analysis: Simulation, Treat.: Pancreatic 2-6Sia 1.5

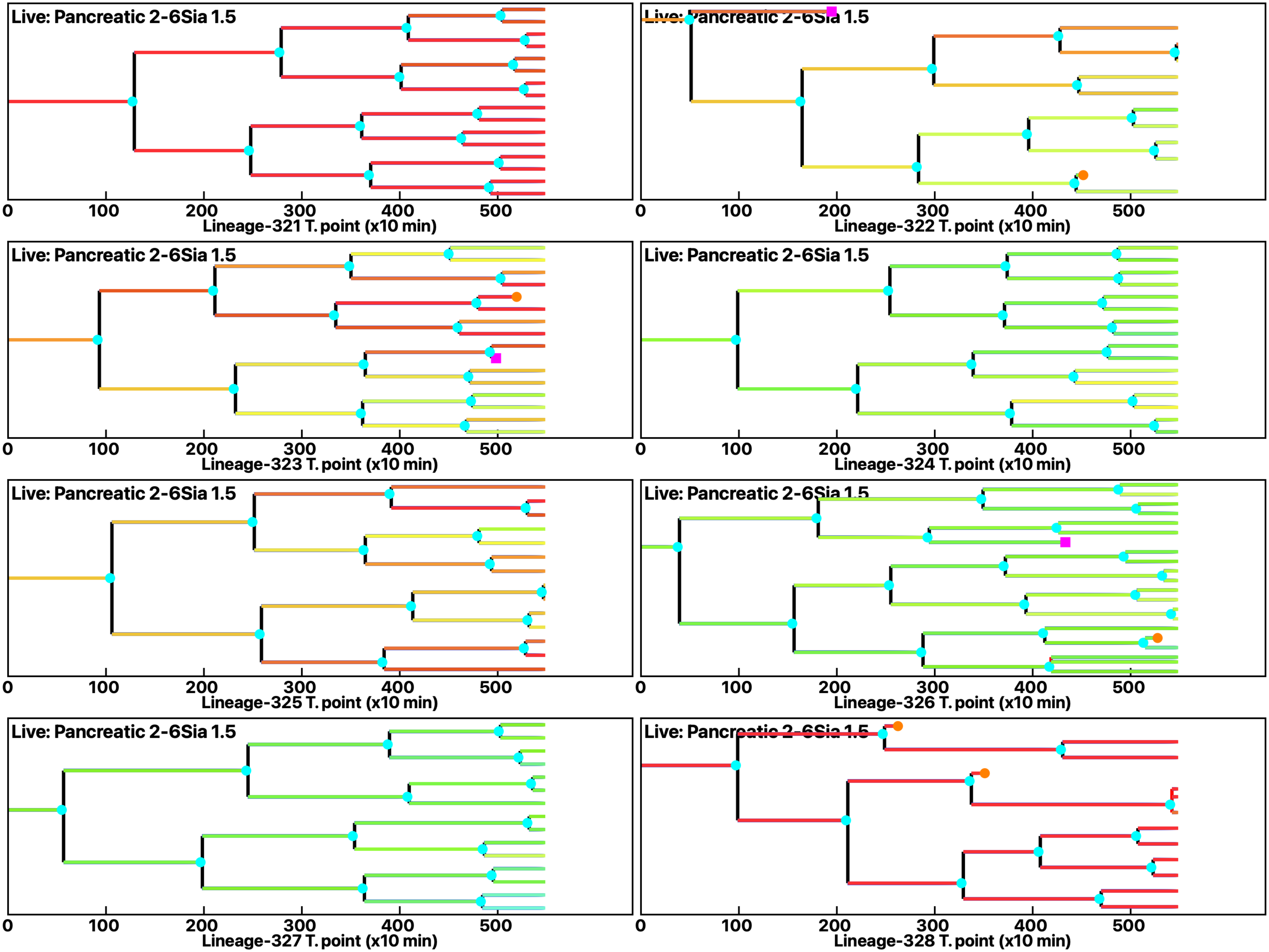

**Analysis: Simulation, Treat.: Pancreatic 2-6Sia 1.5**

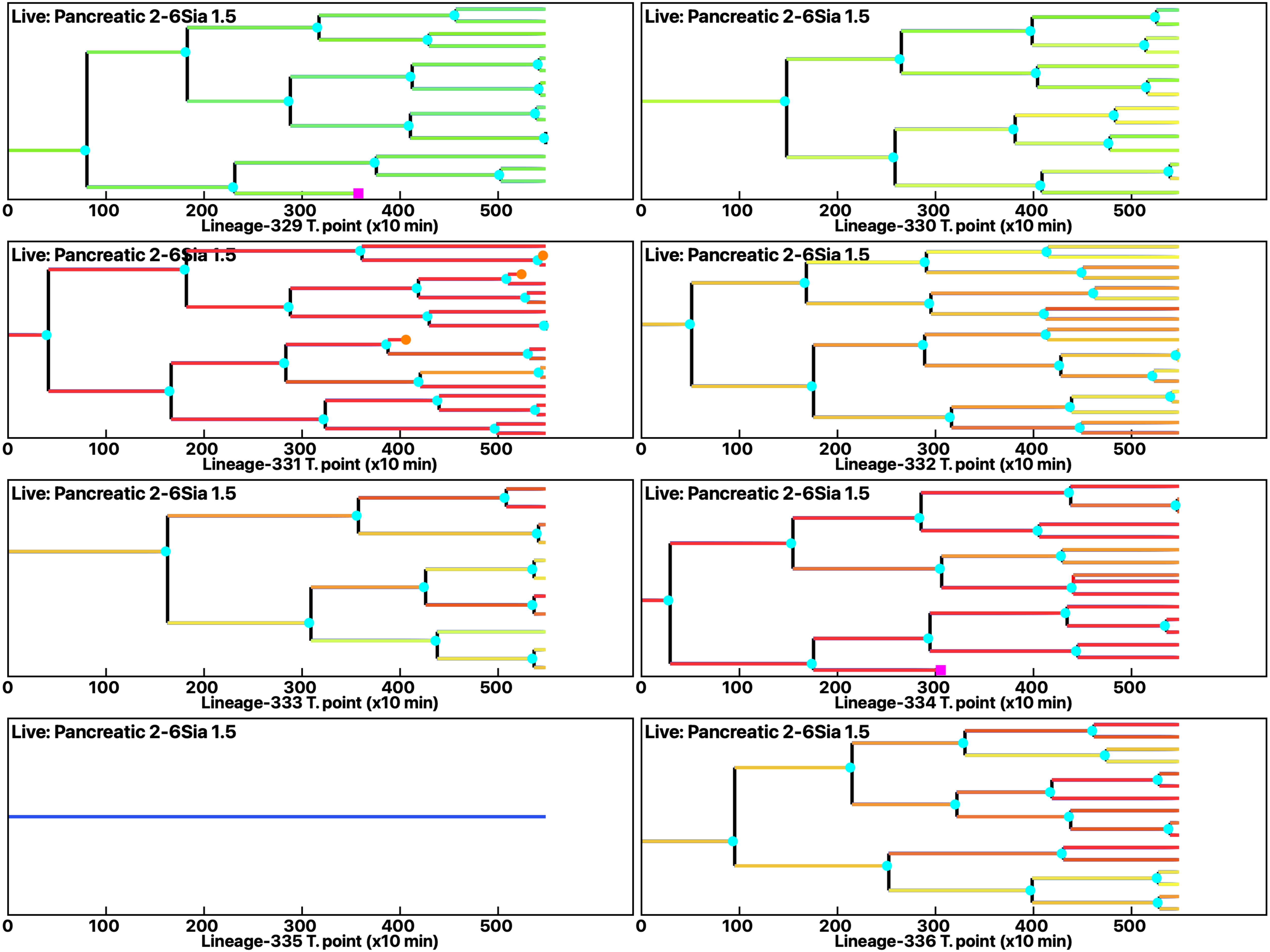

Analysis: Simulation, Treat.: Pancreatic 2-6Sia 1.5

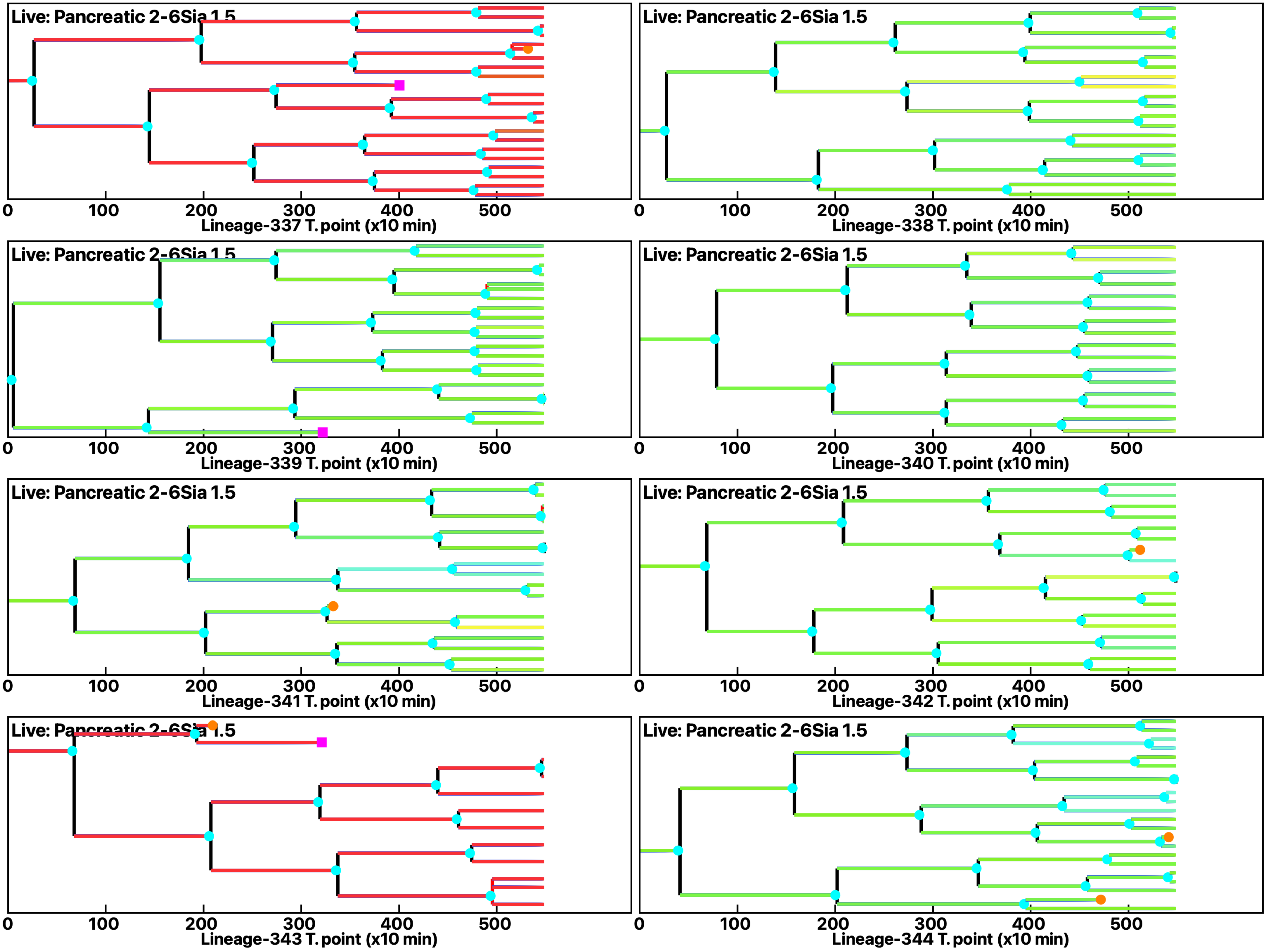

Analysis: Simulation, Treat.: Pancreatic 2-6Sia 1.5

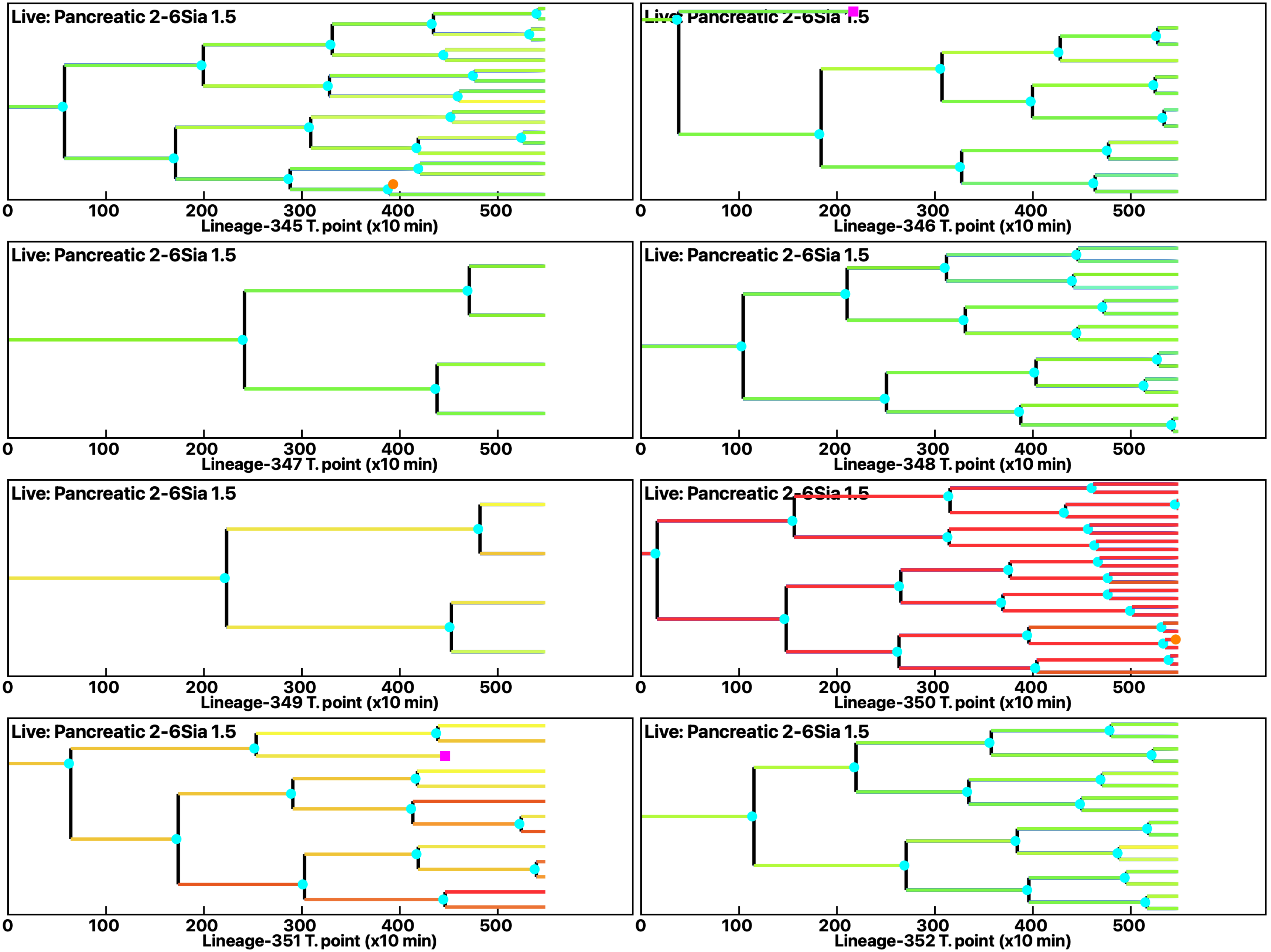

Analysis: Simulation, Treat.: Pancreatic 2-6Sia 1.5

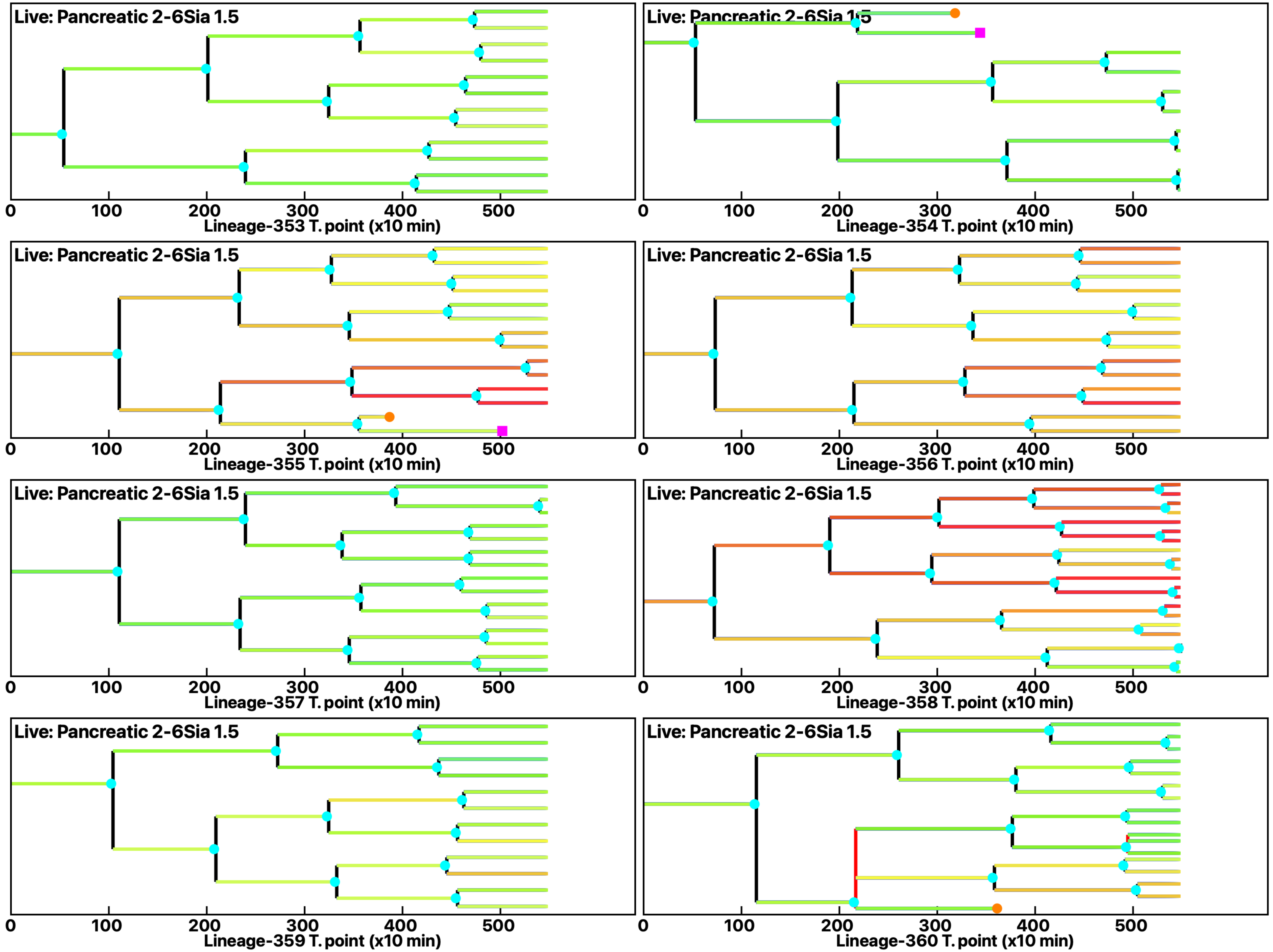

Analysis: Simulation, Treat.: Pancreatic 2-6Sia 1.5

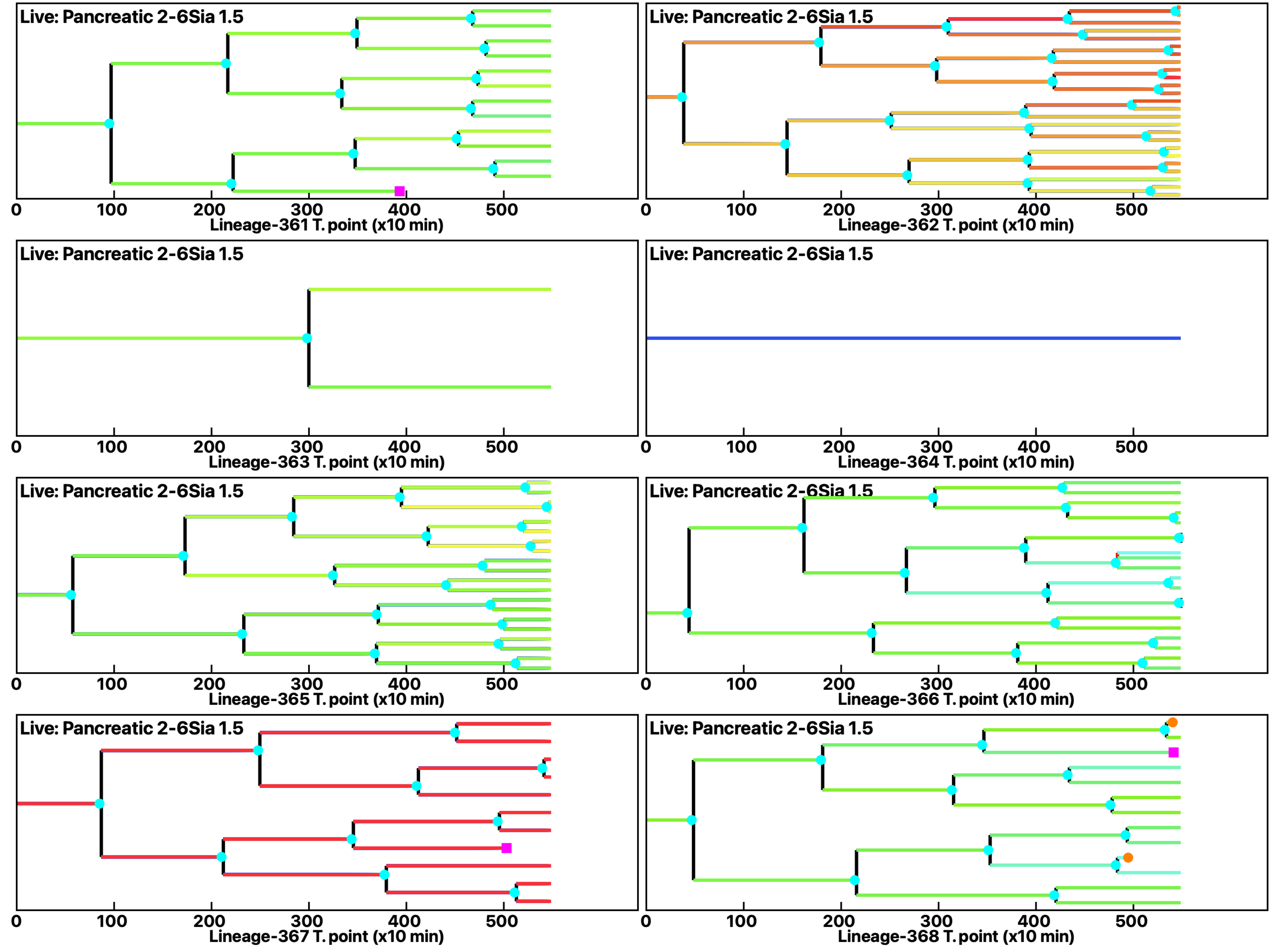

Analysis: Simulation, Treat.: Pancreatic 2-6Sia 1.5

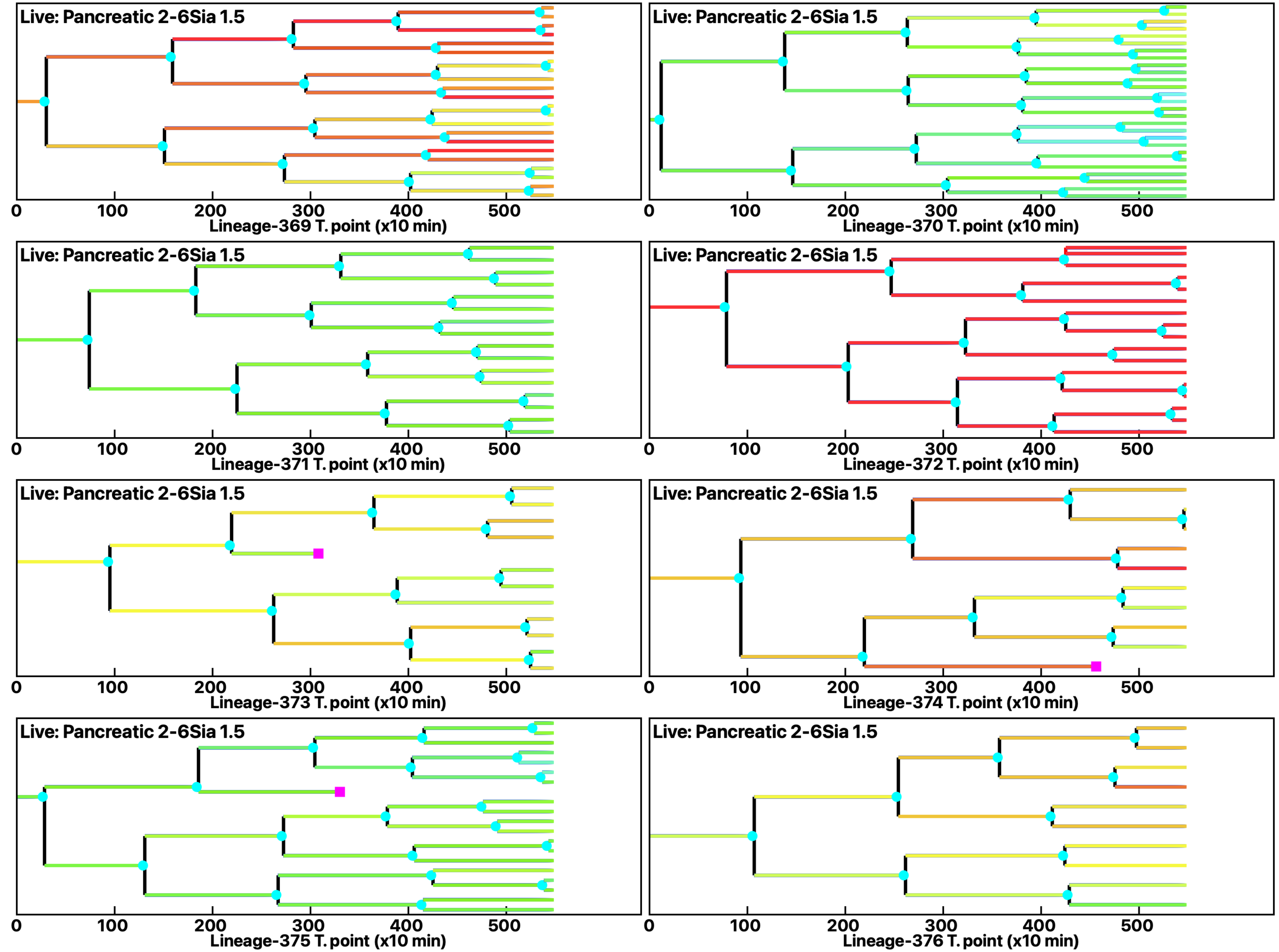

Analysis: Simulation, Treat.: Pancreatic 2-6Sia 1.5

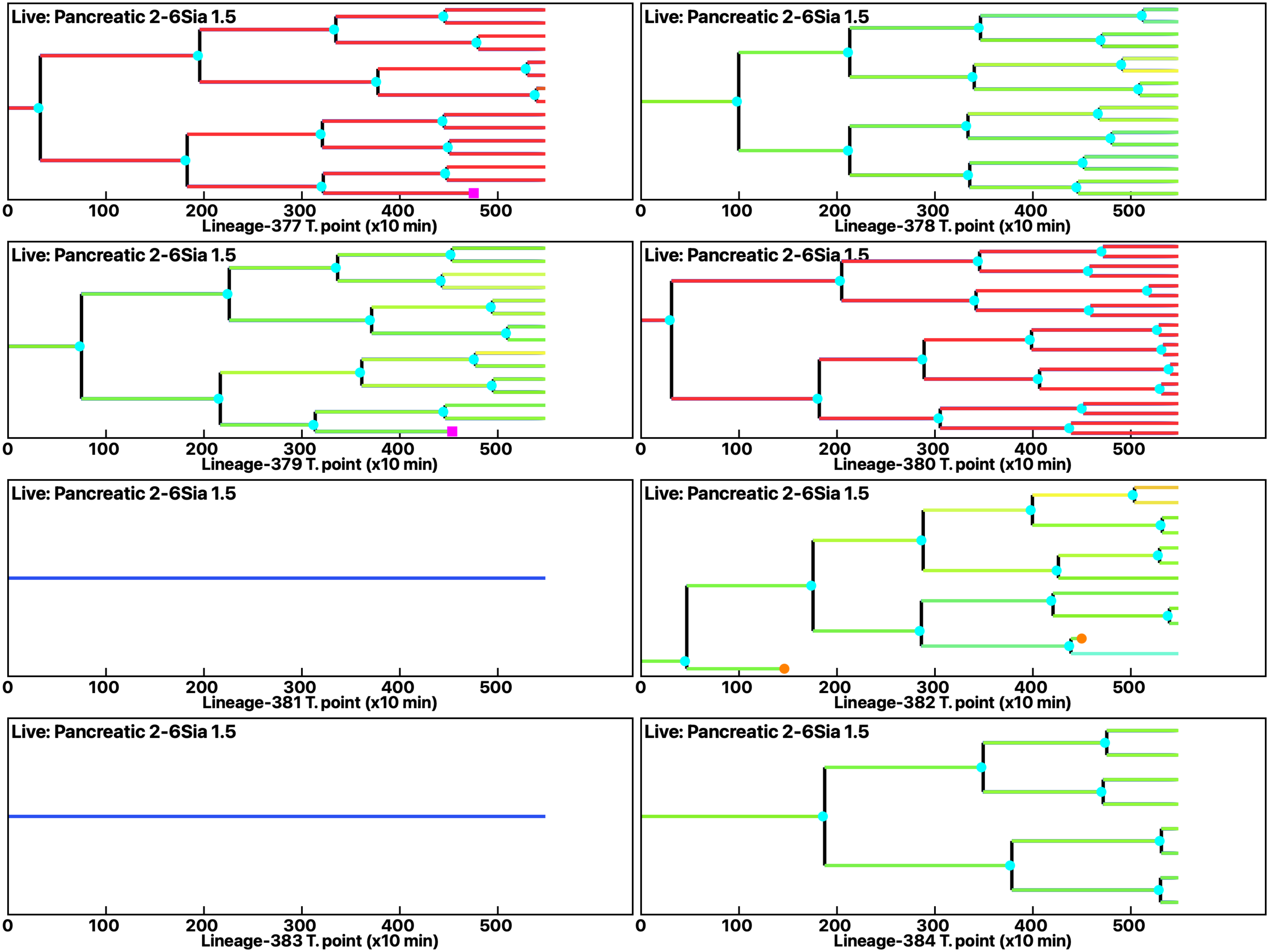

Analysis: Simulation, Treat.: Pancreatic 2-6Sia 1.5

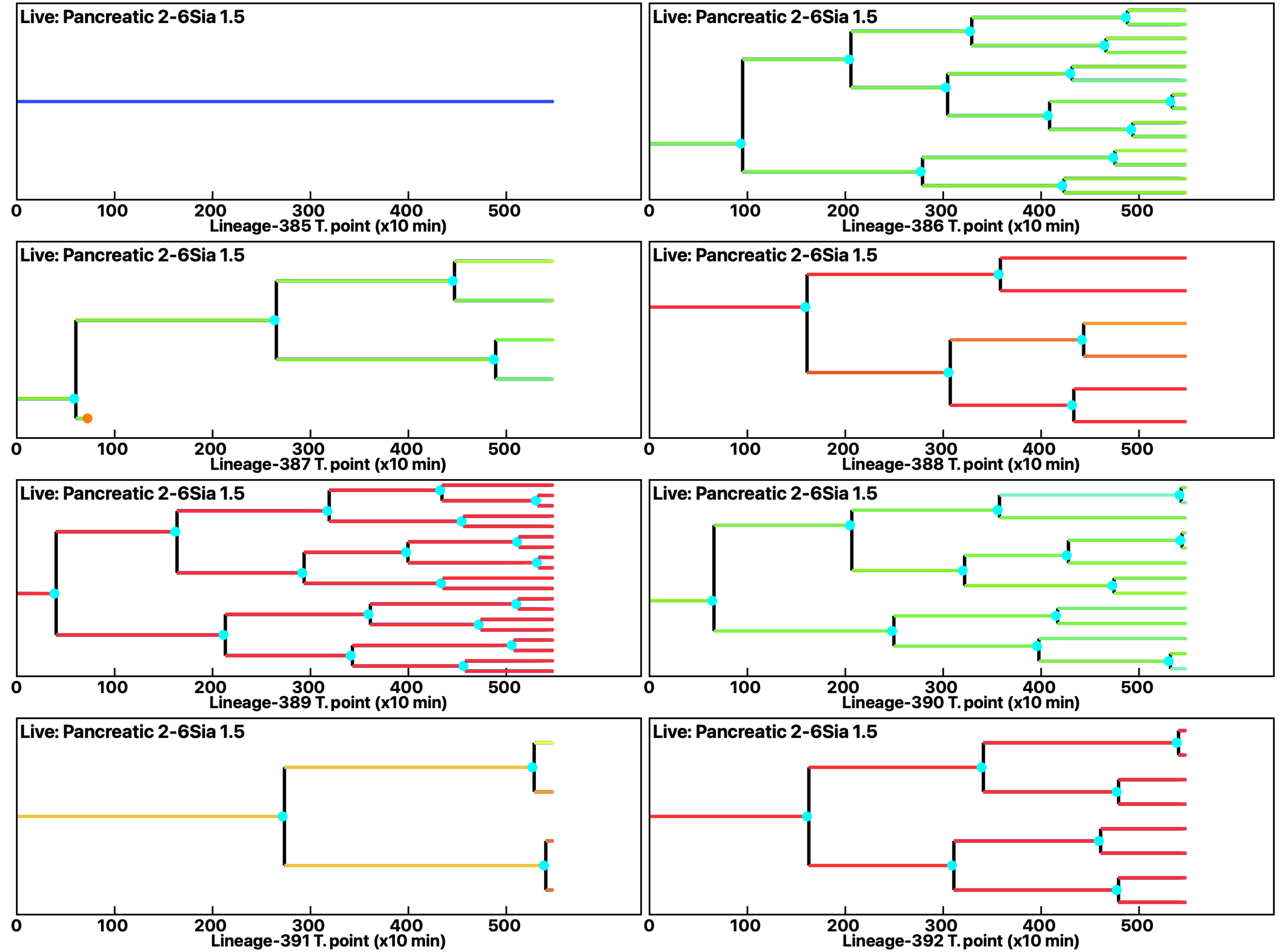

Analysis: Simulation, Treat.: Pancreatic 2-6Sia 1.5

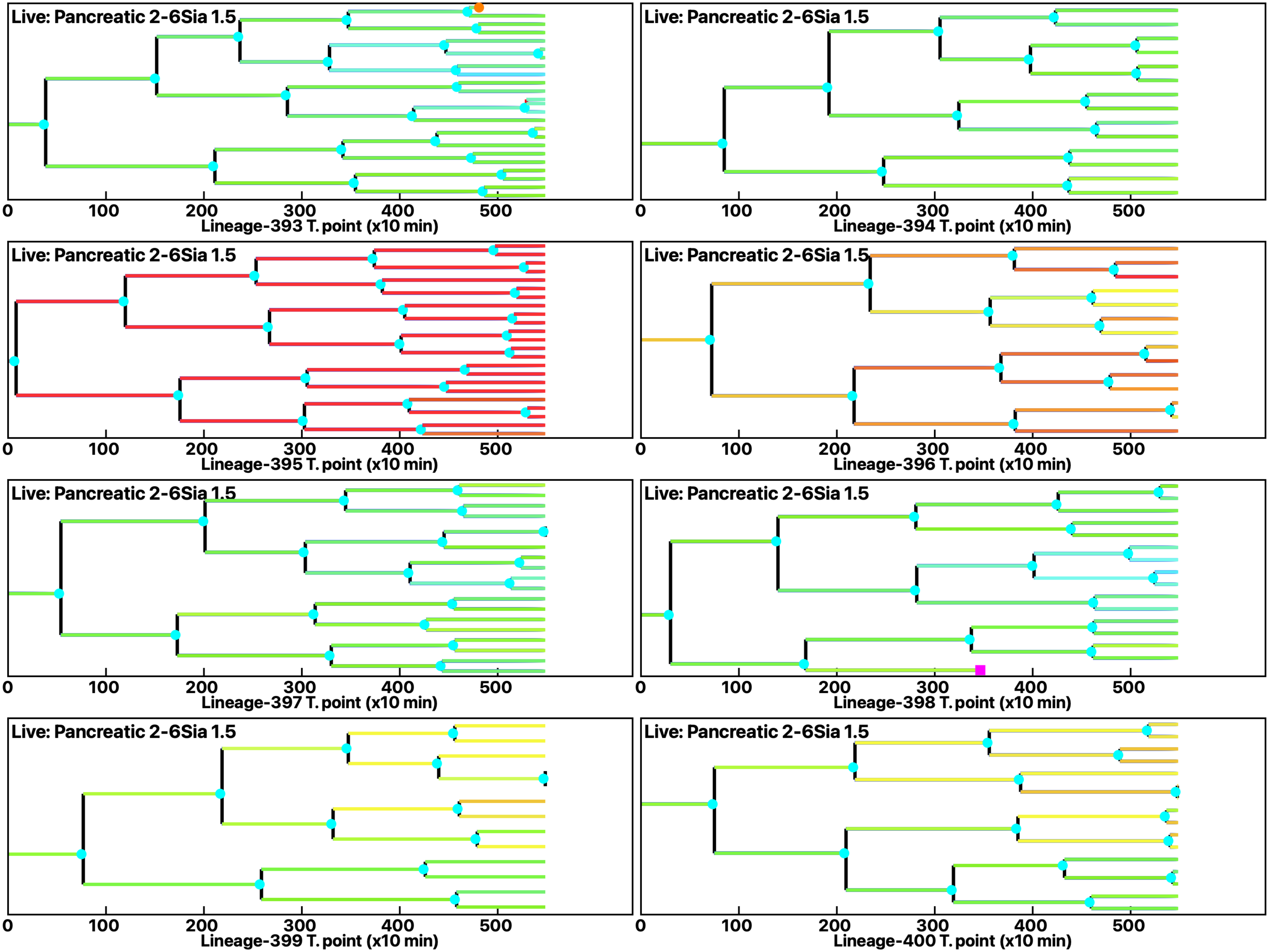

Analysis: Simulation, Treat.: Pancreatic 2-6Sia 1.5

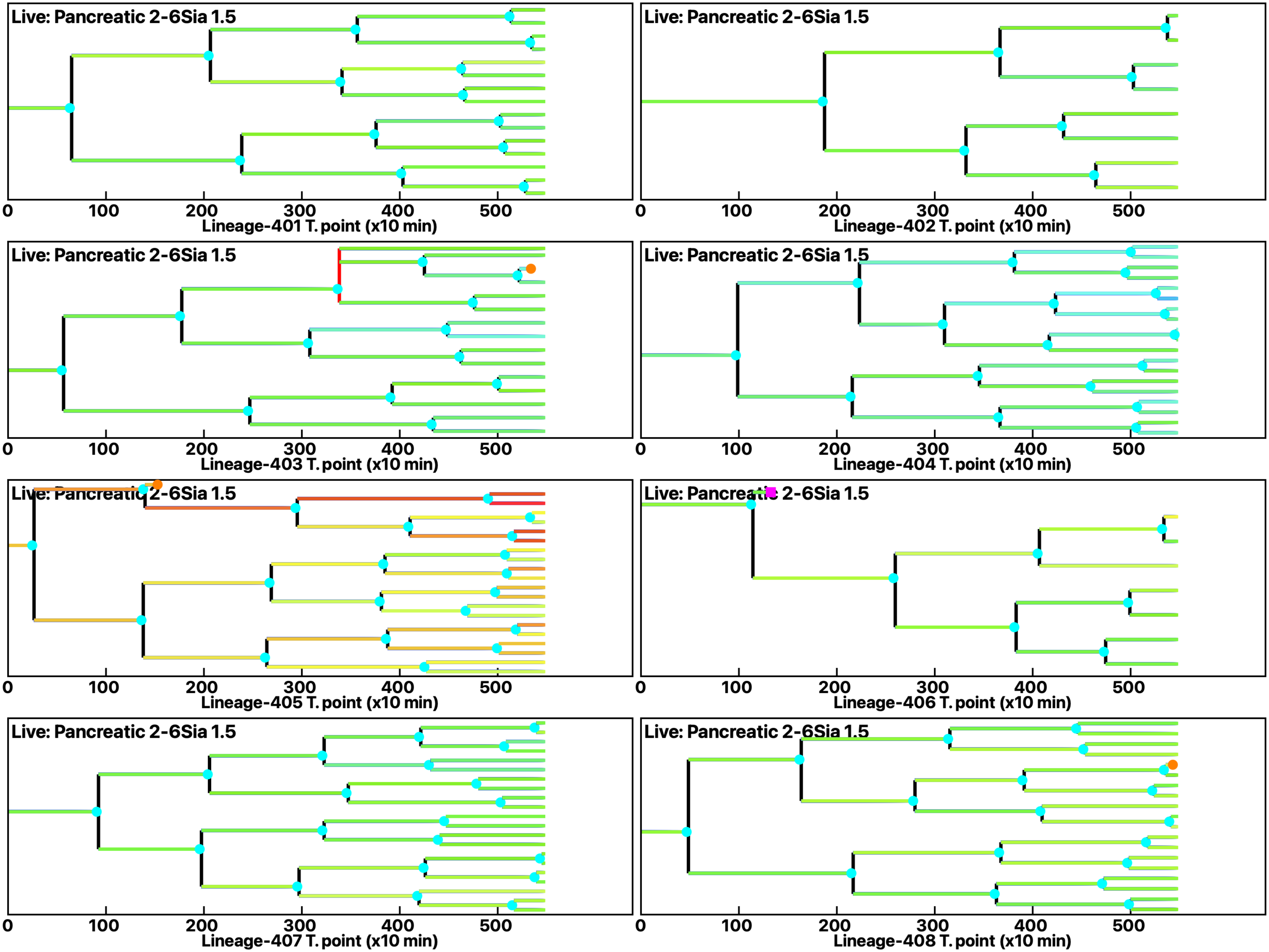

**Analysis: Simulation, Treat.: Pancreatic 2-6Sia 1.5**

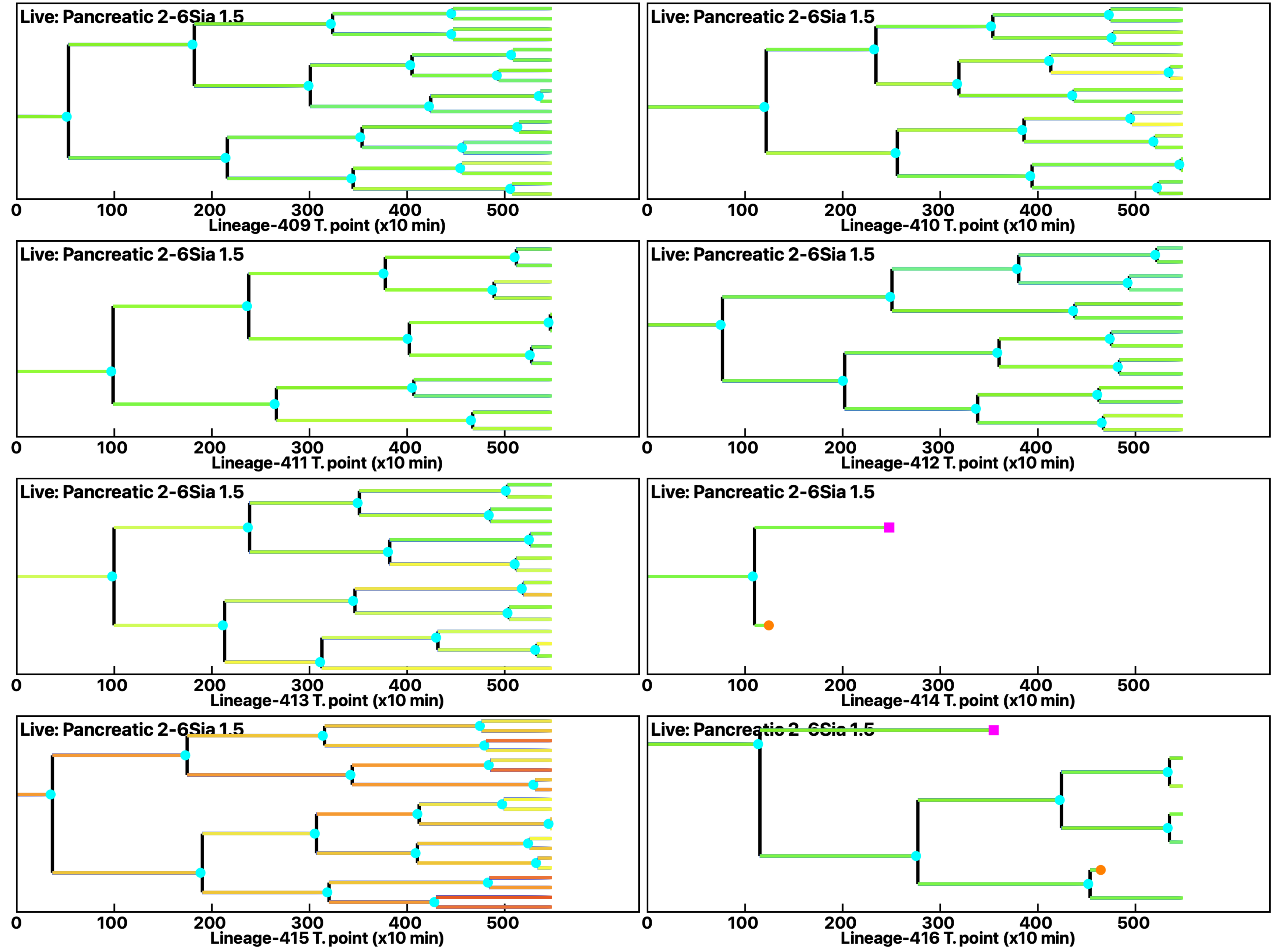

Analysis: Simulation, Treat.: Pancreatic 2-6Sia 1.5

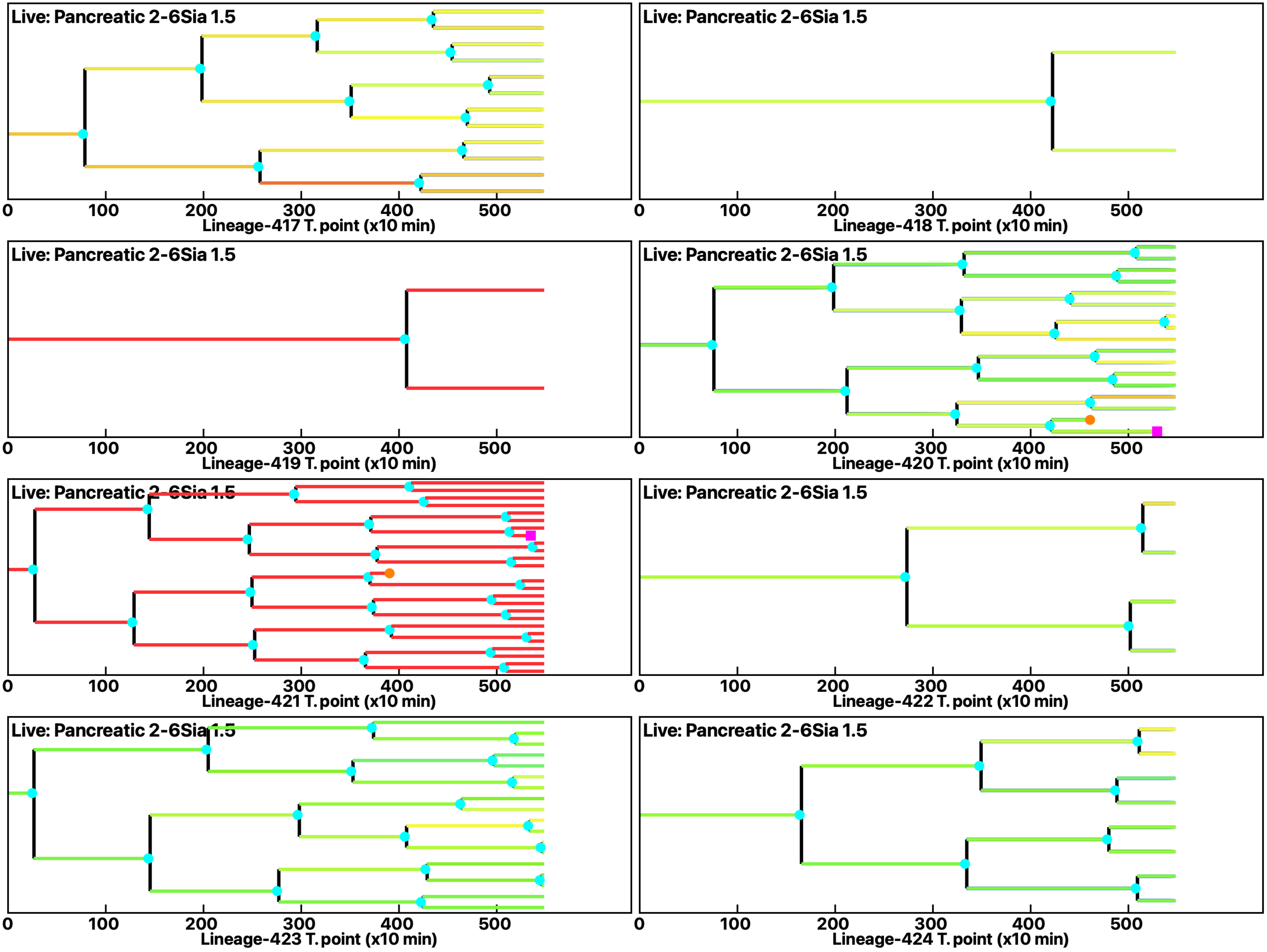

Analysis: Simulation, Treat.: Pancreatic 2-6Sia 1.5

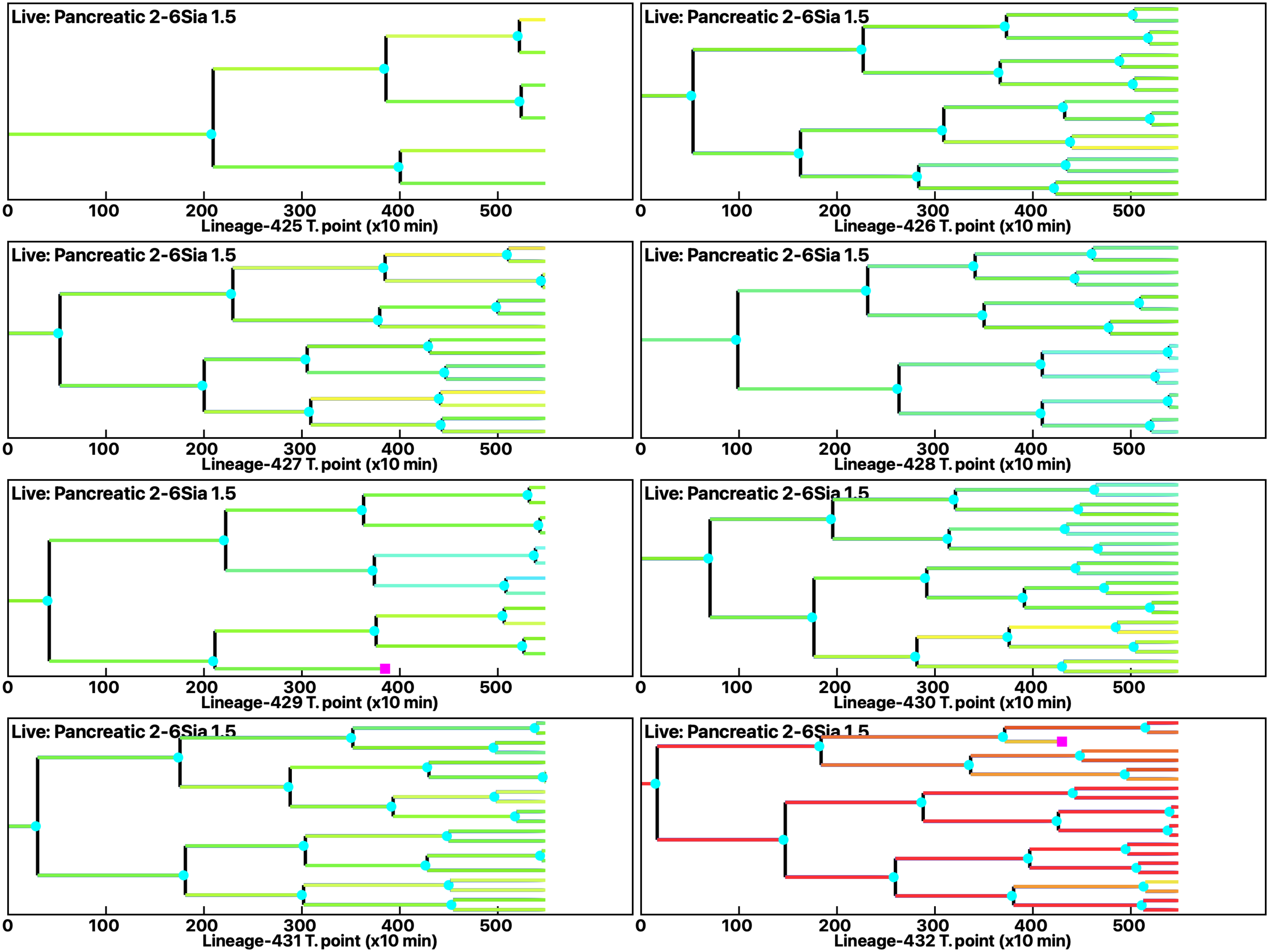

Analysis: Simulation, Treat.: Pancreatic 2-6Sia 1.5

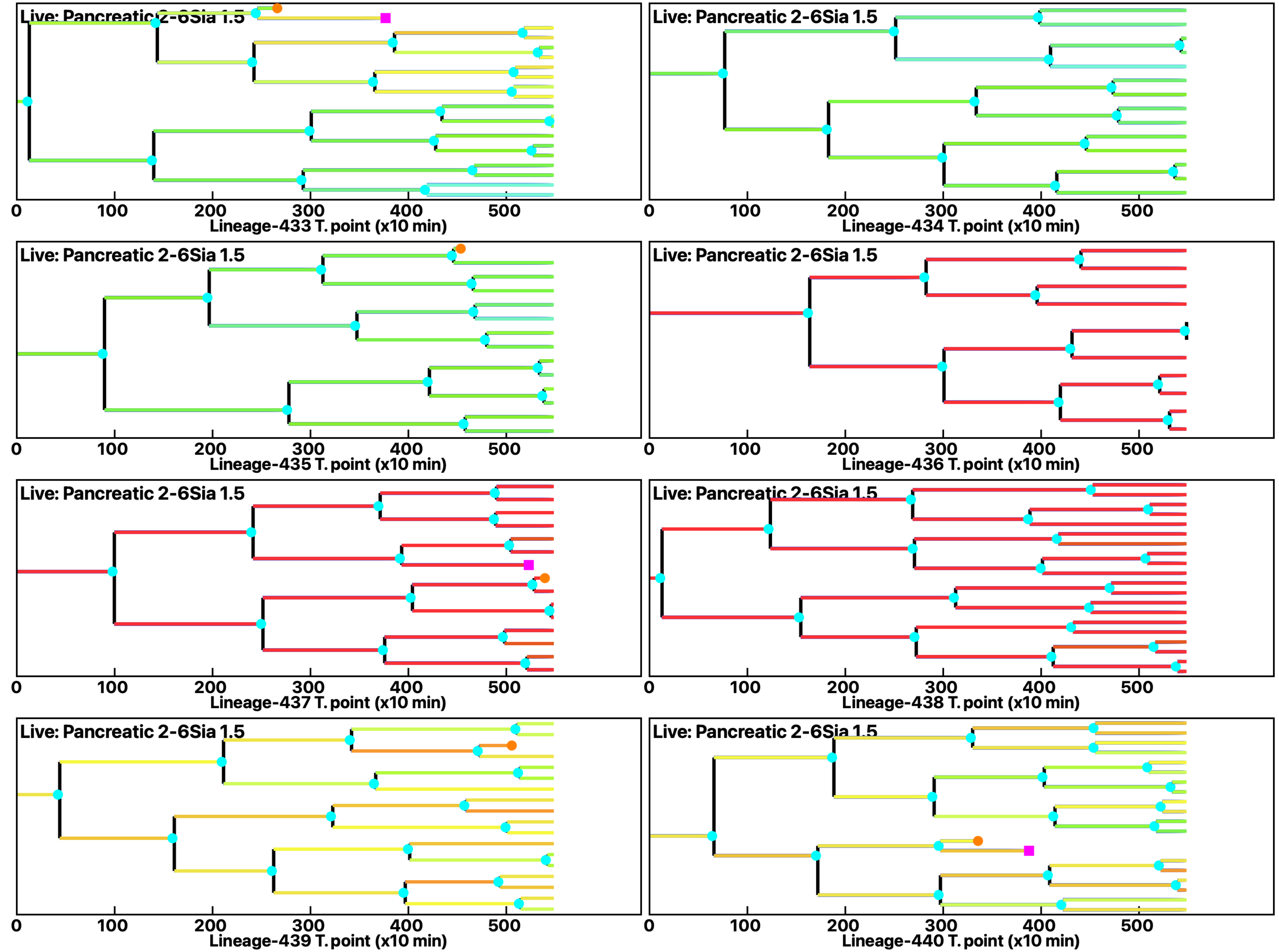

Analysis: Simulation, Treat.: Pancreatic 2-6Sia 1.5

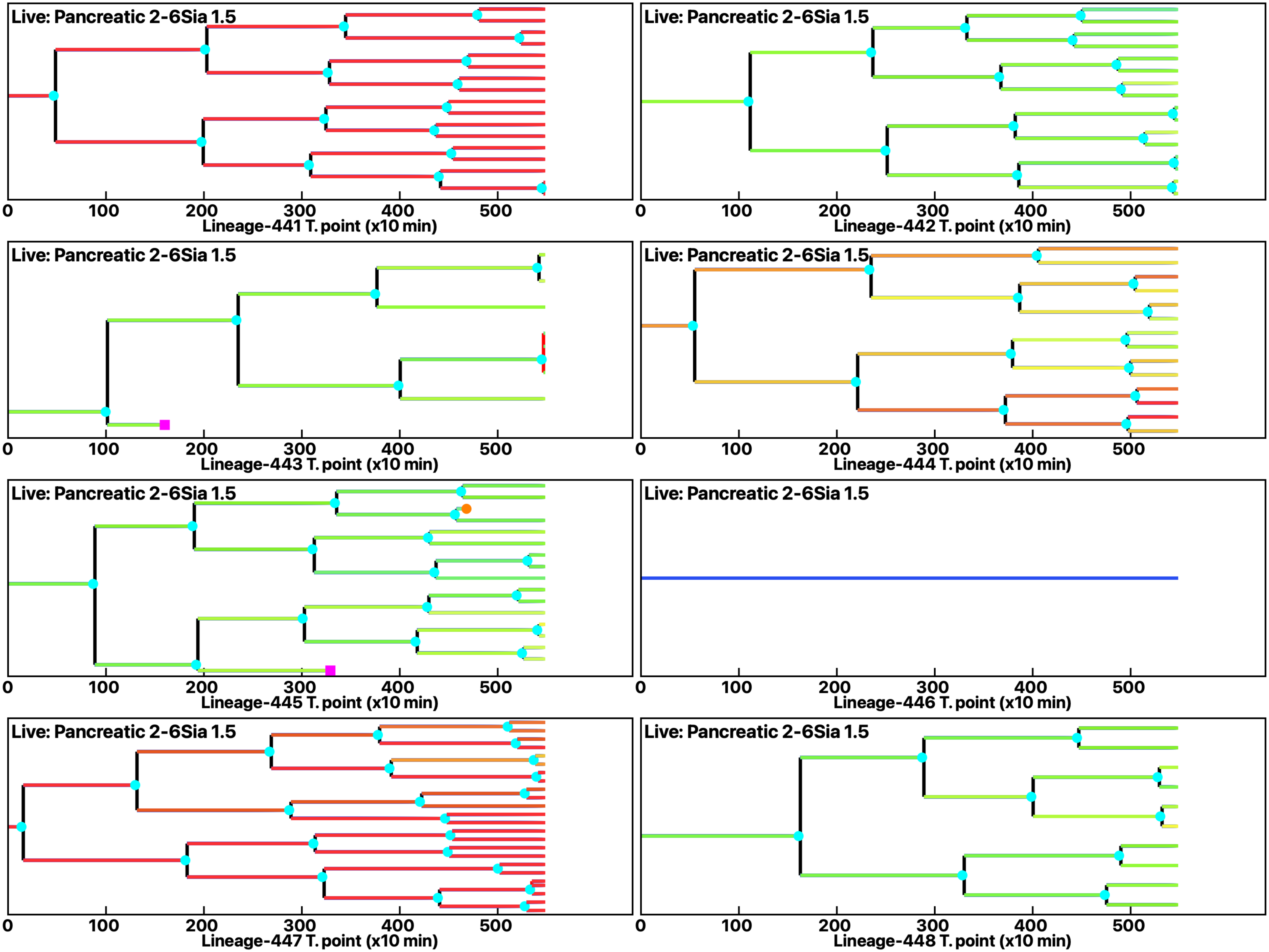

Analysis: Simulation, Treat.: Pancreatic 2-6Sia 1.5

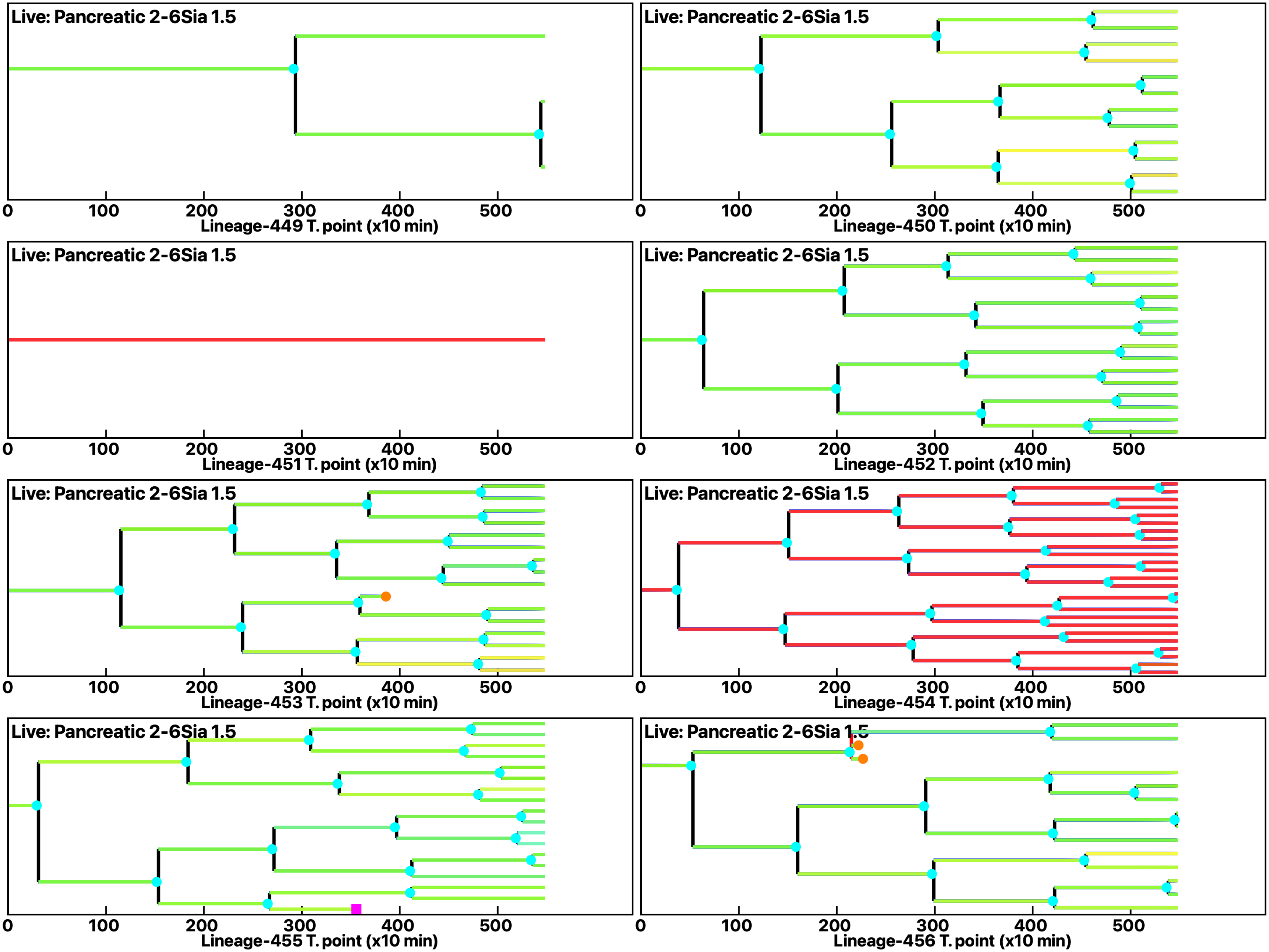

Analysis: Simulation, Treat.: Pancreatic 2-6Sia 1.5

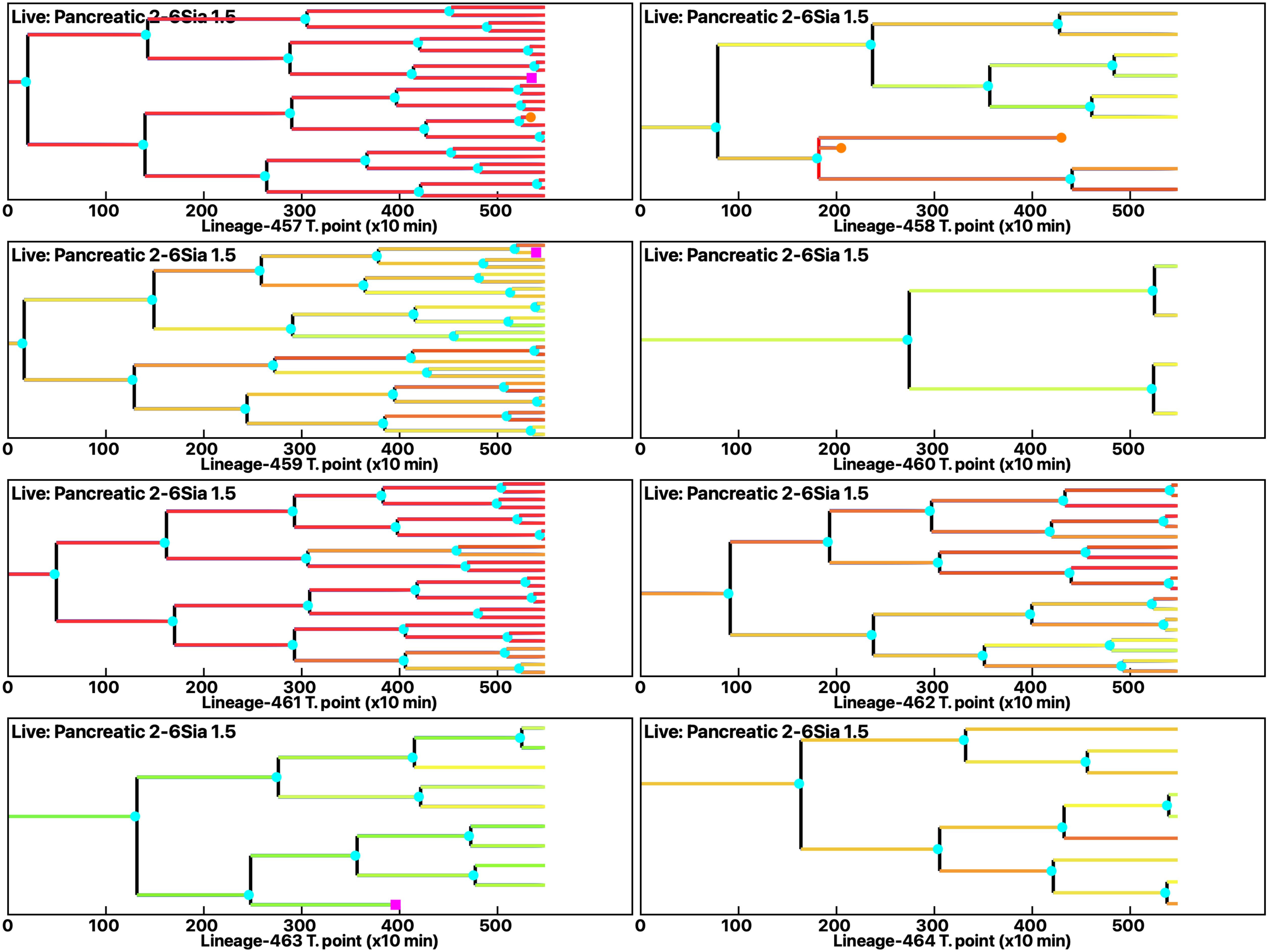

Analysis: Simulation, Treat.: Pancreatic 2-6Sia 1.5

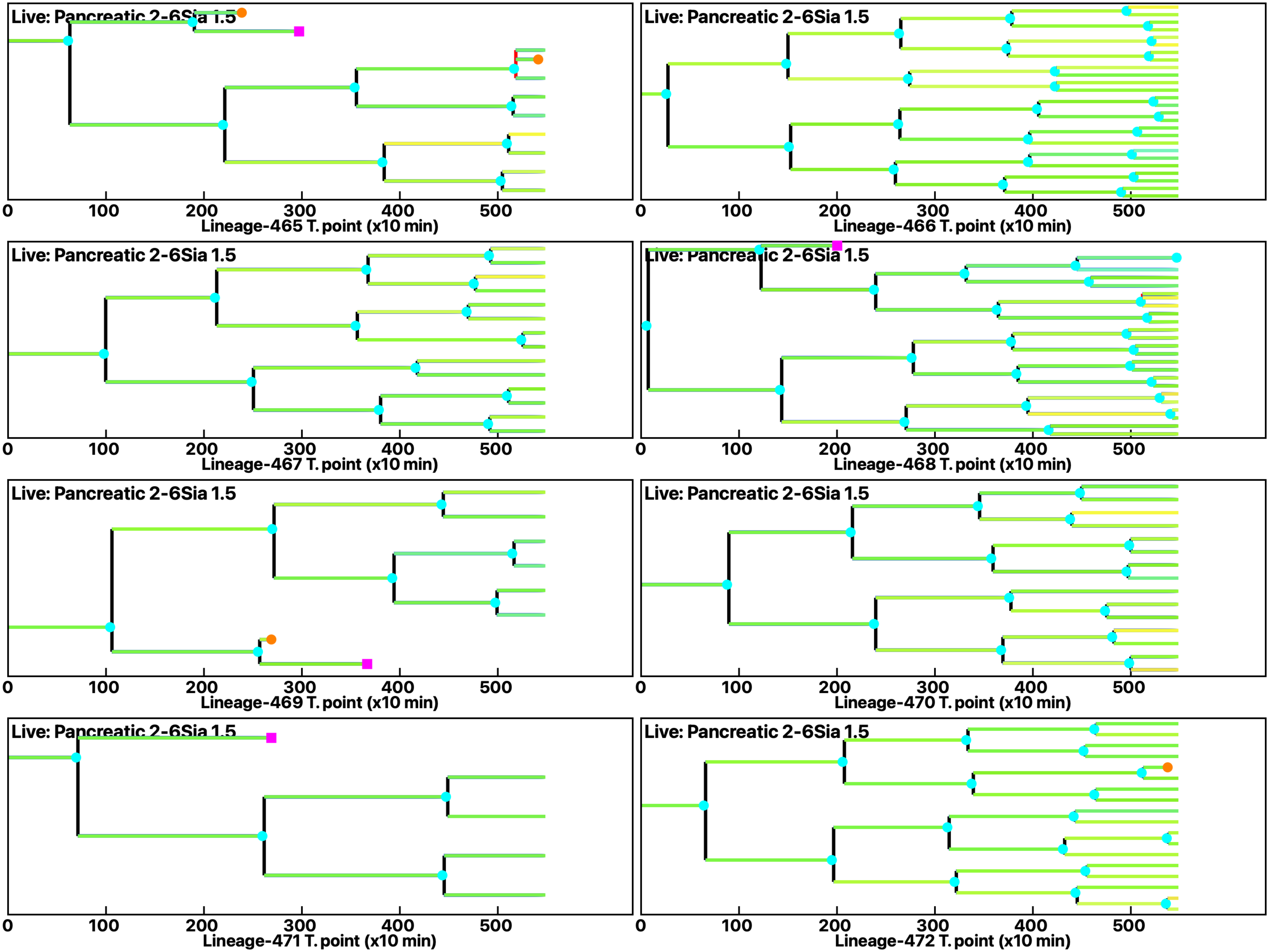

Analysis: Simulation, Treat.: Pancreatic 2-6Sia 1.5

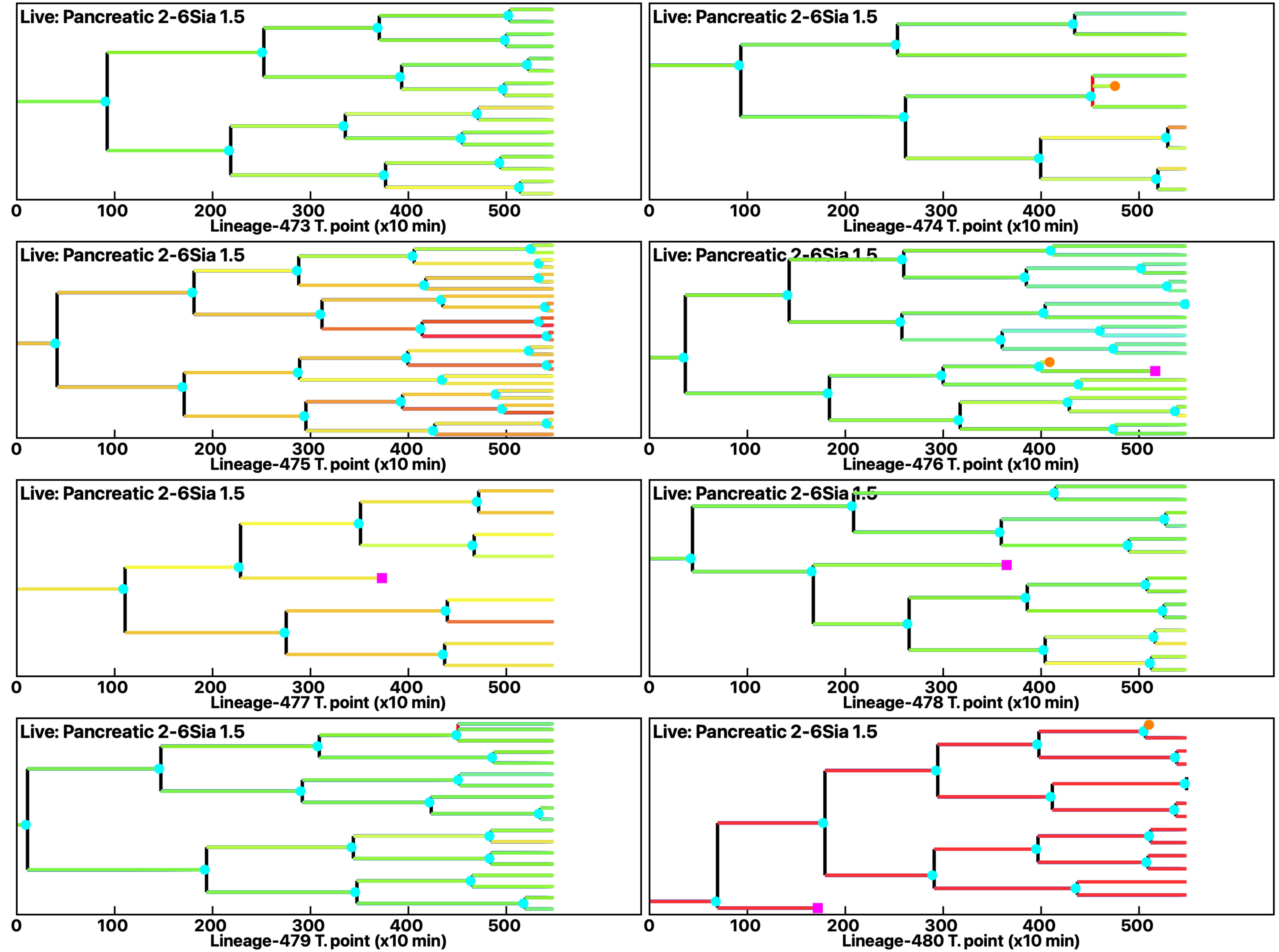

Analysis: Simulation, Treat.: Pancreatic 2-6Sia 1.5

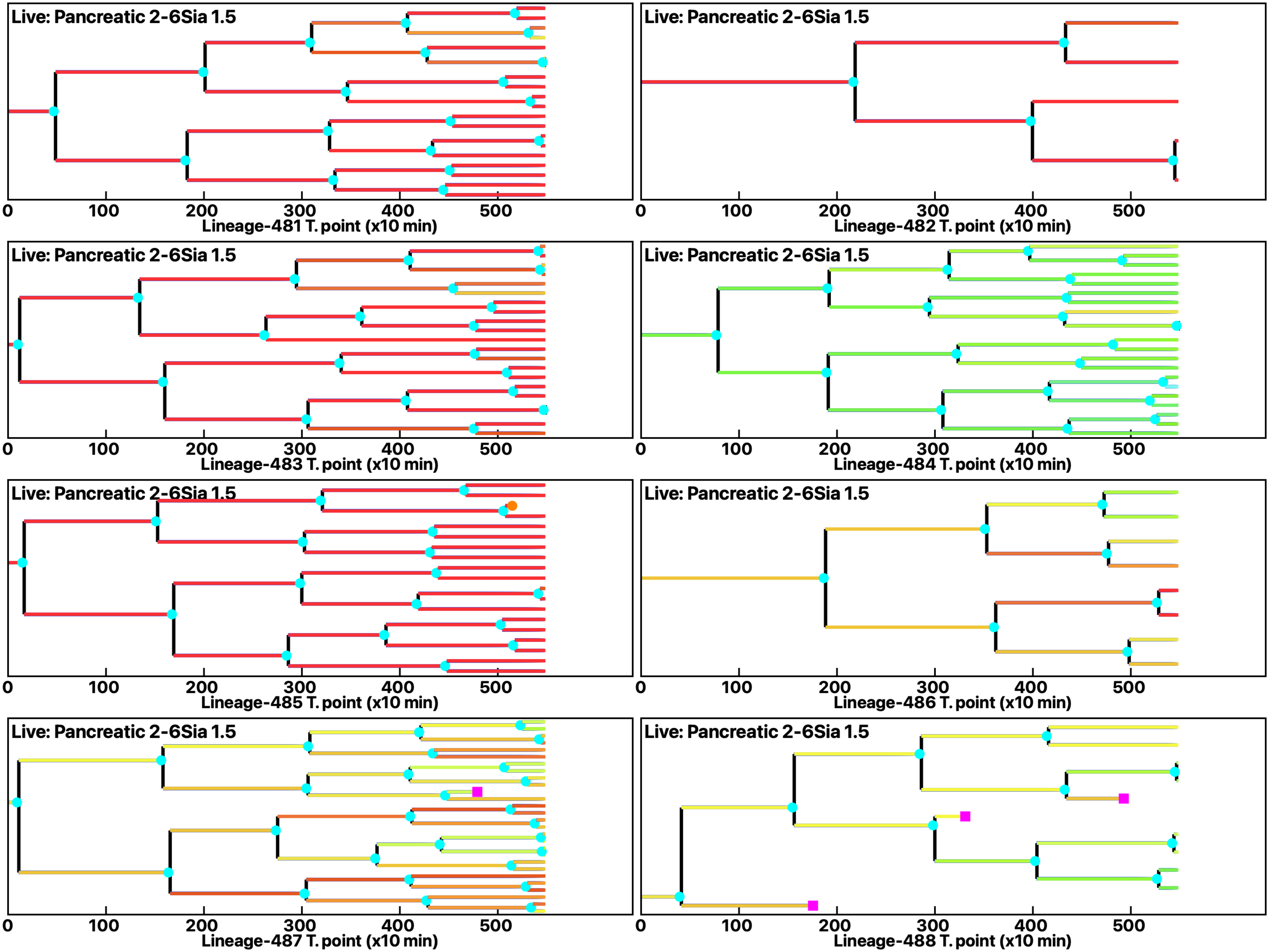

Analysis: Simulation, Treat.: Pancreatic 2-6Sia 1.5

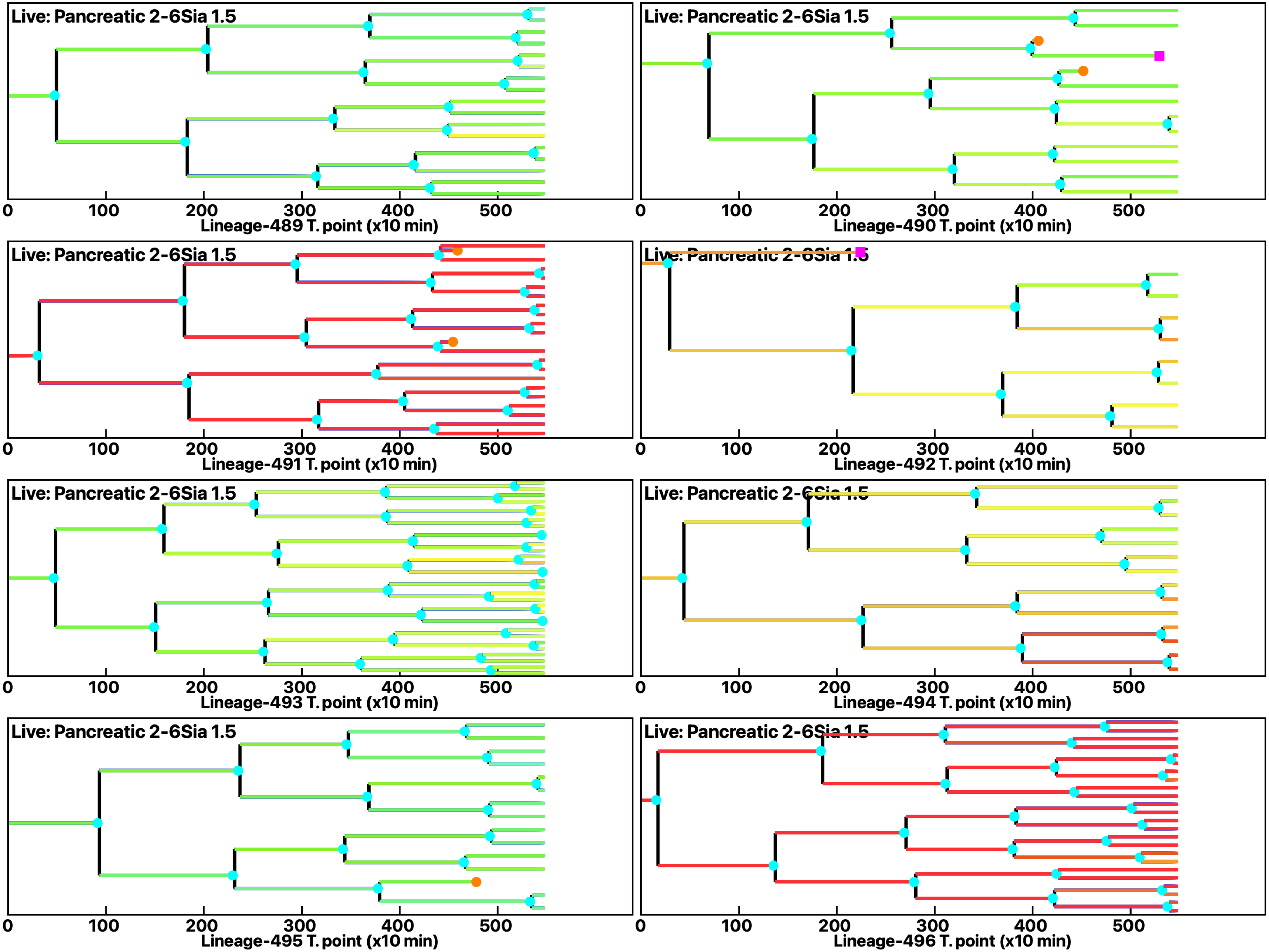

Analysis: Simulation, Treat.: Pancreatic 2-6Sia 1.5

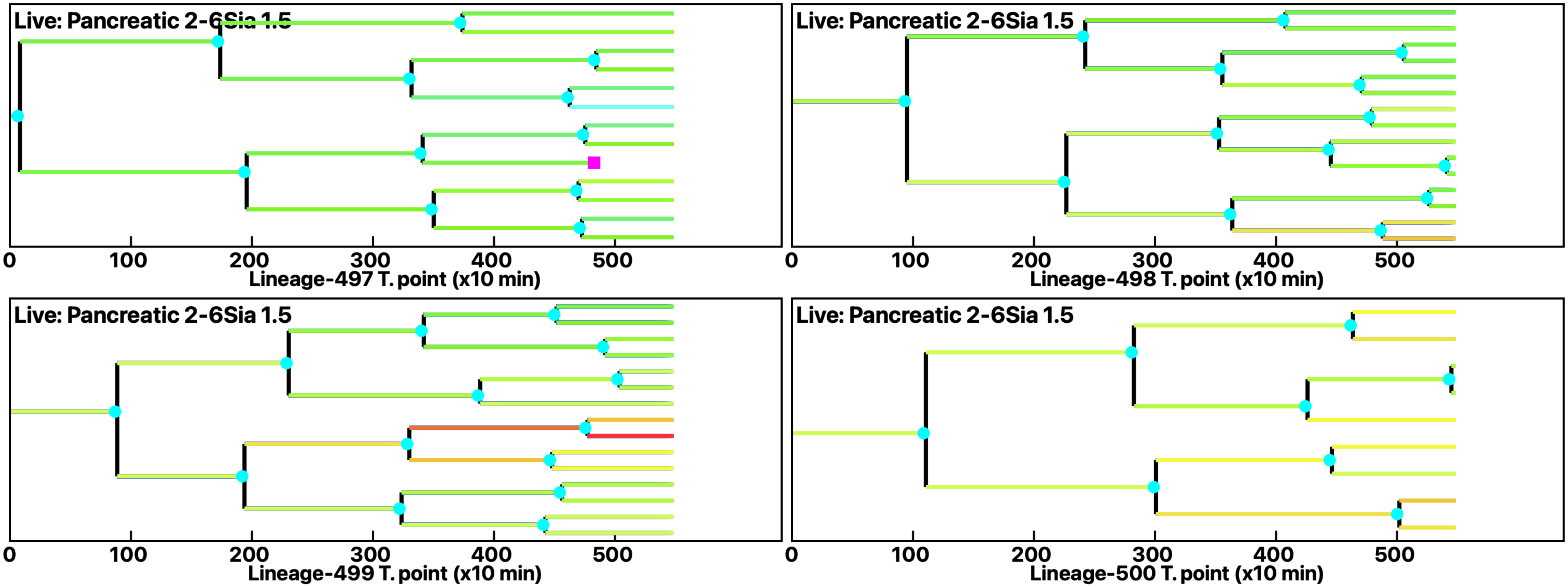
