## Supplementary Data 5 for "Cell Fate Simulation Reveals Cancer Cell Features in the Tumor Microenvironment"

### Cervical cancer

| | | | Cell population size | | | | 2-6Sia levels | | | | Number of reproductive cells produced through multipolar cell division | | | | Supressive | Permissive | Lethal |
|---|---|---|---|---|---|---|---|---|---|---|---|---|---|---|---|---|---|
| ID | Best call | PFS | 2-6Sia 1.5 | 2-6Sia 1.0 | 2-6Sia 0.5 | 2-6Sia 0.25 | 2-6Sia 1.5 | 2-6Sia 1.0 | 2-6Sia 0.5 | 2-6Sia 0.25 | 2-6Sia 1.5 | 2-6Sia 1.0 | 2-6Sia 0.5 | 2-6Sia 0.25 | % | % | % |
| TCGA-EK-A2H1 | 1 | 540 | 2176 | 2177.09617 | 2384.44195 | 2628.69364 | 34998 | 13186.3196 | 9252.45832 | 0 | 0 | 2.90021692 | 7.84355903 | 2.77385892 | 30.3384982 | 11.9347605 | 28.9418072 |
| TCGA-EK-A2H0 | 1 | 463 | 2193 | 2137.45987 | 2244.23833 | 2690.64315 | 34245 | 15717.2422 | 9010.28844 | 0 | 0 | 1.93347795 | 0 | 1.84923928 | 30.7022515 | 13.4808049 | 27.1833828 |
| TCGA-ZJ-A8QQ | 1 | 0 | 2414 | 2439.08243 | 2622.69005 | 2750.74343 | 37911 | 15435.9212 | 10127.0152 | 0 | 0 | 0 | 3.92177952 | 1.84923928 | 29.8384343 | 11.615669 | 29.4526366 |
| TCGA-ZJ-AB0I | 1 | 186 | 2250 | 2093.95662 | 2267.76901 | 2873.71784 | 33002 | 13092.5459 | 8670.07407 | 0 | 0 | 2.90021692 | 3.92177952 | 0.92461964 | 27.7834775 | 14.6430289 | 29.7158808 |
| TCGA-DS-A0VL | 1 | 4483 | 2205 | 2247.66811 | 2591.31582 | 2722.08022 | 35274 | 14765.0043 | 10070.1494 | 0 | 0 | 2.90021692 | 7.84355903 | 0.92461964 | 31.6866309 | 13.708928 | 28.6181104 |
| TCGA-EK-A2RE | 1 | 4879 | 2966 | 3080.99711 | 3063.89025 | 3293.49516 | 42673 | 21029.4729 | 12002.6062 | 0 | 0 | 4.83369487 | 7.84355903 | 0.92461964 | 25.4733723 | 4.10099908 | 37.0506124 |
| TCGA-C5-A7XC | 1 | 2226 | 2147 | 2471.95155 | 2522.68467 | 2807.14523 | 32717 | 16279.8843 | 9703.46297 | 0 | 0 | 6.76717281 | 3.92177952 | 2.77385892 | 32.7473045 | 12.9806297 | 28.7190958 |
| TCGA-GH-A9DA | 1 | 1221 | 2375 | 2378.17787 | 2625.63139 | 2680.47234 | 37938 | 18028.7151 | 10286.8277 | 0 | 0 | 1.93347795 | 0 | 0.92461964 | 30.7915827 | 11.6314351 | 29.3164522 |
| TCGA-MU-A51Y | 1 | 90 | 2309 | 2245.73463 | 2217.76632 | 2712.83402 | 34772 | 12863.4288 | 8456.33708 | 0 | 0 | 0 | 0 | 1.84923928 | 32.5259247 | 11.5300449 | 29.3671111 |
| TCGA-EK-A2IP | 1 | 586 | 2046 | 2129.72596 | 2445.22953 | 2858.92393 | 34070 | 14300.0029 | 9862.29504 | 0 | 0 | 1.93347795 | 7.84355903 | 1.84923928 | 30.5423682 | 14.2385095 | 29.0754164 |
| TCGA-MY-A913 | 1 | 8 | 2227 | 2455.51699 | 2533.46957 | 2832.10996 | 34876 | 16493.5336 | 10040.736 | 0 | 0 | 3.86695589 | 3.92177952 | 0 | 32.4327025 | 11.7908442 | 27.4424388 |
| TCGA-VS-A9V2 | 1 | 27 | 2311 | 2473.88503 | 2621.70961 | 2827.48686 | 36213 | 16162.9089 | 10459.386 | 0 | 0 | 0 | 3.92177952 | 0.92461964 | 33.1538636 | 12.2567167 | 28.1794521 |
| TCGA-FU-A2QG | 1 | 607 | 2181 | 2067.85466 | 2459.9362 | 2301.37828 | 30983 | 13716.0926 | 9775.03544 | 0 | 0 | 1.93347795 | 0 | 1.84923928 | 31.1682197 | 14.9372181 | 26.9059288 |
| TCGA-VS-A9U7 | 1 | 1203 | 2202 | 2296.00506 | 2624.65094 | 2841.35615 | 35679 | 14761.1374 | 10046.6187 | 0 | 0 | 4.83369487 | 0 | 0.92461964 | 32.5430155 | 12.5071869 | 28.1026463 |
| TCGA-C5-A2LY | 1 | 954 | 2078 | 2190.63051 | 2316.79125 | 2730.4018 | 35245 | 16453.8973 | 8992.64043 | 0 | 0 | 1.93347795 | 3.92177952 | 0.92461964 | 33.2348143 | 12.9872163 | 26.7566215 |
| TCGA-EA-A44S | 1 | 3442 | 2090 | 2174.19595 | 2455.03398 | 2753.51729 | 31075 | 15428.1873 | 9653.46028 | 0 | 0 | 2.90021692 | 7.84355903 | 0.92461964 | 31.9293733 | 12.9234347 | 28.4402551 |
| TCGA-4J-AA1J | 1 | 747 | 2443 | 2498.05351 | 2769.75678 | 2955.08437 | 37116 | 16667.5466 | 11039.8093 | 0 | 0 | 0 | 3.92177952 | 0 | 27.1453085 | 9.60784202 | 34.2171105 |
| TCGA-C5-A1BI | 1 | 803 | 2174 | 2376.2444 | 2535.43046 | 2952.31051 | 34244 | 16379.4584 | 9990.73332 | 0 | 0 | 4.83369487 | 7.84355903 | 0 | 33.0991018 | 12.1053141 | 28.2811574 |
| TCGA-FU-A5XV | 1 | 2496 | 2462 | 1987.61533 | 2627.59228 | 2557.49793 | 36036 | 12802.5242 | 10384.8722 | 0 | 0 | 1.93347795 | 7.84355903 | 1.84923928 | 31.8770223 | 11.9799696 | 28.6988876 |
| TCGA-EX-A3L1 | 1 | 27 | 2696 | 2754.23933 | 2942.31508 | 3254.66113 | 40591 | 18020.0145 | 11722.199 | 0 | 0 | 2.90021692 | 3.92177952 | 2.77385892 | 29.7501367 | 8.25914556 | 30.7550547 |
| TCGA-VS-A9UC | 1 | 1367 | 2264 | 2540.59002 | 2733.48032 | 2972.65214 | 33995 | 17761.8952 | 10563.3131 | 0 | 0 | 3.86695589 | 3.92177952 | 2.77385892 | 31.4454554 | 12.9435541 | 29.3868786 |
| TCGA-C5-A1BL | 1 | 2669 | 2134 | 2173.22921 | 2504.05622 | 3024.43084 | 33856 | 12910.799 | 9757.38744 | 0 | 0 | 4.83369487 | 0 | 1.84923928 | 32.897909 | 13.2599805 | 26.4519776 |
| TCGA-EK-A2RC | 1 | 103 | 2159 | 2115.22487 | 2585.43315 | 2812.69295 | 35592 | 15148.7997 | 10261.3361 | 0 | 0 | 1.93347795 | 0 | 1.84923928 | 32.7092618 | 12.541301 | 29.4470001 |
| TCGA-MA-AA41 | 1 | 1415 | 2276 | 2116.19161 | 2641.3185 | 2690.64315 | 33935 | 13084.812 | 10056.4231 | 0 | 0 | 1.93347795 | 0 | 0.92461964 | 31.7476413 | 13.2334753 | 28.2263949 |
| TCGA-ZJ-AAXN | 1 | 3988 | 2147 | 2116.19161 | 2574.64825 | 2772.00968 | 33046 | 13933.6088 | 10130.9369 | 0 | 0 | 4.83369487 | 3.92177952 | 1.84923928 | 29.2209327 | 15.1568221 | 29.8362595 |
| TCGA-C5-A1MP | 1 | 652 | 2421 | 2322.10701 | 2649.16206 | 2769.23582 | 41081 | 16276.9841 | 10400.5593 | 0 | 0 | 0 | 3.92177952 | 0.92461964 | 32.4132008 | 10.7060203 | 29.9765582 |
| TCGA-RA-A741 | 1 | 828 | 2477 | 2290.20463 | 2476.60376 | 2665.67842 | 38005 | 16110.705 | 9809.35101 | 0 | 0 | 0 | 0 | 0 | 33.3296005 | 12.1814392 | 27.9727007 |
| TCGA-C5-A1BJ | 1 | 986 | 2319 | 2295.03832 | 2500.13444 | 2768.3112 | 36250 | 15553.8633 | 9697.5803 | 0 | 0 | 1.93347795 | 7.84355903 | 0 | 32.4092973 | 11.1718976 | 29.675679 |
| TCGA-ZJ-AAXD | 1 | 940 | 2363 | 2467.11786 | 2624.65094 | 2859.84855 | 39326 | 16234.4476 | 10213.2943 | 0 | 0 | 1.93347795 | 0 | 2.77385892 | 32.1456771 | 10.438814 | 32.6545253 |
| TCGA-C5-A902 | 1 | 1667 | 2135 | 1988.58207 | 2146.19384 | 2672.15076 | 31763 | 13398.0354 | 8188.67563 | 0 | 0 | 0 | 0 | 0 | 34.5077163 | 16.1807332 | 26.4542786 |
| TCGA-LP-A4AU | 1 | 43 | 2066 | 2405.24657 | 2408.95307 | 2790.50207 | 31156 | 16672.3803 | 9184.80763 | 0 | 0 | 1.93347795 | 7.84355903 | 0 | 31.6735454 | 13.0920111 | 29.9906299 |
| TCGA-JW-A5VG | 1 | 383 | 1889 | 2274.7368 | 2288.35835 | 2557.49793 | 31100 | 15765.5792 | 9115.19604 | 0 | 0 | 1.93347795 | 7.84355903 | 1.84923928 | 32.0238565 | 14.7626852 | 25.6865762 |
| TCGA-FU-A3YQ | 1 | 632 | 2114 | 2371.4107 | 2623.6705 | 2857.07469 | 32968 | 15525.8279 | 10825.0919 | 0 | 0 | 2.90021692 | 7.84355903 | 0.92461964 | 32.9422711 | 11.1562859 | 28.5485253 |
| TCGA-EA-A3HS | 1 | 1103 | 2371 | 2513.52133 | 2316.79125 | 2922.72268 | 35278 | 15498.7592 | 9077.93913 | 0 | 0 | 3.86695589 | 2.94133464 | 1.84923928 | 31.0361961 | 12.4298407 | 29.6619217 |
| TCGA-C5-A1MQ | 1 | 954 | 2454 | 2374.31092 | 2715.83231 | 2662.90456 | 34888 | 15504.5597 | 10291.7299 | 0 | 0 | 2.90021692 | 0 | 3.69847856 | 33.7606119 | 11.1885569 | 27.5061572 |
| TCGA-EK-A2PL | 1 | 621 | 1927 | 1945.07881 | 2518.76289 | 2658.28147 | 29448 | 12008.8315 | 9670.12784 | 0 | 0 | 2.90021692 | 3.92177952 | 0.92461964 | 32.4717257 | 14.5625651 | 27.5384844 |
| TCGA-C5-A8YT | 1 | 45 | 2019 | 1810.7021 | 2439.34686 | 2797.89903 | 33679 | 10733.7028 | 9549.53312 | 0 | 0 | 4.83369487 | 1.96088976 | 2.77385892 | 29.6453815 | 15.9033935 | 32.0757939 |
| TCGA-VS-A8EG | 1 | 18 | 2271 | 2312.43962 | 2450.13175 | 2758.14039 | 39022 | 15537.4288 | 9601.4967 | 0 | 0 | 2.90021692 | 3.92177952 | 2.77385892 | 33.0420004 | 11.7137114 | 30.9619054 |
| TCGA-C5-A2LV | 2 | 799 | 2205 | 2271.83659 | 2533.46957 | 2795.12517 | 31750 | 14607.4259 | 9945.63285 | 0 | 0 | 2.90021692 | 0 | 0.92461964 | 30.269412 | 10.8613244 | 29.4532272 |
| TCGA-C5-A1BK | 2 | 1847 | 2087 | 2263.13594 | 2470.7211 | 2789.57746 | 29146 | 15925.0911 | 9725.03275 | 0 | 0 | 6.76717281 | 0 | 0 | 34.1152299 | 12.5666722 | 26.7445767 |
| TCGA-Q1-A73O | 2 | 2056 | 2213 | 2043.68619 | 2714.85187 | 2656.43223 | 35352 | 15172.9682 | 11023.1418 | 0 | 0 | 0 | 7.84355903 | 0 | 32.835116 | 14.4715868 | 26.9438254 |
| TCGA-JW-A5VL | 2 | 0 | 2317 | 2127.79248 | 2528.56734 | 2981.89834 | 31087 | 12622.7108 | 9969.16353 | 0 | 0 | 1.93347795 | 3.92177952 | 0.92461964 | 32.2629079 | 13.574749 | 27.3415519 |
| TCGA-JX-A5QV | 2 | 81 | 2258 | 2431.34852 | 2318.75214 | 2761.83887 | 37634 | 17536.645 | 8874.0066 | 0 | 0 | 1.93347795 | 0 | 0.92461964 | 31.8499394 | 13.183727 | 26.6327565 |
| TCGA-C5-A1M7 | 2 | 57 | 2349 | 2274.7368 | 2526.60645 | 2907.00415 | 33808 | 14675.0976 | 9992.69421 | 0 | 0 | 1.93347795 | 7.84355903 | 0.92461964 | 33.1103521 | 9.58197506 | 29.4400347 |
| TCGA-IR-A3L7 | 2 | 1551 | 2289 | 2260.23572 | 2618.76827 | 2919.0242 | 35396 | 15243.5401 | 10658.4163 | 0 | 0 | 3.86695589 | 3.92177952 | 2.77385892 | 27.429426 | 10.5411934 | 39.2854238 |
| TCGA-C5-A905 | 2 | 854 | 2440 | 2641.13087 | 2475.62332 | 2826.56224 | 34458 | 15480.3912 | 9884.84527 | 0 | 0 | 3.86695589 | 3.92177952 | 0.92461964 | 32.8433013 | 10.7009635 | 32.2274021 |
| TCGA-EA-A97N | 2 | 202 | 2589 | 2443.91612 | 2605.04204 | 3220.45021 | 39928 | 18362.2401 | 10230.9423 | 0 | 0 | 6.76717281 | 3.92177952 | 2.77385892 | 28.4746554 | 8.89350562 | 34.539616 |
| TCGA-VS-A8EI | 2 | 524 | 2503 | 2223.49964 | 2607.00293 | 2721.1556 | 37082 | 15122.6978 | 9714.24786 | 0 | 0 | 1.93347795 | 7.84355903 | 0 | 33.4072201 | 12.5982712 | 28.3864515 |
| TCGA-HG-A2PA | 2 | 555 | 2264 | 2311.47289 | 2834.46615 | 3030.90318 | 33354 | 16595.0412 | 11069.2227 | 0 | 0 | 1.93347795 | 3.92177952 | 0 | 31.740363 | 12.7901034 | 29.5096583 |
| TCGA-VS-A94Z | 2 | 579 | 1917 | 2026.28489 | 2380.52017 | 2901.45643 | 29667 | 13305.2285 | 9604.43803 | 0 | 0 | 0 | 0 | 0 | 34.3902733 | 13.6772007 | 26.9393107 |
| TCGA-VS-A8EL | 2 | 1472 | 1987 | 2279.5705 | 2617.78783 | 2640.71369 | 30168 | 15736.577 | 10483.8971 | 0 | 0 | 1.93347795 | 7.84355903 | 0.92461964 | 32.2979296 | 12.9348944 | 27.1118619 |
| TCGA-C5-A2LT | 2 | 2383 | 2389 | 2174.19595 | 2405.03129 | 2758.14039 | 38211 | 12970.7368 | 9633.85138 | 0 | 0 | 1.93347795 | 7.84355903 | 0 | 31.8267276 | 12.8239035 | 28.4080335 |
| TCGA-EA-A5O9 | 2 | 369 | 2106 | 2233.16703 | 2254.04278 | 2631.4675 | 33099 | 15769.4461 | 8945.57908 | 0 | 0 | 1.93347795 | 3.92177952 | 0.92461964 | 31.4198053 | 13.2172148 | 28.8040523 |
| TCGA-VS-A94W | 2 | 542 | 2178 | 2300.83876 | 2674.65363 | 2778.48202 | 30104 | 14785.3059 | 10741.7541 | 0 | 0 | 2.90021692 | 3.92177952 | 1.84923928 | 33.3149239 | 11.2547281 | 27.971759 |
| TCGA-LP-A5U3 | 2 | 1112 | 2140 | 2180.96312 | 2413.85529 | 2738.72337 | 32258 | 13508.2437 | 9759.34833 | 0 | 0 | 0.96673897 | 0 | 0.92461964 | 31.2851325 | 13.2986248 | 27.1183127 |
| TCGA-EK-A2RM | 2 | 321 | 2709 | 2583.12654 | 2808.97458 | 3139.08368 | 37485 | 18640.6609 | 11137.8538 | 0 | 0 | 4.83369487 | 3.92177952 | 0.92461964 | 26.8677598 | 7.36270333 | 34.249729 |
| TCGA-EA-A5ZD | 2 | 825 | 2531 | 2302.77223 | 2788.38524 | 3018.88313 | 38241 | 14252.6327 | 10637.8269 | 0 | 0 | 1.93347795 | 3.92177952 | 0 | 30.4789805 | 9.93504939 | 30.5995862 |
| TCGA-EA-A3Y4 | 2 | 5271 | 2102 | 1995.34924 | 2445.22953 | 2651.80913 | 27000 | 14401.5105 | 9921.12173 | 0 | 0 | 1.93347795 | 3.92177952 | 0.92461964 | 33.4714857 | 14.7764242 | 25.5775839 |
| TCGA-IR-A3LC | 2 | 129 | 2225 | 2229.30007 | 2447.19042 | 2916.25035 | 36654 | 15209.7043 | 9370.11171 | 0 | 0 | 2.90021692 | 0 | 0.92461964 | 29.9286375 | 13.1411537 | 30.0836358 |
| TCGA-EX-A69L | 2 | 279 | 2415 | 2502.8872 | 2547.1958 | 2951.38589 | 34374 | 14737.9356 | 9774.055 | 0 | 0 | 4.83369487 | 1.96088976 | 0.92461964 | 33.4892935 | 12.1411106 | 26.976836 |
| TCGA-C5-A8ZZ | 2 | 0 | 2788 | 2751.33912 | 2872.7035 | 3263.90733 | 43890 | 18523.6855 | 11275.1161 | 0 | 0 | 6.76717281 | 3.92177952 | 3.69847856 | 27.6991073 | 5.95128254 | 38.2183837 |
| TCGA-XS-A8TJ | 2 | 109 | 2183 | 2323.07375 | 2410.91396 | 2847.82849 | 32793 | 16691.7151 | 9894.64972 | 0 | 0 | 1.93347795 | 0 | 0.92461964 | 30.5231307 | 11.0302053 | 30.7463839 |
| TCGA-JW-A852 | 2 | 444 | 2368 | 2196.43095 | 2688.37986 | 2715.60788 | 35435 | 14337.7057 | 10743.715 | 0 | 0 | 3.86695589 | 7.84355903 | 1.84923928 | 33.473382 | 11.9914442 | 28.9847277 |
| TCGA-VS-A9UU | 2 | 4385 | 2245 | 2401.37961 | 2371.69616 | 2800.67289 | 30674 | 14349.3066 | 9344.62014 | 0 | 0 | 2.90021692 | 3.92177952 | 2.77385892 | 34.359387 | 12.2858855 | 28.1663886 |
| TCGA-R2-A69V | 2 | 149 | 2292 | 2294.07158 | 2354.04815 | 2848.75311 | 31494 | 14063.1518 | 9322.06991 | 0 | 0 | 2.90021692 | 0 | 2.77385892 | 33.0446756 | 11.9780454 | 26.8144652 |
| TCGA-MY-A5BE | 2 | 343 | 2370 | 2121.99205 | 2436.40552 | 2872.79322 | 33765 | 13428.9711 | 9499.53043 | 0 | 0 | 6.76717281 | 0 | 0.92461964 | 34.9532721 | 12.9377605 | 25.8119989 |
| TCGA-VS-A950 | 2 | 834 | 2545 | 2302.77223 | 2810.93547 | 3085.45574 | 37787 | 15098.5293 | 11123.1472 | 0 | 0 | 0 | 3.92177952 | 1.84923928 | 32.2705981 | 9.95047363 | 30.5798252 |
| TCGA-VS-A8QF | 2 | 861 | 2289 | 2315.33984 | 2721.71498 | 2821.01452 | 31291 | 17209.8872 | 10775.0892 | 0 | 0 | 1.93347795 | 7.84355903 | 1.84923928 | 33.3251874 | 10.5415756 | 28.5903662 |
| TCGA-ZJ-AAXJ | 2 | 959 | 2247 | 2239.9342 | 2560.92202 | 2926.42116 | 37533 | 16153.2415 | 9752.48521 | 0 | 0 | 1.93347795 | 7.84355903 | 2.77385892 | 33.8276778 | 12.4301224 | 27.480232 |
| TCGA-EK-A2PK | 2 | 1031 | 2184 | 2222.5329 | 2558.96113 | 2616.67358 | 33517 | 15909.6233 | 10036.8142 | 0 | 0 | 1.93347795 | 3.92177952 | 0.92461964 | 33.7551012 | 13.9867403 | 26.8654367 |
| TCGA-PN-A8MA | 2 | 13 | 2304 | 2093.95662 | 2600.13982 | 2958.78285 | 34482 | 14793.0398 | 10052.5013 | 0 | 0 | 1.93347795 | 0 | 0.92461964 | 32.7915292 | 12.4811243 | 31.0647009 |
| TCGA-FU-A3WB | 2 | 1386 | 2526 | 2314.3731 | 2535.43046 | 2661.97994 | 39785 | 15026.9906 | 10299.5735 | 0 | 0 | 0 | 0 | 0.92461964 | 30.7308063 | 12.9496618 | 33.5786121 |
| TCGA-MA-AA3Z | 2 | 2234 | 2284 | 2115.22487 | 2410.91396 | 2611.12586 | 35359 | 16036.2661 | 9269.12589 | 0 | 0 | 1.93347795 | 3.92177952 | 0.92461964 | 33.5049512 | 14.7478832 | 25.5767852 |
| TCGA-EA-A43B | 2 | 5385 | 2157 | 2087.18944 | 2469.74065 | 3102.09889 | 32263 | 14957.3854 | 9399.52505 | 0 | 0 | 1.93347795 | 0 | 2.77385892 | 32.7875455 | 12.1280989 | 27.8776889 |
| TCGA-C5-A1M8 | 2 | 428 | 1895 | 2573.45915 | 2699.16475 | 2721.1556 | 32623 | 15700.8077 | 10694.6927 | 0 | 0 | 2.90021692 | 3.92177952 | 0 | 29.6748567 | 10.0645854 | 33.2602726 |
| TCGA-EA-A3HU | 2 | 474 | 2195 | 1976.9812 | 2489.34955 | 2556.57331 | 33849 | 11826.1179 | 9729.93498 | 0 | 0 | 1.93347795 | 3.92177952 | 0.92461964 | 34.1725838 | 14.0146787 | 26.3175813 |
| TCGA-C5-A7CH | 2 | 636 | 2532 | 2730.07086 | 3073.6947 | 3103.02351 | 41181 | 20021.1641 | 12649.6998 | 0 | 0 | 3.86695589 | 7.84355903 | 1.84923928 | 27.7996353 | 8.64160885 | 32.6336826 |
| TCGA-EX-A8YF | 2 | 1409 | 2147 | 2256.36876 | 2341.30237 | 2806.22061 | 34188 | 15811.0159 | 8726.93987 | 0 | 0 | 4.83369487 | 7.84355903 | 2.77385892 | 34.2149713 | 13.35678 | 26.7092651 |
| TCGA-Q1-A6DW | 2 | 11 | 2473 | 2104.59074 | 2665.82963 | 2960.63209 | 39787 | 14777.5719 | 10424.09 | 0 | 0 | 4.83369487 | 0 | 0.92461964 | 31.3015186 | 13.1359489 | 30.3239707 |
| TCGA-C5-A3HE | 2 | 729 | 2139 | 2290.20463 | 2546.21535 | 2750.74343 | 33684 | 15721.1092 | 9962.30042 | 0 | 0 | 1.93347795 | 3.92177952 | 2.77385892 | 33.0548774 | 13.09006 | 27.691197 |
| TCGA-EK-A2PI | 2 | 773 | 2470 | 2481.61894 | 2737.4021 | 3026.28008 | 38590 | 14244.8988 | 10791.7568 | 0 | 0 | 1.93347795 | 7.84355903 | 0 | 27.4087394 | 11.4346953 | 36.5550931 |
| TCGA-VS-A8QA | 2 | 1015 | 2180 | 2127.79248 | 2226.59032 | 2427.12656 | 33739 | 13471.5076 | 8927.93107 | 0 | 0 | 0 | 0 | 2.77385892 | 33.7837971 | 15.5806307 | 26.2383416 |
| TCGA-EA-A50E | 2 | 1992 | 2678 | 2625.66305 | 2873.68394 | 2791.42669 | 37794 | 16465.4982 | 11175.1107 | 0 | 0 | 4.83369487 | 3.92177952 | 0 | 28.1676125 | 9.03136093 | 33.3780133 |
| TCGA-EK-A2R7 | 2 | 788 | 2212 | 2427.48156 | 2497.19311 | 2686.02006 | 36404 | 17822.7997 | 9864.25593 | 0 | 0 | 4.83369487 | 7.84355903 | 0 | 33.6996062 | 12.3173879 | 29.0096503 |
| TCGA-FU-A3NI | 2 | 1243 | 2218 | 2369.47722 | 2549.15669 | 2808.06985 | 31990 | 15562.564 | 10126.0347 | 0 | 0 | 4.83369487 | 0 | 0.92461964 | 30.8098248 | 11.1464883 | 31.1420755 |
| TCGA-EK-A2RN | 2 | 25 | 2619 | 2383.97831 | 2720.73454 | 2793.27593 | 45109 | 14100.8547 | 10782.9328 | 0 | 0 | 0 | 3.92177952 | 0.92461964 | 31.1505451 | 10.1851424 | 29.3274552 |
| TCGA-LP-A4AX | 2 | 50 | 1863 | 2102.65727 | 2571.70692 | 2925.49654 | 24761 | 15792.6479 | 10144.6632 | 0 | 0 | 3.86695589 | 3.92177952 | 1.84923928 | 33.1291471 | 13.2508326 | 27.1112645 |
| TCGA-JW-A5VI | 2 | 830 | 2205 | 2278.60376 | 2568.76558 | 2798.82365 | 36084 | 16941.1338 | 10373.1068 | 0 | 0 | 0 | 7.84355903 | 0 | 32.4988407 | 12.7592863 | 29.5509095 |
| TCGA-EA-A3QE | 2 | 1122 | 2227 | 2115.22487 | 2415.81618 | 2865.39627 | 32611 | 12698.1164 | 9440.70374 | 0 | 0 | 1.93347795 | 0 | 2.77385892 | 34.5387542 | 13.5521398 | 25.9132046 |
| TCGA-C5-A7UE | 2 | 3935 | 2404 | 2466.15112 | 2520.72378 | 2702.66321 | 37066 | 17437.0709 | 9784.83989 | 0 | 0 | 4.83369487 | 0 | 0 | 34.2211921 | 11.2043461 | 29.1764954 |
| TCGA-Q1-A73Q | 2 | 602 | 2517 | 2380.11135 | 2767.79589 | 2938.44122 | 39120 | 15756.8785 | 10745.6759 | 0 | 0 | 1.93347795 | 7.84355903 | 2.77385892 | 33.3828125 | 9.01591102 | 31.4100199 |
| TCGA-JW-AAVH | 2 | 636 | 2042 | 1846.47144 | 2240.31655 | 2466.8852 | 30843 | 11640.504 | 8429.86507 | 0 | 0 | 2.90021692 | 7.84355903 | 0.92461964 | 32.9687883 | 16.6253853 | 29.9922459 |
| TCGA-MU-A8JM | 2 | 890 | 2209 | 2109.42444 | 2566.80469 | 2836.73306 | 34868 | 15946.3594 | 10122.1129 | 0 | 0 | 0 | 0 | 0.92461964 | 32.3779245 | 12.5278262 | 28.5778304 |
| TCGA-DS-A3LQ | 2 | 156 | 2502 | 2590.86045 | 2712.89098 | 3122.44053 | 38441 | 18178.5597 | 10775.0892 | 0 | 0 | 1.93347795 | 0 | 1.84923928 | 29.654886 | 8.74291742 | 32.6490991 |
| TCGA-EA-A1QS | 2 | 442 | 2462 | 2623.72957 | 2714.85187 | 2874.64246 | 37907 | 18240.4309 | 10764.3043 | 0 | 0 | 3.86695589 | 0 | 0.92461964 | 28.824711 | 9.23519656 | 35.1915358 |
| TCGA-VS-A958 | 2 | 596 | 2044 | 2213.83225 | 2282.47568 | 2926.42116 | 30475 | 14465.3153 | 8713.21364 | 0 | 0 | 3.86695589 | 3.92177952 | 2.77385892 | 33.771408 | 13.136828 | 25.7233242 |
| TCGA-DG-A2KM | 2 | 1066 | 2246 | 2077.52205 | 2406.99218 | 2522.36238 | 34076 | 12283.3854 | 9498.54999 | 0 | 0 | 0 | 0 | 0 | 34.1721494 | 12.8437659 | 25.39314 |
| TCGA-EK-A3GN | 2 | 1800 | 2462 | 2754.23933 | 2698.18431 | 3004.08921 | 38023 | 17612.0506 | 10796.659 | 0 | 0 | 1.93347795 | 3.92177952 | 1.84923928 | 30.1126453 | 9.27500486 | 35.3687802 |
| TCGA-DS-A1OC | 2 | 0 | 2144 | 2190.63051 | 2646.22073 | 2959.70747 | 34260 | 14924.5163 | 10452.5229 | 0 | 0 | 4.83369487 | 0 | 0.92461964 | 31.9238294 | 12.6320909 | 27.7852134 |
| TCGA-C5-A907 | 2 | 12 | 2341 | 2400.41287 | 2528.56734 | 2803.44675 | 37432 | 16809.6573 | 9914.25862 | 0 | 0 | 4.83369487 | 0 | 2.77385892 | 30.2970177 | 14.0387666 | 28.6221744 |
| TCGA-FU-A3HY | 2 | 491 | 2578 | 2296.00506 | 2519.74334 | 2963.40595 | 39975 | 13978.0788 | 9886.80616 | 0 | 0 | 1.93347795 | 3.92177952 | 0.92461964 | 29.2573437 | 11.927059 | 30.5284396 |
| TCGA-EA-A1QT | 2 | 595 | 2224 | 2426.51482 | 2257.96456 | 2093.33887 | 34104 | 14658.6631 | 8890.67416 | 0 | 0 | 4.83369487 | 3.92177952 | 0 | 30.7637399 | 11.4459995 | 30.1167946 |
| TCGA-DS-A0VN | 2 | 791 | 2276 | 2465.18438 | 2022.65779 | 2173.78077 | 35543 | 16124.2393 | 7972.97776 | 0 | 0 | 1.93347795 | 0 | 0 | 31.0773949 | 12.9382293 | 28.5417221 |
| TCGA-ZJ-AB0H | 2 | 919 | 2225 | 2564.7585 | 2422.6793 | 2443.76971 | 34173 | 16583.4403 | 9183.82718 | 0 | 0 | 3.86695589 | 3.92177952 | 0 | 33.1271483 | 10.8655268 | 28.5676139 |
| TCGA-EK-A2RB | 2 | 1013 | 2314 | 2353.04266 | 1957.94842 | 2245.90111 | 34116 | 14380.2422 | 7220.97653 | 0 | 0 | 1.93347795 | 3.92177952 | 0 | 31.9766216 | 14.4654772 | 29.1092843 |
| TCGA-EK-A2R8 | 2 | 4694 | 2377 | 2611.16197 | 2275.61256 | 2367.9509 | 38882 | 16567.0058 | 9048.52579 | 0 | 0 | 6.76717281 | 0 | 0.92461964 | 29.7655737 | 12.8879346 | 29.1224133 |
| TCGA-MA-AA43 | 2 | 473 | 2597 | 2748.4389 | 2594.25715 | 2350.38313 | 35860 | 18407.6768 | 10593.7069 | 0 | 0 | 4.83369487 | 0 | 0 | 29.5154597 | 7.98467648 | 33.0241152 |
| TCGA-MA-AA3Y | 2 | 534 | 2593 | 2496.12003 | 2306.00636 | 2087.79115 | 37405 | 16538.9704 | 8977.93376 | 0 | 0 | 6.76717281 | 0 | 0.92461964 | 32.1452041 | 11.9726078 | 27.8694664 |
| TCGA-BI-A0VS | 2 | 548 | 2179 | 2535.75633 | 2348.16549 | 2367.9509 | 33010 | 16922.7657 | 9112.25471 | 0 | 0 | 3.86695589 | 0 | 0 | 32.4113778 | 10.6289703 | 28.9948397 |
| TCGA-EK-A2IR | 2 | 1099 | 3201 | 3117.73319 | 3151.14984 | 3285.17358 | 46416 | 21164.8163 | 12770.2945 | 0 | 0 | 6.76717281 | 7.84355903 | 2.77385892 | 23.2912145 | 2.77812716 | 46.4219085 |
| TCGA-BI-A20A | 2 | 44 | 2033 | 2671.09978 | 2115.80005 | 2322.64454 | 32310 | 16617.2762 | 7921.01418 | 0 | 0 | 1.93347795 | 3.92177952 | 0.92461964 | 33.3024422 | 11.6365437 | 28.9344877 |
| TCGA-EA-A6QX | 2 | 380 | 2055 | 2430.38178 | 2245.21877 | 2151.5899 | 34118 | 17091.945 | 8858.31948 | 0 | 0 | 4.83369487 | 0 | 0.92461964 | 33.8052836 | 11.9611014 | 28.2507174 |
| TCGA-C5-A1MF | 2 | 747 | 2365 | 2510.62111 | 2379.53972 | 1989.78147 | 37252 | 15594.4664 | 9574.04424 | 0 | 0 | 4.83369487 | 0 | 0 | 33.5881198 | 12.4724596 | 27.4821656 |
| TCGA-IR-A3LH | 2 | 761 | 2176 | 2314.3731 | 2442.28819 | 2281.96127 | 31809 | 12124.8402 | 9360.30726 | 0 | 0 | 4.83369487 | 2.94133464 | 0 | 32.8755284 | 11.7344168 | 27.2680393 |
| TCGA-DS-A1OD | 2 | 4738 | 2123 | 2411.047 | 2219.72721 | 2426.20194 | 32031 | 14868.4454 | 8437.70863 | 0 | 0 | 1.93347795 | 0 | 0 | 34.6179362 | 13.540216 | 26.4174595 |
| TCGA-MA-AA3X | 2 | 284 | 2581 | 2719.43673 | 2499.154 | 2381.82019 | 39378 | 18994.4873 | 10058.384 | 0 | 0 | 4.83369487 | 3.92177952 | 0 | 29.5389521 | 9.08212814 | 30.667946 |
| TCGA-EA-A411 | 2 | 552 | 2398 | 2835.44541 | 2720.73454 | 2568.59336 | 32984 | 19679.9053 | 10677.0447 | 0 | 0 | 1.93347795 | 7.84355903 | 0.92461964 | 30.4540318 | 8.71900228 | 33.857652 |
| TCGA-EA-A439 | 2 | 699 | 2403 | 2497.08677 | 2194.23564 | 2451.16667 | 40526 | 17045.5416 | 8348.48814 | 0 | 0 | 2.90021692 | 3.92177952 | 0 | 28.203803 | 12.4696234 | 33.1773388 |
| TCGA-DG-A2KJ | 2 | 1525 | 2306 | 2195.46421 | 2331.49792 | 2451.16667 | 36057 | 13928.7751 | 9000.48399 | 0 | 0 | 1.93347795 | 3.92177952 | 0.92461964 | 34.7894742 | 12.4401359 | 27.704724 |
| TCGA-EK-A3GJ | 2 | 1946 | 2172 | 2299.87202 | 2095.21071 | 2124.77593 | 30698 | 14654.7961 | 7959.25153 | 0 | 0 | 4.83369487 | 0 | 0.92461964 | 31.4986709 | 13.4731807 | 26.2510629 |
| TCGA-ZJ-AAXU | 2 | 27 | 2015 | 2504.82068 | 2278.5539 | 2112.75588 | 33198 | 16201.5785 | 8805.37546 | 0 | 0 | 3.86695589 | 0 | 0.92461964 | 36.3500869 | 12.9715085 | 26.1572963 |
| TCGA-EK-A2RJ | 2 | 376 | 2310 | 2571.52567 | 2236.39477 | 2253.29806 | 34128 | 17685.5228 | 8595.56025 | 0 | 0 | 4.83369487 | 3.92177952 | 0.92461964 | 32.3422806 | 13.396594 | 27.075999 |

ID: Patients ID
PFS: Progress-free survival

Sato et al.

| | | | | | | | | | | | | | | | | | |
|---|---|---|---|---|---|---|---|---|---|---|---|---|---|---|---|---|---|
| TCGA-HM-A4S6 | 2 | 448 | 2192 | 2193.53073 | 2436.40552 | 2071.14799 | 38023 | 12799.624 | 9331.87436 | 0 | 0 | 2.90021692 | 7.84355903 | 0 | 31.3007129 | 12.7800991 | 28.578027 |
| TCGA-EA-A410 | 2 | 1243 | 2399 | 2652.73174 | 2082.46492 | 2415.1065 | 36945 | 17526.9776 | 8237.69787 | 0 | 0 | 3.86695589 | 3.92177952 | 0.92461964 | 29.071521 | 11.9098544 | 31.6306007 |
| TCGA-C5-A3HD | 2 | 3609 | 2271 | 2283.43745 | 2249.14055 | 2498.32227 | 32453 | 13721.893 | 8266.13077 | 0 | 0 | 3.86695589 | 0 | 0.92461964 | 30.1273679 | 13.4723583 | 28.1017581 |
| TCGA-DG-A2KK | 2 | 0 | 2227 | 2547.35719 | 2260.90589 | 2403.08645 | 32338 | 15142.0325 | 8488.69176 | 0 | 0 | 2.90021692 | 7.84355903 | 0 | 31.6233253 | 12.2572522 | 28.6773159 |
| TCGA-DS-A0VM | 2 | 9 | 2568 | 2238.96746 | 2407.97262 | 2531.60858 | 37523 | 12185.7448 | 9299.51968 | 0 | 0 | 0 | 3.92177952 | 0 | 29.3626823 | 11.4661283 | 31.3713841 |
| TCGA-LP-A4AW | 2 | 44 | 2326 | 2666.26609 | 2605.04204 | 2298.60443 | 38017 | 18269.4331 | 10486.8384 | 0 | 0 | 6.76717281 | 0 | 0 | 30.2371819 | 8.92584025 | 35.1154404 |
| TCGA-VS-A9U6 | 2 | 346 | 1931 | 2351.10918 | 2171.68541 | 2210.76556 | 25195 | 13743.1612 | 8353.39037 | 0 | 0 | 1.93347795 | 3.92177952 | 0 | 31.8331592 | 12.3083329 | 29.5387867 |
| TCGA-DG-A2KL | 2 | 542 | 2447 | 2607.29501 | 2445.22953 | 2440.99585 | 33839 | 17777.363 | 9628.94916 | 0 | 0 | 4.83369487 | 7.84355903 | 0.92461964 | 28.8506375 | 9.68510156 | 32.0371683 |
| TCGA-VS-A9UH | 2 | 1735 | 2662 | 2556.05785 | 2269.72989 | 2234.80567 | 38924 | 15379.8503 | 8823.02347 | 0 | 0 | 2.90021692 | 0 | 0 | 32.7819053 | 11.6649806 | 28.2541623 |
| TCGA-BI-A0VR | 2 | 720 | 2251 | 2459.38395 | 2344.24371 | 2329.11687 | 33941 | 17141.2487 | 9133.82449 | 0 | 0 | 4.83369487 | 7.84355903 | 0.92461964 | 33.2467312 | 12.5644056 | 27.9095203 |
| TCGA-DR-A0ZL | 2 | 730 | 2569 | 2564.7585 | 2532.48912 | 2505.71923 | 35491 | 16194.8113 | 9708.36519 | 0 | 0 | 4.83369487 | 7.84355903 | 1.84923928 | 33.4939372 | 8.73905988 | 30.4383147 |
| TCGA-ZJ-AAXT | 2 | 1617 | 2573 | 2562.82502 | 2517.78245 | 2279.18741 | 41907 | 15535.4953 | 10135.8392 | 0 | 0 | 0 | 3.92177952 | 0 | 31.1631379 | 11.0679009 | 28.9259546 |
| TCGA-EA-A78R | 2 | 2394 | 2258 | 2580.22632 | 2502.09533 | 2080.39419 | 37699 | 15977.295 | 9717.1892 | 0 | 0 | 4.83369487 | 0 | 0 | 31.3689589 | 10.82166 | 29.7843257 |
| TCGA-MA-AA3W | 2 | 3874 | 2209 | 2320.17354 | 2096.19115 | 2247.75035 | 35816 | 13039.3753 | 7778.84967 | 0 | 0 | 1.93347795 | 0 | 0.92461964 | 33.5414065 | 12.7696542 | 26.9260798 |
| TCGA-EA-A5ZE | 2 | 617 | 2376 | 2660.46565 | 2373.65705 | 2367.02628 | 31396 | 16169.6761 | 9117.15693 | 0 | 0 | 3.86695589 | 0 | 0 | 31.9588375 | 11.3940103 | 30.7194877 |
| TCGA-VS-A9UB | 2 | 965 | 2142 | 2420.71439 | 2192.27475 | 2152.51452 | 36849 | 16510.9349 | 8570.06869 | 0 | 0 | 2.90021692 | 3.92177952 | 1.84923928 | 32.7968615 | 12.8875121 | 25.3579156 |
| TCGA-C5-A2LX | 2 | 2893 | 2358 | 2485.4859 | 2324.63481 | 2220.01176 | 32286 | 17675.8554 | 8917.14617 | 0 | 0 | 6.76717281 | 0 | 0 | 34.62158 | 12.4441814 | 26.5687623 |
| TCGA-Q1-A6DT | 2 | 3 | 2561 | 2738.77151 | 2720.73454 | 2490.00069 | 40359 | 18574.9226 | 10815.2875 | 0 | 0 | 4.83369487 | 7.84355903 | 0 | 29.4948308 | 8.91088698 | 31.4312454 |
| TCGA-FU-A3TQ | 2 | 5 | 2457 | 2430.38178 | 2342.28282 | 2409.55878 | 37474 | 15181.6688 | 9258.34099 | 0 | 0 | 1.93347795 | 0 | 0 | 32.3806097 | 10.7501972 | 29.862241 |
| TCGA-VS-A8EC | 2 | 53 | 2533 | 2752.30586 | 2544.25446 | 2551.02559 | 36923 | 18351.6059 | 10532.9193 | 0 | 0 | 4.83369487 | 2.94133464 | 1.84923928 | 26.1605895 | 9.59113547 | 36.7279082 |
| TCGA-C5-A7UH | 2 | 454 | 2261 | 2511.58785 | 2182.4703 | 2342.06155 | 35187 | 18111.8547 | 8411.23662 | 0 | 0 | 4.83369487 | 3.92177952 | 0.92461964 | 31.4969394 | 11.2206568 | 29.878367 |
| TCGA-C5-A7CG | 2 | 1582 | 2320 | 2592.79393 | 2302.08458 | 1948.17358 | 35336 | 15002.8221 | 8679.87851 | 0 | 0 | 2.90021692 | 3.92177952 | 0.92461964 | 34.5056802 | 13.2953575 | 26.0813415 |
| TCGA-JW-A5VJ | 2 | 3589 | 2081 | 2565.72523 | 2464.83843 | 2372.574 | 29645 | 16403.6269 | 9582.86825 | 0 | 0 | 0 | 3.92177952 | 0.92461964 | 32.4265314 | 10.1298278 | 29.7162595 |
| TCGA-EA-A5ZF | 2 | 1320 | 2945 | 3023.95951 | 2900.15595 | 2751.66805 | 42847 | 21108.7455 | 11327.0797 | 0 | 0 | 0 | 3.92177952 | 0.92461964 | 25.5394516 | 6.13006197 | 38.4554336 |
| TCGA-VS-A8EH | 2 | 1427 | 2255 | 2404.27983 | 2357.96993 | 2223.71024 | 32342 | 14611.2928 | 9010.28844 | 0 | 0 | 2.90021692 | 3.92177952 | 0 | 30.5166704 | 13.2055483 | 30.6849308 |
| TCGA-EA-A3HR | 2 | 1505 | 2466 | 2485.4859 | 2307.96725 | 2459.48824 | 38885 | 16284.718 | 8698.50697 | 0 | 0 | 0 | 2.94133464 | 0.92461964 | 30.0610182 | 11.6174413 | 32.4312206 |
| TCGA-VS-A9U5 | 2 | 0 | 2334 | 2522.22198 | 2070.69958 | 2120.15284 | 35230 | 16078.8026 | 8100.43559 | 0 | 0 | 1.93347795 | 3.92177952 | 1.84923928 | 34.5942268 | 12.8430715 | 26.6515347 |
| TCGA-ZJ-AAX8 | 2 | 410 | 2500 | 2351.10918 | 2332.47837 | 2319.87068 | 42588 | 16404.5936 | 8647.52383 | 0 | 0 | 1.93347795 | 0 | 0.92461964 | 32.3865244 | 11.7928456 | 29.7040845 |
| TCGA-EK-A2R9 | 2 | 685 | 2302 | 2396.54591 | 2226.59032 | 2388.29253 | 36099 | 15404.9855 | 8811.25813 | 0 | 0 | 4.83369487 | 0 | 1.84923928 | 31.3516156 | 12.552279 | 30.1046714 |
| TCGA-EX-A1H5 | 2 | 829 | 2454 | 2098.79031 | 2451.1122 | 2137.72061 | 36747 | 11883.1555 | 9512.27622 | 0 | 0 | 6.76717281 | 3.92177952 | 0 | 32.8855893 | 12.0436068 | 31.8818674 |
| TCGA-EK-A2RO | 2 | 911 | 2651 | 2392.67896 | 2614.84649 | 2391.06639 | 38766 | 16450.0304 | 10479.9753 | 0 | 0 | 6.76717281 | 0 | 0 | 27.8572411 | 12.6826826 | 32.65741 |
| TCGA-EA-A3QD | 2 | 2526 | 2571 | 2207.06508 | 2286.39746 | 2465.03596 | 38282 | 13491.8091 | 9168.14006 | 0 | 0 | 2.90021692 | 3.92177952 | 0 | 33.8088375 | 12.0074794 | 27.6574062 |
| TCGA-FU-A23L | 2 | 795 | 2666 | 2235.10051 | 2689.3603 | 2335.58921 | 38054 | 16308.8865 | 10520.1736 | 0 | 0 | 1.93347795 | 0 | 2.77385892 | 33.1578019 | 13.4454086 | 27.3538454 |
| TCGA-MU-A5YI | 2 | 6408 | 2599 | 2615.02892 | 2470.7211 | 2196.89627 | 38089 | 16488.6999 | 9385.79883 | 0 | 0 | 4.83369487 | 7.84355903 | 0 | 30.6893084 | 11.0026724 | 33.3096214 |
| TCGA-IR-A3LL | 2 | 1535 | 2235 | 2065.92119 | 2454.05353 | 2004.57538 | 32215 | 14606.4591 | 9391.6815 | 0 | 0 | 1.93347795 | 3.92177952 | 2.77385892 | 33.0042504 | 13.505127 | 25.8550914 |
| TCGA-EA-A3HQ | 2 | 0 | 2538 | 2346.27549 | 2811.91591 | 2490.00069 | 38425 | 16693.6486 | 10931.9604 | 0 | 0 | 4.83369487 | 0 | 0 | 30.5766334 | 10.5974758 | 30.1553582 |
| TCGA-DR-A0ZM | 2 | 4 | 2542 | 1816.50253 | 2520.72378 | 2234.80567 | 38416 | 12222.4808 | 9867.19726 | 0 | 0 | 1.93347795 | 3.92177952 | 0 | 33.3879196 | 13.3482208 | 27.1263415 |
| TCGA-ZJ-AAXF | 2 | 619 | 2484 | 2288.27115 | 2522.68467 | 1984.23375 | 36982 | 17177.0181 | 9912.29773 | 0 | 0 | 6.76717281 | 5.88266927 | 0.92461964 | 34.3524192 | 11.5851797 | 28.9277376 |
| TCGA-EX-A69M | 2 | 2 | 2274 | 2194.49747 | 2223.64899 | 2214.46404 | 34681 | 15051.1591 | 8504.37888 | 0 | 0 | 1.93347795 | 3.92177952 | 0 | 32.773634 | 13.1266628 | 26.8062709 |
| TCGA-VS-A9V3 | 2 | 397 | 2585 | 2123.92552 | 2355.0286 | 2268.09198 | 39458 | 9853.97035 | 9542.67001 | 0 | 0 | 0 | 3.92177952 | 0.92461964 | 33.5248087 | 11.5716566 | 27.9536347 |
| TCGA-MY-A5BD | 2 | 725 | 2354 | 2251.53507 | 2194.23564 | 2055.42946 | 34286 | 13265.5922 | 8492.61354 | 0 | 0 | 4.83369487 | 3.92177952 | 0 | 32.526425 | 14.6884671 | 28.894062 |
| TCGA-LP-A4AV | 2 | 1053 | 2371 | 2352.07592 | 2440.3273 | 2414.18188 | 34521 | 15497.7925 | 9549.53312 | 0 | 0 | 2.90021692 | 7.84355903 | 0.92461964 | 33.4598557 | 12.116772 | 28.2132846 |
| TCGA-JX-A3Q0 | 2 | 1106 | 2653 | 2126.82574 | 2278.5539 | 2318.94606 | 36134 | 11236.4071 | 8724.97898 | 0 | 0 | 0 | 7.84355903 | 1.84923928 | 32.9675169 | 12.0608235 | 27.2608158 |
| TCGA-ZJ-AAXA | 2 | 1136 | 3024 | 2731.0376 | 3083.49914 | 2832.10996 | 44675 | 18110.8879 | 12577.1469 | 0 | 0 | 6.76717281 | 3.92177952 | 3.69847856 | 26.9979737 | 6.16298552 | 37.1818739 |
| TCGA-ZX-AA5X | 2 | 1791 | 2702 | 2224.46638 | 2756.03055 | 2415.1065 | 36291 | 14783.3724 | 10792.7372 | 0 | 0 | 1.93347795 | 5.88266927 | 0.92461964 | 31.0301247 | 10.2623337 | 31.8115422 |
| TCGA-EK-A2GZ | 2 | 0 | 2549 | 1956.67968 | 2283.45612 | 2302.3029 | 37884 | 11773.914 | 8665.17184 | 0 | 0 | 1.93347795 | 0 | 0 | 32.3911614 | 13.6415335 | 26.7520388 |
| TCGA-HM-A3JK | 2 | 186 | 2502 | 2542.5235 | 2483.46688 | 2433.59889 | 36736 | 17703.8908 | 9597.57492 | 0 | 0 | 0 | 3.92177952 | 0.92461964 | 31.1811818 | 9.9526276 | 33.134703 |
| TCGA-FU-A3HZ | 2 | 540 | 2481 | 1959.5799 | 2374.6375 | 2103.50968 | 40600 | 13889.1388 | 9728.95453 | 0 | 0 | 0 | 3.92177952 | 0 | 30.4161762 | 13.5574415 | 29.4809277 |
| TCGA-VS-A957 | 2 | 0 | 2740 | 2304.70571 | 2647.20117 | 2268.09198 | 39845 | 17432.2372 | 10318.2019 | 0 | 0 | 4.83369487 | 3.92177952 | 0.92461964 | 30.917076 | 13.3810536 | 28.6339184 |
| TCGA-ZJ-AAXI | 2 | 6375 | 2452 | 2268.93637 | 2169.72452 | 2090.56501 | 38544 | 14959.3189 | 8362.21437 | 0 | 0 | 4.83369487 | 0 | 0.92461964 | 32.6249567 | 13.7627745 | 28.0104221 |
| TCGA-EK-A2RK | 2 | 119 | 2477 | 2286.33767 | 2453.07309 | 2082.24343 | 35744 | 15513.2603 | 9434.82107 | 0 | 0 | 0 | 3.92177952 | 0 | 31.8359012 | 13.0254433 | 28.4348971 |
| TCGA-VS-A954 | 2 | 1688 | 2776 | 2304.70571 | 2648.18162 | 2305.07676 | 41616 | 13573.9819 | 10310.3583 | 0 | 0 | 6.76717281 | 0 | 0.92461964 | 32.1958552 | 11.270431 | 29.8624005 |
| TCGA-MY-A5BF | 2 | 0 | 2599 | 2376.2444 | 2847.21193 | 2448.39281 | 39329 | 15599.3001 | 11487.8726 | 0 | 0 | 6.76717281 | 7.84355903 | 0.92461964 | 31.0168613 | 9.47500152 | 34.1257238 |
| TCGA-EA-A3HT | 2 | 13 | 2635 | 2563.79176 | 2720.73454 | 2336.51383 | 39055 | 17497.9754 | 11141.7756 | 0 | 0 | 4.83369487 | 0 | 0.92461964 | 29.4806303 | 9.6430898 | 32.1943832 |
| TCGA-C5-A7CO | 2 | 1714 | 2455 | 2310.50615 | 2565.82425 | 2192.27317 | 32330 | 16231.5474 | 10050.5405 | 0 | 0 | 1.93347795 | 0 | 0 | 32.5280271 | 13.0960407 | 29.6824521 |
| TCGA-C5-A8XI | 2 | 4482 | 2459 | 2082.35575 | 2588.37448 | 2018.44467 | 34402 | 12620.7773 | 9890.72794 | 0 | 0 | 2.90021692 | 3.92177952 | 0 | 34.1733629 | 13.1930011 | 27.8847514 |
| TCGA-IR-A3LK | 2 | 254 | 2689 | 2172.26247 | 2671.7123 | 2216.31328 | 34790 | 14443.0803 | 10170.1547 | 0 | 0 | 1.93347795 | 0 | 0.92461964 | 30.8719543 | 12.2966326 | 27.2378287 |
| TCGA-VS-A9UD | 2 | 808 | 2630 | 2023.38467 | 2401.10951 | 2126.62517 | 36362 | 11835.7852 | 9330.89391 | 0 | 0 | 1.93347795 | 7.84355903 | 0 | 33.9221877 | 12.847476 | 25.3668532 |
| TCGA-EK-A2RA | 2 | 739 | 2571 | 2183.86334 | 2440.3273 | 1859.4101 | 39019 | 12575.3406 | 9308.34368 | 0 | 0 | 0 | 7.84355903 | 0 | 33.5673958 | 13.8992754 | 25.4112168 |
| TCGA-C5-A8XJ | 2 | 1246 | 2650 | 2630.49675 | 2575.6287 | 2351.30775 | 38309 | 16668.5134 | 10224.0792 | 0 | 0 | 1.93347795 | 3.92177952 | 1.84923928 | 31.1583291 | 12.3606109 | 32.9450634 |
| TCGA-MA-AA42 | 2 | 4467 | 2291 | 2187.7303 | 2321.69347 | 2132.17289 | 32829 | 13184.3861 | 8974.01198 | 0 | 0 | 2.90021692 | 0 | 0 | 34.7735105 | 12.9329663 | 25.2805999 |
| TCGA-DS-A1O9 | 2 | 259 | 2488 | 2343.37527 | 2548.17624 | 2027.69087 | 34475 | 15785.8807 | 10032.8924 | 0 | 0 | 4.83369487 | 3.92177952 | 0.92461964 | 30.4009783 | 12.1444875 | 29.0491779 |
| TCGA-C5-A3HL | 2 | 266 | 2575 | 2001.14967 | 2573.66781 | 2309.69986 | 36932 | 13265.5922 | 10220.1574 | 0 | 0 | 1.93347795 | 7.84355903 | 0 | 31.5542475 | 12.9593273 | 29.2376768 |
| TCGA-VS-A9UI | 2 | 1246 | 2284 | 2275.70354 | 2427.58152 | 2188.57469 | 30231 | 14874.2458 | 9582.86825 | 0 | 0 | 0 | 3.92177952 | 0 | 33.4141433 | 12.6564092 | 26.4093474 |
| TCGA-VS-A959 | 2 | 1561 | 2407 | 2367.54375 | 2567.78514 | 2305.07676 | 36702 | 16018.8648 | 9909.35639 | 0 | 0 | 0 | 0 | 1.84923928 | 34.3928358 | 11.2060843 | 29.0986188 |
| TCGA-DS-A1OA | 2 | 879 | 2481 | 2228.33333 | 2498.17355 | 2246.82573 | 38816 | 15574.1649 | 9994.6551 | 0 | 0 | 6.76717281 | 3.92177952 | 0 | 32.4092153 | 11.8847376 | 29.0035634 |
| TCGA-FU-A3TX | 2 | 1630 | 2504 | 2315.33984 | 2693.28208 | 2310.62448 | 38317 | 14339.6392 | 10514.2909 | 0 | 0 | 1.93347795 | 7.84355903 | 0.92461964 | 31.8479581 | 12.3238255 | 28.6728157 |
| TCGA-VS-A8Q9 | 2 | 512 | 2263 | 2246.70137 | 2451.1122 | 2126.62517 | 38637 | 14474.0159 | 9499.53043 | 0 | 0 | 4.83369487 | 0 | 0 | 32.0335976 | 14.0421097 | 29.486057 |
| TCGA-DS-A5RQ | 2 | 21 | 2558 | 2287.30441 | 2456.99487 | 2391.99101 | 38025 | 15639.9031 | 9658.3625 | 0 | 0 | 1.93347795 | 0 | 0.92461964 | 31.576337 | 11.1120227 | 29.6088214 |
| TCGA-EK-A2PM | 3 | 634 | 2706 | 2572.49241 | 2819.75947 | 2746.12033 | 42174 | 18148.5907 | 11361.3953 | 0 | 0 | 3.86695589 | 0 | 0.92461964 | 23.3080313 | 8.1789609 | 40.0306115 |
| TCGA-ZJ-AAX4 | 4 | 71 | 2604 | 2295.03832 | 2414.83574 | 2432.67427 | 37697 | 15318.9458 | 9188.72941 | 0 | 0 | 0 | 3.92177952 | 1.84923928 | 32.1141111 | 11.4638872 | 28.5147831 |

### Pancreatic cancer

| | | | Cell population size | | | | 2-6Sia levels | | | | Number of reproductive cells produced through multipolar cell division | | | | Supressive | Permissive | Lethal |
|---|---|---|---|---|---|---|---|---|---|---|---|---|---|---|---|---|---|
| ID | Best call | PFS | 2-6Sia 1.5 | 2-6Sia 1.0 | 2-6Sia 0.5 | 2-6Sia 0.25 | 2-6Sia 1.5 | 2-6Sia 1.0 | 2-6Sia 0.5 | 2-6Sia 0.25 | 2-6Sia 1.5 | 2-6Sia 1.0 | 2-6Sia 0.5 | 2-6Sia 0.25 | % | % | % |
| TCGA-IB-AAUU | 1 | 245 | 3966 | 4280.07692 | 4278.57016 | 4952.84353 | 277387 | 170021.25 | 160732.475 | 71111.1745 | 15 | 21.5769231 | 10.2726774 | 33.8673511 | 29.8901174 | 7.429147 | 30.0125763 |
| TCGA-2J-AABO | 1 | 440 | 4148 | 4265.36538 | 4550.79611 | 4807.11129 | 291324 | 181840.5 | 157545.89 | 68174.9778 | 11 | 19.6153846 | 5.13633872 | 27.7096509 | 31.2908171 | 8.58084107 | 31.10845 |
| TCGA-L1-A7W4 | 1 | 164 | 4208 | 4192.78846 | 4450.12387 | 4519.75195 | 276949 | 179931.923 | 171153.079 | 81257.0119 | 24 | 15.6923077 | 15.4090162 | 42.0776181 | 27.9938098 | 7.17828737 | 32.7332587 |
| TCGA-M8-A5N4 | 1 | 584 | 4296 | 3671.01923 | 4392.59688 | 5010.3154 | 303470 | 196921.788 | 163313.999 | 77246.2965 | 31 | 13.7307692 | 2.05453549 | 35.9199179 | 28.5543419 | 7.27650997 | 30.3261032 |
| TCGA-HV-AA8V | 1 | 920 | 4372 | 4391.88462 | 4382.3242 | 4868.6883 | 287217 | 159542.712 | 170615.818 | 75046.9713 | 26 | 25.5 | 10.2726774 | 28.7359343 | 29.8548804 | 7.89668903 | 31.2986427 |
| TCGA-HZ-A77Q | 1 | 33 | 4397 | 4285.96154 | 4917.53069 | 4747.58686 | 293022 | 163724.712 | 155540.664 | 76239.5125 | 11 | 0 | 16.4362839 | 33.8673511 | 32.2998154 | 8.50691476 | 28.8545926 |
| TCGA-2J-AABI | 1 | 969 | 4485 | 4029.98077 | 4609.35037 | 4785.55934 | 277901 | 167740.962 | 155065.039 | 70318.8838 | 22 | 1.96153846 | 14.3817484 | 23.6045175 | 34.1690069 | 8.51420687 | 27.6771755 |
| TCGA-XD-AAUL | 1 | 498 | 4492 | 4760.65385 | 4755.22239 | 4878.95113 | 262779 | 176055.923 | 159942.506 | 70803.2895 | 18 | 21.5769231 | 13.3544807 | 40.0250513 | 29.2903112 | 6.86685061 | 30.8975311 |
| TCGA-2J-AABP | 1 | 463 | 4530 | 4202.59615 | 4472.72376 | 4447.91211 | 276775 | 178841.308 | 152629.387 | 71877.8082 | 19 | 9.80769231 | 15.4090162 | 20.5256674 | 31.318564 | 10.1028273 | 28.921614 |
| TCGA-YB-A89D | 1 | 350 | 4554 | 4459.55769 | 4306.30639 | 4741.42916 | 298325 | 181633.558 | 138126.421 | 73013.9039 | 32 | 17.6538462 | 11.2999452 | 22.5782341 | 29.8305383 | 7.53912159 | 30.8687796 |
| TCGA-HZ-7922 | 1 | 4 | 4648 | 4092.75 | 4757.27693 | 4741.42916 | 298042 | 181093.154 | 156613.131 | 74745.2439 | 22 | 19.6153846 | 13.3544807 | 34.8936345 | 31.035349 | 7.69144336 | 30.7507192 |
| TCGA-US-A776 | 1 | 1216 | 4726 | 4464.46154 | 4054.62579 | 4892.29281 | 284385 | 180230.077 | 154093.243 | 66318.4312 | 29 | 24.5192308 | 7.19087421 | 28.7359343 | 31.5987249 | 9.39206504 | 34.1353507 |
| TCGA-2J-AABH | 1 | 1287 | 4690 | 4660.61538 | 4278.57016 | 4892.29281 | 312670 | 188660.769 | 177760.465 | 76227.1971 | 28 | 34.3269231 | 11.2999452 | 49.2616016 | 29.2315704 | 6.59312912 | 31.1589118 |
| TCGA-IB-A6UF | 1 | 666 | 5428 | 4160.42308 | 4563.12332 | 5141.67967 | 334552 | 169605.404 | 160758.157 | 76240.5388 | 19 | 15.6923077 | 8.21814196 | 32.8410678 | 27.2456221 | 6.60785809 | 32.7231853 |
| TCGA-Q3-AA2A | 1 | 95 | 4689 | 4683.17308 | 4911.36709 | 4409.93963 | 294739 | 190136.827 | 174134.21 | 68022.0616 | 16 | 34.3269231 | 26.7089614 | 26.6833676 | 29.9393965 | 8.19030256 | 30.6098214 |
| TCGA-LB-A8F3 | 1 | 379 | 5379 | 5243.19231 | 5205.16566 | 5074.97125 | 338426 | 208523.308 | 186485.05 | 73950.9006 | 32 | 19.6153846 | 42.1179775 | 42.0776181 | 27.7584812 | 7.21217478 | 32.5026282 |
| TCGA-IB-7644 | 1 | 347 | 4692 | 5013.69231 | 4670.98643 | 4302.17988 | 295044 | 203618.481 | 161125.918 | 65308.5684 | 21 | 22.5576923 | 32.8725678 | 32.8410678 | 29.7643342 | 8.93859297 | 30.9937712 |
| TCGA-HZ-7925 | 1 | 361 | 5153 | 4834.21154 | 5173.32036 | 4408.91335 | 323031 | 190136.827 | 177477.966 | 64549.1187 | 13 | 11.7692308 | 40.063442 | 40.0250513 | 28.4206848 | 7.03502723 | 31.3200334 |
| TCGA-XD-AAUG | 1 | 420 | 4939 | 4659.63462 | 4800.42217 | 4970.29035 | 311585 | 183728.481 | 172516.263 | 72731.676 | 34 | 11.7692308 | 34.9271033 | 34.8936345 | 30.9584206 | 7.32593935 | 32.3245763 |
| TCGA-XN-A8T3 | 1 | 951 | 5202 | 4789.09615 | 4609.35037 | 4229.31376 | 325537 | 193669.558 | 172178.292 | 63209.8189 | 32 | 13.7307692 | 13.3544807 | 13.3416838 | 29.7651402 | 9.857866 | 30.7405784 |
| TCGA-HZ-8317 | 1 | 16 | 5325 | 5301.05769 | 5737.29035 | 4461.2538 | 336881 | 213906.75 | 199163.589 | 66070.0706 | 32 | 14.7115385 | 29.7907646 | 29.7622177 | 27.8443932 | 7.08587353 | 32.8886698 |
| TCGA-HZ-A9TJ | 1 | 603 | 4639 | 4869.51923 | 4964.78501 | 4923.08131 | 286499 | 194727.808 | 176294.554 | 74493.8045 | 15 | 11.7692308 | 34.9271033 | 34.8936345 | 28.3946868 | 6.94197691 | 32.0377688 |
| TCGA-FB-A78T | 1 | 1 | 5325 | 5084.30769 | 5163.04768 | 4735.27146 | 331983 | 203786.192 | 181017.931 | 71350.2986 | 21 | 34.3269231 | 36.9816388 | 36.9462012 | 30.1612677 | 7.42229371 | 29.9730201 |
| TCGA-2J-AABE | 1 | 676 | 4973 | 4985.25 | 5325.35599 | 4818.40041 | 308775 | 199621.846 | 189088.146 | 72289.3478 | 15 | 30.4038462 | 35.9543711 | 35.9199179 | 30.182156 | 6.46222446 | 30.7336045 |
| TCGA-OE-A75W | 1 | 110 | 4727 | 4522.32692 | 4262.13387 | 4101.02834 | 295958 | 184255.154 | 154245.279 | 59774.8485 | 13 | 9.80769231 | 32.8725678 | 32.8410678 | 31.1593549 | 11.1648714 | 28.3093413 |
| TCGA-3A-A9J0 | 1 | 743 | 4738 | 4562.53846 | 4834.32201 | 4214.94579 | 297568 | 192476.942 | 171982.084 | 63314.4998 | 29 | 7.84615385 | 45.1997808 | 45.1564682 | 31.5397716 | 8.03543287 | 30.3063732 |
| TCGA-F2-A8YN | 1 | 517 | 5141 | 5359.90385 | 5200.02932 | 4947.71211 | 323038 | 215357.308 | 178837.042 | 76261.0645 | 25 | 17.6538462 | 24.6544259 | 24.6308008 | 28.8296627 | 7.31774978 | 31.9956323 |
| TCGA-HZ-A49I | 1 | 0 | 4729 | 4808.71154 | 4684.34092 | 3841.37864 | 295316 | 195450.635 | 160024.687 | 58643.8842 | 29 | 13.7307692 | 9.2454097 | 9.23655031 | 32.2603867 | 7.89534753 | 28.7107079 |
| TCGA-HZ-A77O | 1 | 11 | 4704 | 5046.05769 | 4774.74048 | 4509.48912 | 291427 | 205962.519 | 168359.938 | 67096.354 | 25 | 11.7692308 | 10.2726774 | 10.2628337 | 29.8608622 | 7.92647394 | 29.8488862 |
| TCGA-HZ-7289 | 1 | 240 | 5188 | 4893.05769 | 4744.94971 | 4048.68789 | 329142 | 189654.288 | 169942.958 | 61779.1799 | 21 | 34.3269231 | 22.5998904 | 22.5782341 | 29.1869485 | 10.7833755 | 30.633683 |
| TCGA-F2-A44G | 1 | 153 | 5117 | 5360.88462 | 5017.17566 | 5207.36181 | 316777 | 212228.654 | 180071.817 | 78199.7138 | 27 | 31.3846154 | 40.063442 | 40.0250513 | 28.1591343 | 6.32485377 | 31.2490918 |
| TCGA-HZ-8638 | 1 | 91 | 4786 | 4743 | 4478.88737 | 3986.0846 | 305121 | 189173.712 | 158638.903 | 59674.2727 | 26 | 24.5192308 | 19.5180871 | 19.499384 | 31.3332669 | 10.4517207 | 28.9754049 |
| TCGA-FB-AAQ6 | 1 | NA | 4293 | 4965.63462 | 4087.49836 | 4695.24641 | 274022 | 199567.904 | 147568.039 | 70373.2768 | 14 | 14.7115385 | 8.21814196 | 38.998768 | 30.7738981 | 8.23309193 | 32.3610453 |
| TCGA-IB-7890 | 1 | NA | 5141 | 5130.40385 | 4276.51562 | 4818.40041 | 326296 | 199842.519 | 154038.798 | 74936.1326 | 19 | 22.5576923 | 17.4635517 | 44.1301848 | 29.1321963 | 6.85718265 | 32.5544692 |
| TCGA-IB-A5SQ | 1 | NA | 4783 | 4821.46154 | 4384.37873 | 4395.57166 | 297204 | 196952.192 | 156196.061 | 67652.5996 | 41 | 16.6730769 | 13.3544807 | 21.5519507 | 31.371112 | 7.64763483 | 30.1880526 |
| TCGA-IB-AAUN | 1 | NA | 4513 | 4543.90385 | 4282.67923 | 5276.12279 | 282875 | 178785.404 | 154877.049 | 79213.6817 | 28 | 29.4230769 | 15.4090162 | 44.1301848 | 30.897942 | 8.0554286 | 30.3706688 |
| TCGA-FB-AAPU | 1 | NA | 4587 | 5117.65385 | 4563.12332 | 4984.65832 | 286530 | 201519.635 | 165055.218 | 74023.7667 | 22 | 33.3461538 | 19.5180871 | 35.9199179 | 32.4697597 | 7.29486894 | 31.8803046 |
| TCGA-IB-7893 | 1 | NA | 4655 | 4596.86538 | 4652.49562 | 5309.99014 | 294495 | 184164.923 | 169515.614 | 78886.2973 | 22 | 5.88461538 | 16.4362839 | 42.0776181 | 28.661101 | 7.50107375 | 31.7904658 |
| TCGA-2J-AABA | 1 | NA | 4678 | 4810.67308 | 4209.74322 | 4838.92608 | 295966 | 191565.808 | 152571.86 | 71245.6177 | 43 | 28.4423077 | 16.4362839 | 36.9462012 | 32.4477626 | 8.68093805 | 31.8172496 |
| TCGA-2J-AABU | 1 | NA | 4539 | 4774.38462 | 4227.20677 | 4775.29651 | 285558 | 191470.673 | 156165.243 | 70574.4283 | 28 | 26.4807692 | 15.4090162 | 12.3154004 | 32.1302205 | 9.6773299 | 30.279631 |
| TCGA-2L-AAQA | 1 | NA | 4665 | 5133.34615 | 4252.88846 | 5632.24312 | 297960 | 200577.115 | 154806.167 | 83997.1885 | 32 | 19.6153846 | 11.2999452 | 46.1827515 | 28.8192047 | 7.15244269 | 31.7757009 |
| TCGA-2L-AAQI | 1 | NA | 4390 | 4509.57692 | 3798.83612 | 4790.69076 | 277345 | 174378.808 | 136846.445 | 72843.5409 | 18 | 29.4230769 | 0 | 36.9462012 | 29.4981672 | 7.86537024 | 31.5789907 |
| TCGA-2L-AAQL | 1 | NA | 4490 | 5298.11538 | 4088.52562 | 5065.7347 | 277471 | 211467.577 | 143342.887 | 75504.6936 | 41 | 18.6346154 | 11.2999452 | 43.1039014 | 30.4724141 | 9.54242125 | 29.9584361 |
| TCGA-3A-A9IB | 1 | NA | 4520 | 5292.23077 | 4754.19512 | 4938.47556 | 283239 | 211378.327 | 170510.009 | 73905.7441 | 22 | 36.2884615 | 20.5453549 | 34.8936345 | 28.2465228 | 7.49746232 | 32.3103056 |
| TCGA-3A-A9IC | 1 | NA | 4521 | 5028.40385 | 4469.64196 | 4989.78973 | 284695 | 199800.346 | 160464.358 | 75734.5811 | 27 | 18.6346154 | 16.4362839 | 37.9724846 | 29.2110201 | 8.66822905 | 31.1777311 |
| TCGA-3A-A9IZ | 1 | NA | 4496 | 5321.65385 | 4753.16785 | 4711.66694 | 284220 | 212713.154 | 167847.331 | 72501.7885 | 15 | 28.4423077 | 20.5453549 | 21.5519507 | 27.940225 | 8.64379725 | 30.91039 |
| TCGA-FB-AAPP | 1 | NA | 4331 | 4610.59615 | 4304.25185 | 5032.89363 | 270272 | 182656.5 | 157610.608 | 74615.9322 | 27 | 9.80769231 | 3.08180323 | 35.9199179 | 28.767492 | 10.0135215 | 33.6882539 |
| TCGA-FB-AAPQ | 1 | NA | 4494 | 4923.46154 | 4524.08715 | 5039.05133 | 283364 | 200369.192 | 158719.03 | 72417.6333 | 17 | 25.5 | 23.6271581 | 50.287885 | 29.7612634 | 8.2832864 | 31.0736836 |
| TCGA-FB-AAQ2 | 1 | NA | 4450 | 4929.34615 | 4129.61633 | 4903.58193 | 280941 | 193701.923 | 147661.52 | 73928.3224 | 26 | 8.82692308 | 12.3272129 | 32.8410678 | 32.2287036 | 7.3183346 | 31.0087086 |
| TCGA-HZ-8005 | 1 | NA | 4351 | 4577.25 | 4845.62195 | 4844.05749 | 278954 | 183167.481 | 168887.954 | 73386.4448 | 20 | 24.5192308 | 16.4362839 | 41.0513347 | 29.2055908 | 9.90535508 | 29.0378345 |
| TCGA-IB-7654 | 1 | NA | 4820 | 4792.03846 | 4397.73321 | 4970.29035 | 303532 | 192261.173 | 161295.418 | 73194.5298 | 35 | 21.5769231 | 16.4362839 | 38.998768 | 29.9719103 | 7.68210619 | 32.561211 |
| TCGA-IB-7887 | 1 | NA | 4335 | 4472.30769 | 4504.56906 | 4881.0037 | 276201 | 181606.096 | 160701.657 | 73938.5852 | 37 | 16.6730769 | 9.2454097 | 29.7622177 | 30.7172868 | 9.88578572 | 29.4839193 |
| TCGA-IB-A5SS | 1 | NA | 4766 | 4633.15385 | 4305.27912 | 5180.67844 | 300847 | 188084.077 | 157067.184 | 77862.0665 | 19 | 18.6346154 | 14.3817484 | 29.7622177 | 29.0700392 | 8.24073496 | 30.5729533 |
| TCGA-IB-A7LX | 1 | NA | 4537 | 4743.98077 | 4139.88901 | 4856.3729 | 284461 | 198973.558 | 150216.335 | 70283.9901 | 35 | 22.5576923 | 19.5180871 | 51.3141684 | 30.2763498 | 9.01840833 | 30.3325614 |
| TCGA-US-A779 | 1 | NA | 5079 | 4976.42308 | 5309.94697 | 5250.46571 | 319269 | 193907.885 | 191524.826 | 75883.3922 | 29 | 13.7307692 | 14.3817484 | 46.1827515 | 29.4935461 | 5.94956219 | 33.3640877 |
| TCGA-FB-A7DR | 2 | 166 | 4364 | 4873.44231 | 4168.65251 | 5305.88501 | 277752 | 190962.635 | 161017.028 | 67891.7236 | 20 | 20.5961538 | 15.4090162 | 32.8410678 | 28.7974898 | 8.51656605 | 31.755495 |
| TCGA-2J-AAB4 | 2 | 729 | 4274 | 4651.78846 | 4496.35092 | 4993.89487 | 294089 | 163332.404 | 160093.514 | 77939.0378 | 18 | 15.6923077 | 18.4908194 | 27.7096509 | 30.3381646 | 9.09809762 | 29.4444435 |
| TCGA-HZ-7919 | 2 | 20 | 4374 | 4240.84615 | 4251.86119 | 4628.53799 | 259397 | 180803.827 | 159052.892 | 77852.83 | 16 | 14.7115385 | 16.4362839 | 23.6045175 | 30.7189213 | 9.37816408 | 30.0135652 |
| TCGA-IB-8127 | 2 | 522 | 4603 | 4540.96154 | 4493.26911 | 4705.50924 | 286657 | 168951.231 | 155954.653 | 68714.8029 | 20 | 21.5769231 | 10.2726774 | 19.499384 | 31.4665127 | 9.1925499 | 29.0805106 |
| TCGA-YY-A8LH | 2 | 2016 | 4822 | 4205.53846 | 3924.16278 | 4912.81848 | 275395 | 161518.962 | 160425.322 | 65532.2982 | 33 | 23.5384615 | 10.2726774 | 21.5519507 | 33.08638 | 9.1040978 | 29.4601109 |
| TCGA-HV-A5A6 | 2 | 1953 | 5104 | 4254.57692 | 5018.20293 | 4477.67433 | 318426 | 172605.577 | 176250.381 | 66495.9782 | 41 | 21.5769231 | 44.172513 | 44.1301848 | 27.7171649 | 8.1740659 | 31.9322759 |
| TCGA-3A-A9IH | 2 | 1021 | 4755 | 4637.07692 | 5303.78337 | 4747.58686 | 300216 | 189964.212 | 188674.158 | 72699.8612 | 20 | 19.6153846 | 22.5998904 | 22.5782341 | 29.9265584 | 7.01337159 | 31.6251498 |
| TCGA-3E-AAAY | 2 | 2285 | 4708 | 4838.13462 | 5012.03933 | 4382.22998 | 294593 | 189236.481 | 174953.97 | 64321.2838 | 13 | 16.6730769 | 27.7362291 | 27.7096509 | 30.6830943 | 8.99924838 | 29.8344833 |
| TCGA-HV-A7OL | 2 | 252 | 5187 | 4170.23077 | 5074.70266 | 4279.60164 | 325597 | 165666.635 | 179000.377 | 62459.6057 | 25 | 7.84615385 | 43.1452453 | 43.1039014 | 34.2506702 | 9.22452741 | 26.6123781 |
| TCGA-2J-AAB8 | 2 | 80 | 4992 | 5054.88462 | 4547.71431 | 4445.85955 | 317525 | 201770.712 | 160379.095 | 64962.7109 | 15 | 30.4038462 | 21.5726226 | 21.5519507 | 30.6760277 | 8.04606975 | 31.154618 |
| TCGA-IB-A5SO | 2 | 329 | 4780 | 4563.51923 | 4957.59414 | 4300.12731 | 300416 | 183609.808 | 175282.695 | 67006.0411 | 31 | 36.2884615 | 28.7634968 | 28.7359343 | 33.1455411 | 9.0158289 | 29.4846951 |
| TCGA-2J-AABK | 2 | 484 | 4376 | 4739.07692 | 5180.51124 | 4095.89692 | 273457 | 184984.846 | 180292.68 | 63786.5901 | 11 | 31.3846154 | 34.9271033 | 34.8936345 | 30.9757623 | 9.04744705 | 30.0065865 |
| TCGA-HV-A5A4 | 2 | 232 | 5010 | 4857.75 | 4839.45834 | 4301.15359 | 311594 | 193296.865 | 172665.217 | 64564.5129 | 22 | 34.3269231 | 13.3544807 | 13.3416838 | 30.2575896 | 8.87497969 | 30.6639896 |
| TCGA-FB-A545 | 2 | 385 | 5003 | 4829.30769 | 4808.64031 | 4078.4501 | 317610 | 201354.865 | 165841.077 | 64191.9721 | 24 | 15.6923077 | 18.4908194 | 18.4731006 | 28.6319173 | 7.70167418 | 35.4002255 |
| TCGA-LB-A7SX | 2 | 127 | 4557 | 4988.19231 | 5063.40271 | 4327.83696 | 284816 | 196527.519 | 179824.246 | 66514.4513 | 21 | 11.7692308 | 29.7907646 | 29.7622177 | 33.792597 | 8.90510921 | 28.8584768 |
| TCGA-FB-A5VM | 2 | 449 | 5075 | 4438.96154 | 4565.17786 | 4005.58398 | 319429 | 179056.096 | 161616.952 | 60980.7314 | 22 | 36.2884615 | 29.7907646 | 29.7622177 | 32.5653267 | 9.62306692 | 28.0463938 |
| TCGA-HZ-A4BH | 2 | 194 | 4951 | 4493.88462 | 4815.83119 | 4387.3614 | 307465 | 179490.577 | 172725.826 | 64998.6308 | 35 | 11.7692308 | 26.7089614 | 26.6833676 | 32.1418021 | 9.14464934 | 29.9266245 |
| TCGA-S4-A8RO | 2 | 525 | 5444 | 5132.36538 | 5090.11167 | 4988.76345 | 338122 | 198511.615 | 182049.308 | 74973.0789 | 30 | 18.6346154 | 44.172513 | 44.1301848 | 29.1202467 | 6.7097949 | 31.1411216 |
| TCGA-HZ-8637 | 2 | 0 | 4950 | 4871.48077 | 5139.42053 | 4169.78932 | 311350 | 196647.173 | 178166.236 | 65274.701 | 31 | 27.4615385 | 27.7362291 | 27.7096509 | 33.0219095 | 9.75290274 | 26.995335 |
| TCGA-IB-A7M4 | 2 | 483 | 4614 | 4282.03846 | 4283.70649 | 4071.26612 | 287400 | 169781.942 | 154338.76 | 61721.708 | 19 | 5.88461538 | 34.9271033 | 34.8936345 | 33.5558157 | 9.55118732 | 27.9778042 |
| TCGA-2J-AAB6 | 2 | NA | 4982 | 4952.88462 | 4734.67703 | 4863.55688 | 306905 | 200203.442 | 169158.125 | 70906.9441 | 35 | 30.4038462 | 9.2454097 | 42.0776181 | 28.9905561 | 6.6133539 | 31.0282532 |
| TCGA-FB-AAQ0 | 2 | NA | 4434 | 4453.67308 | 4326.85174 | 4757.84969 | 280007 | 177251.481 | 152411.606 | 68938.5326 | 12 | 16.6730769 | 17.4635517 | 22.5782341 | 28.8917618 | 8.68024449 | 33.9201314 |
| TCGA-2J-AABF | 2 | NA | 4650 | 5578.61538 | 4400.81502 | 4929.23901 | 291472 | 223736.019 | 153897.035 | 72071.7758 | 24 | 23.5384615 | 9.2454097 | 23.6045175 | 30.6304071 | 8.42281576 | 28.9338649 |
| TCGA-2L-AAQE | 2 | NA | 4898 | 4726.32692 | 4290.89737 | 4461.2538 | 306275 | 195947.885 | 154073.725 | 65703.6875 | 25 | 15.6923077 | 14.3817484 | 23.6045175 | 34.137865 | 8.96894719 | 30.1164919 |
| TCGA-3A-A9IU | 2 | NA | 4370 | 5292.23077 | 4431.63305 | 5001.07885 | 275484 | 206151.808 | 154302.806 | 73082.6649 | 24 | 23.5384615 | 9.2454097 | 31.8147844 | 30.4801116 | 7.64109082 | 30.7433688 |
| TCGA-F2-6879 | 2 | NA | 5210 | 4793.01923 | 5146.6114 | 5038.02505 | 329150 | 193378.269 | 186204.606 | 74159.2361 | 26 | 34.3269231 | 24.6544259 | 34.8936345 | 27.8333196 | 7.83959363 | 32.3307309 |
| TCGA-F2-A7TX | 2 | NA | 4000 | 5130.40385 | 4371.02425 | 4199.55154 | 254851 | 199481.596 | 153352.583 | 59808.7158 | 17 | 31.3846154 | 14.3817484 | 41.0513347 | 32.2598763 | 10.4563071 | 27.9066409 |
| TCGA-FB-AAQ3 | 2 | NA | 4626 | 4739.07692 | 4110.09825 | 4406.86078 | 297176 | 187446.577 | 149534.229 | 65640.0579 | 35 | 13.7307692 | 19.5180871 | 31.8147844 | 29.3849898 | 9.41125205 | 30.1364876 |
| TCGA-HV-AA8X | 2 | NA | 4678 | 5341.26923 | 4689.47725 | 5312.04271 | 294600 | 213327.115 | 166008.522 | 79589.3014 | 26 | 17.6538462 | 17.4635517 | 34.8936345 | 29.5999793 | 6.27558868 | 32.7282936 |
| TCGA-IB-7646 | 2 | NA | 4487 | 4623.34615 | 4829.18567 | 4662.40534 | 289310 | 180558.635 | 169737.504 | 70244.9914 | 24 | 36.2884615 | 12.3272129 | 36.9462012 | 28.2779565 | 8.33764218 | 30.2318238 |
| TCGA-IB-7651 | 2 | NA | 4400 | 4571.36538 | 4322.74267 | 5279.20164 | 278041 | 175821.519 | 159454.554 | 80141.4419 | 25 | 24.5192308 | 19.5180871 | 37.9724846 | 32.0033211 | 9.32765783 | 29.2618269 |
| TCGA-IB-7886 | 2 | NA | 4637 | 5198.07692 | 4202.55234 | 4813.26899 | 294015 | 204733.615 | 149420.203 | 73460.3372 | 24 | 17.6538462 | 16.4362839 | 21.5519507 | 29.8951239 | 9.31123517 | 29.7017614 |
| TCGA-IB-AAUO | 2 | NA | 4136 | 4979.36538 | 4539.49616 | 4638.80082 | 261269 | 200722.269 | 162725.374 | 70595.9803 | 29 | 23.5384615 | 13.3544807 | 26.6833676 | 30.6063496 | 10.0264508 | 27.8813988 |
| TCGA-IB-AAUQ | 2 | NA | 4790 | 4492.90385 | 4281.65196 | 4868.6883 | 301569 | 174871.154 | 157389.746 | 73572.2021 | 23 | 17.6538462 | 12.3272129 | 55.4193018 | 30.0864099 | 7.85892294 | 32.1016173 |
| TCGA-US-A77E | 2 | NA | 4909 | 5245.15385 | 4825.0766 | 5254.57084 | 303252 | 211528.385 | 170840.79 | 77617.8111 | 27 | 24.5192308 | 3.08180323 | 30.788501 | 29.0061455 | 5.40365381 | 33.5998977 |
| TCGA-US-A77G | 2 | NA | 4620 | 5145.11538 | 4638.11387 | 4976.44805 | 285170 | 206164.558 | 162797.283 | 74683.6669 | 15 | 19.6153846 | 23.6271581 | 26.6833676 | 29.5940467 | 7.40939524 | 32.1961636 |
| TCGA-3A-A9IL | 3 | 2741 | 4044 | 4442.88462 | 3920.05371 | 4498.2 | 282986 | 190798.846 | 170978.443 | 70924.391 | 27 | 15.6923077 | 5.13633872 | 40.0250513 | 28.1641139 | 7.76563055 | 34.8896889 |
| TCGA-3A-A9IX | 3 | 1037 | 4117 | 4300.67308 | 4237.47945 | 4864.58316 | 258243 | 182595.692 | 140364.837 | 68785.6164 | 21 | 13.7307692 | 14.3817484 | 10.2628337 | 33.2847552 | 9.1961448 | 29.2178762 |
| TCGA-RB-AA9M | 3 | 286 | 4132 | 4393.84615 | 4424.44218 | 5216.59836 | 288783 | 184430.712 | 155645.445 | 71147.0945 | 26 | 30.4038462 | 14.3817484 | 24.6308008 | 30.6283474 | 8.91575626 | 31.052811 |
| TCGA-IB-AAUV | 3 | 404 | 4318 | 4499.76923 | 3836.84503 | 4719.87721 | 270420 | 150222.462 | 156085.116 | 73842.1146 | 15 | 7.84615385 | 9.2454097 | 28.7359343 | 32.4726271 | 9.60606678 | 29.3683032 |
| TCGA-IB-7652 | 3 | 1116 | 4357 | 4029 | 4320.68813 | 4382.22998 | 300384 | 192007.154 | 152957.085 | 71728.9971 | 28 | 36.2884615 | 4.10907098 | 36.9462012 | 30.0843381 | 6.69870981 | 30.6845274 |
| TCGA-XD-AAUI | 3 | 202 | 4385 | 4455.63462 | 4001.20787 | 4784.53306 | 282901 | 161488.558 | 159211.091 | 71379.0345 | 26 | 23.5384615 | 17.4635517 | 30.788501 | 33.3703435 | 7.78707363 | 29.9285317 |
| TCGA-2L-AAQM | 3 | 1383 | 4452 | 4415.42308 | 4812.74938 | 5461.88008 | 275776 | 163720.788 | 154920.194 | 64453.6743 | 34 | 22.5576923 | 18.4908194 | 32.8410678 | 30.2648549 | 12.2748625 | 31.4158582 |
| TCGA-XN-A8T5 | 3 | 720 | 4436 | 4531.15385 | 4538.4689 | 4711.66694 | 303139 | 171761.135 | 136884.454 | 74230.0497 | 26 | 21.5769231 | 14.3817484 | 34.8936345 | 34.3952764 | 10.1269116 | 29.5284184 |
| TCGA-F2-7273 | 3 | 360 | 4388 | 4109.42308 | 4014.56235 | 4948.7384 | 269744 | 198859.788 | 153636.109 | 78960.1897 | 28 | 24.5192308 | 18.4908194 | 45.1564682 | 30.5976173 | 8.3895014 | 30.3720342 |
| TCGA-H6-8124 | 3 | 392 | 4533 | 3885.80769 | 4581.61414 | 4823.53183 | 295290 | 176427.635 | 142821.035 | 73680.9881 | 9 | 11.7692308 | 14.3817484 | 33.8673511 | 30.3771824 | 9.34852031 | 29.5956471 |
| TCGA-HZ-7926 | 3 | 8 | 4469 | 4447.78846 | 4095.7165 | 4433.54415 | 279933 | 168883.558 | 153277.593 | 71502.1885 | 13 | 29.4230769 | 8.21814196 | 32.8410678 | 33.1646877 | 8.63655308 | 29.0229475 |
| TCGA-XD-AAUH | 3 | 395 | 4529 | 4173.17308 | 4789.12223 | 4955.92238 | 281569 | 156575.885 | 168621.891 | 73256.1068 | 12 | 28.4423077 | 7.19087421 | 31.8147844 | 34.0193815 | 9.46406046 | 29.7386308 |
| TCGA-HZ-8636 | 3 | 5 | 4541 | 4415.42308 | 4272.40655 | 5009.28912 | 280819 | 173688.346 | 176204.154 | 68376.1294 | 33 | 12.75 | 19.5180871 | 22.5782341 | 29.7507448 | 7.40791017 | 30.2952021 |
| TCGA-HZ-A49G | 3 | 660 | 4569 | 3993.69231 | 4777.82228 | 5075.99754 | 279558 | 193152.692 | 162709.965 | 72923.591 | 10 | 13.7307692 | 15.4090162 | 27.7096509 | 30.6223643 | 8.34146536 | 29.8510453 |
| TCGA-3A-A9IV | 3 | 1103 | 4696 | 4024.09615 | 4310.41546 | 4927.18645 | 313352 | 197859.404 | 173755.148 | 74236.2074 | 9 | 16.6730769 | 14.3817484 | 18.4731006 | 26.349839 | 6.66053071 | 36.3325594 |
| TCGA-RL-AAAS | 3 | 9 | 4561 | 4046.65385 | 4538.4689 | 5220.70349 | 266216 | 168512.827 | 142942.252 | 67101.4854 | 23 | 9.80769231 | 9.2454097 | 21.5519507 | 33.4050114 | 10.2350437 | 28.7841734 |
| TCGA-2J-AABR | 3 | 438 | 4665 | 4248.69231 | 4035.1077 | 4945.65955 | 259386 | 175916.654 | 146654.798 | 71435.4801 | 23 | 9.80769231 | 18.4908194 | 47.2090349 | 32.069929 | 9.98705413 | 28.8189477 |
| TCGA-HZ-7924 | 3 | 840 | 4720 | 5043.11538 | 4149.13442 | 4573.11869 | 267060 | 170040.865 | 146770.879 | 70679.1092 | 18 | 13.7307692 | 7.19087421 | 43.1039014 | 32.877079 | 9.02004158 | 29.3847927 |
| TCGA-HZ-8002 | 3 | 24 | 4699 | 4437.98077 | 4459.36928 | 4920.00246 | 269487 | 173883.519 | 162211.74 | 70685.2669 | 19 | 20.5961538 | 14.3817484 | 35.9199179 | 30.9392826 | 9.27880473 | 30.1259306 |
| TCGA-3A-A9IN | 3 | 2084 | 4823 | 4496.82692 | 4308.36092 | 5328.46324 | 304832 | 183459.75 | 153838.481 | 78662.5676 | 23 | 12.75 | 10.2726774 | 43.1039014 | 28.8732176 | 8.3760525 | 34.6241698 |
| TCGA-IB-AAUW | 3 | 179 | 4922 | 4860.69231 | 4877.46725 | 4938.47556 | 295374 | 207734.769 | 146230.536 | 68337.1306 | 17 | 19.6153846 | 13.3544807 | 10.2628337 | 32.1355745 | 8.96370402 | 29.7736854 |
| TCGA-IB-A5ST | 3 | 635 | 4578 | 4775.36538 | 4978.13949 | 4619.30144 | 288670 | 183817.731 | 182634.851 | 69704.14 | 26 | 10.7884615 | 33.8998356 | 33.8673511 | 34.0570842 | 8.72022199 | 28.4557915 |
| TCGA-3A-A9IR | 3 | 1542 | 5123 | 4920.51923 | 4962.73047 | 4104.10719 | 322570 | 197142.462 | 182455.079 | 60962.2583 | 19 | 32.3653846 | 25.6816936 | 25.6570842 | 29.8844013 | 8.2030674 | 32.8174628 |
| TCGA-HZ-8003 | 3 | 21 | 4737 | 4777.32692 | 5203.11113 | 4331.94209 | 297509 | 195474.173 | 183107.394 | 66474.4263 | 20 | 24.5192308 | 34.9271033 | 34.8936345 | 32.0233946 | 9.28455222 | 28.4895575 |
| TCGA-IB-AAUT | 3 | 287 | 4570 | 4536.05769 | 5086.0026 | 4101.02834 | 286380 | 180699.865 | 181919.872 | 61898.2287 | 25 | 18.6346154 | 21.5726226 | 21.5519507 | 31.8281594 | 10.2500828 | 29.4043209 |
| TCGA-HZ-7923 | 3 | 314 | 4126 | 4976.42308 | 4491.21458 | 4326.81068 | 256325 | 198263.481 | 156150.861 | 65994.1257 | 3 | 13.7307692 | 17.4635517 | 17.4468172 | 32.1427509 | 9.27895524 | 29.113133 |
| TCGA-3A-A9IJ | 3 | 1854 | 5697 | 4965.63462 | 5247.28364 | 4801.97988 | 356745 | 195374.135 | 182197.234 | 73439.8115 | 13 | 26.4807692 | 49.3088517 | 49.2616016 | 27.2968537 | 5.97003407 | 33.8853766 |
| TCGA-S4-A8RM | 3 | 737 | 5017 | 4917.57692 | 4681.25911 | 4066.1347 | 315178 | 197230.731 | 167949.031 | 60963.2846 | 19 | 25.5 | 18.4908194 | 18.4731006 | 31.0952852 | 8.86045975 | 31.4312022 |
| TCGA-3A-A9IS | 3 | 998 | 5108 | 4835.19231 | 5410.61921 | 4220.07721 | 317952 | 194354.135 | 189083.01 | 63361.7088 | 16 | 20.5961538 | 43.1452453 | 43.1039014 | 28.5023821 | 7.98291137 | 32.8467872 |
| TCGA-IB-AAUP | 3 | 431 | 4433 | 4867.55769 | 4468.61469 | 4340.15236 | 275051 | 192162.115 | 159224.446 | 66856.2037 | 25 | 17.6538462 | 34.9271033 | 34.8936345 | 35.4755935 | 10.071797 | 27.852181 |
| TCGA-HZ-A49H | 3 | 491 | 4950 | 4255.55769 | 4369.99699 | 3951.19097 | 311471 | 169947.692 | 158831.002 | 59443.3589 | 30 | 13.7307692 | 20.5453549 | 20.5256674 | 32.2068558 | 9.86625685 | 28.6678543 |

ID: Patients ID
PFS: Progress-free surviva

Sato et al.

| | | | | | | | | | | | | | | | | |
|---|---|---|---|---|---|---|---|---|---|---|---|---|---|---|---|---|---|
| TCGA-F2-6880 | 3 | 295 | 4730 | 4468.38462 | 4954.51233 | 4807.11129 | 297744 | 182378.942 | 174633.462 | 72150.7996 | 29 | 19.6153846 | 39.0361743 | 38.998768 | 28.2588772 | 8.97184911 | 34.0628089 |
| TCGA-HZ-7920 | 3 | 0 | 4979 | 4886.19231 | 5002.79392 | 4056.89815 | 311758 | 191589.346 | 177868.328 | 61959.8057 | 36 | 22.5576923 | 29.7907646 | 29.7622177 | 32.4558275 | 9.47381399 | 29.3019602 |
| TCGA-2J-AABT | 3 | 319 | 4946 | 4670.42308 | 4685.36818 | 4725.00862 | 308531 | 187616.25 | 162266.186 | 72382.7396 | 2 | 11.7692308 | 30.8180323 | 30.788501 | 29.9341548 | 8.22476909 | 30.7270583 |
| TCGA-HZ-7918 | 3 | 969 | 4307 | 3759.28846 | 4917.53069 | 3947.08583 | 273238 | 159200.423 | 174476.29 | 58447.8641 | 9 | 16.6730769 | 10.2726774 | 10.2628337 | 32.380093 | 10.4148647 | 28.4528961 |
| TCGA-HZ-8519 | 3 | 454 | 4834 | 4998.98077 | 4511.75993 | 4515.64682 | 300011 | 199347.231 | 159269.646 | 66135.7528 | 38 | 23.5384615 | 16.4362839 | 16.4205339 | 30.3242005 | 9.69679733 | 30.9421835 |
| TCGA-HZ-A8P0 | 3 | 0 | 4604 | 4830.28846 | 5028.47561 | 4397.62423 | 287016 | 198390.981 | 177784.092 | 65240.8337 | 6 | 17.6538462 | 26.7089614 | 26.6833676 | 30.659223 | 7.76704262 | 31.6694159 |
| TCGA-IB-AAUM | 3 | 8 | 4577 | 4727.30769 | 4200.49781 | 4208.78809 | 290419 | 190867.5 | 155401.983 | 65092.0226 | 17 | 21.5769231 | 45.1997808 | 45.1564682 | 32.3571161 | 9.76217822 | 30.1413985 |
| TCGA-HZ-A77P | 3 | 330 | 5141 | 5241.23077 | 4914.44889 | 4358.62546 | 321573 | 204942.519 | 174870.761 | 64876.5031 | 32 | 28.4423077 | 26.7089614 | 26.6833676 | 29.2150331 | 8.87114039 | 31.7911536 |
| TCGA-IB-7885 | 3 | 1257 | 4373 | 4878.34615 | 4855.89463 | 4296.02218 | 278913 | 194199.173 | 175846.665 | 63456.1269 | 17 | 21.5769231 | 25.6816936 | 25.6570842 | 31.5727083 | 8.61958445 | 29.3946619 |
| TCGA-IB-8126 | 3 | 462 | 4584 | 4915.61538 | 4286.7883 | 4374.01971 | 286583 | 196870.788 | 151657.592 | 67177.4304 | 4 | 30.4038462 | 39.0361743 | 38.998768 | 32.3777843 | 10.8387703 | 29.2986999 |
| TCGA-HZ-A8P1 | 3 | 7 | 4900 | 5112.75 | 4968.89408 | 4653.16879 | 305410 | 204412.904 | 173615.44 | 67744.9651 | 26 | 32.3653846 | 35.9543711 | 35.9199179 | 29.1141828 | 7.40924234 | 33.065403 |
| TCGA-3A-A9I9 | 3 | NA | 4635 | 4679.25 | 4058.73486 | 4592.61807 | 293905 | 190585.038 | 146580.834 | 67558.1815 | 20 | 27.4615385 | 18.4908194 | 17.4468172 | 29.9378714 | 8.9806442 | 29.6339358 |
| TCGA-3E-AAAZ | 3 | NA | 4541 | 5144.13462 | 4888.7672 | 5069.83984 | 287989 | 206632.385 | 170457.619 | 76537.1347 | 20 | 14.7115385 | 12.3272129 | 33.8673511 | 30.1563732 | 7.46869319 | 32.8314378 |
| TCGA-HV-A5A3 | 3 | NA | 4231 | 4714.55769 | 4363.83338 | 4957.97495 | 269895 | 183748.096 | 155992.662 | 72963.616 | 24 | 26.4807692 | 20.5453549 | 29.7622177 | 30.9986762 | 9.27408494 | 29.0424623 |
| TCGA-IB-7888 | 3 | NA | 4230 | 4803.80769 | 4205.63415 | 4691.14127 | 267187 | 194616 | 152051.035 | 71689.9984 | 13 | 22.5576923 | 5.13633872 | 31.8147844 | 33.1190247 | 8.97804256 | 29.0469168 |
| TCGA-2J-AABV | 3 | NA | 3959 | 4928.36538 | 4705.91354 | 4453.04353 | 247432 | 195587.942 | 162851.728 | 65498.4308 | 15 | 25.5 | 16.4362839 | 31.8147844 | 31.10433 | 10.2132025 | 29.5414284 |
| TCGA-F2-7276 | 3 | NA | 4200 | 4686.11538 | 4423.41491 | 4626.48542 | 268028 | 186137.25 | 161021.137 | 68754.8279 | 30 | 13.7307692 | 7.19087421 | 47.2090349 | 31.9835869 | 9.00607179 | 29.4044412 |
| TCGA-IB-7645 | 3 | NA | 4305 | 4778.30769 | 4427.52398 | 4680.87844 | 276144 | 192091.5 | 157469.873 | 69976.1051 | 24 | 19.6153846 | 6.16360647 | 19.499384 | 31.6887847 | 8.75195198 | 29.6585994 |
| TCGA-IB-7649 | 3 | NA | 4747 | 4616.48077 | 3906.69923 | 5118.07515 | 302030 | 181421.712 | 136720.091 | 78231.5285 | 12 | 20.5961538 | 11.2999452 | 30.788501 | 31.6594698 | 9.24734434 | 29.6985741 |
| TCGA-IB-7889 | 3 | NA | 4042 | 4739.07692 | 4188.17059 | 4523.85708 | 256034 | 192015.981 | 149873.228 | 65542.561 | 29 | 15.6923077 | 9.2454097 | 29.7622177 | 32.924491 | 9.50363479 | 28.7003096 |
| TCGA-IB-7897 | 3 | NA | 4014 | 4925.42308 | 4110.09825 | 4242.65544 | 251483 | 197125.788 | 152707.459 | 65253.1491 | 7 | 34.3269231 | 11.2999452 | 12.3154004 | 31.061377 | 9.75343247 | 29.81038 |
| TCGA-IB-A6UG | 3 | NA | 4141 | 4316.36538 | 4051.54398 | 4924.1076 | 261497 | 174495.519 | 149013.404 | 74553.329 | 14 | 11.7692308 | 13.3544807 | 38.998768 | 33.4496953 | 8.67138133 | 29.581299 |
| TCGA-US-A77J | 3 | NA | 3945 | 4780.26923 | 4455.26021 | 4254.97084 | 253021 | 192129.75 | 157780.107 | 62546.8398 | 23 | 21.5769231 | 19.5180871 | 32.8410678 | 34.7355653 | 9.44736928 | 28.1865324 |
| TCGA-2L-AAQJ | 3 | NA | 4630 | 4887.17308 | 4563.12332 | 4970.29035 | 292930 | 199657.154 | 159899.361 | 75135.2316 | 13 | 28.4423077 | 10.2726774 | 12.3154004 | 32.5852179 | 8.1114855 | 30.5229389 |
| TCGA-HV-A7OP | 4 | 978 | 4509 | 4625.30769 | 4846.64922 | 4065.10842 | 285169 | 185424.231 | 171538.304 | 59632.1951 | 11 | 18.6346154 | 24.6544259 | 24.6308008 | 33.1977705 | 10.4307878 | 26.6055031 |
| TCGA-PZ-A5RE | 6 | 247 | 4143 | 4164.34615 | 3999.15333 | 4463.30637 | 298024 | 177824.25 | 166335.193 | 72406.3441 | 16 | 13.7307692 | 17.4635517 | 33.8673511 | 30.425719 | 7.61920258 | 29.0493854 |
| TCGA-3A-A9I7 | 6 | 1323 | 4247 | 4190.82692 | 4088.52562 | 4720.90349 | 276866 | 169425.923 | 155083.53 | 68632.7002 | 12 | 7.84615385 | 19.5180871 | 27.7096509 | 34.1375256 | 9.50141159 | 28.8268572 |
| TCGA-IB-7891 | 6 | 648 | 4293 | 4143.75 | 4396.70595 | 4564.90842 | 273383 | 174163.038 | 136623.528 | 72697.8086 | 8 | 29.4230769 | 8.21814196 | 25.6570842 | 31.7496018 | 8.99112211 | 29.3389731 |
| TCGA-H8-A6C1 | 6 | 671 | 4438 | 4304.59615 | 4435.74212 | 4920.00246 | 285509 | 171143.25 | 154059.344 | 72808.6472 | 25 | 11.7692308 | 16.4362839 | 32.8410678 | 31.7710496 | 7.15172132 | 30.1280345 |
| TCGA-HZ-8315 | 6 | 28 | 4496 | 4103.53846 | 4344.31529 | 4761.95483 | 295988 | 165841.212 | 171134.588 | 72856.8825 | 14 | 7.84615385 | 14.3817484 | 36.9462012 | 30.7820143 | 9.19260372 | 29.1771931 |
| TCGA-FB-AAPZ | 6 | 716 | 4468 | 4720.44231 | 4408.00589 | 4844.05749 | 276466 | 181445.25 | 142974.097 | 71760.8119 | 12 | 16.6730769 | 11.2999452 | 25.6570842 | 30.9226275 | 9.29628178 | 29.2865268 |
| TCGA-FB-A4P6 | 6 | 767 | 4417 | 4085.88462 | 4331.98808 | 4690.11499 | 254171 | 174312.115 | 150867.623 | 74763.717 | 8 | 16.6730769 | 12.3272129 | 38.998768 | 31.1351998 | 8.81989865 | 30.9027101 |
| TCGA-HV-A5A5 | 6 | 289 | 4655 | 4517.42308 | 4222.07043 | 5050.34045 | 284927 | 168929.654 | 175961.719 | 74295.7318 | 20 | 21.5769231 | 14.3817484 | 43.1039014 | 30.9788555 | 8.15633317 | 30.6824316 |
| TCGA-FB-AAPS | 6 | 228 | 4821 | 4720.44231 | 4608.3231 | 5319.22669 | 279272 | 171843.519 | 144565.335 | 73991.952 | 22 | 24.5192308 | 3.08180323 | 22.5782341 | 32.4615176 | 9.06794562 | 30.3984826 |
| TCGA-LB-A9Q5 | 6 | 155 | 4894 | 4725.34615 | 4824.04933 | 5143.73224 | 272146 | 189027.577 | 157989.67 | 73578.3598 | 26 | 19.6153846 | 15.4090162 | 31.8147844 | 29.8725791 | 8.07627609 | 29.2985666 |
| TCGA-F2-A44H | 6 | 586 | 4562 | 4902.86538 | 5222.62921 | 4628.53799 | 287850 | 196061.654 | 179530.447 | 71859.3351 | 24 | 17.6538462 | 15.4090162 | 15.3942505 | 28.8481355 | 7.24593842 | 31.2262032 |
| TCGA-IB-AAUR | 6 | 338 | 4344 | 4711.61538 | 4632.97753 | 3953.24353 | 271629 | 188971.673 | 168473.965 | 61532.8719 | 28 | 15.6923077 | 17.4635517 | 17.4468172 | 35.202393 | 10.1479839 | 27.8554845 |
| TCGA-FB-A4P5 | 6 | 4 | 4891 | 4432.09615 | 4203.57961 | 4143.10595 | 305900 | 177782.077 | 153161.512 | 63388.3922 | 31 | 19.6153846 | 22.5998904 | 22.5782341 | 34.0694579 | 10.3291373 | 28.382596 |
| TCGA-HZ-8001 | 6 | 706 | 5172 | 4701.80769 | 4734.67703 | 3862.9306 | 326321 | 190708.615 | 168439.038 | 60336.2255 | 33 | 19.6153846 | 24.6544259 | 24.6308008 | 32.7079779 | 9.35505294 | 28.5639432 |
| TCGA-HZ-A4BK | 6 | 657 | 5027 | 5101.96154 | 5004.84845 | 4220.07721 | 314617 | 208240.846 | 173851.711 | 64025.7142 | 31 | 20.5961538 | 14.3817484 | 14.3679671 | 31.8062431 | 8.88467884 | 29.6231218 |
| TCGA-IB-A5SP | 6 | 482 | 4900 | 4459.55769 | 4673.04097 | 4595.69692 | 306390 | 174926.077 | 160948.201 | 70361.9877 | 23 | 19.6153846 | 29.7907646 | 29.7622177 | 29.5088379 | 8.59477439 | 29.8908335 |
| TCGA-RB-A7B8 | 6 | 36 | 4783 | 4817.53846 | 4682.28638 | 4416.09733 | 304908 | 185050.558 | 168377.401 | 68106.2168 | 12 | 27.4615385 | 18.4908194 | 18.4731006 | 30.7295825 | 8.77653634 | 30.669068 |
| TCGA-IB-AAUS | 6 | 225 | 4942 | 4608.63462 | 4781.93135 | 5027.76222 | 309880 | 180995.077 | 171750.949 | 77441.2903 | 38 | 17.6538462 | 32.8725678 | 32.8410678 | 30.8280069 | 8.85842277 | 29.3036186 |
| TCGA-H6-A45N | 6 | 233 | 4637 | 4724.36538 | 4749.05878 | 4234.44517 | 294002 | 195009.288 | 171642.058 | 65274.701 | 23 | 18.6346154 | 21.5726226 | 21.5519507 | 31.8285282 | 9.54114626 | 29.7430061 |
| TCGA-2J-AAB1 | 6 | NA | 4415 | 4875.40385 | 3909.78104 | 4892.29281 | 282500 | 192581.885 | 140847.653 | 75372.3031 | 41 | 9.80769231 | 16.4362839 | 36.9462012 | 31.2413315 | 9.30211249 | 30.0133349 |
| TCGA-2J-AAB9 | 6 | NA | 4527 | 4466.42308 | 4411.0877 | 4921.02875 | 289181 | 174595.558 | 160404.777 | 71003.4148 | 30 | 17.6538462 | 7.19087421 | 44.1301848 | 30.9858633 | 8.82215179 | 30.6480145 |
| TCGA-FB-AAPY | 6 | NA | 4318 | 4616.48077 | 3522.5011 | 4805.05873 | 269908 | 185798.885 | 121765.128 | 70423.5647 | 11 | 10.7884615 | 2.05453549 | 32.8410678 | 33.2697526 | 9.01222526 | 28.4966611 |
| TCGA-FB-AAQ1 | 6 | NA | 4926 | 4717.5 | 4704.88627 | 4744.50801 | 311563 | 192527.942 | 165331.553 | 71092.7014 | 31 | 25.5 | 5.13633872 | 29.7622177 | 30.0085328 | 7.79114044 | 30.2432463 |
| TCGA-S4-A8RP | 6 | NA | 4593 | 4835.19231 | 3900.53563 | 4514.62053 | 291175 | 199814.077 | 140686.372 | 65124.8637 | 24 | 17.6538462 | 9.2454097 | 13.3416838 | 29.6557348 | 8.81995889 | 30.5865829 |
| TCGA-US-A774 | 6 | NA | 4194 | 4403.65385 | 4094.68923 | 5074.97125 | 263515 | 175134 | 146535.635 | 77367.3979 | 12 | 7.84615385 | 13.3544807 | 57.4718686 | 32.1718919 | 9.04687416 | 28.9279785 |
