## Supplementary material for "Cell Fate Simulation Reveals Cancer Cell Features in the Tumor Microenvironment": Legends for Supplementary Videos-Figs

### Legends for Supplementary Videos, Figs and Data

**Supplementary Video 1. Single-cell tracking of HeLa cell.**

A video for single-cell tracking of HeLa cells is shown (time points 1–693). A unique color was assigned to each cell lineage. The total number of tracked cells was 11,640.

**Supplementary Video 2. Single-ell tracking of MiaPaCa2 cell.**

A video for single-cell tracking of MiaPaCa2 cells is shown (time points 1–597). A unique color was assigned to each cell lineage. The total number of tracked cells was 6,987.

**Supplementary Fig. 1. Computerized single-cell lineage tracking analysis system.**

The computerized single-cell lineage tracking analysis system included Computer 1 to control the microscope and Computer 2 to perform image processing, single-cell tracking, and data analysis. Commercially available image acquisition software (Metamorph) was installed on Computer 1 to control the microscope and create image files. Computer 2 employed various custom software programs; *Image Processing Controller* controlled other software programs; *File Transfer* and *Map* communicated between Computers 1 and 2; *File Converter* imported images created by other microscopes; *Name Assignment*, *Focal Image Selection*, and *Contrast Set* produced live cell videos; *Data Backup* controlled file archiving and backup; *Outline Drawing* performed image segmentation and created the object segmentation library; *Movie Viewer* played movies and fine-tuned image quality; *Object Tracking Controller* created the cell lineage database by controlling *Automatic Object Tracking*, and facilitated data verification; and

*Data Analysis* provided various options for data analysis. DIC/FL; differential interference contrast/fluorescence. FOV; field of view.

**Supplementary Fig. 2. Image segmentation.**

**A.** Image segmentation was performed using the Stepwise Area Expansion method. Pixels with values exceeding predefined thresholds (THs) ranging from TH1 to TH4 were extracted from the NIR-DIC image and assigned a grayscale value of 150 (ranging from 0 to 255 grayscale, such as 200, 170, 140, and 110 for TH1, 2, 3, and 4, respectively). The segmented results are displayed, with areas enclosed by blue circles representing the segmented regions. **B.** Connectivity analysis was conducted using TH1 to identify connected pixel edges, which were marked with circles (referred to as 'edge circles,' as shown in **a**: Edge circles, indicated by pink lines). These circles were overlaid onto the NIR-DIC image, and lines were extended from pixels within these regions toward the 12, 3, 6, and 9 o'clock directions (**b**: Line extension, depicted in red, yellow, light blue, and blue regions, respectively, along with a magnified view of Line extension). The endpoints of each line were linked (**c**: Edge linking, and a magnified view of Edge linking), creating new edge circles by connecting the linked lines (**d**: Creation of new areas, marked by green lines). These newly formed edge circles were then overlaid onto the TH2 image (**e**: represented by green circles), and pixels that did not overlap with green circles were identified (**f**: indicated by pink circles). The processing of pink circles followed a similar procedure as for TH1 (**f-h**). The green circles were expanded by 1–4 pixels outside the edge circles (**k** and **l**, along with an enlarged view of Edge expansion). Finally, the green and red edge circles were overlaid to detect overlaps (**i**: Overlay), which were subsequently eliminated (**j**: Removal of overlaps). These processes were replicated for the TH3 and TH4 images.

**Supplementary Fig. 3. Object tracking (single-cell tracking).**

**A.** At Time A, the segmented areas identified as cell representations are denoted by orange characters (cell lineage numbers). The cell being tracked is highlighted by a blue area with a white asterisk. Additionally, the yellow and magenta areas represent segmented regions corresponding to neighboring cells. **B.** At Time A+1, there were changes in the positions of cells and segmentation patterns (indicated by black characters representing segmented area numbers). **C.** The blue area from Time A was overlaid onto the Time A+1 image. However, the blue segmented area observed at Time A was not present in Time A+1; instead, it was partially overlaid with the light blue and white segmented areas. **D.** To identify which segmented area corresponds to the tracked cell, the yellow and magenta areas were overlaid on the Time A+1 image. This process revealed that the light blue area was associated with the yellow area. Furthermore, the magenta area overlapped with the white area, with the larger portion of the white area overlapping with the blue area. **E.** Consequently, it was determined that the white area represents the segmented region corresponding to the tracked cell. This systematic process was repeated for all cells recorded in the database.

**Supplementary Fig. 4. Outline of the Generation of Deduced Cell Populations.**

**A.** A list containing the 2-6Sia expression levels of all analyzed HeLa or MiaPaCa2 cells obtained through single-cell tracking was compiled. This list served as a reference when assigning 2-6Sia expression levels to deduced cells. Subsequently, a progenitor cell was created by assigning the length of time until a cellular event occurred and the type of that event using the cell fate simulation algorithm. In the case of bipolar cell division, two daughter cells were generated, and each was assigned a 2-6Sia expression level using the 2-6Sia expression level list. These expression levels were later modified to introduce variations in 2-6Sia expression levels. **B.** The initial step in generating the deduced cell population involved creating a list of 2-6Sia expression levels. SNA1 binding levels of individual cells, measured at the end of the live cell, are displayed on the cell lineage map using a heatmap scale, where blue to red corresponds to low to high expression. Based on the 2-6Sia expression levels of cells, we traced back along the cell lineage map. Subsequently, a list of 2-6Sia expression levels for all tracked HeLa or MiaPaCa2 cells was compiled. **C**. The deduced cell population was generated using a cell fate simulation algorithm. The algorithm assigned the length of time for the First event (the event that occurred in the progenitor cell) to the progenitors. Then, a cellular event was assigned to a progenitor cell. If a bipolar cell division was assigned, two daughter cells were created. In Example 1, a longer time interval until the Next event was assigned compared to Example 2. After the assignment, the algorithm referred to the cell doubling time (the time between bipolar cell divisions) of 2-6Sia-expressing cells. It checked whether any of the 2-6Sia-expressing cells had cell doubling times within $\pm$10% of the time assigned to daughter cells. If no such cells were found, 2-6Sia expression levels were not assigned to the daughter cells (Example 1). If some cells met this criterion, 2-6Sia expression levels were assigned to the daughter cells (Example 2). **D.** The algorithm assigned 2-6Sia expression levels to the daughter cells by referencing the 2-6Sia expression level list. A random value was generated to select one value from the list, which was then assigned to one of the daughter cells, followed by assignment to the second daughter cell. Finally, 2-6Sia expression levels were modified to create deduced cell populations with various levels of 2-6Sia expression. For additional details, see the Supplemental Materials and Methods.

**Supplementary Fig. 5. Validation 3D TME simulations.**

**A-H**: Validation simulations of the 3D TME were conducted by varying the ratio of cancer cells to Suppressive, Permissive, or Lethal cells. **A-C**: Suppressive, Permissive, or Lethal cells were introduced alongside Cervical 2-6Sia 1.0 cells in the 3D TME. (**A**) Suppressive cells, (**B**) Permissive cells, and (**C**) Lethal cells. Simulations were performed by changing the ratio from 1:0 to 1:0.9 for Suppressive and Permissive cells and from 1:0 to 1:0.46 for Lethal cells. **D**: Both Suppressive and Permissive cells were placed in the 3D TME, with a cancer cell to Suppressive cell ratio of 1:0.2, and the ratio of Permissive cells changed from 1:0 to 1:0.5. **E-H**: Suppressive, Permissive, or Lethal cells were introduced alongside Pancreatic 2-6Sia 1.0 cells in the 3D TME. (**E**) Suppressive cells, (**F**) Permissive cells, and (**G**) Lethal cells. Simulations were performed by changing the ratio from 1:0 to 1:0.9 for Suppressive and Permissive cells and from 1:0 to 1:0.46 for Lethal cells. **H**: Both Suppressive and Permissive cells were placed in the 3D TME, with a cancer cell to Suppressive cell ratio of 1:0.2, and the ratio of Permissive cells changed from 1:0 to 1:0.5. **I-L**: Control simulations of the 3D TME were conducted by varying the Suppressive, Permissive, and Lethal strength. Suppressive, Permissive, or Lethal cells were placed alongside Cervical 2-6Sia 1.0 cells in the 3D TME. (**I**) Suppressive cells, (**J**) Permissive cells, and (**K**) Lethal cells. Simulations involved changing the Suppressive strength (doubling time increase) from 0% to 90% and the Permissive strength (doubling time decrease) from 0% to 90%. For Lethal cells, the simulation varied the chance of inducing cell death from 0% to 44.5%. **L**: Both Suppressive and Permissive cells were placed in the 3D TME, with a 32.5% increase in the doubling time for Suppressive cells, and the ratio of Permissive cell doubling time changed from 0% to 100%. **M-P**: Suppressive, Permissive, or Lethal cells were introduced alongside

Pancreatic 2-6Sia 1.0 cells in the 3D TME. (**M**) Suppressive cells, (**N**) Permissive cells, and (**O**) Lethal cells. **P**: Both Suppressive and Permissive cells were placed in the 3D TME.

ancer case.

**Supplementary Data 1. Cell lineage maps of HeLa cells.**

Cell-lineage maps of HeLa cells were generated using data from the cell-lineage database. In these maps, the following symbols are used to represent specific events: Light blue circle, Mitosis; Pink square, Cell death; Orange circle and orange vertical line, Cell fusion; Black vertical line, Bipolar cell division; and Red vertical line, Multipolar cell division. Levels of SNA1 binding to each cell were shown at the end of the cell lineage map using a heat map scale from blue to red, indicating low to high binding.

**Supplementary Data 2. Cell lineage maps of MiaPaCa2 cells.**

Cell-lineage maps of MiaPaCa2 cells were generated using data from the cell-lineage database. In these maps, the following symbols are used to represent specific events: Light blue circle, Mitosis; Pink square, Cell death; Orange circle and orange vertical line, Cell fusion; Black vertical line, Bipolar cell division; and Red vertical line, Multipolar cell division. Levels of SNA1 binding to each cell were shown at the end of the cell lineage map using a heat map scale from blue to red, indicating low to high binding.

**Supplementary Data 3. Cell lineage maps of Cervical 2-6Sia 1.5 cells.**

Cell-lineage maps of Cervical 2-6Sia 1.5 cells were generated using data from the cell-lineage database. In these maps, the following symbols are used to represent specific events: Light blue circle, Mitosis; Pink square, Cell death; Orange circle and orange vertical line, Cell fusion; Black vertical line, Bipolar cell division; and Red vertical line, Multipolar cell division. Levels of 2-6Sia expression of individual cells were shown using a heat map scale from blue to red, indicating low to high binding.

**Supplementary Data 4. Cell lineage maps of Pancreatic 2-6Sia 1.5 cells.**

Cell-lineage maps of Pancreatic 2-6Sia 1.5 cells were generated using data from the cell-lineage database. In these maps, the following symbols are used to represent specific events: Light blue circle, Mitosis; Pink square, Cell death; Orange circle and orange vertical line, Cell fusion; Black vertical line, Bipolar cell division; and Red vertical line, Multipolar cell division. Levels of 2-6Sia expression of individual cells were shown using a heat map scale from blue to red, indicating low to high binding.

**Supplementary Data 5. Summary of results (Cervical 2-6Sia 1.5, 1.0, 0.5 and 0.25, and Pancreatic 2-6Sia 1.5, 1.0, 0.5 and 0.25).**

The summary table comprises the following information: patient ID, the Best call of immune cell landscape (C1 to C6), Progression-free survival (PFS), cell population size, 2-6Sia levels, the number of reproductive cells generated through multipolar cell division (MD), and the relative composition of Suppressive, Permissive and Lethal cells.
